# Biocatalytic Tetrapeptide Macrocyclization by Cryptic Penicillin-binding Protein-type Thioesterases

**DOI:** 10.1101/2024.11.16.623930

**Authors:** Paisley L. Jeannette, Zachary L. Budimir, Lucas O. Johnson, Elizabeth I. Parkinson

**Affiliations:** James Tarpo Jr. and Margaret Tarpo Department of Chemistry, Purdue University, West Lafayette, IN 47906; Borch Department of Medicinal Chemistry and Molecular Pharmacology, Purdue University, West Lafayette, IN 47906

## Abstract

Cyclic tetrapeptides (CTPs) are a diverse class of natural products with a broad range of biological activities. However, they are challenging to synthesize due to the ring strain associated with their small ring size. While chemical methods have been developed to access CTPs, they generally require the presence of certain amino acids, limiting their substrate scopes. Herein, we report the first bioinformatics-guided discovery of a thioesterase from a cryptic biosynthetic gene cluster for peptide cyclization. Specifically, we hypothesized that predicted penicillin-binding type thioesterases (PBP-TEs) from cryptic nonribosomal peptide synthetase (NRPS) gene clusters containing four adenylation domains would catalyze tetrapeptide cyclization. We found that one of the predicted PBP-TEs, WP516, efficiently cyclizes a wide variety of tetrapeptide substrates. To date, it is only the second stand-alone enzyme capable of cyclizing tetrapeptides, and its substrate scope greatly surpasses that of the only other reported tetrapeptide cyclase Ulm16. AlphaFold modeling, covalent docking, molecular dynamics, and mutational analyses were used to rationalize the broad substrate scope of WP516. Overall, the bioinformatics guided workflow outlined in this paper, and the discovery of WP516, represent promising tools for the biocatalytic production of head-to-tail CTPs, as well as a more general strategy for discovery of enzymes for peptide cyclization.

## Introduction

Peptide-based therapeutics have garnered significant interest in pharmaceutical development, resulting in hundreds of peptides currently undergoing clinical trials and a global market value exceeding tens of billions of dollars.^[1]^ Clinically relevant peptide drugs include the antibiotic murepavadin, the antifungal rezafungin, the immunosuppressant cyclosporine, and the anticancer agent octreotide.^[2]^ Unfortunately, linear and unmodified peptides often display poor therapeutic properties due to their intrinsic instability.^[3]^ Peptidyl natural products exhibit astounding structural diversity, which endows them with a range of biological activities.^[4]^ This diversity is generated by biosynthetic enzymes that construct core scaffolds and perform peripheral modifications, introducing pharmacophores in stereoselective and regioselective fashion. These modifications can include methylation (N, O, C), halogenation, oxidation, and glycosylation.^[5]^ A particularly promising modification is head-to-tail macrocyclization, which enhances membrane permeability by eliminating the zwitterionic C and N termini, while also increasing potency, specificity, and proteolytic stability through defined secondary structures.^[6]^ This enables these peptides to occupy a ‘Goldilocks zone’ between small molecules and biologics.^[7]^ Despite these advantages, macrocyclization remains a synthetically challenging transformation. Epimerization at the α-carbon of the C-terminal residue, competing oligomerization, and conformational rigidity preclude remote residue coupling leading to poor yields and scalability issues.^[8],[9]^ These problems are exacerbated as ring size decreases. Specifically, for tetrapeptides, the ground-state E geometry of the peptide bond impedes the adoption of a conformation favorable to cyclization, while the transannular ring strain further complicates the synthesis and derivatization of pharmaceutically or industrially relevant small cyclic peptides, at times rendering them inaccessible.^[9]^ This severely limits our ability to access a class of cyclic peptides, cyclic tetrapeptides (CTPs), which show improved PK properties and oral bioavailability compared to their larger counterparts.^[10]^ CTP natural products have a variety of interesting bioactivities including inhibition of the chloroplast F1-ATPase (tentoxin), selective blockage of calcium channels by the cyclic peptide family oncychocins, antagonism of kappa opioid receptors by tetrapeptide CJ-15208 and inhibition of histone deacetylases (chlamydocin, trapoxin A, apicidin, microsporins and others), further motivating their investigation.^[11–14]^ While several synthetic strategies have been employed to gain access to CTPs,^[15]^ such as ring contraction via Ser/Thr ligation mediated peptide cyclization^[16]^, on resin cyclization via anchoring of a Glu/Asp^[17]^, pseudo proline protecting groups^[18]^, anion assisted cyclization^[19]^, and direct aminolysis,^[20]^ they are either incompatible with Fmoc solid phase peptide synthesis (SPPS) or restrict amino acid sequence.^[9]^ Additionally, there have been attempts at derivatizing these naturally occurring CTPs to incorporate D amino acids to improve their selectivity, and proteolytic stability.^[21]^ However, even with these turn inducing elements high yielding cyclization still proves to be a challenge with many falling below 50% or resulting in exclusively oligomerization.^[22][23]^

Recently, there has been increased interest in utilizing natural product biosynthetic machinery as biocatalysts for head-to-tail macrocyclization.^[24]^ Many peptidyl natural products from soil dwelling bacteria and fungi are produced by multimodular enzyme complexes known as Non-Ribosomal Peptide Synthetases (NRPSs). NRPSs construct peptides in an assembly line fashion through the use of multiple domains with three being necessary: 1) The adenylation (A) domain, which selects, activates and attaches an amino acid to 2) the carrier protein (T, PCP) domain, and 3) the condensation (C) domain, which catalyzes peptide bond formation and grows the peptide chain.^[25]^ These complexes are frequently implicated in the production of small cyclic peptides including tetra- and pentapeptides.^[26–29]^ Offloading from the complex and cyclization are typically catalyzed by C-terminal domains which are *in cis* to the NRPS, such as thioesterase domains (TE) in bacteria or condensation termination domains (C_T_) in fungi (**Figure 1A**).^[30]^ While these domains can efficiently catalyze the cyclization of cyclic peptides *in vivo* and have been studied as biocatalysts^[31]^, they have failed to reach widespread use due to their low substrate promiscuity,^[32]^ an inability to accurately predict cyclization modes (e.g. head-to-tail) via bioinformatics,^[33]^ and in the case of C_T_ domains, the strict requirement for a peptidyl carrier protein (PCP) tethered substrate.^[30],[34]^ However, a recently discovered thioesterase family sharing sequence similarity to the penicillin binding protein (PBP-TEs) have garnered significant attention for their use as biocatalysts for head-to-tail peptide macrocyclization as they act *in trans* to the NRPS and display unprecedented substrate promiscuity.^[35]^ The most extensively studied enzyme from this family is SurE from *Streptomyces albidoflavus*,^[36]^ which natively catalyzes the cyclization of structurally unrelated octa- and decapeptides. SurE has demonstrated great biocatalytic promise *in vitro*, catalyzing the cyclization of peptides ranging from 5-10 amino acids varying widely in sequence and peptide mimics with the equivalent length of 23 amino acids.^[37–41]^ Recently, we characterized Ulm16 from *Streptomyces* sp. KCB13F003, which natively catalyzes the cyclization of the hexapeptide antibiotics the ulleungmycins (**Figure 1B**).^[42]^ While Ulm16 has a narrower substrate scope with respect to peptide length compared to SurE, it has demonstrated the unique ability to cyclize tetrapeptides *in vitro* with many substrates having a cycloselectivity above 20 (*i*.*e*. a cyclic:hydrolyzed ratio above 20:1) and high catalytic efficiencies up to 10^6^ M^-1^s^-1^.^[43]^ However, its substrate scope was limited to the cyclization of DDLL and DDDL tetrapeptides (here, and for the remainder of this article, peptides will be described from C to N-terminus). Additionally, it had a relatively limited scope for amino acids at the third position. These limitations drove us to find an alternative biocatalyst for cyclization of tetrapeptides. Herein, we describe the bioinformatics identification of WP_043619516.1 (herein named WP516) a PBP-TE from a cryptic biosynthetic gene cluster (**Figure 1C**), which is a capable of cyclizing a much wider scope of diastereomers, including DLDL and DLLL peptides, and has a greatly extended amino acid substrate scope compared to that of Ulm16.

**Figure 1.**
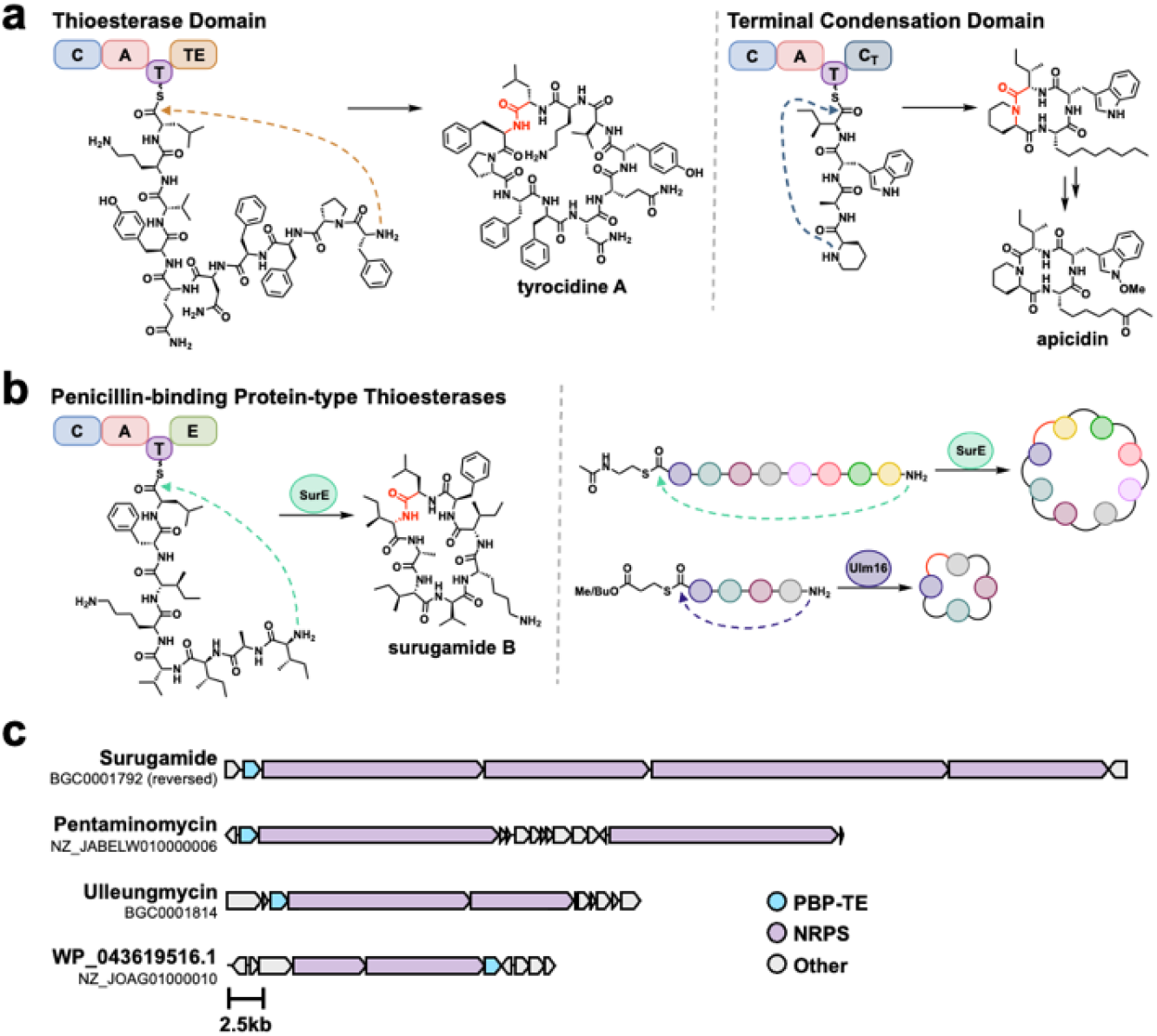
Head-to-tail offloading mechanisms in non-ribosomal peptide synthesis. **A.)** Thioesterase domain (TE) catalyzed cyclization and release^25^ and condensation Termination domain (CT) catalyzed cyclization and release from fungal NRPSs are common in peptide cyclization.^20^ Both act *in-cis* (connected) to the NRPS. **B.)** Penicillin-binding protein type thioesterase (PBP-TE) catalyze peptide cyclization and release *in-trans* (not connected) to the NRPS, with SurE and Ulm16 showing broad catalytic activity.^32,36^ **C.)** Biosynthetic gene clusters (BGCs) of known natural products produced by PBP-TEs along with the BGC of WP_04319516.1 a cryptic tetrapeptide BGC with a PBP-TE.

### Identification and Initial Evaluation of PBP-TEs WP516 and SEC28031.1

To-date, no known cyclic tetrapeptide natural products have been associated with a PBP-TE; however, previous bioinformatics has identified multiple PBP-TEs colocalized with cryptic NRPS biosynthetic gene clusters (BGCs) that are predicted to produce a tetrapeptide (*i*.*e*. have 4 adenylation, domains).^[44]^ While expression of traditional bioinformatically identified TEs is inherently risky due to their low substrate promiscuity and our inability to predict what reaction they will catalyze (hydrolysis, head-to-tail, or head-to-sidechain cyclization),^[33]^ PBP-TEs have displayed large substrate tolerance and have only been shown to catalyze head-to-tail cyclization. Because of these characteristics, we chose to explore cryptic PBP-TEs as potential biocatalysts for tetrapeptide macrocyclization. We conducted a bioinformatics search utilizing BiG-SCAPE CORASON^[45]^ to identify PBP-TEs that likely cyclize tetrapeptides (see **Figure S1** for workflow and **Figure S2** for results). From this, we identified 10 predicted tetrapeptide PBP-TEs. Two PBP-TEs were chosen for expression due to their predicted substrates: WP_031183424.1 from *Streptomyces seoulensis* and WP_043619516.1 from *Nonomuraea candida* HMC10 (**SI Figures S2-S3**). WP_031183424.1 was hypothesized to cyclize a tetrapeptide with DLDL stereochemistry and clustered next to FlkO, a recently validated PBP-TE with a native hexapeptide substrate possessing DLDLDL stereochemistry.^[46]^ Unfortunately, attempts to express WP_031183424.1 were unsuccessful, preventing further investigation or characterization (**Figure S4**).

We were able to express and characterize WP_043619516.1 (**Figure S4**, here after referred to as WP516) which was predicted to natively cyclize a tetrapeptide with a C-terminal D-serine followed by D-enduracididine^[47]^ at the 2-position, L-phenylalanine at the 3-position, and an N-terminal L-valine. Additionally, we conducted a BlastP search of WP516 and identified SEC28301.1 (from *Streptomyces* sp. TLI_105) which, despite having the same predicted substrate as WP516, shared only 54% sequence identity (**Figure S5**). We hypothesized that this may cause them to possess differing substrate scopes from each other. In our initial evaluation of WP516 and SEC28301.1, we set out to confirm their activities as tetrapeptide cyclases and investigate their tolerance of DLDL stereochemistry, a characteristic that resulted in reduced cyclization activity with Ulm16. Utilizing a substrate previously tested with Ulm16 (**1**), both WP516 and SEC28301.1 demonstrated the ability to cyclize a tetrapeptide possessing DLDL stereochemistry (**Figure 2A, Figure S7**). Due to SEC28301.1 having poorer solubility and a narrower substrate scope with respect to some C-terminal residues (**Figure S8**), we chose to focus on WP516.

**Figure 2.**
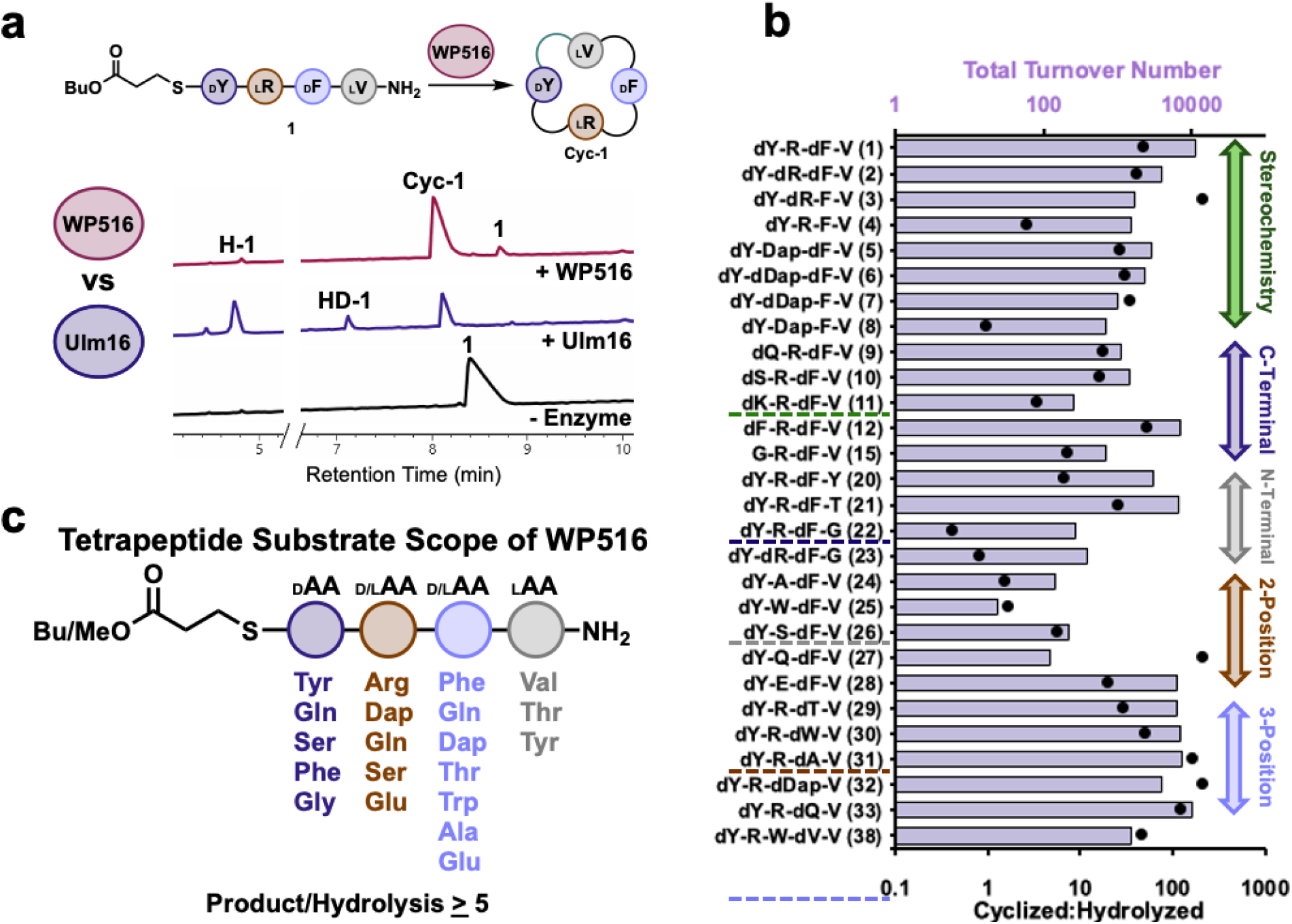
Substrate scope of WP516. **A.)** comparison between WP516 and Ulm16 on DLDL peptide substrate **1**. Reaction conditions can be found in the methods section and **SI Table 1. Cyc-1** is head-to-tail cyclic peptide of **1, H-1** represents hydrolyzed **1** and **HD-1** represents **1** which has been dimerized and hydrolyzed. Traces were taken at a wavelength of 214 nm. Full UPLC traces can be found in **Figure S7. B.)** Substrate scope of WP516 featuring total turnover number (TTN, top, bars) and cyclized to hydrolyzed ratio (bottom, dots) for all substrates that were cyclized. For a full list of tested substrates see **SI Table 1**. All assays were carried out in triplicate. **C.)** Representative substrate scope of WP516 featuring all amino acids and stereochemistries that yielded a cyclized:hydrolyzed ratio >5:1.

### Substrate Scope Investigation

To fully explore the stereochemical tolerance of WP516, we synthesized diastereomers of **1** with DDDL (**2**), DDLL (**3**), or DLLL (**4**) stereochemistry. Excitingly, a high cycloselectivity was maintained with these diastereomers. This differs greatly from Ulm16, where changing the stereochemistry to DLDL or DLLL greatly reduced or completely eliminated cyclization activity, respectively. Interestingly, we found the DLDL peptide (**1**) to be the best WP516 substrate with a TTN of 11219 and cycloselectivity >20 (**SI Table 1, Figure 2B, Figure S9**). This is despite the fact that the natural substrate for WP516 is predicted to have DDLL stereochemistry (**3**). Modifying the stereochemistry from DLDL to DDDL (**2**) or the predicted DDLL (**3**) resulted in modest decreases in TTNs compared to the DLDL substrate but good retention of cycloselectivity (>20, **Figure S11** and **S12**). With the DLLL diastereomer (**4**), WP516 displayed a greater propensity for hydrolysis with cycloselectivity dropping to just 2.6 and an approximately eight-fold reduction in TTN compared to **1** (**Figure S13**). However, it is a great improvement over Ulm16, which is unable to cyclize DLLL peptides (**Figure 3A, Figure S14**).

**Figure 3.**
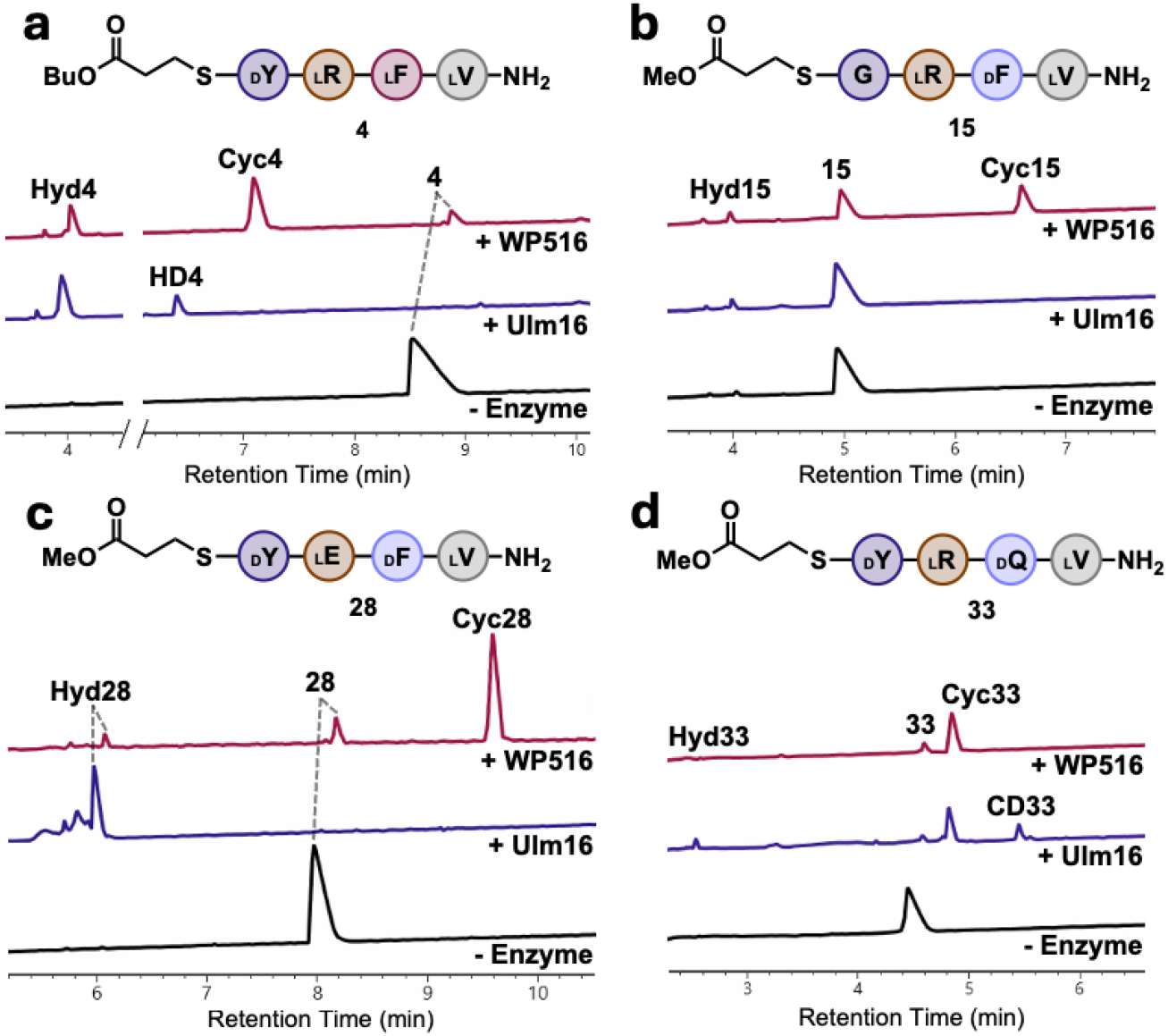
Comparison of WP516 and Ulm16 enzyme reactions with tetrapeptide substrates. **A.)** Ulm16 and WP516 with peptide **4. B.)** Ulm16 and WP516 with peptide **15. C.)** Ulm16 and WP516 with peptide **28. D.)** Ulm16 and WP516 with peptide **33**. UV traces (214 nm) are from UPLC analysis. Full list of reaction conditions and traces can be found in the supplementary information. All reactions were run in triplicate with a representative 214 nm trace used for each reaction. CycX is head-to-tail cyclized peptide, HydX is hydrolyzed peptide, and CDx represents a peptide that has been dimerized then cyclized.

To further examine the ability of WP516 to cyclize substrates poorly tolerated by Ulm16, we investigated its activity toward a substrate with a 2,3-diaminopropionic acid (dap) at the 2 position and DLDL stereochemistry (**5**). Excitingly, **5** was well tolerated by WP516 (TTN: 2901, cycloselectivity >20) and exhibited a catalytic efficiency of 3.7 × 10^4^ M^-1^s^-1^ (**Figure 4A, Figure S15, SI Table 1**). In the same manner as before, we tested diastereomers of **5** with DDDL (**6**), DDLL (**7**), and DLLL (**9**) stereochemistries (**SI Table 1**). Similarly to diastereomers of **1**, the DDDL (**6**) and DDLL (**7**) diastereomers were well tolerated, with modest decreases in activity as observed in the Michaelis-Menten kinetics (**Figure 4, Figure S16** and **S17**) but excellent cycloselectivities (>20). The DLLL substrate (**8**) was still cyclized by WP516, but its cycloselectivity was significantly diminished, dropping to 0.9 (**Figure S18**).

**Figure 4.**
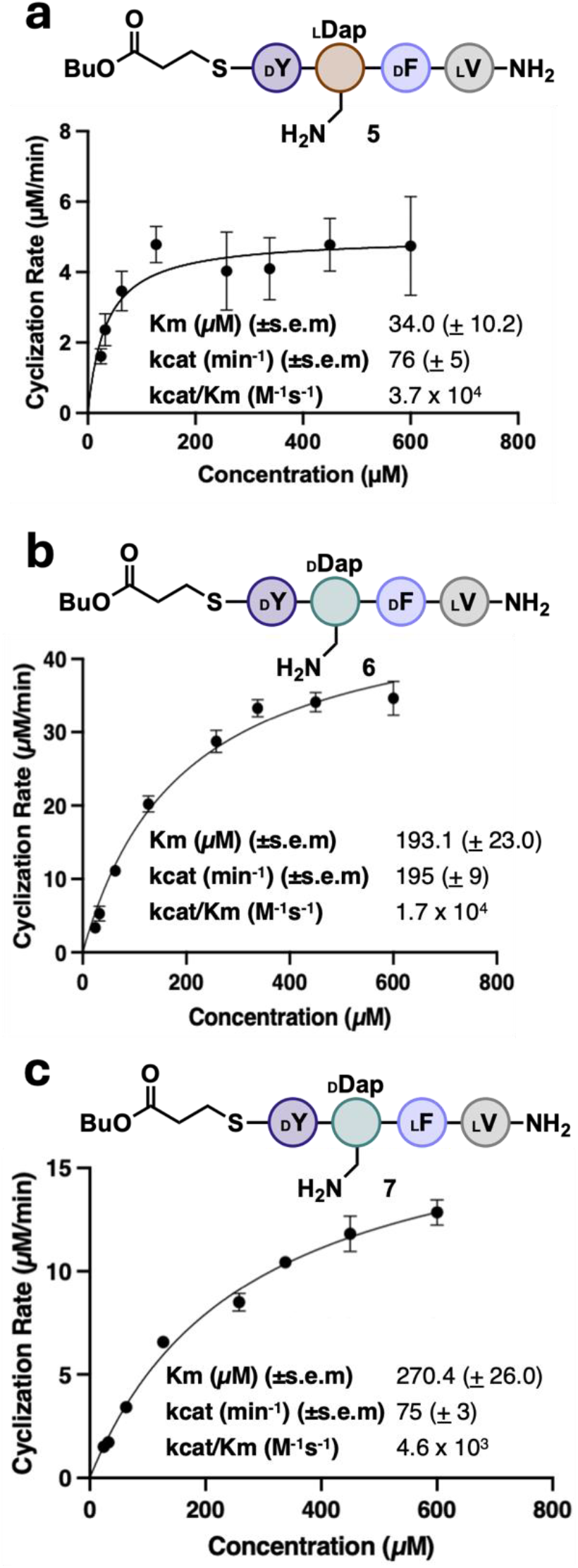
Michalis-Menten kinetics for WP516. **A.)** peptide **5 B.)** peptide **6 C.)** peptide **7**. Methods can be found in the supplementary information All reactions were run in triplicate. s.e.m., standard error of the mean.

To fully elucidate the substrate scope of WP516, we used **1** as our model sequence, modifying each position and determining TTNs. WP516 was tolerant of a wide range of substitutions of the C-terminal residue as long as D-chirality was maintained (peptides **9-15, Figure 2B, SI Table 1**). Polar uncharged residues at the C-terminus, such as D-glutamine (**9**) or D-serine (**10**), decreased catalytic activity compared to D-tyrosine but maintained cycloselectivities above 15. We do not attribute this decrease in activity to nonenzymatic factors such as glutarimide formation or epimerization due to their absence in UPLC spectra in enzymatic reactions. A positive charge at the C-terminus, such as D-lysine (**11**), was not well tolerated by WP516, resulting in a dramatically reduced TTN and greater propensity for hydrolysis (**Figure S21**).

Substitution of D-tyrosine for D-phenylalanine (**12**) was well tolerated by WP516 with good catalytic efficiency (*k*_cat_/*K*_M_ = 2.7 × 10^4^ M^-1^s^-1^) and little change in the cycloselectivity (**Figure S22** and **S23**). Substitution for D-valine (**13**) resulted in a substrate that was not processed by WP516 (**Figure S24**), possibly due to the increase in steric bulk at the β-carbon. Negatively charged D-glutamate (**14**) at the C-terminus also resulted in a substrate that WP516 could not process (**Figure S25**), suggesting that the negatively charged amino acid may prevent the initial attack of the active site serine on the thioester. Notably, WP516 was tolerant of glycine (**15**) at the C-terminus, with a cycloselectivity of 7 (**Figure 3B, Figure S26**). This is particularly noteworthy because many PBP-TEs are unable to tolerate a C-terminal glycine without further engineering.^[39]^ Interestingly, Ulm16 failed to cyclize a tetrapeptide with a C-terminal glycine residue (**Figure 3B, Figure S27**) despite previously showing the ability to cyclize a hexapeptide substrate bearing a C-terminal glycine, making WP516 the only tetrapeptide cyclase capable of tolerating a C-terminal glycine.^[43]^ On the basis of this finding, we chose to probe the ability of WP516 to cyclize an all L tetrapeptide with glycine at the C terminus (GLLL, **16**), but only hydrolysis was observed (**Figure S28**).

Tolerance for substitutions at the N-terminus (peptides **17-24, Figure 2B, SI Table 1**) was more limited, with WP516 displaying a preference for small, uncharged residues. Substituting L-valine for L-proline (**17**), which converted the N-terminus to a secondary amine, resulted exclusively in hydrolysis by WP516 (**Figure S29**). Similarly, a charged residue at the N-terminus, either L-histidine (**18**) or L-glutamate (**19**), yielded only the hydrolyzed product (**SI Figures 30**-**31**). Substitution for L-tyrosine (**20**) was better tolerated, resulting in an approximately three-fold reduction in TTN and a moderate cycloselectivity of 6 (**Figure S32**). An N-terminal L-threonine (**21**) resulted in an approximately two-fold reduction in TTN compared to L-valine, although no significant decrease in cycloselectivity was observed compared to **1** (>20) (**Figure S33**). Notably, WP516 exhibited cyclization activity toward substrates possessing an N-terminal glycine, with either D-D-D-G (**22**) or D-L-D-G (**23**) stereochemistry. However, this resulted in a 30-fold and 40-fold reduction in TTN, respectively, compared to L-valine and preference for hydrolysis over cyclization (**SI Figures 34** and **35**). Regardless, cyclization of **22** is the first example of a homochiral peptide being cyclized by this class of enzymes demonstrating the utility of WP516 and how it can be used as a starting point for future engineering approaches.

At the 2 position (Peptides **24**-**28, Figure 2B, SI Table 1**), WP516 displays a preference for charged residues. Replacement of L-arginine with non-polar residues, such as L-alanine (**24**) or L-tryptophan (**25**), substantially increased hydrolysis with cycloselectivity almost approaching 1, and drastically decreased the ability of WP516 to turnover (**SI Figures 36** and **37**). Polar uncharged amino acids similarly reduced catalytic activity but maintained better cycloselectivities, with L-serine (**26**) having a cycloselectivity >5 (**Figure S38**) and for L-glutamine (**27**) complete selectivity for cyclized product was observed (**Figure S39**). Substitution for L-glutamate (**28**) at the 2 position was much better tolerated by WP516, with a cycloselectivity of 20 and a good retention of catalytic activity (**Figure S40**). In contrast, incubation of this substrate with Ulm16 resulted exclusively in hydrolysis (**Figure 3C, Figure S41**).

Substitutions made at the 3 position were very well tolerated by WP516 with a wide variety of residues accepted (Peptides **29**-**34**). Replacement of D-phenylalanine with D-threonine (**29**), D-tryptophan (**30**), D-alanine (**31**), or D-diaminopropionic acid (D-dap, **32**) yielded substrates that retained moderate to high TTNs with low levels of hydrolysis (**SI Figures 42**-**45**). D-glutamine (**33**) was specifically well tolerated with little change in TTN compared to D-phenylalanine and virtually complete selectivity for the cyclic product (**Figure S46**). Again, these results strongly contrast with Ulm16 which, when incubated with this substrate, produced cyclic dimer and increased levels of the hydrolyzed product alongside the cyclic peptide (**Figure 3D, Figure S47**). Similarly to what was observed at the 2 position, WP516 was also tolerant of a negatively charged D-glutamate (**34**) at the 3 position with minimal hydrolysis. Unfortunately, the cyclic peptide was too insoluble to determine TTN, though we still observed a cycloselectivity above 20 (**Figure S48)**.

Prompted by the previously observed ring size promiscuity of PBP-TEs, we explored the tolerance of WP516 to substrates ranging from 3 to 6 amino acids. Specifically, we investigated its ability to cyclize 3 tripeptide substrates with varying stereochemistry (**35**-**37**) and three larger substrates: a modified PenA pentapeptide substrate that is efficiently cyclized by Ulm16 (**38**), a derivative of the Ulm16 hexapeptide substrate (**39**), and a modified DsaJ hexapeptide substrate (**40**). While the tri- and hexapeptide substrates resulted exclusively in hydrolysis by WP516 (**SI Figures 49**-**53**), the PenA pentapeptide substrate was successfully cyclized (**Figure S54**), with a cycloselectivity >20. While WP516 was notably less efficient with this substrate when compared to its activity toward tetrapeptide substrates (>10-fold reduction in TTN and 10^2^-fold reduction in catalytic efficiency, **Figure S55**), the minimal levels of hydrolysis highlight tolerance for pentapeptides in addition to tetrapeptides.

To test the utility of WP516 on a larger scale, we assayed WP516 on a 44 milligram scale with peptide **1**. Given the high degree of purity (>90%) of these peptide thioesters post thiolysis and deprotection (**SI Figure 56**, T=0) we were able to directly cyclize the crude thioesters, limiting the process to 1 round of HPLC purification after the enzymatic reaction. While full conversion was observed, the final yield of cyclic peptide was 50%, likely due to loss of product during purification (**SI Figures S56-S58**). To the best of our knowledge, this is the first demonstration of any PBP-TE being used on an isolable scale for cyclic peptide production. In all, these results highlight the utility of WP516 as a biocatalyst for cyclic peptide production.

### Rationalization of WP516’s Substrate Tolerance: Computational Modeling

The distinct tetrapeptide substrate scope of WP516 compared to Ulm16 prompted us to investigate the basis of this unique substrate tolerance. Structurally, PBP-TEs have been shown to display two domains: a PBP domain containing an α-β-hydrolase fold and a lipocalin-like domain containing an eight-strand antiparallel β-barrel fold, connected by an unstructured loop (**Figure 5**). Previously, the crystal structure of Ulm16 (PDB: 8FEK) revealed that, while the PBP domains of Ulm16 and SurE (PDB: 6KSU and 6KSV) are highly similar, the orientation of the lipocalin domains differed between the two enzymes, suggesting that the lipocalin domain may play an important role in the enzymes ability to cyclize shorter peptides.^[43]^ Docking and subsequent mutagenesis studies with Ulm16 previously demonstrated that the lipocalin domain does play a key role in cyclization, with residue Arg431(Arg438 of WP516) on the lipocalin domain being essential to cyclization.^[43]^ To investigate if a similar trend of lipocalin domain differences holds true for WP516, we attempted to solve its crystal structure. Unfortunately, crystallization efforts were unsuccessful due to insufficient protein solubility, prompting us to instead generate and compare its AlphaFold3^[48]^ model to the crystal structures of SurE and Ulm16. Consistent with previous structural comparisons among this family of proteins, the α/β-hydrolase domains of Ulm16, SurE and WP516 are highly similar (**Figure 5, Figure S59**), sharing approximately 47% sequence identity and a root mean square deviation (RMSD) of approximately 0.950 Å with each other (**Figure S60**). Interestingly, in the 70 amino acids (N to C terminus) leading into the unstructured loop between the two domains, the proteins exhibit close to 60% sequence identity, followed by a conserved XDVG motif (**Figure S61A** and **C**). However, after this motif, the proteins lose nearly all sequence identity, despite all ending in a lipocalin domain (**Figure S61 B** and **D**). Of note, WP516 and Ulm16 are predicted to have similar lipocalin domain angles yet only share 25% sequence identity. Looking at the lipocalin domain density between the crystal structure of Ulm16 and the Alpha Fold 3 model of WP516 revealed that WP516 has increased steric bulk from the addition of multiple tryptophan residues (W374, W394), resulting in a much shallower lipocalin domain, which possibly forces shorter peptides to turn sooner (**Figure S62**).

**Figure 5.**
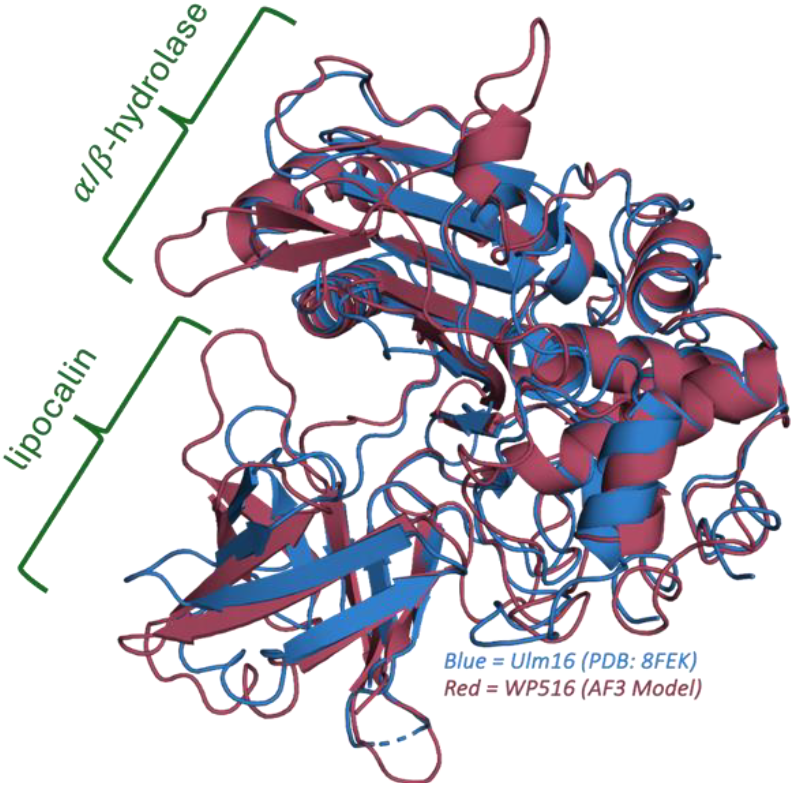
WP516 Alpha Fold 3 model comparison with Ulm16 crystal structure (PDB: 8FEK) highlighting the similarity between there α/β-hydrolase domain and lipocalin domain angle. RMSD 1.023

To further investigate the substrate tolerance and cyclization mechanism of WP516, we conducted covalent docking studies with its AlphaFold model. We began by docking tetrapeptide **5** to gain insight into unique ability of WP516 to cyclize peptides with a 2-position Dap and DLDL stereochemistry. Consistent with previous observations that linked variability in the docking outputs with hydrolysis,^[43]^ the top outputs when docking **5** with WP516 were highly uniform, with many taking a cyclic shape, while docking **5** with Ulm16 led to a peptide lacking uniformity and not taking a cyclic shape (**Figure S63**). Alternatively, the hexapeptide **40** resulted in a cyclic shape when docked with Ulm16 but not with WP516 (**Figure S64-S65**). These differences were hypothesized to be due to differences in the lipocalin domain.

We next conducted molecular dynamics (MD) simulations to better understand the protein– substrate interactions. We performed simulations on the Ulm16 crystal structure and the WP516 AlphaFold model, each covalently docked with peptide **1**. For each model, we ran four independent 100 nanosecond (ns) simulation replicates to ensure comprehensive conformational sampling. The GROMACS MD simulation suite was used to compute the RMSD of the ligand and protein backbone, as well as the RMS fluctuation of the backbone per residue for each simulation to ensure system stability (**SI Figures S66-S68**). Generally, the WP516 simulations had lower backbone RMSD values than those of Ulm16; however, no notable differences in these stability parameters were observed. To further explore potential differences that could explain their divergent substrate scopes, we measured the distance between the nucleophile and electrophile involved in the macrocyclization reaction (the nitrogen of the N-terminus and the carbonyl carbon of the C-terminus), as well as the angle between the N, C, and carbonyl O atoms throughout the simulations (**SI Figures S69-S70**). We hypothesized that a shorter distance and an angle closer to 180° would reflect a productive macrocyclization geometry, and that Ulm16, which poorly tolerates peptide **1**, would be an outlier. Indeed, the Ulm16 simulations with peptide **1** exhibited larger distances between these atoms compared to WP516. By comparison, simulations of both Ulm16 and WP516 with substrate **3**, a peptide cyclized efficiently by both enzymes, showed similarly large interatomic distances and angles between the reactive termini (**Figures S66-S69**).

In both enzymes, the peptide gradually shifted from initial contacts with the lipocalin domain toward β-strands 6 and 7 of the α/β-hydrolase domains, causing it to be challenging to determine whether the interactions with lipocalin domain or the α/β-hydrolase domains were key for cyclization. (**Figure 6, Figure S71**). Interestingly, the peptides adopted a similar trajectory regardless of the protein they were simulated with, ultimately localizing near β-strands 6 and 7, further implicating this region of the α/β-hydrolase domain in substrate positioning (**Figures S72**). Examination of specific protein–substrate interactions and ligand conformations over the course of the simulations revealed differences between

**Figure 6.**
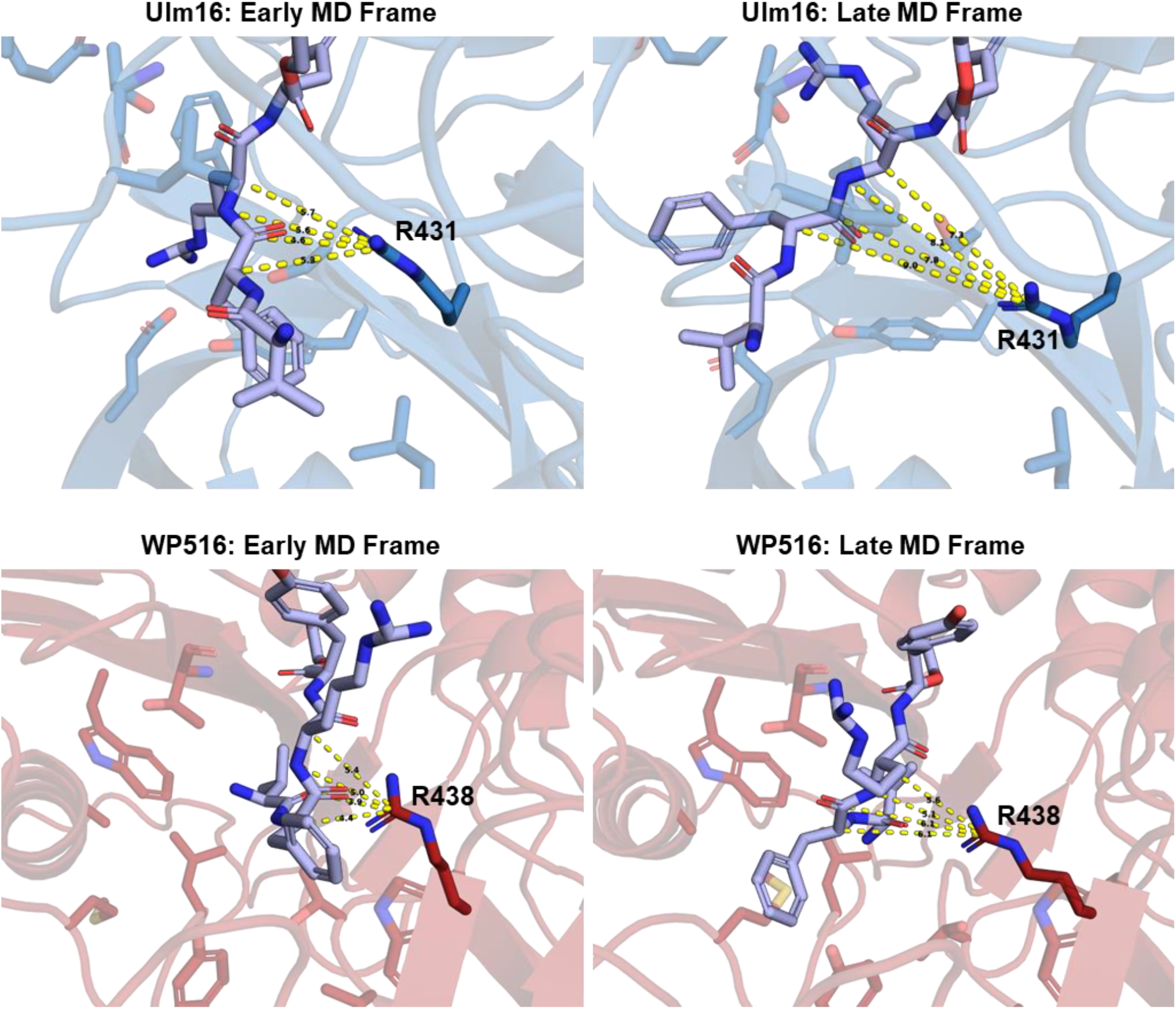
key frames from MD simulations with DLDL peptide 1 docked to Ulm16 (top, blue) and WP516 (bottom, red). Simulation closest to the average of the 4 were chosen with early frame images taken to give a representative view of how the substrate looked post equilibration, and late view is to give a representative view of how the substrate looked towards the end of the MD simulation. Residues in the lipocalin previously implicated in the docking (see **Figure S64**), are shown in transparent, excluding R431/R438 due to its known importance in cyclization. New residues in the α/β-hydrolase domain are shown in stick.

Ulm16 and WP516 that may underlie their distinct substrate specificities. In simulations with Ulm16, peptide **1** was generally unable to adopt or maintain a cyclization-competent conformation. In contrast, WP516 positioned the substrate into a cyclic geometry early in the simulation and retained this conformation throughout most of the trajectory. Compared to Ulm16, the substrate in the WP516 simulations was within hydrogen-bonding distance to a larger number of residues, including the key lipocalin arginine (R438 in WP516), potentially contributing to increased stabilization within the active site. The enhanced positioning observed in WP516 may also be facilitated by reduced steric bulk at position 296, where a glycine in the α/β-hydrolase domain (G296) replaces a leucine present in Ulm16 (L300). This substitution appears to permit an alternative binding orientation, allowing the backbone amide N–H and side chain of T295 on β-strand 6 to form hydrogen bonds with the C-terminal carbonyl oxygen and N-terminal amine of the substrate, respectively. These interactions effectively “lock” the substrate in a cyclic conformation, consistent with WP516’s high cyclization efficiency for peptide **1**. Interestingly, we have previously found that a mutant Ulm16 L300G resulted in approximately 3-fold improved cyclization of DLDL peptides compared to wild type Ulm16, providing experimental support to this finding.^[43]^

### Mutant Testing

Given the computational modeling results suggested the potential importance of both the lipocalin domain and the α/β-hydrolase domain for substrate specificity, we sought to experimentally evaluate their roles. To determine the effect of the lipocalin domain, we generated chimeric proteins with swapped lipocalin domains. We constructed a chimeric variant of Ulm16 (Ulm16 Chimera, Ulm16C) containing its native α/β-hydrolase domain fused to the lipocalin domain and active site loop of WP516 (**Figure S73**). This construct also incorporated the previously mentioned L300G mutation that enhances cyclization of DLDL peptides. Conversely, a WP516 chimeric protein (WP516 Chimera, WP516C) was generated by combining the WP516 α/β-hydrolase domain with the lipocalin domain of Ulm16 (**Figure S74-S75**). We hypothesized that these swaps would invert substrate specificity. To test this, we evaluated both chimeras against substrates with known divergent outcomes. First, two hexapeptides (**39** and **40**), which are efficiently cyclized by Ulm16 but are not tolerated by WP516, were tested. Contrary to our expectations, both chimeras exhibited substrate profiles identical to their respective wild-type proteins (**Figures S76** and **S77**). Testing these chimeras against tetrapeptides further supported this trend. Specifically, we evaluated their ability to cyclize peptide **1**, a substrate that WP516 cyclizes cleanly without hydrolysis, and peptide **4**, which is cyclized by WP516 but undergoes dimerization with Ulm16.

Once again, the chimeras mirrored the behavior of their respective parental enzymes (**Figure 7, Figures S78** and **S79**). These results suggest that, while the lipocalin domain is essential for cyclization,^[43]^ lipocalin domain swapping is insufficient to alter substrate specificity between Ulm16 and WP516.

**Figure 7.**
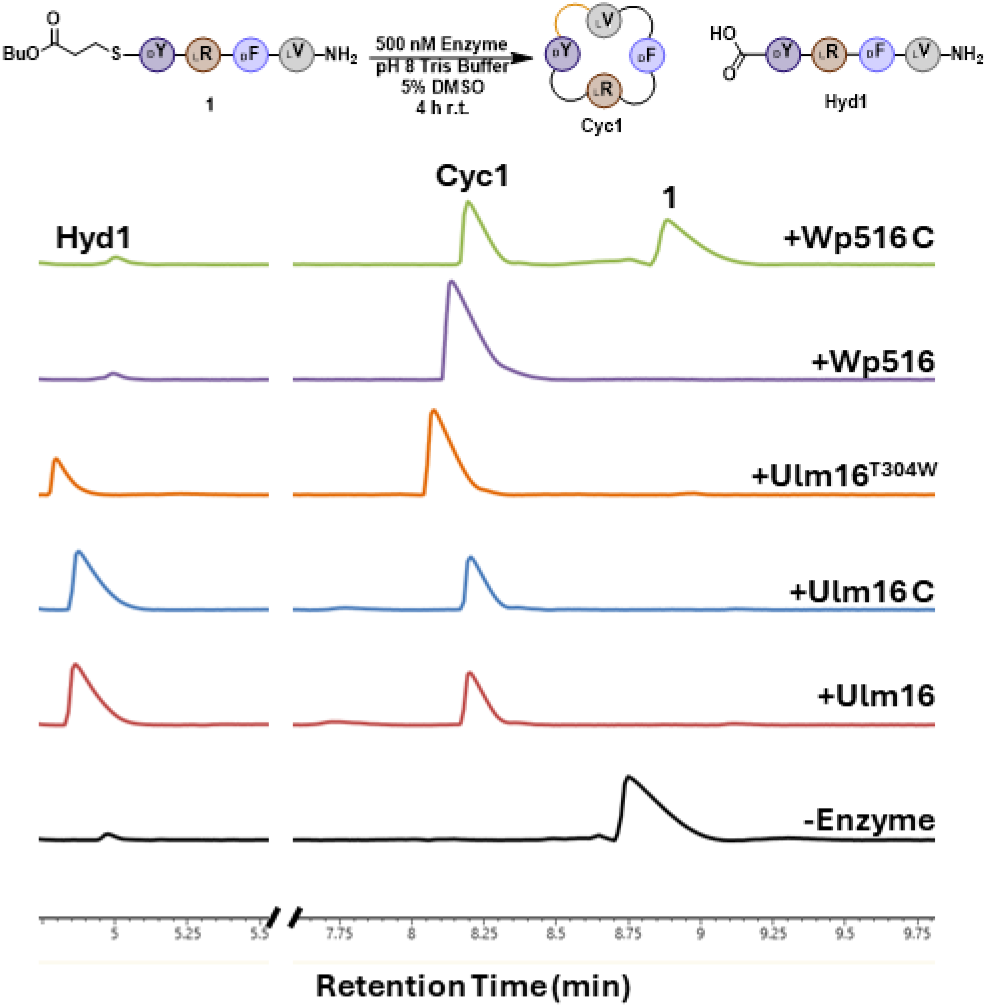
Comparison between WT and mutant enzyme reactions with DLDL peptide substrate 1. Reaction conditions can be found in the methods section. **Cyc1** is head-to-tail cyclic peptide of **1** and **Hyd1** represents hydrolyzed **1**. Traces were taken at a wavelength of 214 nm. Full UPLC traces can be found in **Figure S78**.

Given the lipocalin chimera results, this suggests that the α/β-hydrolase domain likely plays a dominant role in determining substrate length and stereochemical preference. We examined all sequence differences within the α/β-hydrolase domains, excluding conservative mutations such as Ile to Leu. This analysis revealed 107 amino acid differences. Structural analysis identified only one residue, Thr304 in Ulm16 and Trp300 in WP516, that was positioned in proximity to the substrate (**Figure S80**). In addition, the residue located directly behind this position in the structure was also substituted, with Gln327 in Ulm16 and Ala323 in WP516. This change likely accommodates the bulkier Trp side chain in WP516 (**Figure S80**). To evaluate the functional significance of the identified tryptophan in WP516, we generated two point mutants: Ulm16^T304W^ and WP516^W300T^. Although we were unable to obtain soluble protein for the WP516^W300T^ variant, the Ulm16^T304W^ mutant was successfully expressed, purified and tested with the same substrates used to evaluate the chimeric proteins. Remarkably, this single-residue substitution greatly enhanced cyclization efficiency for the tetrapeptides. Cycloselectivity of the DLDL peptide **1** increased approximately five-fold (**Figure 7, Figure S81**), and, for the first time, Ulm16 was able to successfully cyclize the DLLL peptide **4** (**Figure S82**). Interestingly, these improvements occurred without significant loss of cyclization of the native hexapeptides **39** and **40** (**Figures S83** and **S84**). We also note that this enhancement could potentially be further improved by introducing an additional L300G substitution, which previously led to a ∼3-fold increase in peptide **1** cyclization.^[43]^ Overall, these mutational studies suggest that the α/β-hydrolase domain plays a central role in determining both cyclization efficiency and substrate length specificity.

## Discussion

The chemical synthesis of cyclic tetrapeptides (CTPs) is a persistent challenge due to their high levels of ring strain. Even peptides with turn inducing elements such as proline, and multiple D-amino acids remain a synthetic challenge.^[21–23,49]^ Current methods that alleviate the problems with accessing CTPs often severely limit the peptide sequence and/or are incompatible with Fmoc-based solid-phase peptide synthesis (Fmoc-SPPS), thereby restricting access to a class of highly valuable scaffolds with diverse biological activities of interest across various industries. To address this, we explored a biocatalytic method to improve their accessibility. While excised thioesterases and ribosomally synthesized post-translationally modified peptide (RiPP) cyclases are efficient at cyclizing large peptides^[43]^ and even proteins^[50–52]^, none have demonstrated the ability to efficiently produce CTPs. We sought to expand upon the previous discovery of Ulm16, the first PBP-TE capable of efficiently producing CTPs, by conducting a bioinformatics search to identify a cyclase with a broader substrate scope. However, up to this point, the literature has lacked precedence on a workflow for the discovery, expression, and validation of TE domains belonging to cryptic BGCs as biocatalysts, despite this approach yielding many novel and synthetically useful enzymes of other classes.^[5,53]^ This is due, in part, to the poor substrate promiscuity of TE domains and lack of knowledge about the reaction they catalyze based on primary amino acid sequence.^[33]^ To circumvent these issues we aimed to exploit the broad substrate promiscuity, ease of bioinformatics identification, clustering based on peptide length, and native selectivity for head-to-tail macrocyclization of PBP-TEs towards the development of a workflow capable of identifying head-to-tail tetrapeptide cyclases.^[35,44]^ In doing so, we have successfully expanded the biocatalytic toolbox of PBP-TEs by demonstrating the utility of cryptic PBP-TEs, highlighted by the discovery of WP_043619516.1 (referred to as ‘WP516’). To the best of our knowledge, WP516 represents the first instance of a PBP-TE discovered entirely through bioinformatics, and one that is not associated with a known natural product. The substrate scope of WP516 surpassed all previously reported PBP-TEs with respect to tetrapeptides, favoring cyclization over hydrolysis for all stereochemistries, as long as heterochirality is maintained at the C- and N-termini. Interestingly, WP516 is one of the few PBP-TEs capable of utilizing a C-terminal glycine, accepting a GLDL substrate. The only other examples are Ulm16 and WolJ, both of which catalyze the cyclization of the hexapeptide desotamides—a substrate WP516 does not accept. Additionally, WP516 tolerates a wide range of amino acid substitutions at the 2nd and 3rd positions, beyond what Ulm16 has previously been shown to accommodate. This substrate promiscuity was leveraged to access a DLDL, cyclic peptide that Ulm16 was incapable of producing, on preparative scale further showcasing its utility beyond previous characterized tetrapeptide cyclases.

To rationalize the broad tetrapeptide substrate scope of WP516, we performed computational modeling with two substrates that displayed varying degrees of cyclization efficiency. Despite the lipocalin domain angle being highly similar between Ulm16 and WP516, we observed multiple residue substitutions in WP516 that increase steric bulk. These observations, in conjunction with observations from covalent docking studies, led us to hypothesize that the increased bulk of the lipocalin domain may contribute to the increased tetrapeptide substrate promiscuity of WP516. To experimentally evaluate the contribution of the lipocalin domain to the differing substrate scopes of these enzymes, we generated chimeric mutants, Ulm16 Chimera and WP516 Chimera, that featured swapped lipocalin domains and tested the activity of these proteins with substrates with known divergent outcomes between the wild-type enzymes. Surprisingly, we found the chimeras to possess substrate specificity identical to their respective wild-type proteins. This suggests that while the lipocalin domain is required for peptide cyclizaiton, it is not the central factor dictating substrate selectivity of these enzymes. Instead, the α/β-hydrolase domain plays a much larger role in this than previously thought. To investigate beyond the static snapshot that covalent docking provides, we conducted MD simulations on the protein-substrate complexes in which we observed a trend of the peptide substrates shifting away from the lipocalin domain toward the α/β-hydrolase domain. Examination of sequence differences between the Ulm16 Chimera and WP516 within their α/β-hydrolase domains led to the identification of a residue near the substrate binding site, Thr304 in Ulm16 and Trp300 in WP516. Given that this was the only notable difference in proximity to the substrate between the α/β-hydrolase domains of these proteins, we generated and tested the activity of point mutant Ulm16^T304W^. Excitingly Ulm16^T304W^ displayed an increased tolerance towards a DLDL tetrapeptide substrate and, notably, enabled Ulm16 to cyclize a tetrapeptide with DLLL stereochemistry for the first time. This highlights the fundamental importance of this residue for tetrapeptide cyclization. Taken together, these results illuminate the importance of the α/β-hydrolase domain in determining substrate preference of PBP-TEs and provide a valuable foundation to guide future engineering efforts toward the development of novel biocatalytic methods for highly efficient CTP cyclization.

## Data Availability

Detailed experimental procedures, characterization data, NMR spectra of compounds, detailed computational results and calculated structures are available within the Supplementary Information. Protein structures used in this paper are available at the PDB (SurE: 6KSU and Ulm16: 8FEK). Protein sequences used in this study are from the NCBI database: Ulm16 (accession ATU31793.1), SurE (BBZ90014.1), FlkO (AGI87381.1), WP516 (WP_04319516.1), WP_031183424.1, and SEC28301.1.

## Acknowledgements

We wish to thank Prof. Chittaranjan Das (Purdue University) and his students Rishi Patel and Gunaratne (Jayamini) Dhanapala for attempts to obtain a crystal structure of WP516, and Dr. César Aguilar for his help with bioinformatics. This work was funded by the National Institutes of Health (grant no. 1R35GM138002 to E.I.P.). Z.L.B. acknowledges the National Science Foundation for support under the Graduate Research Fellowship Program under grant no. DGE-1842166. We acknowledge the support from the Purdue Center for Cancer Research, NIH grant no. P30 CA023168.

## Author contributions

P.L.J., Z.L.B., and L.O.J. synthesized all the peptides used in this study, performed nuclear magnetic resonance analyses of the peptides, expressed the proteins, and performed the protein assays. P.L.J. and Z.L.B. performed the docking studies. P.L.J., Z.L.B., and E.I.P. conceived of the ideas and wrote the paper, with input from all authors. Project management and funding was the responsibility of E.I.P.

## Ethics declarations

The authors declare no competing interests

## Supplementary Information

Materials and methods, along with supplementary figures, tables, NMRs, and mass spectra can be found in the Supplementary Information.

## Notes

### Competing Interest Statement

The authors have declared no competing interest.

### Summary of Updates

Molecular Dynamics to further explore the alpha fold model Further mutational analysis of the enzymes to explore reasons for substrate scopes Scale up of enzyme reaction

