## Supplemental Files for "Biocatalytic Tetrapeptide Macrocyclization by Cryptic Penicillin-binding Protein-type Thioesterases"

Methods

**Reagents and materials**

Reagents were purchased from commercial sources and used without further purification. Dimethylformamide (DMF), dichloromethane (DCM), and acetonitrile (high performance liquid chromatography (HPLC) grade) were purchased from Fisher Scientific. Amino acids were purchased from either Chem-Impex or AAPTECC. Unless stated otherwise, all other reagents were purchased Sigma-Aldrich.

**Bioinformatic Search Using BiG-SCAPE CORASON and Substrate prediction.**

BiG-SCAPE CORASON output was generated following a previously established protocol with SurE as the input.[1] The CORASON output was analyzed in Interactive Tree of Life (iTOL) v6,[2] and selected PBP-TEs were chosen for further analysis with PRISM and expression. For the identification of SEC28301.1 a BlastP[3] search was conducted using the WP_043619516.1 amino acid sequence. For each selected PBP-TE their coding DNA sequence was searched on NCBI and the flanking 50,000 base pairs up and downstream of the PBP-TE were selected, downloaded, and uploaded into PRISM4.[4] The resulting output was then used to inform peptide thioester synthesis, and no additional modifications to the peptide (e.g. hydroxylation, chlorination) were assumed. In the case of an unnatural/noncommercially available amino acid prediction a close homolog of it was chosen (e.g. for enduracididine, arginine was used). If the predicted C-Terminal residue was a glycine we choose not to go forward with expression in fear of a narrower substrate scope. See **SI Figure 1** for a graphical representation of the workflow.

**WP516 a cloning for activity assays**

*Nonomuraea candida* HMC10 was obtained from the USDAs ARS Culture Collection (NRRL-B-24552) and grown in liquid ATCC 172 medium. Once the culture was sufficiently dense (~5 days) the genomic DNA was extracted following a previously reported protocol.[5] WP516 was amplified from the genomic DNA using the primer pair WP516-Fwd (5'-CAGCAGCCATCATCATCATCATCACatggacgatgtgatcgcacgg-3') and WP516-Rev (5'CTTTGTTAGCAGCCGGATCTCAGTGctagccatgccgccgg-3'). The uppercase letters represent sequence homology to the plasmid pET28b, and the lowercase letters represent sequence homology to the gene to be amplified. PCR amplification was performed with Q5 polymerase master mix. Each reaction contained 12.5 µL of the master mix (Q5 buffer, dNTPs, Q5 polymerase), 500 nM of forward and reverse primer, NRRL-B-24552 genomic DNA (1µL, 142 ng), GC enhancer (5 µL), and nuclease free water to a final volume of 25 µL. Thermocyclers were operated under the following program: (1) initial denaturation at 98 °C for 30 s; (2) 98 °C for 10 s; (3) 72 °C for 20 s; (4) 72 °C for 40 s ; (5) repeat steps 2–4 for a total of 34 cycles with a -0.3 °C in annealing temperature (step 4) per cycle; (6) 72 °C for 2 min and (7) hold at 4 °C. The products were analyzed by electrophoresis through a 1.5% agarose gel with the Gene Ruler 1kb Plus Ladder (Thermo Fisher Scientific) used as a control and showed only a single band at 1.5 kb. The PCR product was cleaned utilizing the GeneJET PCR Purification Kit (Thermo Fisher Scientific) using the standard protocol. The plasmid pET28b was PCR amplified using primers pET28-Fwd (5′-GTGATGATGATGATGATGGCTGCTGCCCATGG -3′) and pET28B-Rev (5′-CACTGAGATCCGGCTGCTAACAAAGCCCG-3′). PCR amplification was performed with Q5 polymerase. Each reaction contained the following: 5 µl of 5× Q5 buffer, 200 µM dNTPs, 500 nM of forward and reverse primer, 50 ng pET28b, 5 µl of Q5 High GC Enhancer, 0.5 µl Q5 polymerase and nuclease free water to a final volume of 25 µl. Thermocyclers were operated under the following program: (1) initial denaturation at 98 °C for 30 s; (2) 98 °C for 10 s; (3) 72 °C for 20 s; (4) 72 °C for 2 min 45 s ; (5) repeat steps 2–4 for a total of 34 cycles; (6) 72 °C for 2 min and (7) hold at 4 °C. The products were analyzed by electrophoresis through a 1.5% agarose gel with the Gene Ruler 1kb Plus Ladder used as a control and showed only a single band at 5 kb. The PCR product was cleaned utilizing the GeneJET PCR Purification Kit using the standard protocol. The gene fragment was then cloned into pET28b using NEB Gibson Assembly master mix to yield pET28b-WP516. Specifically, 65 ng of WP516 and 51 ng of linearized pET28b were combined (~5:1 insert:vector) with an equal volume of master mix and incubated at 50 °C for 60 minutes. NEB High Efficiency cells were then transformed with 2 µl of the mix before plating on LB plates supplemented with kanamycin (50 µg $mL^{-1}$). Colonies were then grown overnight in 5 mL of LB supplemented with kanamycin at 37 °C on an orbital shaker. Plasmids were isolated using the Thermo Scientific GeneJET Plasmid Miniprep Kit and verified by Sanger sequencing (GeneWiz). All DNA and amino acid sequences used in this study are reported in supplementary notes 1 and 2

SEC28301.1, WP516 Chimera, Ulm16 Chimera, and Ulm16$^{T304W}$ were purchased as a geneblock from *thermofisher* in a pET151 vector (SEC28031.1) or Twist Bioscience in a pET28a vector (all others) and used for expression without further modificiation.

**Protein overexpression and purification**

pET28b-*WP516*, or pET151-SEC28301.1 was introduced into BL21(DE3) competent cells by electroporation while pET28a-WP516 Chimera, pET28a-Ulm16 Chimera, and pET28a-Ulm16$^{T304W}$ were electroporated into T7 Express competent cells. Transformed cells were selected on LB agar plates supplemented with kanamycin (50 µg $mL^{-1}$). Single colonies were picked and inoculated in 5 mL LB containing 50 µg $mL^{-1}$ kanamycin and grown for 16 h at 37 °C on an orbital shaker. This overnight culture was then used to inoculate 1 L LB with 50 µg $mL^{-1}$ kanamycin, and the culture was then grown at 37 °C with shaking at 250 r.p.m. until an optical density ($OD_{600}$) of 0.6 was achieved. The culture was cooled on ice prior to the addition of 0.2 mM isopropyl-1-thio-β-D-galactopyranoside (IPTG) and then grown at 18 °C for 18 h with shaking at 250 r.p.m. Cells were then collected via centrifugation (5,000g, 10 min at 4 °C) and frozen at -80 °C overnight. Cell pellets were then thawed on ice for 30 min prior to the addition of 25 mL of lysis buffer containing 50 mM Tris-HCl, 300 mM NaCl, 10 mM imidazole, and 10% glycerol (v/v) at pH 8.0. The resulting suspension was then mixed with 1 mg $mL^{-1}$ lysozyme, 85 µg RNase A, 300 units of DNase I, and 100 µM phenylmethylsulfonyl fluoride at 4 °C for 30 min. The cell lysate was then clear via centrifugation (17,000g, 30 min at 4°C) and the supernatant was collected and mixed with 1 mL HisPur Ni-NTA resin (Thermo Scientific) on an orbital shaker at 4 °C. The resin was loaded onto a 10 mL Pierce centrifuge column and washed with 20 mL of wash buffer containing 50 mM Tris-HCl, 300 mM NaCl, 20 mM imidazole, and 10% glycerol (v/v) at pH 8.0. The protein was eluted using 20 mL of elution buffer containing 50 mM Tris-HCl, 300 mM NaCl, 300 mM imidazole, and 10% glycerol (v/v) at pH 8.0 with 1 mL fractions collected. The fractions were assessed by SDS-PAGE using 4-15% TGX Mini-PROTEAN gel (Bio-Rad) and stained with Coomassie stain. Fractions that contained pure protein were combined and concentrated using an Amicon Ultra-15 centrifugal filter with a 30 kDa cutoff (EMD Millipore). The resulting samples were exchanged into storage buffer containing 25 mM Tris-HCl, 50 mM NaCl, 5% glycerol (v/v), and 1 mM DTT at pH 8.0. The Pierce BCA Protein Assay Kit (Thermo Scientific) was used to determine protein concentration following the standard protocol. Ulm16 was prepared as previously reported.[6]

**Peptide synthesis**

All Peptides reported in this paper were synthesized according to this standard protocol. Peptides 1, 2, 3, 38, and 39 were used from a previously reported synthesis and their spectral data can be found there.[2] In this study, UPLC and/or analytical HPLC were utilized to assess the purity of the peptide prior to assay or purification by monitoring absorbance at 214 nm. The analytical HPLC analysis was performed on a Luna Omega5 µm Polar C18 100 Å 150x4.6 mm (Phenomenex) column, while the UPLC analysis was carried out on a CORTECS T3 Column, 120Å, 1.6 µm, 2.1 mm X 50 mm (Waters) column. For purification, semi-preparative was employed using a Luna Omega 5 µm Polar C18 100 Å 150x21.2 mm (Phenomenex) column. The specific gradient and flow rate for each peptide can be found in their respective sections. NMRs were taken on a Bruker AV800 spectrometer and analyzed on mestrenova.

- <u>Attachment of Fmoc-MeDbz-OH</u>

  Here, 0.1 mmol Fmoc-Gly Rink amide resin (Chem-Impex), in a 5 mL fritted polypropylene syringe (Torviq), was allowed to swell in DMF (5 mL) for 20 minutes. The resin was then subjected to 20% piperidine in DMF (4 mL) for 15 minutes. The solution was then drained, and the resin was washed with DMF (3 x 4 mL) and DCM (3 x 4 mL). The Kaiser ninhydrin test was used to verify successful Fmoc deprotection. The resin was then washed with DMF (3 x 4 mL) and a solution containing Fmoc-MeDbz-OH (synthesized according to a previously published route[7]) (0.3 mmol), PyOxim (Chem-Impex) (0.5 mmol), and i-$Pr_2$EtN (0.7 mmol) in 4 mL of DMF was added to the resin and allowed to shake for 1.5 h. The solution was drained, and the resin was washed with DMF (3 x 4 mL) and DCM (3 x 4 mL). Based on a negative Kaiser ninhydrin test, the loading was assumed to be 100%. The resin was washed with DMF (3 x 4 mL) and used in solid-phase peptide synthesis.

- <u>SPPS</u>
  Manual solid-phase peptide synthesis (SPPS) was performed in 5 mL fritted polypropylene syringes. The resin was treated with 20% piperidine in DMF (4 mL) for 15 minutes. The solution was then drained, and the resin washed with DMF (3 x 4 mL), DCM (3 x 4 mL), and DMF (3 x 4 mL). A solution of Fmoc-AA-OH (0.5 mmol), PyOxim (0.5 mmol), and i-$Pr_2$EtN (0.7 mmol) in 3 mL of DMF was added and the resin was allowed to shake for 1 h. The solution was drained, and the resin washed with DMF (3 x 4 mL), DCM (3 x 4 mL), and DMF (3 x 4 mL). Deprotection and coupling cycles were repeated until the desired peptide sequence was achieved. Following the coupling of the first amino acid, the Kaiser ninhydrin test was used to determine deprotection and coupling reaction completion. The last amino acid in the peptide sequence was N-Boc protected.
- <u>Activation and thiolysis</u>
  Following the coupling of the last amino acid, the resin was washed with DCM (4 x 5 mL). The resin was treated with 3 mL of 0.5 M 4-nitrophenyl chloroformate in DCM and allowed to shake for 1.5 h. The solution was the drained and the resin washed with DCM (3 x 5 mL) and DMF (3 x 5 mL). The resin was then subjected to 4 mL of 1 M i-$Pr_2$EtN in DMF and allowed to shake for 15 min. The solution was drained, and the resin washed with DMF (3 x 5 mL) and treated with more 1 M i-$Pr_2$EtN in DMF solution. This process was repeated until the solution no longer turned yellow. The resin was then treated with a 1:2 thiol:DMF solution (3 mL) and allowed to shake overnight. The solution was then drained, and the resin washed with DCM (3 x 4 mL). The filtrate and washes were combined and subjected to rotary evaporation and lyophilization.
- <u>Global deprotection</u>
  The crude peptide thioester was treated with a 90:10 trifluoroacetic acid:triisopropylsilane solution (5 mL) for 1.5 h. The solution was then removed by a stream of nitrogen and diethyl ether was added to precipitate the peptide thioester. The precipitate was collected via centrifugation and then lyophilized. The resulting dry peptide thioester was purified via HPLC and characterized by UPLC-MS.

**WP516 kinetic assays**

A 100 µL reaction mixture containing Tris-HCl buffer (pH 8.0), 5% DMSO, 24-600 µM substrate, and a concentration of WP516 dependent on the peptide was left to stand at room temperature (21-23 °C) for 5 min. The reaction was then quenched with 50 µL of acetonitrile and centrifuged at 21,000g for 10 min. 5 µL of the supernatant was loaded onto a CORTECS T3 Column, 120 Å, 1.6 µm, 2.1 × 50 mm (Waters) and separated by UPLC using mobile phases of $H_2O$ + 0.1% formic acid and acetonitrile + 0.1% formic acid with monitoring at 214 nm. Unless stated otherwise, peptides were eluted using an 11 min 0-40% mobile phase B gradient at a flow rate of 0.5 mL $min^{-1}$. All reactions were performed in triplicate. The spectra were analyzed using the waters_connect (Unifi) software v.3.0.0.15. The extinction coefficient $\varepsilon$ (214 nm) of the peptide thioester substrates was used to calculate the cyclization rate, with the assumption that the $\varepsilon$ (214 nm) values of the peptide thioesters were equal to that of their corresponding cyclic peptides. The $\varepsilon$ (214 nm) value of SMMP-dY-dR-dF-V was used to calculate the cyclization rate of SMMP-dY-R-dF-V. The Michaelis–Menten equation and PRISM v.10 were used to estimate the kinetic parameters.

**WP516 TTN assays**

A 100 µL reaction mixture containing Tris-HCl buffer (pH 8.0), 5% DMSO, 400 µM substrate, and a concentration of WP516 dependent on the peptide was left to stand at room temperature or incubated at 30 °C, depending on the substrate, for 4 h. The reaction was then quenched with 100 µL of acetonitrile and centrifuged at 21,000g for 10 min. 5 µL of the supernatant was loaded onto a CORTECS T3 Column, 120 Å, 1.6 µm, 2.1 × 50 mm (Waters) and separated by UPLC using mobile phases of $H_2O$ + 0.1% formic acid and acetonitrile + 0.1% formic acid with monitoring at 214 nm. Unless stated otherwise, peptides were eluted using a 0-40% mobile phase B in 11 min gradient at a flow rate of 0.5 mL $min^{-1}$. All reactions were performed in triplicate and TTNs were calculated according to the following equation:

$$\mathrm{TTN} = \frac{Area_{Product}}{Area_{Product} + Area_{Hydrolysis} + Area_{Starting\ Material}} \times \frac{[Substrate]_{Total}}{[Enzyme]_{Total}}$$

**WP516 Scale Up Reaction**

A 40 mL reaction mixture containing Tris-HCl buffer (pH 8.0), 2.5% DMSO, 1.2 mM crude substrate, and 0.15 µM WP516 was left to stand at room temperature for 2 hours. Aliquots were taken from the reaction mixture at 5 min, 20 min, 45 min, 60 min, and 120 min, quenched with acetonitrile, and analyzed via UPLC as previously described for the TTN assays to assess reaction progress (**SI Figure S56**). After 2 hours, the reaction was split into two 20 mL aliquots which were each quenched with 20 mL acetonitrile and subjected to lyophilization. Following combination of the resulting solids, ~4 mL of methanol was added, and the mixture was subjected to sonication followed by centrifugation. The supernatant was loaded onto a Luna Omega 5 µm Polar C18 100 Å 150x21.2 mm (Phenomenex) column to be purified by reverse-phase semi-preparative HPLC with mobile phases of $H_2O$ + 0.05% TFA (A) and acetonitrile + 0.05% TFA (B). The peptide was eluted using a gradient of 0-60% mobile phase B over 22 minutes at a 20 mL/min flow rate. The column was equilibrated with 0% mobile phase B for 1 minute before and 5 minutes after the gradient. This process was repeated on the remaining pellet and continued until no cyclic peptide remained in the pellet. This process ultimately yielded the cyclic peptide product as an off-white solid (16 mg, 50% yield) that was characterized by UPLC-MS and NMR spectroscopy (**SI Figures S57 and S58**).

**$^1$H NMR** (800 MHz, DMSO) δ 9.18 (s, 1H), 8.35 (d, *J* = 9.2 Hz, 1H), 8.26 (d, *J* = 9.2 Hz, 1H), 7.57 (t, *J* = 5.7 Hz, 1H), 7.35 (q, *J* = 9.6 Hz, 3H), 7.40-6.79 (m, 13H), 6.63-6.56 (m, 2H), 4.56-4.40 (m, 2H), 4.27 (q, *J* = 7.8 Hz, 1H), 3.90 (t, *J* = 10.0 Hz, 1H), 3.08-2.86 (m, 4H), 2.82 (dd, *J* = 8.7, 14.2 Hz, 1H), 2.65 (dd, *J* = 8.1, 14.2 Hz, 1H), 1.89-1.78 (m, 1H), 1.60-1.20 (m, 4H), 0.76 (d, *J* = 6.6 Hz, 3H), 0.65 (d, *J* = 6.6 Hz, 3H).

**$^{13}$C NMR** (201 MHz, DMSO) δ 173.65, 173.39, 173.19, 172.98, 157.18, 156.14, 138.37, 130.25, 129.41, 128.47, 128.34, 126.64, 115.35, 59.92, 53.98, 53.36, 52.70, 40.72, 34.34, 33.89, 27.82, 26.89, 25.40, 19.34, 18.92.

**Mass spec:** expected neutral mass for $C_{29}H_{39}N_7O_5$ (Da): 565.30127, observed neutral mass (Da): 566.3071, mass error (ppm): -2.6.

**Multiple Sequence Alignment**

Ulm16 (Accession ATU31793.1), SurE (BBZ90014.1), and the amino acid sequence for WP_043619516.1 were used as inputs for a Clustal Omega multiple sequence alignment (MSA). The alignment was performed and analyzed in EMBL-EBI Job Dispatcher.[8]

**Alphafold3 Model Generation**

WP_043619516.1 amino acid sequence was used as the input for the modeling and carried out utilizing the standard settings in the alphafold 3 server.[9] The highest scoring output was use for all modeling and comparison with other PBP-TEs.

**Docking**

For the covalent docking simulation, the structure of Ulm16 (PDB 8FEK) and WP516 (Alphafold 3 model) was refined using the Protein Preparation Wizard in Maestro (Schrödinger Suite 2022-4, Schrödinger LLC), using PROPKA in the hydrogen bond assignment step with the pH set at 8.0. Peptides docked in this study were made as C-Terminal Acyl fluorides and suitable 3D coordinates and states were generated using LigPrep. Peptides were covalently docked using Glide through precision mode and nucleophilic C-F substitution as the reaction type. The center of the grid box for the simulation was set around the active site serine of Ulm16 and post-docking minimization was conducted for 200 poses. The top 5 poses (by MMGBSA) for each peptide were chosen an analyzed using Pymol.

**Molecular Dynamics Simulations**

To obtain the models for molecular dynamics (MD) simulations, covalent docking was performed as earlier described, with the missing loops in the Ulm16 crystal structure modeled in using homology modeling during protein preparation. The pose to use for simulation for each model was chosen based on docking score and, if the output poses varied, conformations seemingly conducive to catalysis. To obtain parameters for these models, Gaussian 16[10] was used to performed optimizations of the ligand at the level of B3LYP[11]/6-31G(d) and electrostatic potential calculations. Restrained electrostatic potential charge fitting and generation of parameters of the bonds, angles, dihedral angles, and van der Waals radii was conducted using the Antechamber package within the AmberTools program.[12] The general Amber force field (GAFF)[13]was used for the ligand and the ff14SB[14] force was used for the protein. Following tleap processing of the parameterized ligand and protein complex, the models were solvated within an octahedral box of TIP3P water molecules and sodium ions were added to neutralize the systems. GROMACS (version 2022.3) was used to perform the MD simulations. The Particle Mesh Ewald (PME) method[15] was used to consider long-range electrostatic interactions and the LINCS[16] constraint algorithm was employed. The non-bonding cutoff value was set to 10 Å. To prepare the systems for simulation, first, a 50 ps energy minimization was performed to ensure a reasonable starting structure. A 100 ps NVT equilibration followed by a 50 ps NPT equilibration were then conducted. The final frame of the NPT equilibration was used as the starting structure for production MD. For each model, four independent simulation replicates were performed starting from the same structure.

### Synthesis of Peptide Thioester Substrates

SBMP-dY-R-F-V

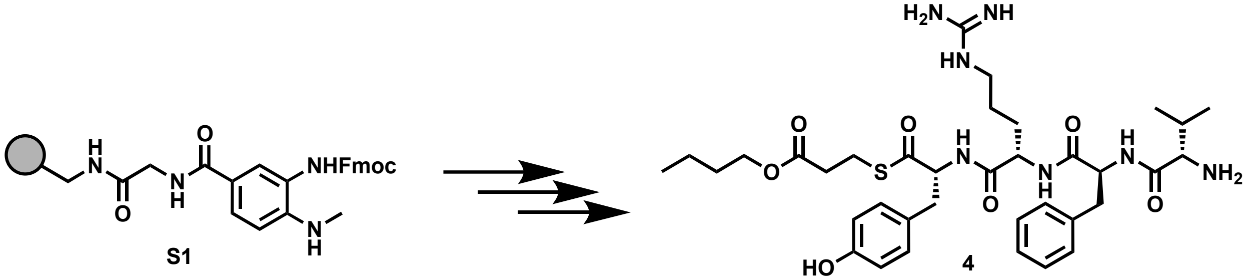

Peptide **4** was synthesized following the peptide synthesis protocol outlined above. Starting with Fmoc-Me-Dbz-OH loaded resin **S1** (0.1 mmol), amino acids Fmoc-D-Tyr(tBu)-OH, Fmoc-L-Arg(Pbf)-OH, Fmoc-D-Phe-OH, and Boc-L-Val-OH were used with butyl 3-mercaptopropionate as the cleaving thiol. The crude peptide was purified by reverse-phase semi-preparative HPLC with mobile phases of $H_2O$ + 0.05% TFA (A) and acetonitrile + 0.05% TFA (B). The peptide was eluted using a gradient of 0-55% mobile phase B over 25 minutes at a 20 mL/min flow rate. The column was equilibrated with 0% mobile phase B for 1 minute before and 5 minutes after the gradient, yielding an off-white solid (52 mg, 54% yield).

**$^1$H NMR** (800 MHz, DMSO) δ 9.32 (s, 1H), 8.69 (d, $J$ = 7.5 Hz, 1H), 8.56 (d, $J$ = 6.9 Hz, 1H), 8.32 (d, $J$ = 6.6 Hz, 1H), 8.00 (m, 8.06-7.91 (m, 3H), 7.72 (s, 1H), 7.59-6.93 (m, 11H), 6.65 (d, $J$ = 7.4 Hz, 2H), 4.66 (t, $J$ = 9.7 Hz, 1H), 4.47 (t, $J$ = 9.2 Hz, 1H), 4.37 (dd, $J$ = 7.1, 13.7 Hz, 1H), 4.00 (t, $J$ = 6.1 Hz, 2H), 3.61 (s, 1H), 3.00 (dd, $J$ = 7.3, 11.0 Hz, 6H), 2.80 (t, $J$ = 11.2 Hz, 1H), 2.72 (t, $J$ = 11.7 Hz, 1H), 2.56 (t, $J$ = 5.6 Hz, 2H), 2.09 (dt, $J$ = 7.8, 13.5 Hz, 1H), 1.61-1.44 (m, 3H), 1.43-1.15 (m, 5H), 0.99-0.79 (m, 10H).

**$^{13}$C NMR** (201 MHz, DMSO) δ 200.59, 171.91, 171.60, 170.93, 168.48, 157.36, 156.55, 137.94, 130.58, 129.67, 128.57, 127.35, 126.80, 115.49, 64.37, 61.45, 57.48, 54.50, 52.41, 40.73, 37.82, 36.53, 34.07, 30.57, 30.40, 29.79, 25.22, 23.89, 19.04, 18.85, 17.53, 14.00.

**Mass spec:** expected neutral mass for $C_{36}H_{53}N_7O_7S$ (Da): 727.3727, observed neutral mass (Da): 727.3744, mass error (ppm): 2.3.

**UPLC Trace** Obtained with mobile phases of $H_2O$ + 0.1% formic acid (A) and acetonitrile + 0.1% formic acid (B). The peptide was eluted using a gradient of 0-70% mobile phase B over 4 min at a 0.5 mL/min flow rate. Mobile phase B was held at 70% for 1 min. The column was equilibrated with 0% mobile phase for 1 minute before and 2 minutes after the gradient. The peptide purity was determined to be 98%.

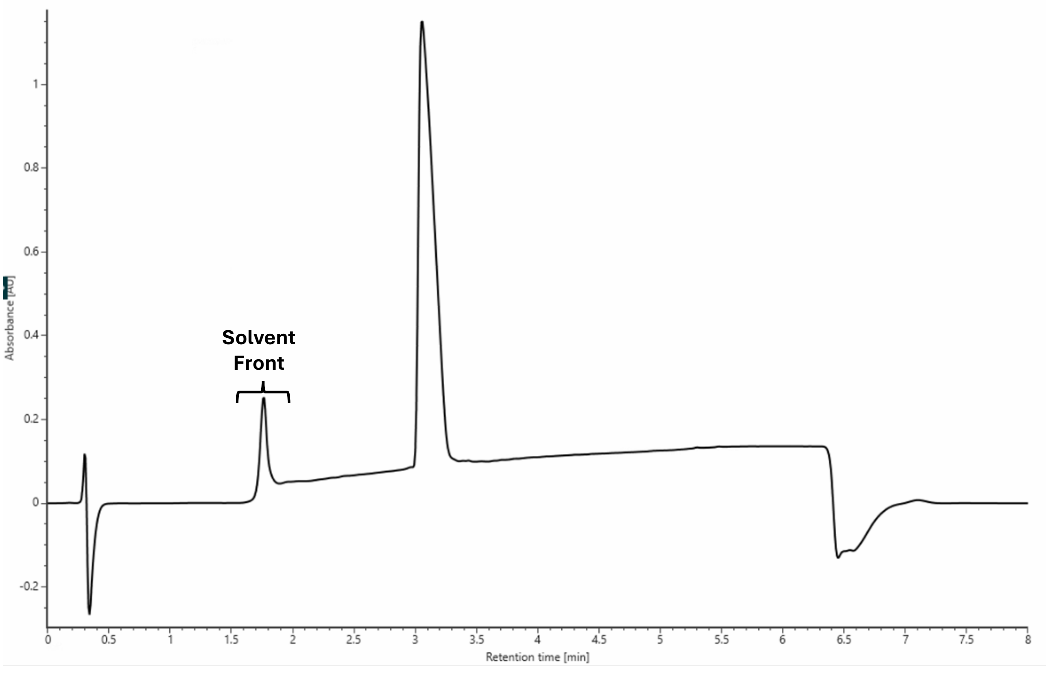

SBMP-dY-Dap-dF-V

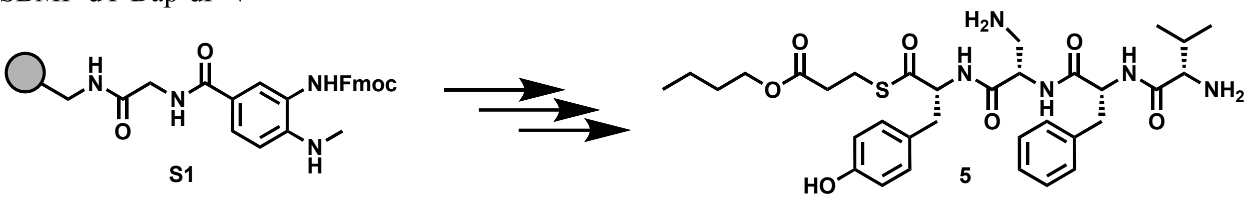

Peptide **5** was synthesized following the peptide synthesis protocol outlined above. Starting with Fmoc-Me-Dbz-OH loaded resin S1 (0.1 mmol), amino acids Fmoc-D-Tyr(tBu)-OH, Fmoc-L-Dap(Boc)-OH, Fmoc-D-Phe-OH, and Boc-L-Val-OH were used with butyl 3-mercaptopropionate as the cleaving thiol. The crude peptide was purified by reverse-phase semi-preparative HPLC with mobile phases of $H_2O$ + 0.05% TFA (A) and acetonitrile + 0.05% TFA (B). The peptide was eluted using a gradient of 0-55% mobile phase B over 25 minutes at a 20 mL/min flow rate. The column was equilibrated with 0% mobile phase B for 1 minute before and 5 minutes after the gradient, yielding an off-white solid (24 mg, 27% yield).

**$^1$H NMR** (800 MHz, DMSO) δ 8.89 (d, *J* = 7.6 Hz, 1H), 8.77 – 8.71 (m, 2H), 8.13 (s, 3H), 7.94 (s, 3H), 7.31 – 7.21 (m, 4H), 7.17(t, *J* = 6.0 Hz, 1H), 6.99 (d, *J* = 7.8 Hz, 2H), 6.66 (d, *J* = 7.8 Hz, 1H), 4.83 – 4.78 (m, 1H), 4.63 (sextet, *J* = 4.3 Hz, 1H), 4.53 (q, *J* = 6.6 Hz, 1H), 4.03 (t, *J* = 6.2 Hz, 2H), 3.27 (d, *J* = 14.1 Hz, 1H), 3.06 (d, *J* = 7.8 Hz, 1H), 3.02 – 2.95 (m, 3H), 2.79 (dd, *J* = 8.7, 12.6 Hz, 1H), 2.74 (t, *J* = 8.6 Hz, 1H), 2.69 (t, *J* = 13.4 Hz, 1H), 2.56 (t, *J* = 6.2 Hz, 2H), 1.82 (sextet, 6.0 Hz, 1H), 1.54 (quintet, *J* = 7.3 Hz, 2H), 1.32 (quintet, *J* = 7.2 Hz, 2H), 0.88 (t, *J* = 7.1 Hz, 3H), 0.67 (d, *J* = 6.9 Hz, 3H), 0.37 (d, *J* = 7.0 Hz, 3H).

**$^{13}$C NMR** (201 MHz, DMSO) δ 199.85, 171.93, 171.57, 168.85, 168.12, 156.56, 137.93, 130.52, 129.54, 128.43, 126.87, 126.74, 115.48, 64.35, 61.41, 57.62, 54.31, 51.08, 38.61, 36.71, 33.93, 30.52, 29.97, 23.84, 18.98, 18.69, 16.62, 13.93.

Mass spec: expected neutral mass for $C_{33}H_{47}N_5O_7S$ (Da): 657.3196, observed neutral mass (Da): 657.3200, mass error (ppm): 0.5.

UPLC Trace: Obtained with mobile phases of $H_2O$ + 0.1% formic acid (A) and acetonitrile + 0.1% formic acid (B). The peptide was eluted using a gradient of 0-70% mobile phase B over 4 min at a 0.5 mL/min flow rate. Mobile phase B was held at 70% for 1 min. The column was equilibrated with 0% mobile phase for 1 minute before and 2 minutes after the gradient. The peptide purity was determined to be 97%.

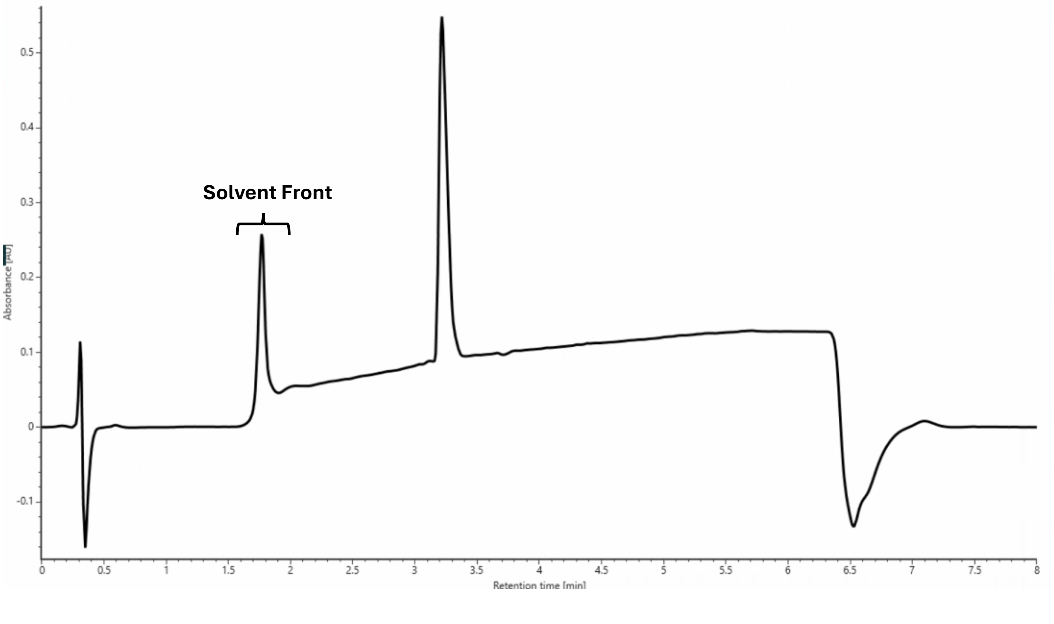

SBMP-dY-dDap-dF-V

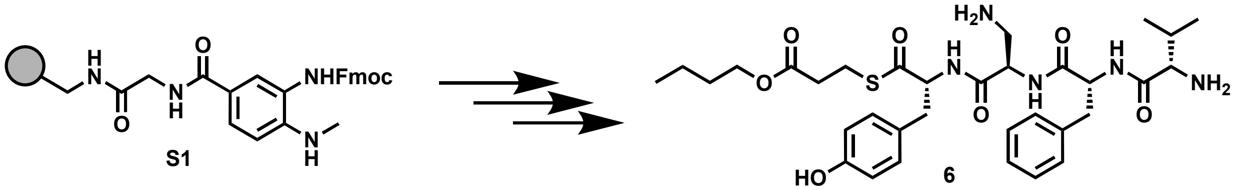

Peptide **6** was synthesized following the peptide synthesis protocol outlined above. Starting with Fmoc-Me-Dbz-OH loaded resin **S1** (0.1 mmol), amino acids Fmoc-D-Tyr(tBu)-OH, Fmoc-D-Dap(Boc)-OH, Fmoc-D-Phe-OH, and Boc-L-Val-OH were used with butyl 3-mercaptopropionate as the cleaving thiol. The crude peptide was purified by reverse-phase semi-preparative HPLC with mobile phases of $H_2O$ + 0.05% TFA (A) and acetonitrile + 0.05% TFA (B). The peptide was eluted using a gradient of 0-55% mobile phase B over 25 minutes at a 20 mL/min flow rate. The column was equilibrated with 0% mobile phase B for 1 minute before and 5 minutes after the gradient, yielding an off-white solid (21 mg, 24% yield).

**$^1$H NMR** (800 MHz, DMSO) δ 8.75 (q, $J$ = 7.2 Hz 2H), 8.69 (d, $J$ = 6.2 Hz, 1H), 8.17 (s, 3H), 7.99 (s, 3H), 7.27 – 7.21 (m, 3H), 7.20 – 7.15 (m, 1H), 7.01 (d, 5.8 Hz, 2H), 6.65 (d, 5.8 Hz, 2H), 4.77 (t, $J$ = 9.8 Hz, 1H), 4.64 (t, $J$ = 5.2 Hz, 1H), 4.50 (t, $J$ = 5.8 Hz, 1H), 4.02 (q, $J$ = 5.5 Hz, 2H), 3.61 (s, 1H), 3.22 (d, $J$ = 11.6 Hz, 2H), 3.03 – 2.89 (m, 4H), 2.86 (dd, $J$ = 7.1, 11.0 Hz, 1H), 2.71 (t, $J$ = 11.1 Hz, 1H), 2.59-2.51 (m, 2H), 1.88 – 1.82 (m, 1H), 1.53 (q, $J$ = 6.2 Hz, 2H), 1.31 (q, $J$ = 6.8 Hz, 2H), 0.88 (q, $J$ = 6.3 Hz, 3H), 0.69 (d, $J$ = 5.1 Hz, 3H), 0.41 (d, $J$ = 5.2 Hz, 3H).

**$^{13}$C NMR** (201 MHz, DMSO) δ 200.01, 171.85, 171.53, 169.22, 168.15, 156.53, 137.80, 130.43, 129.51, 128.46, 126.90, 126.75, 115.49, 64.32, 61.58, 57.62, 54.20, 51.28, 39.88, 38.30, 36.37, 33.91, 30.51, 30.00, 23.80, 18.96, 18.67, 16.64, 13.91.

**Mass spec:** expected neutral mass for $C_{33}H_{47}N_5O_7S$ (Da): 657.3196, observed neutral mass (Da): 657.3201, mass error (ppm): 0.7.

**UPLC Trace:** Obtained with mobile phases of $H_2O$ + 0.1% formic acid (A) and acetonitrile + 0.1% formic acid (B). The peptide was eluted using a gradient of 0-70% mobile phase B over 4 min at a 0.5 mL/min flow rate. Mobile phase B was held at 70% for 1 min. The column was equilibrated with 0% mobile phase for 1 minute before and 2 minutes after the gradient. The peptide purity was determined to be 98%.

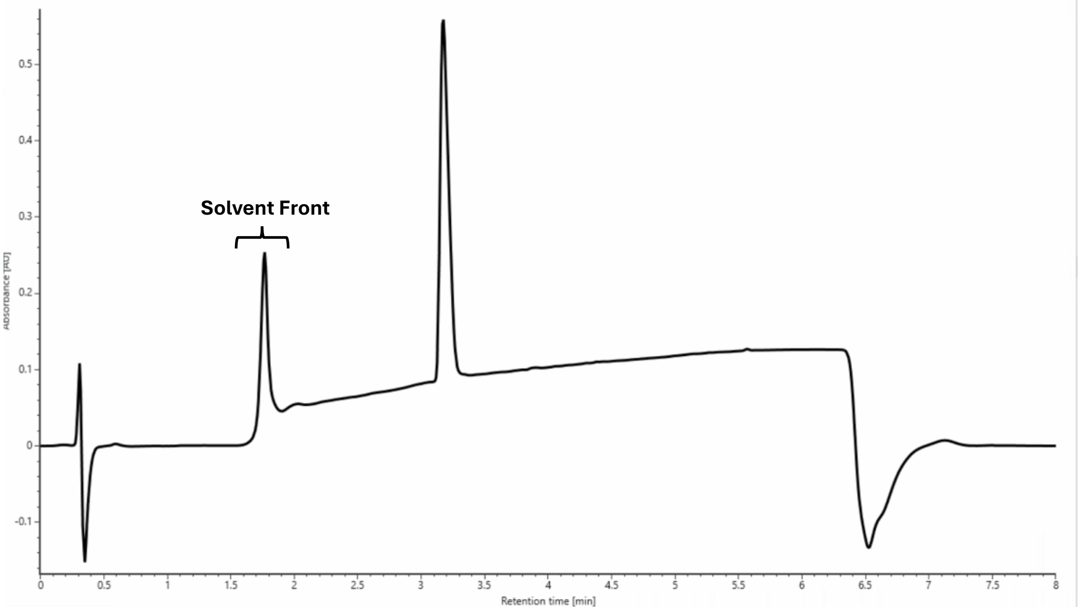

SBMP-dY-dDap-F-V

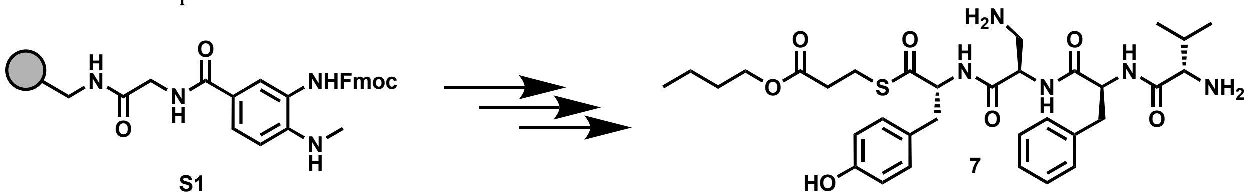

Peptide **7** was synthesized following the peptide synthesis protocol outlined above. Starting with Fmoc-Me-Dbz-OH loaded resin **S1** (0.1 mmol), amino acids Fmoc-D-Tyr(tBu)-OH, Fmoc-D-Dap(Boc)-OH, Fmoc-L-Phe-OH, and Boc-L-Val-OH were used with butyl 3-mercaptopropionate as the cleaving thiol. The crude peptide was purified by reverse-phase semi-preparative HPLC with mobile phases of $H_2O$ + 0.05% TFA (A) and acetonitrile + 0.05% TFA (B). The peptide was eluted using a gradient of 0-55% mobile phase B over 25 minutes at a 20 mL/min flow rate. The column was equilibrated with 0% mobile phase B for 1 minute before and 5 minutes after the gradient, yielding an off-white solid (22 mg, 25% yield).

**$^1$H NMR** (800 MHz, DMSO) δ 9.34 (s, 1H), 8.83 – 8.77 (m, 2H), 8.50 (t, *J* = 5.3 Hz, 1H), 7.97-8.14 (m, 5H), 7.30 (dd, *J* = 15.9, 7.2 Hz, 4H), 7.22 (d, *J* = 6.8 Hz, 1H), 7.01 (t, *J* = 6.6 Hz, 2H), 6.65 (t, *J* = 6.5 Hz, 2H), 4.73 – 4.49 (m, 2H), 4.43 (quintet, *J* = 7.0 Hz, 1H), 4.03 (q, *J* = 5.9 Hz, 2H), 3.60 (s, 1H), 3.22 – 3.16 (m, 2H), 3.01 – 2.91 (m, 3H), 2.92 – 2.77 (m, 3H), 2.53 (d, *J* = 6.9 Hz, 2H), 2.05 (quintet, *J* = 6.5 Hz, 1H), 1.55 (q, *J* = 7.1 Hz, 2H), 1.32 (quintet, *J* = 7.0 Hz, 2H), 0.96 – 0.83 (m, 9H).

**$^{13}$C NMR** (201 MHz, DMSO) δ 199.95, 171.79, 171.50, 169.46, 168.92, 158.92, 158.77, 156.51, 137.89, 130.44, 129.52, 128.57, 127.07, 126.84, 115.50, 64.32, 61.79, 57.40, 55.16, 50.67, 40.09, 39.99, 39.89, 39.78, 39.68, 36.89, 36.42, 33.94, 30.51, 23.73, 18.98, 18.80, 17.78, 13.93.

**Mass Spec:** expected neutral mass for $C_{33}H_{47}N_5O_7S$ (Da): 657.3196, observed neutral mass (Da): 657.3209, mass error (ppm): 1.9.

**UPLC Trace:** Obtained with mobile phases of $H_2O$ + 0.1% formic acid (A) and acetonitrile + 0.1% formic acid (B). The peptide was eluted using a gradient of 0-70% mobile phase B over 4 min at a 0.5 mL/min flow rate. Mobile phase B was held at 70% for 1 min. The column was equilibrated with 0% mobile phase for 1 minute before and 2 minutes after the gradient. The peptide purity was determined to be 98%.

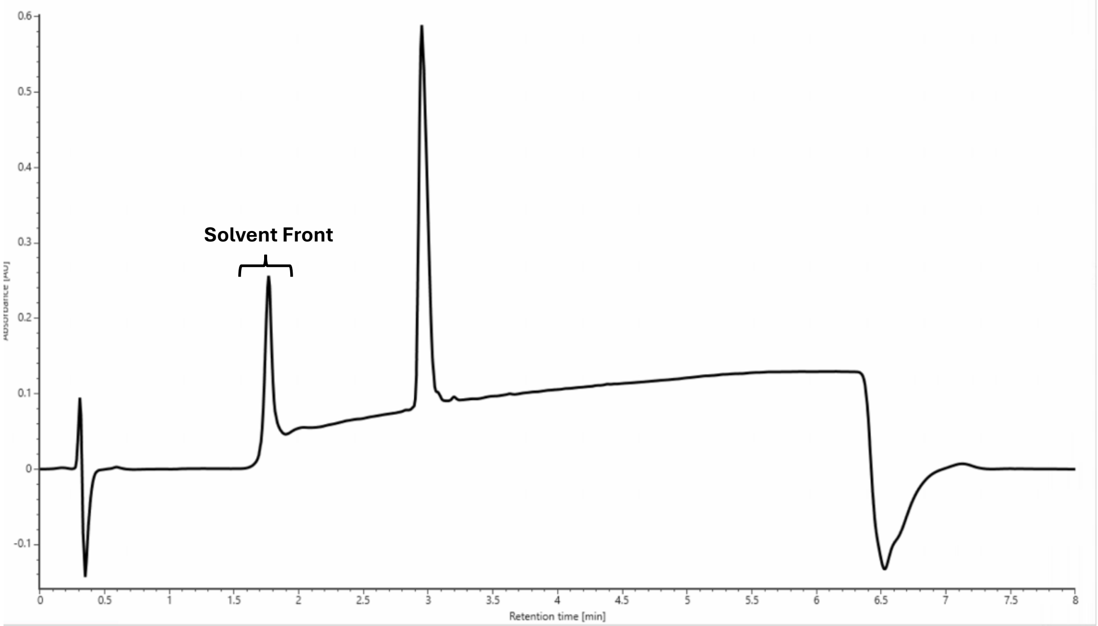

SBMP-dY-Dap-F-V

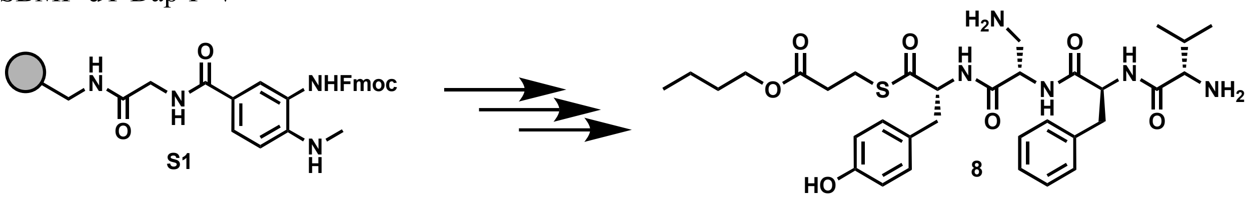

Peptide **8** was synthesized following the peptide synthesis protocol outlined above. Starting with Fmoc-Me-Dbz-OH loaded resin **S1** (0.1 mmol), amino acids Fmoc-D-Tyr(tBu)-OH, Fmoc-L-Dap(Boc)-OH, Fmoc-L-Phe-OH, and Boc-L-Val-OH were used with butyl 3-mercaptopropionate as the cleaving thiol. The crude peptide was purified by reverse-phase semi-preparative HPLC with mobile phases of $H_2O$ + 0.05% TFA (A) and acetonitrile + 0.05% TFA (B). The peptide was eluted using a gradient of 0-55% mobile phase B over 25 minutes at a 20 mL/min flow rate. The column was equilibrated with 0% mobile phase B for 1 minute before and 5 minutes after the gradient, yielding an off-white solid (24 mg, 27% yield).

**$^1$H NMR** (800 MHz, DMSO) δ 9.36 (s, 1H), 8.88 (d, *J* = 8.2 Hz, 1H), 8.67 (d, *J* = 8.1 Hz, 1H), 8.59 (d, *J* = 7.3 Hz, 1H), 8.15-8.02 (m, 6H), 7.31-7.22 (m, 4H), 7.20 (t, *J* = 6.8 Hz, 1H), 7.01 (d, *J* = 7.9, 2H), 6.66 (d, *J* = 8.1, 2H), 4.70-4.59 (m, 2H), 4.52 (dd, *J* = 7.3, 13.9 Hz, 1H), 4.01 (t, *J* = 6.7 Hz, 2H), 3.13 (d, *J* = 13.2 Hz, 1H), 3.08-2.96 (m, 4H), 2.87 (dd, *J* = 9.2, 13.2 Hz, 1H), 2.79 (dd, *J* = 9.0, 13.1, 2H), 2.57 (t, *J* = 6.6 Hz, 2H), 2.09 (dt, *J* = 6.1, 12.2 Hz, 1H), 1.53 (quintet, *J* = 6.8 Hz, 2H), 1.32 (sextet, *J* = 7.5 Hz, 2H), 0.95 (d, *J* = 6.7 Hz, 3H), 0.92-0.85 (m, 6H).

**$^{13}$C NMR** (201 MHz, DMSO) δ 199.94, 171.60, 171.49, 169.04, 168.48, 156.60, 137.69, 130.58, 130.52, 129.80, 128.57, 127.10, 126.84, 115.54, 64.40, 61.56, 57.53, 54.60, 51.03, 37.66, 36.44, 34.03, 30.58, 23.93, 18.92, 17.64, 14.00.

**Mass spec:** expected neutral mass for $C_{33}H_{47}N_5O_7S$ (Da): 657.3196, observed neutral mass (Da): 657.3283, mass error (ppm): 2.1.

**UPLC Trace** Obtained with mobile phases of $H_2O$ + 0.1% formic acid (A) and acetonitrile + 0.1% formic acid (B). The peptide was eluted using a gradient of 0-70% mobile phase B over 4 min at a 0.5 mL/min flow rate. Mobile phase B was held at 70% for 1 min. The column was equilibrated with 0% mobile phase for 1 minute before and 2 minutes after the gradient. The peptide purity was determined to be 97%.

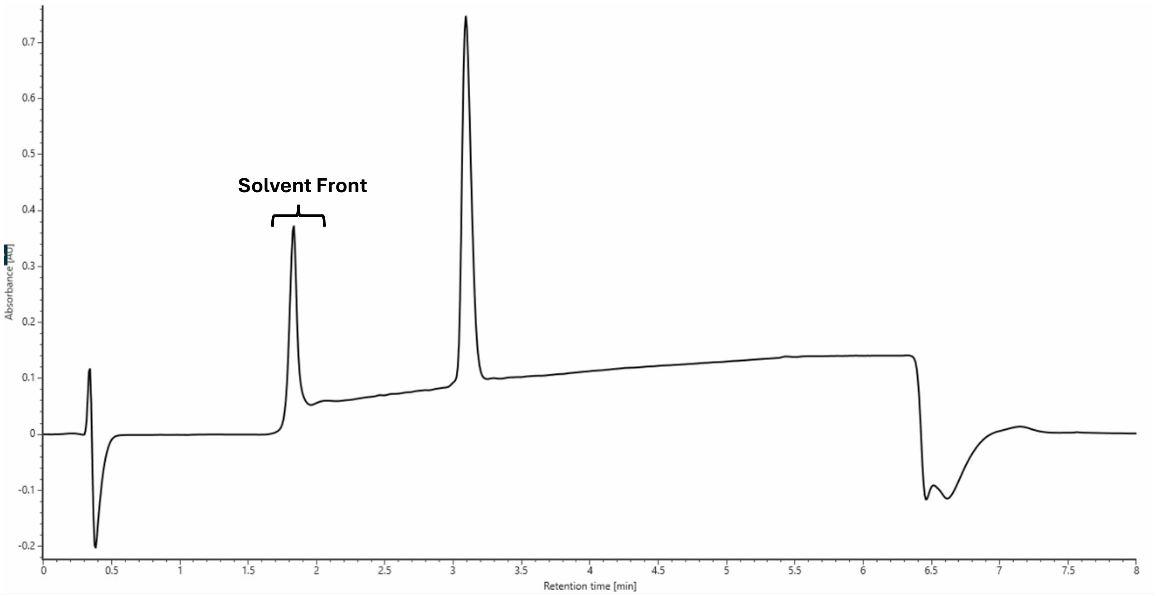

SMMP-dQ-R-dF-V

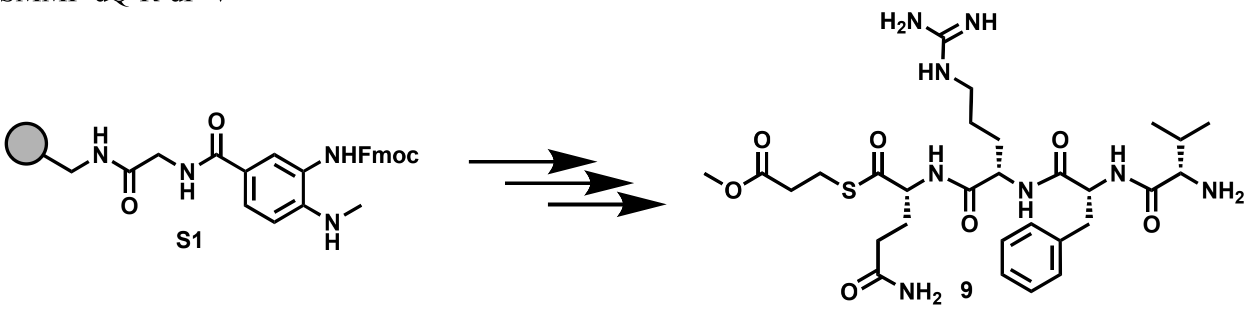

Peptide **9** was synthesized following the peptide synthesis protocol outlined above. Starting with Fmoc-Me-Dbz-OH loaded resin **S1** (0.1 mmol), amino acids Fmoc-D-Gln(Trt)-OH, Fmoc-L-Arg(Pbf)-OH, Fmoc-D-Phe-OH, and Boc-L-Val-OH were used with methyl 3-mercaptopropionate as the cleaving thiol. The crude peptide was purified by reverse-phase semi-preparative HPLC with mobile phases of $H_2O$ + 0.05% TFA (A) and acetonitrile + 0.05% TFA (B). The peptide was eluted using a gradient of 0-55% mobile phase B over 25 minutes at a 20 mL/min flow rate. The column was equilibrated with 0% mobile phase B for 1 minute before and 5 minutes after the gradient, yielding an off-white solid (17 mg, 19% yield).

**$^1$H NMR** (800 MHz, DMSO) δ 8.75 (d, *J* = 7.9 Hz, 1H), 8.59 (d, *J* = 7.9 Hz, 1H), 8.51 (d, *J* = 7.5 Hz, 1H), 7.99 (s, 3H), 7.78 (t, *J* = 5.6 Hz, 1H), 7.56-6.87 (m, 10H), 4.85 (dd, *J* = 9.5, 13.1 Hz, 1H), 4.39-4.33 (m, 2H), 3.60 (s, 4H), 3.12-2.95 (m, 5H), 2.75 (t, *J* = 12.0 Hz, 1H), 2.63-2.54 (m, 2H), 2.12 (t, *J* = 7.7 Hz, 2H), 1.98 (sextet, *J* = 6.8 Hz, 1H), 1.91-1.73 (m, 2H), 1.68 (heptet, *J* = 5.6 Hz, 1H), 1.59-1.49 (m, 1H), 1.44-1.30 (m, 2H), 0.70 (d, *J* = 7.0 Hz, 3H), 0.47 (d, *J* = 7.0 Hz, 3H).

**$^{13}$C NMR** (201 MHz, DMSO) δ 200.72, 174.02, 172.14, 172.03, 171.17, 168.10, 157.25, 137.52, 129.64, 128.42, 126.80, 59.06, 57.52, 54.35, 52.34, 52.00, 40.71, 38.96, 33.82, 31.20, 27.43, 25.27, 23.67, 18.60, 16.97.

**Mass spec:** expected neutral mass for $C_{29}H_{46}N_8O_7S$ (Da): 650.3210, observed neutral mass (Da): 650.3221, mass error (ppm): 1.7.

**UPLC Trace:** Obtained with mobile phases of $H_2O$ + 0.1% formic acid (A) and acetonitrile + 0.1% formic acid (B). The peptide was eluted using a gradient of 0-70% mobile phase B over 4 min at a 0.5 mL/min flow rate. Mobile phase B was held at 70% for 1 min. The column was equilibrated with 0% mobile phase for 1 minute before and 2 minutes after the gradient. The peptide purity was determined to be 98%.

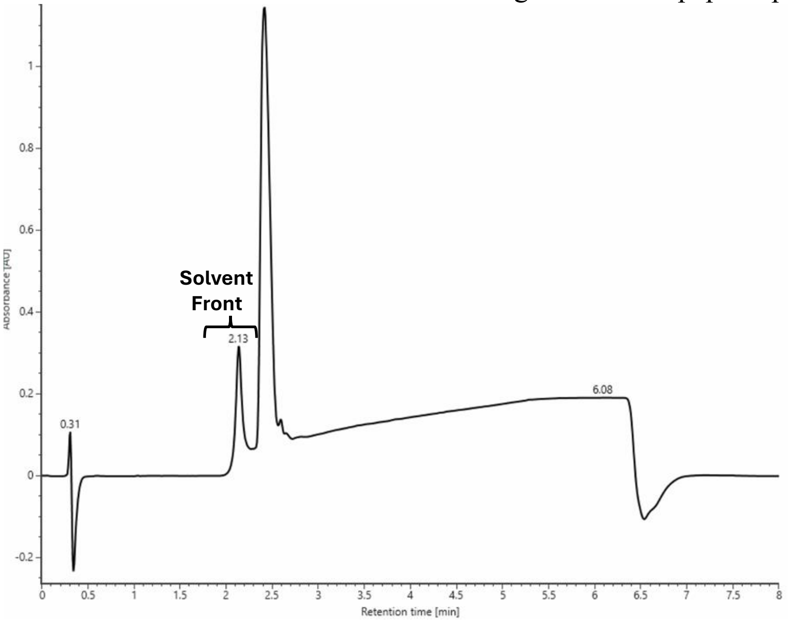

SMMP-dS-R-dF-V

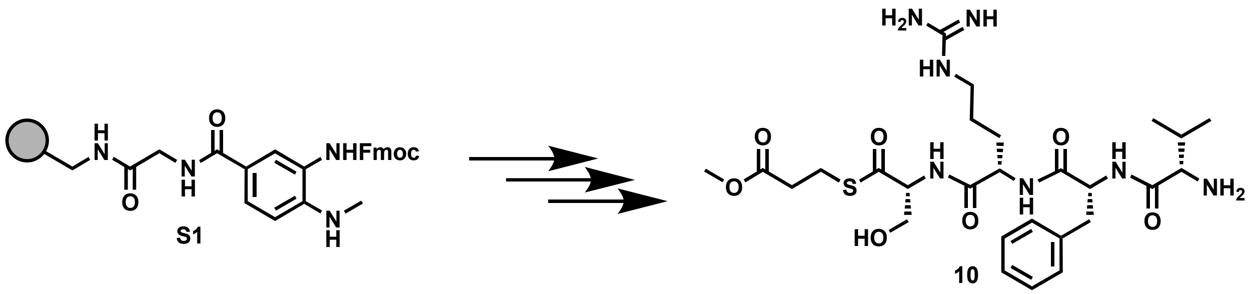

Peptide **10** was synthesized following the peptide synthesis protocol outlined above. Starting with Fmoc-Me-Dbz-OH loaded resin **S1** (0.1 mmol), amino acids Fmoc-D-Ser(tBu)-OH, Fmoc-L-Arg(Pbf)-OH, Fmoc-D-Phe-OH, and Boc-L-Val-OH were used with methyl 3-mercaptopropionate as the cleaving thiol. The crude peptide was purified by reverse-phase semi-preparative HPLC with mobile phases of $H_2O$ + 0.05% TFA (A) and acetonitrile + 0.05% TFA (B). The peptide was eluted using a gradient of 0-55% mobile phase B over 25 minutes at a 20 mL/min flow rate. The column was equilibrated with 0% mobile phase B for 1 minute before and 5 minutes after the gradient, yielding an off-white solid (10 mg, 12% yield).
**$^1$H NMR** (800 MHz, DMSO) δ 8.89-8.35 (m, 3H), 8.13-7.65 (m, 4H), 7.61-6.85 (m, 9H), 5.19 (s, 1H), 4.91 (s, 1H), 4.46 (m, 2H), 3.78-3.28 (m, 8H), 2.73 (t, *J* = 7.5 Hz, 1H), 2.58 (s, 2H), 1.83 (s, 1H), 1.66 (s, 1H), 1.52 (s, 1H), 1.39 (s, 2H), 0.77-0.54 (m, 3H), 0.52-0.30 (m, 3H).
**$^{13}$C NMR** (201 MHz, DMSO) δ 199.38, 172.25, 172.05, 171.03, 168.00, 157.25, 137.62, 129.65, 128.39, 126.76, 61.99, 61.73, 57.56, 54.11, 52.12, 52.00, 40.75, 39.17, 33.80, 30.33, 30.00, 25.23, 23.74, 18.64, 16.86.
**Mass spec:** expected neutral mass for $C_{27}H_{43}N_7O_7S$ (Da): 609.2945, observed neutral mass (Da): 609.2951, mass error (ppm): 1.1.
**UPLC Trace:** Obtained with mobile phases of $H_2O$ + 0.1% formic acid (A) and acetonitrile + 0.1% formic acid (B). The peptide was eluted using a gradient of 0-70% mobile phase B over 4 min at a 0.5 mL/min flow rate. Mobile phase B was held at 70% for 1 min. The column was equilibrated with 0% mobile phase for 1 minute before and 2 minutes after the gradient. The peptide purity was determined to be 96%.

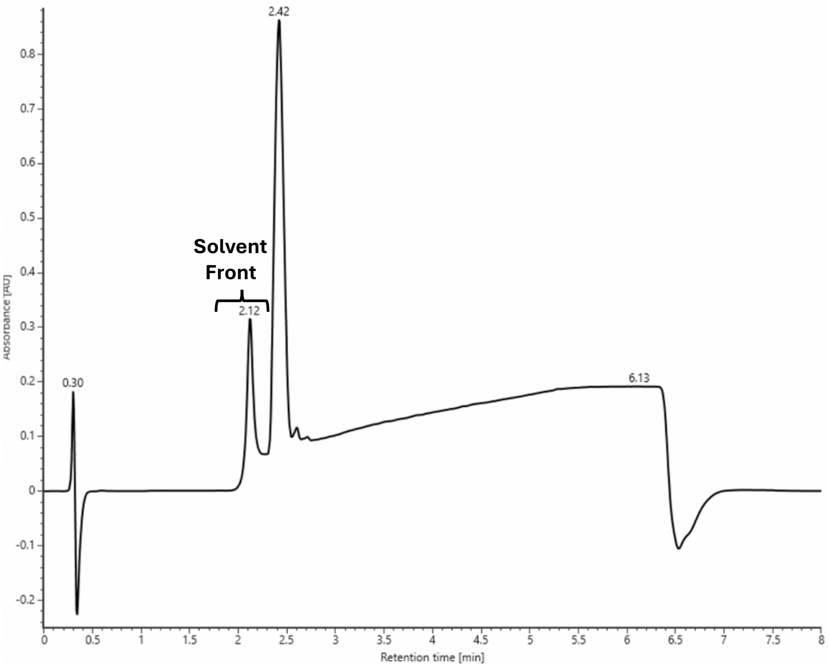

SMMP-dK-R-dF-V

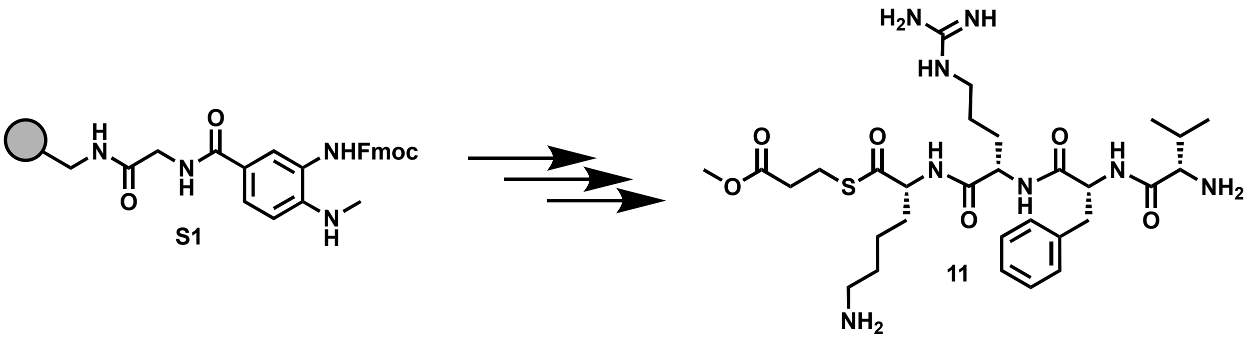

Peptide **11** was synthesized following the peptide synthesis protocol outlined above. Starting with Fmoc-Me-Dbz-OH loaded resin **S1** (0.1 mmol), amino acids Fmoc-D-Lys(Boc)-OH, Fmoc-L-Arg(Pbf)-OH, Fmoc-D-Phe-OH, and Boc-L-Val-OH were used with methyl 3-mercaptopropionate as the cleaving thiol. The crude peptide was purified by reverse-phase semi-preparative HPLC with mobile phases of $H_2O$ + 0.05% TFA (A) and acetonitrile + 0.05% TFA (B). The peptide was eluted using a gradient of 0-55% mobile phase B over 25 minutes at a 20 mL/min flow rate. The column was equilibrated with 0% mobile phase B for 1 minute before and 5 minutes after the gradient, yielding an off-white solid (13 mg, 13% yield).

**$^1$H NMR** (800 MHz, DMSO) δ 8.72 (dd, *J* = 7.9, 14.4 Hz, 2H), 8.56 (d, *J* = 7.5 Hz, 1H), 8.05-7.71 (m, 7H), 7.62-6.92 (m, 8H), 4.93 (t, 10.4 Hz, 1H), 4.46 (dd, *J* = 8.1, 14.0 Hz, 1H), 4.31 (dd, *J* = 8.4, 12.7 Hz, 1H), 3.60 (s, 4H), 3.12-2.95 (m, 5H), 2.82-2.68 (m, 3H), 2.64-2.53 (m, 2H), 1.82 (5.5, 10.9 Hz, 1H), 1.73 (dd, *J* = 8.4, 13.8 Hz, 1H), 1.69-1.47 (m, 5H), 1.46-1.20 (m, 4H), 0.69 (d, *J* = 6.8 Hz, 3H), 0.41 (d, *J* = 6.8 Hz, 3H).

**$^{13}$C NMR** (201 MHz, DMSO) δ $^{13}$C NMR (201 MHz, DMSO) δ 206.91, 200.88, 172.14, 172.02, 171.01, 167.93, 157.26, 137.63, 129.65, 128.39, 126.77, 70.17, 59.29, 57.58, 53.99, 52.01, 40.69, 38.82, 33.82, 31.07, 30.44, 30.00, 26.80, 25.30, 23.70, 22.48, 18.67, 16.78.

**Mass spec:** expected neutral mass for $C_{30}H_{50}N_8O_6S$ (Da): 650.3574, observed neutral mass (Da): 650.3585, mass error (ppm): 1.7.

**UPLC Trace:** Obtained with mobile phases of $H_2O$ + 0.1% formic acid (A) and acetonitrile + 0.1% formic acid (B). The peptide was eluted using a gradient of 0-70% mobile phase B over 4 min at a 0.5 mL/min flow rate. Mobile phase B was held at 70% for 1 min. The column was equilibrated with 0% mobile phase for 1 minute before and 2 minutes after the gradient. peptide purity was determined to be 98%.

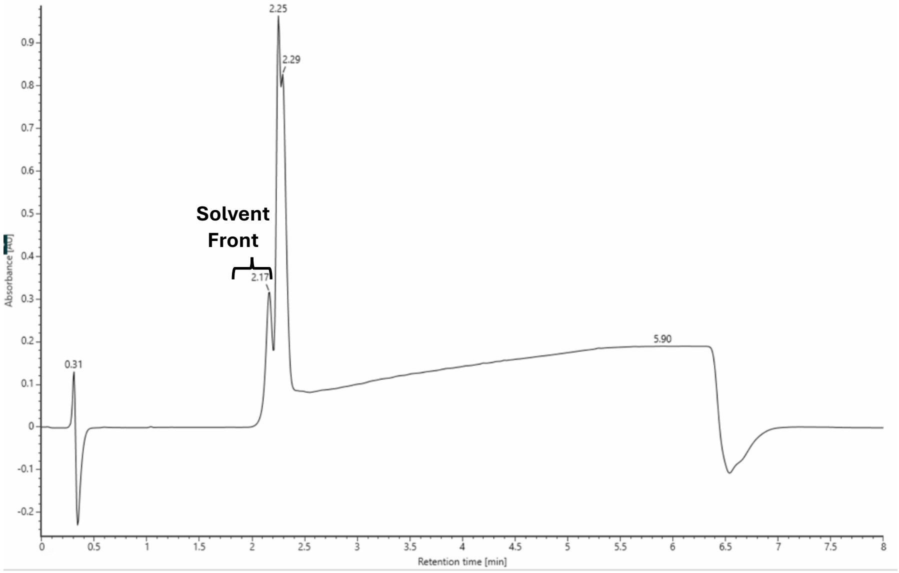

SBMP-dF-R-dF-V

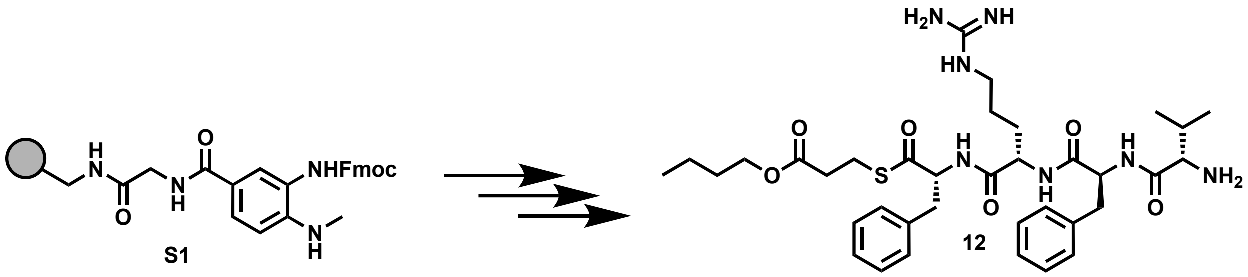

Peptide **12** was synthesized following the peptide synthesis protocol outlined above. Starting with Fmoc-Me-Dbz-OH loaded resin **S1** (0.1 mmol), amino acids Fmoc-D-Phe-OH, Fmoc-D-Dap(Boc)-OH, and Boc-L-Val-OH were used with butyl 3-mercaptopropionate as the cleaving thiol. The crude peptide was purified by reverse-phase semi-preparative HPLC with mobile phases of $H_2O$ + 0.05% TFA (A) and acetonitrile + 0.05% TFA (B). The peptide was eluted using a gradient of 0-55% mobile phase B over 25 minutes at a 20 mL/min flow rate. The column was equilibrated with 0% mobile phase B for 1 minute before and 5 minutes after the gradient, yielding an off-white solid (40 mg, 42% yield).

**$^1$H NMR** (800 MHz, DMSO) δ 8.78 (d, *J* = 7.6 Hz, 1H), 8.71 (d, *J* = 8.0 Hz, 1H), 8.43 (d, *J* = 6.7 Hz, 1H), 7.95 (s, 3H), 7.75-7.72 (m, 1H), 7.57-7.17 (m, 13H), 7.15 (q, *J* = 6.9 Hz, 2H), 4.90 (sextet, *J* = 4.9 Hz, 1H), 4.62 (sextet, *J* = 4.7 Hz, 1H), 4.39 (q, *J* = 6.4 Hz, 1H), 4.03 (t, *J* = 6.3 Hz, 2H), 3.14 (dd, *J* = 4.4, 13.3 Hz, 1H), 3.05-2.97 (m, 3H), 2.90 (octet, *J* = 6.8 Hz, 2H), 2.78 (t, *J* = 10.5 Hz, 1H), 2.69 (t, *J* = 10.9 Hz, 1H), 2.62-2.53 (m, 2H), 1.82 (dt, *J* = 11, 6.4 Hz, 1H), 1.55 (quintet, *J* = 6.8 Hz, 2H), 1.32 (quintet, *J* = 6.8 Hz, 3H), 1.26-1.18 (m, 1H), 1.09 (s, 2H), 0.89 (m, 3H), 0.68 (d, *J* = 6.7, 3H), 0.42 (d, *J* = 7 Hz, 3H)

**$^{13}$C NMR** (201 MHz, DMSO) δ 200.29, 171.82, 171.59, 170.88, 167.91, 157.25, 137.63, 137.20, 129.64, 129.58, 128.60, 128.36, 127.00, 126.73, 64.36, 60.78, 57.55, 53.93, 51.91, 40.69, 39.30, 37.43, 33.93, 30.52, 30.09, 29.99, 24.79, 23.87, 18.98, 18.65, 16.81, 13.93.

**Mass spec:** expected neutral mass for $C_{36}H_{53}N_7O_6S$ (Da): 711.3778, observed neutral mass (Da): 711.3748, mass error (ppm): -4.2.

**UPLC Trace:** Obtained with mobile phases of $H_2O$ + 0.1% formic acid (A) and acetonitrile + 0.1% formic acid (B). The peptide was eluted using a gradient of 0-70% mobile phase B over 4 min at a 0.5 mL/min flow rate. Mobile phase B was held at 70% for 1 min. The column was equilibrated with 0% mobile phase for 1 minute before and 2 minutes after the gradient. The peptide purity was determined to be 97%.

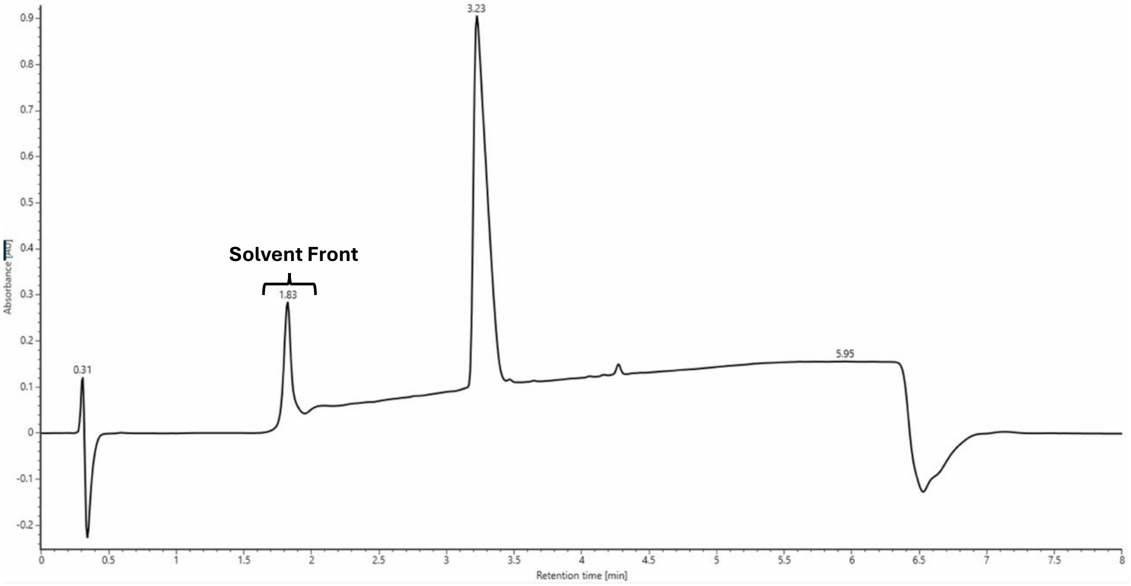

SMMP-dV-R-dF-V

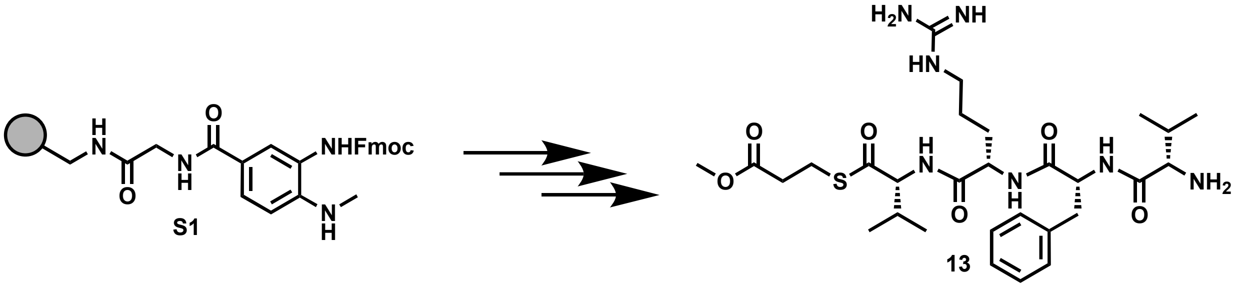

Peptide **13** was synthesized following the peptide synthesis protocol outlined above. Starting with Fmoc-Me-Dbz-OH loaded resin **S1** (0.1 mmol), amino acids Fmoc-D-Val-OH, Fmoc-L-Arg(Pbf)-OH, Fmoc-D-Phe-OH, and Boc-L-Val-OH were used with methyl 3-mercaptopropionate as the cleaving thiol. The crude peptide was purified by reverse-phase semi-preparative HPLC with mobile phases of $H_2O$ + 0.05% TFA (A) and acetonitrile + 0.05% TFA (B). The peptide was eluted using a gradient of 0-55% mobile phase B over 25 minutes at a 20 mL/min flow rate. The column was equilibrated with 0% mobile phase B for 1 minute before and 5 minutes after the gradient, yielding an off-white solid (23 mg, 27% yield).

**$^1$H NMR** (800 MHz, DMSO) δ 8.72 (d, $J$ = 9.4 Hz, 1H), 8.56 (dd, $J$ = 8.4, 16.1 Hz, 2H), 8.03-7.85 (m, 4H), 7.68-6.94 (m, 8H), 4.94 (dt, $J$ = 4.5, 9.7 Hz, 1H), 4.60 (dd, $J$ = 8.4, 13.4 Hz, 1H), 4.30 (t, $J$ = 7.2 Hz, 1H), 3.60 (s, 4H), 3.13-2.97 (m, 5H), 2.72 (t, $J$ = 12.3 Hz, 1H), 2.63-2.53 (m, 2H), 2.13 (sextet, $J$ = 6.5 Hz, 1H), 1.82 (sextet, $J$ = 6.5 Hz, 1H), 1.70-1.60 (m, 1H), 1.58-1.49 (m, 1H), 1.46-1.34 (m, 2H), 0.87 (d, $J$ = 5.8 Hz, 6H), 0.67 (d, $J$ = 7.1 Hz, 3H), 0.42 (d, $J$ = 7.3 Hz,3H).

**$^{13}$C NMR** (201 MHz, DMSO) δ 200.31, 172.43, 171.99, 170.98, 167.90, 157.28, 137.66, 129.66, 128.37, 126.74, 64.41, 57.55, 51.99, 40.70, 39.38, 33.83, 29.99, 25.31, 23.80, 19.49, 18.65, 17.86, 16.81.

**Mass spec:** expected neutral mass for $C_{29}H_{47}N_7O_6S$ (Da): 621.3309, observed neutral mass (Da): 621.3306, mass error (ppm): -0.4.

**UPLC Trace:** Obtained with mobile phases of $H_2O$ + 0.1% formic acid (A) and acetonitrile + 0.1% formic acid (B). The peptide was eluted using a gradient of 0-70% mobile phase B over 4 min at a 0.5 mL/min flow rate. Mobile phase B was held at 70% for 1 min. The column was equilibrated with 0% mobile phase for 1 minute before and 2 minutes after the gradient. The peptide purity was determined to be 99%.

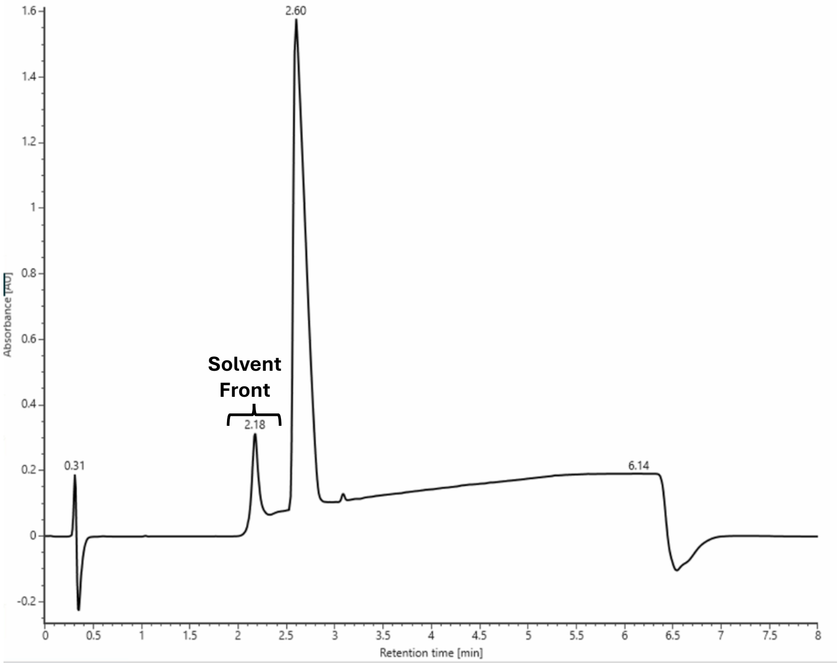

SMMP-dE-R-dF-V

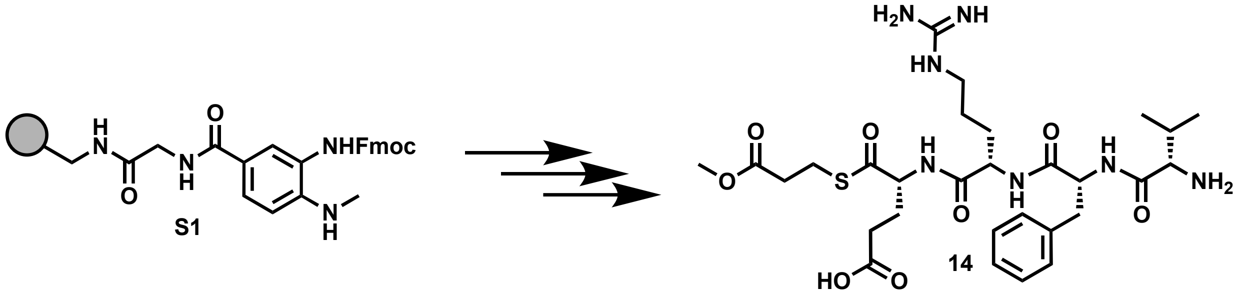

Peptide **14** was synthesized following the peptide synthesis protocol outlined above. Starting with Fmoc-Me-Dbz-OH loaded resin **S1** (0.1 mmol), amino acids Fmoc-D-Glu(tBu)-OH, Fmoc-L-Arg(Pbf)-OH, Fmoc-D-Phe-OH, and Boc-L-Val-OH were used with methyl 3-mercaptopropionate as the cleaving thiol. The crude peptide was purified by reverse-phase semi-preparative HPLC with mobile phases of $H_2O$ + 0.05% TFA (A) and acetonitrile + 0.05% TFA (B). The peptide was eluted using a gradient of 0-55% mobile phase B over 25 minutes at a 20 mL/min flow rate. The column was equilibrated with 0% mobile phase B for 1 minute before and 5 minutes after the gradient, yielding an off-white solid 38 mg, 43% yield).

**$^1$H NMR** (800 MHz, DMSO) δ 8.77-8.68 (m, 2H), 8.55 (d, $J$ = 8.2 Hz, 1H), 8.15-7.73 (m, 3H), 7.61-6.94 (m, 8H), 4.93 (sextet, $J$ = 4.4 Hz, 1H), 4.48-4.36 (m, 2H), 3.44-3.29 (m, 5H), 3.13-2.96 (m, 5H), 2.73 (t, $J$ = 12.2 Hz, 1H), 2.60 (dq, $J$ = 6.9, 16.9 Hz, 2H), 2.29 (dd, $J$ = 6.1, 13.1 Hz, 2H), 2.01 (sextet, $J$ = 7.1 Hz, 1H), 1.84 (dt, $J$ = 6.7, 12.2 Hz, 1H), 1.78 (dt, $J$ = 8.0, 14.1 Hz, 1H), 1.71-1.62 (m, 1H), 1.58-1.49 (m, 1H), 1.48-1.33 (m, 2H), 0.70 (d, $J$ = 6.6 Hz, 3H), 0.44 (d, $J$ = 7.1 Hz, 3H).

**$^{13}$C NMR** (201 MHz, DMSO) δ 200.69, 174.03, 172.23, 172.11, 171.10, 168.03, 157.32, 137.70, 129.73, 128.47, 126.84, 58.77, 57.65, 54.11, 52.18, 52.07, 40.80, 39.37, 33.88, 30.43, 30.10, 30.08, 27.04, 25.36, 23.79, 18.74, 16.90.

**Mass spec:** expected neutral mass for $C_{32}H_{51}N_7O_8S$ (Da): 651.3050, observed neutral mass (Da): 651.3071, mass error (ppm): 3.4.

**UPLC Trace:** Obtained with mobile phases of $H_2O$ + 0.1% formic acid (A) and acetonitrile + 0.1% formic acid (B). The peptide was eluted using a gradient of 0-70% mobile phase B over 4 min at a 0.5 mL/min flow rate. Mobile phase B was held at 70% for 1 min. The column was equilibrated with 0% mobile phase for 1 minute before and 2 minutes after the gradient. The peptide purity was determined to be 95%.

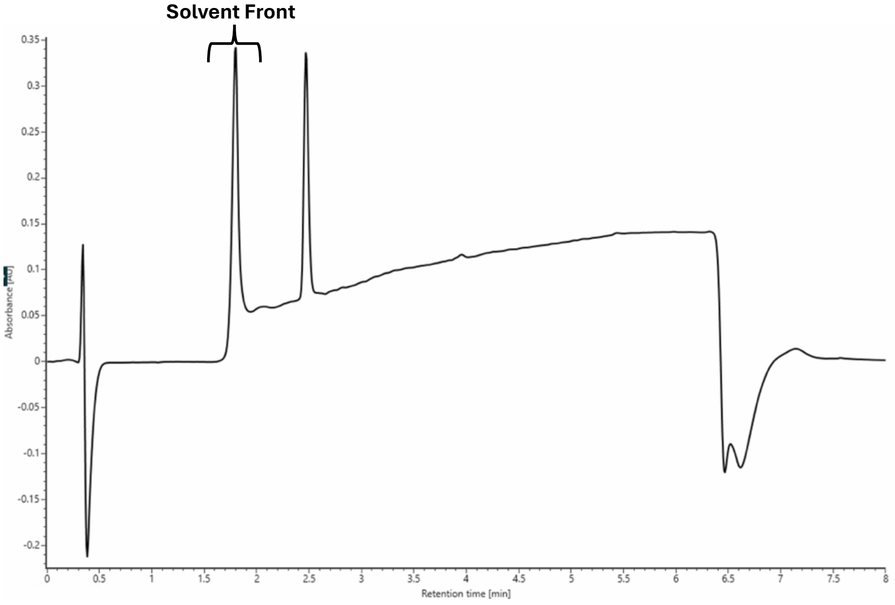

SMMP-G-R-dF-V

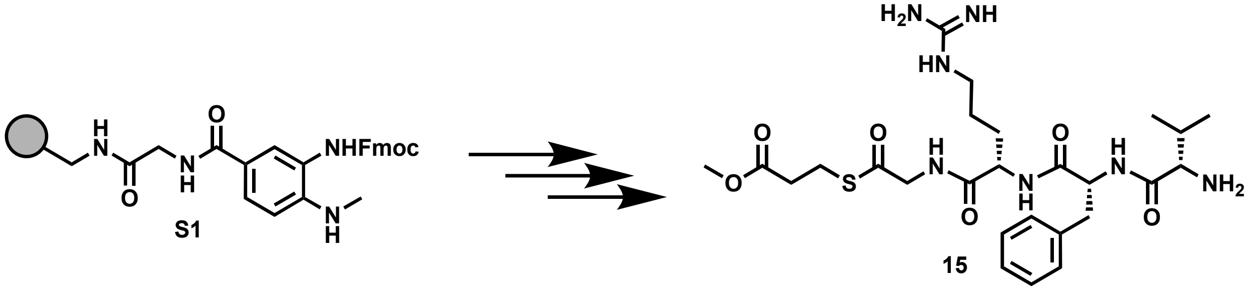

Peptide **15** was synthesized following the peptide synthesis protocol outlined above. Starting with Fmoc-Me-Dbz-OH loaded resin **S1** (0.1 mmol), amino acids Fmoc-Gly-OH, Fmoc-L-Arg(Pbf)-OH, Fmoc-D-Phe-OH, and Boc-L-Val-OH were used with methyl 3-mercaptopropionate as the cleaving thiol. The crude peptide was purified by reverse-phase semi-preparative HPLC with mobile phases of $H_2O$ + 0.05% TFA (A) and acetonitrile + 0.05% TFA (B). The peptide was eluted using a gradient of 0-55% mobile phase B over 25 minutes at a 20 mL/min flow rate. The column was equilibrated with 0% mobile phase B for 1 minute before and 5 minutes after the gradient, yielding an off-white solid (30 mg, 37% yield).

**$^1$H NMR** (800 MHz, DMSO) δ 8.75 (t, $J$ = 4.2 Hz, 2H), 8.60 (d, $J$ = 7.6 Hz, 1H), 7.97 (s, 3H), 7.85 (t, $J$ = 5.2 Hz), 7.55-7.05 (m, 8H), 4.87 (dt, $J$ = 4.1, 9.1 Hz, 1H), 4.38 (dd, $J$ = 7.1, 13.2 Hz, 1H), 4.08 (dd, $J$ = 5.4, 17.1 Hz, 1H), 4.00 (dd, $J$ = 5.5, 17.1 Hz, 1H), 3.63-3.59 (m, 4H), 3.13-2.98 (m, 5H), 2.75 (t, $J$ = 11.9 Hz, 1H), 2.59 (t, $J$ = 6.7 Hz, 2H), 1.84 (sextet, $J$ = 6.5, 1H), 1.72 (septet, $J$ = 5.5 Hz, 1H), 1.59-1.50 (m, 1H), 1.47-1.37 (m, 2H), 0.70 (d, $J$ = 6.9 Hz, 3H), 0.47 (d, $J$ = 6.9 Hz, 3H).

**$^{13}$C NMR** (201 MHz, DMSO) δ 198.21, 172.52, 172.06, 171.19, 168.11, 157.37, 137.68, 129.70, 128.48, 126.85, 57.61, 54.31, 52.24, 52.05, 49.23, 40.76, 39.07, 33.91, 30.09, 29.76, 25.37, 23.55, 18.69, 17.03.

**Mass spec:** expected neutral mass for $C_{26}H_{41}N_7O_6S$ (Da): 579.2839, observed neutral mass (Da): 579.2855, mass error (ppm): 2.7.

**UPLC Trace** Obtained with mobile phases of $H_2O$ + 0.1% formic acid (A) and acetonitrile + 0.1% formic acid (B). The peptide was eluted using a gradient of 0-70% mobile phase B over 4 min at a 0.5 mL/min flow rate. Mobile phase B was held at 70% for 1 min. The column was equilibrated with 0% mobile phase for 1 minute before and 2 minutes after the gradient. The peptide purity was determined to be 97%.

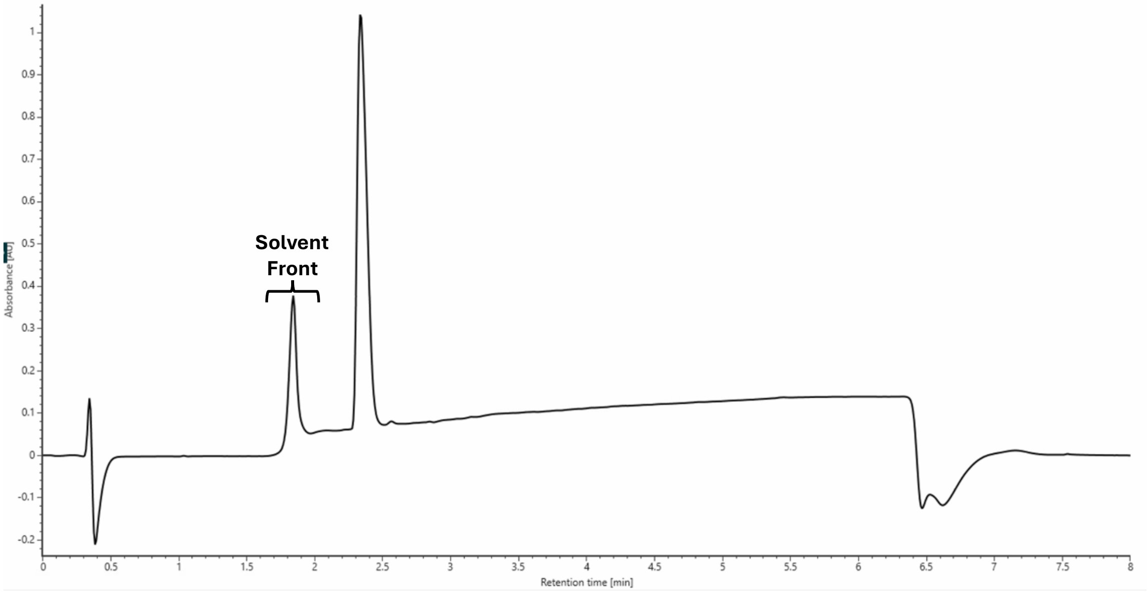

SMMP-G-R-F-V

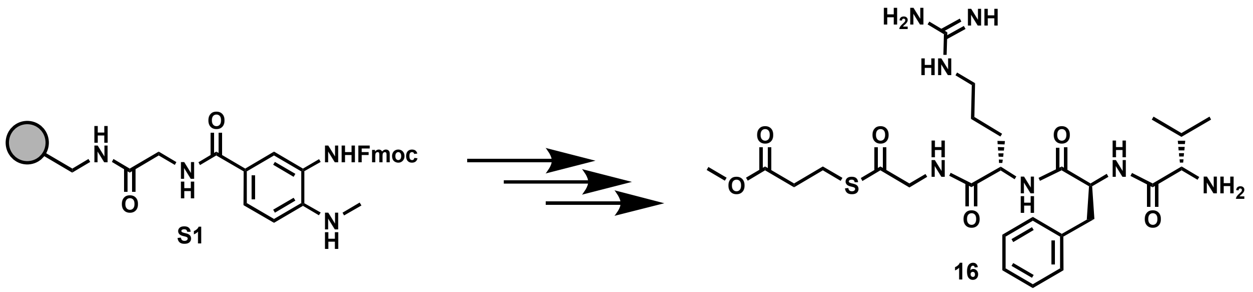

Peptide **16** was synthesized following the peptide synthesis protocol outlined above. Starting with Fmoc-Me-Dbz-OH loaded resin **S1** (0.1 mmol), amino acids Fmoc-Gly-OH, Fmoc-L-Arg(Pbf)-OH, Fmoc-L-Phe-OH, and Boc-L-Val-OH were used with methyl 3-mercaptopropionate as the cleaving thiol. The crude peptide was purified by reverse-phase semi-preparative HPLC with mobile phases of $H_2O$ + 0.05% TFA (A) and acetonitrile + 0.05% TFA (B). The peptide was eluted using a gradient of 0-55% mobile phase B over 25 minutes at a 20 mL/min flow rate. The column was equilibrated with 0% mobile phase B for 1 minute before and 5 minutes after the gradient, yielding an off-white solid (20 mg, 25% yield).
**$^{1}$H NMR** (800 MHz, DMSO) δ 8.66-8.51 (m, 2H), 8.41 (s, 1H), 8.05 (s, 3H), 7.85 (s, 1H), 7.69-7.01 (m, 9H), 4.71-4.61 (m, 1H), 4.36-4.28 (m, 1H), 4.08-3.94 (m, 2H), 3.66-3.61 (m, 1H), 3.54-3.26 (m, 2H), 3.17-2.97 (m, 5H), 2.83 (t, *J* = 11.1 Hz, 1H), 2.61-2.55 (m, 2H), 2.15-2.06 (m, 1H), 1.79-1.70 (m, 1H), 1.63-1.44 (m, 3H), 0.98-0.83 (m, 6H).
**$^{13}$C NMR** (201 MHz, DMSO) δ 198.34, 172.37, 172.04, 171.17, 168.44, 157.38, 137.94, 129.65, 128.60, 126.84, 57.46, 54.53, 52.64, 52.02, 49.25, 40.78, 37.73, 33.92, 30.39, 29.41, 25.48, 23.52, 18.80, 17.60.
**Mass spec:** expected neutral mass for $C_{26}H_{41}N_7O_6S$ (Da): 579.2839, observed neutral mass (Da): 579.2853, mass error (ppm): 2.4.
**UPLC Trace:** Obtained with mobile phases of $H_2O$ + 0.1% formic acid (A) and acetonitrile + 0.1% formic acid (B). The peptide was eluted using a gradient of 0-70% mobile phase B over 4 min at a 0.5 mL/min flow rate. Mobile phase B was held at 70% for 1 min. The column was equilibrated with 0% mobile phase for 1 minute before and 2 minutes after the gradient. The peptide purity was determined to be 99%.

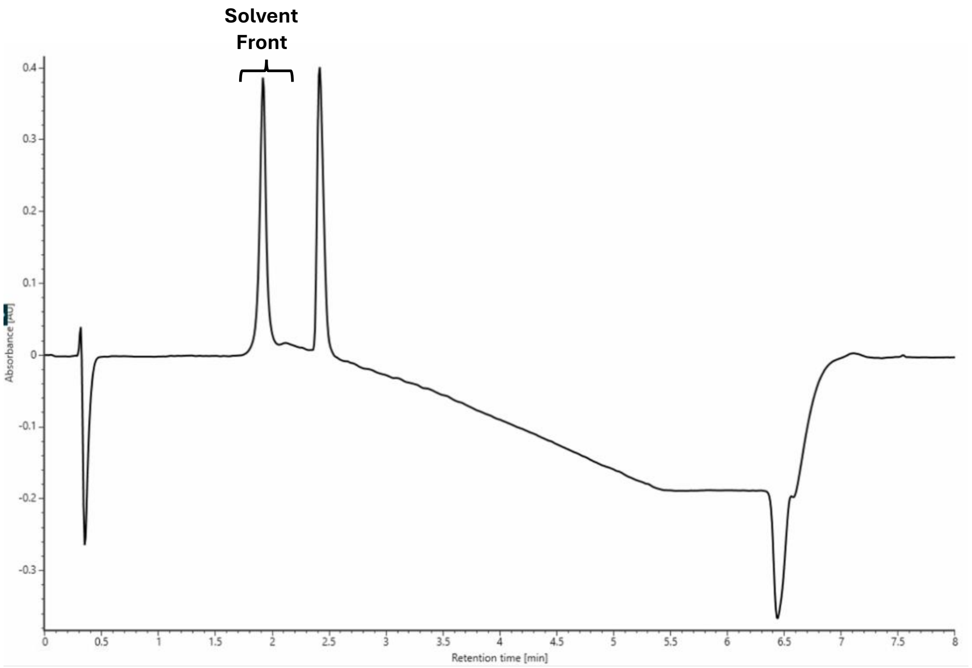

SMMP-dY-R-dF-P

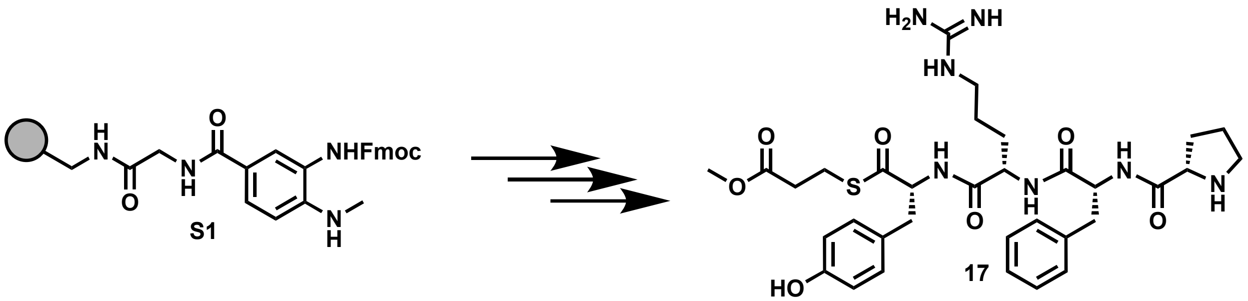

Peptide **17** was synthesized following the peptide synthesis protocol outlined above. Starting with Fmoc-Me-Dbz-OH loaded resin **S1** (0.1 mmol), amino acids Fmoc-D-Tyr(tBu)-OH, Fmoc-L-Arg(Pbf)-OH, Fmoc-D-Phe-OH, and Boc-L-Pro-OH were used with methyl 3-mercaptopropionate as the cleaving thiol. The crude peptide was purified by reverse-phase semi-preparative HPLC with mobile phases of $H_2O$ + 0.05% TFA (A) and acetonitrile + 0.05% TFA (B). The peptide was eluted using a gradient of 0-55% mobile phase B over 25 minutes at a 20 mL/min flow rate. The column was equilibrated with 0% mobile phase B for 1 minute before and 5 minutes after the gradient, yielding an off-white solid (27 mg, 30% yield).

**$^1$H NMR** (800 MHz, DMSO) δ 9.33 (s, 2H), 8.81 (d, *J* = 8.9 Hz, 1H), 8.69 (d, *J* = 8.3 Hz, 1H), 8.49 (d, *J* = 8.3 Hz, 1H), 8.34 (s, 1H), 7.74 (t, *J* = 4.8 Hz, 1H), 7.58-6.99 (m, 10H), 6.67 (d, *J* = 8.2 Hz, 2H), 4.85 (dt, *J* = 4.1, 9.9 Hz, 1H), 4.51 (dt, *J* = 5.7, 8.9 Hz, 1H), 4.41 (dd, *J* = 7.5, 13.5 Hz, 1H), 4.11 (t, *J* = 6.7 Hz, 1H), 3.56-3.28 (m, 3H), 3.15-2.93 (m, 8H), 2.71 (q, *J* = 12.1 Hz, 2H), 2.58 (dd, *J* = 6.5, 9.9 Hz, 2H), 2.07 (dt, *J* = 7.1, 13.2 Hz, 1H), 1.73 (septet, *J* = 6.7 Hz, 1H), 1.53 (septet, *J* = 6.7 Hz, 1H), 1.45 (sextet, *J* = 6.6 Hz, 1H), 1.36-1.16 (m, 4H).

**$^{13}$C NMR** (201 MHz, DMSO) δ 200.51, 172.10, 171.86, 170.78, 168.03, 157.33, 156.57, 137.75, 130.62, 129.83, 128.38, 127.23, 126.81, 115.51, 61.27, 59.45, 54.01, 52.07, 45.96, 40.71, 39.29, 36.75, 33.84, 30.09, 30.04, 25.09, 23.86, 23.57.

**Mass spec:** expected neutral mass for $C_{33}H_{45}N_7O_7S$ (Da): 683.3102, observed neutral mass (Da): 683.3118, mass error (ppm): 2.5.

**UPLC Trace** Obtained with mobile phases of $H_2O$ + 0.1% formic acid (A) and acetonitrile + 0.1% formic acid (B). The peptide was eluted using a gradient of 0-70% mobile phase B over 4 min at a 0.5 mL/min flow rate. Mobile phase B was held at 70% for 1 min. The column was equilibrated with 0% mobile phase for 1 minute before and 2 minutes after the gradient. The peptide purity was determined to be 97%.

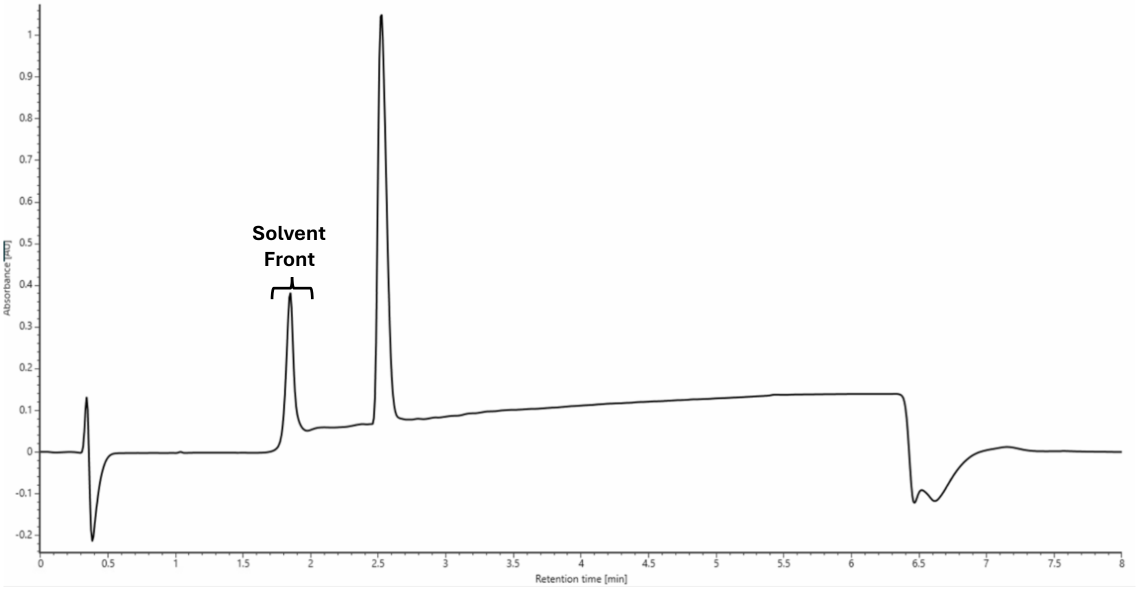

SMMP-dY-R-dF-H

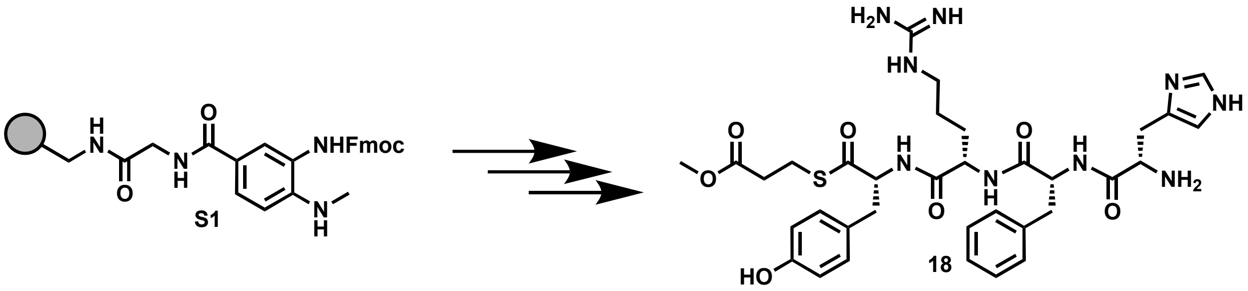

Peptide **18** was synthesized following the peptide synthesis protocol outlined above. Starting with Fmoc-Me-Dbz-OH loaded resin **S1** (0.1 mmol), amino acids Fmoc-D-Tyr(tBu)-OH, Fmoc-L-Arg(Pbf)-OH, Fmoc-D-Phe-OH, and Boc-L-His(Trt)-OH were used with methyl 3-mercaptopropionate as the cleaving thiol. The crude peptide was purified by reverse-phase semi-preparative HPLC with mobile phases of $H_2O$ + 0.05% TFA (A) and acetonitrile + 0.05% TFA (B). The peptide was eluted using a gradient of 0-55% mobile phase B over 25 minutes at a 20 mL/min flow rate. The column was equilibrated with 0% mobile phase B for 1 minute before and 5 minutes after the gradient, yielding an off-white solid (5 mg, 5% yield).
**Mass spec:** expected neutral mass for $C_{34}H_{45}N_9O_7S$ (Da): 723.3163, observed neutral mass (Da): 723.3194, mass error (ppm): 4.3.
**UPLC Trace** Obtained with mobile phases of $H_2O$ + 0.1% formic acid (A) and acetonitrile + 0.1% formic acid (B). The peptide was eluted using a gradient of 0-70% mobile phase B over 4 min at a 0.5 mL/min flow rate. Mobile phase B was held at 70% for 1 min. The column was equilibrated with 0% mobile phase for 1 minute before and 2 minutes after the gradient. The peptide purity was determined to be 95%.

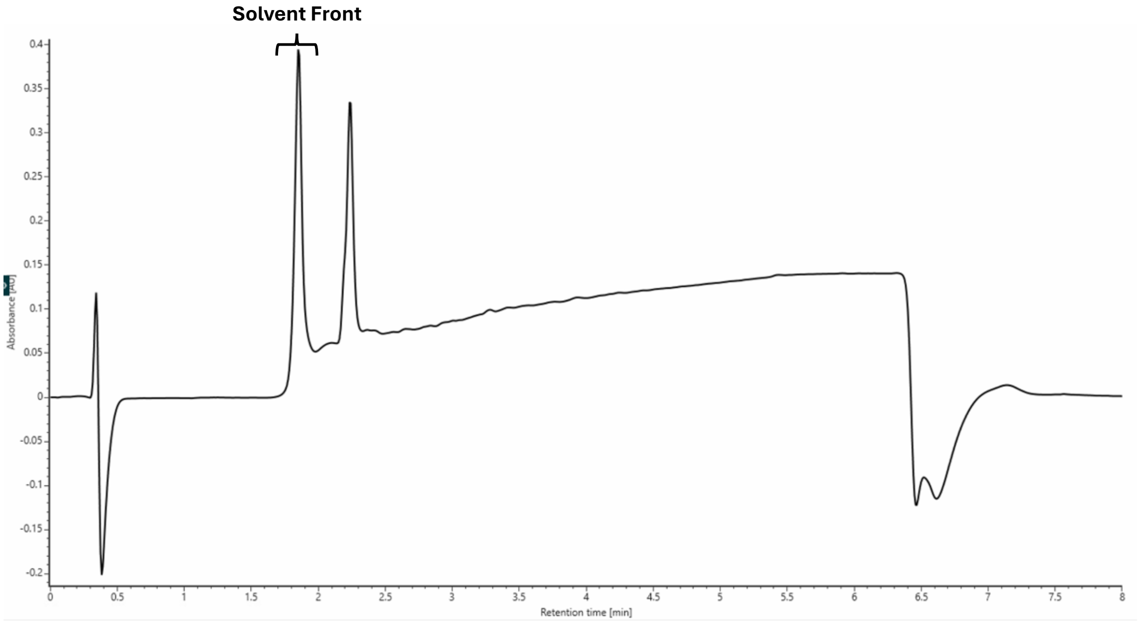

SMMP-dY-R-dF-E

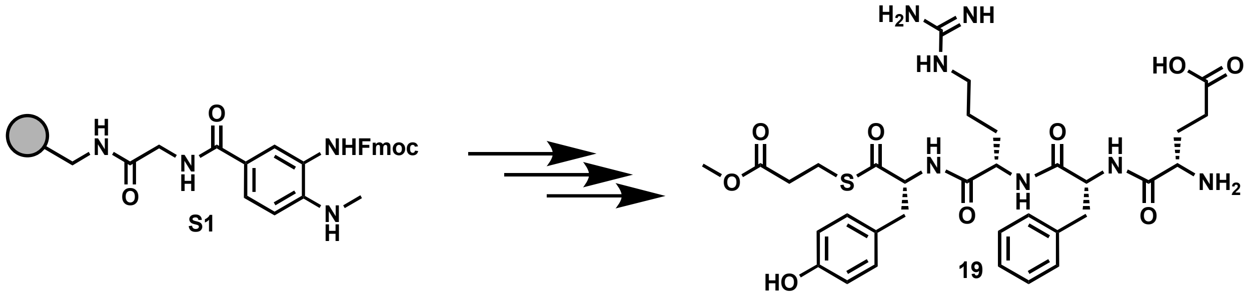

Peptide **19** was synthesized following the peptide synthesis protocol outlined above. Starting with Fmoc-Me-Dbz-OH loaded resin **S1** (0.1 mmol), amino acids Fmoc-D-Tyr(tBu)-OH, Fmoc-L-Arg(Pbf)-OH, Fmoc-D-Phe-OH, and Boc-L-Glu(tBu)-OH were used with methyl 3-mercaptopropionate as the cleaving thiol. The crude peptide was purified by reverse-phase semi-preparative HPLC with mobile phases of $H_2O$ + 0.05% TFA (A) and acetonitrile + 0.05% TFA (B). The peptide was eluted using a gradient of 0-55% mobile phase B over 25 minutes at a 20 mL/min flow rate. The column was equilibrated with 0% mobile phase B for 1 minute before and 5 minutes after the gradient, yielding an off-white solid (10 mg, 11% yield).
**$^1$H NMR** (800 MHz, DMSO) δ 9.30 (s, 1H), 8.79 (d, *J* = 8.8 Hz, 1H), 8.70 (d, *J* = 8.3 Hz, 1H), 8.50 (d, *J* = 8.4 Hz, 1H), 8.01 (s, 2H), 7.61 (t, *J* = 5.0 Hz, 1H), 7.49-6.85 (m, 10H), 6.67 (d, *J* = 7.9 Hz, 2H), 4.89 (dt, *J* = 4.7, 9.1 Hz, 1H), 4.52 (dt, *J* = 5.4, 9.1 Hz, 1H), 4.40 (dd, *J* = 7.8, 12.9 Hz, 1H), 3.80 s, 1H), 3.35-3.27 (m, 7H), 3.05-2.90 (m, 6H), 2.69 (t, *J* = 11.6 Hz, 2H), 2.58 (t, *J* = 5.6 Hz, 2H), 2.00-1.91 (m, 1H), 1.74-1.60 (m, 3H), 1.40 (dt, *J* = 7.3, 13.1 Hz, 1H), 1.32-1.14 (m, 3H).
**$^{13}$C NMR** (201 MHz, DMSO) δ 200.44, 173.66, 172.11, 171.89, 170.87, 168.06, 157.22, 156.57, 137.59, 130.64, 129.69, 128.39, 127.17, 126.92, 115.51, 70.25, 61.23, 53.93, 52.10, 51.96, 51.80, 40.73, 39.44, 36.88, 33.83, 30.16, 28.76, 26.65, 25.03, 23.87.
**Mass spec:** expected neutral mass for $C_{33}H_{45}N_7O_9S$ (Da): 715.2999, observed neutral mass (Da): 715.3014, mass error (ppm): 2.0.
**UPLC Trace** Obtained with mobile phases of $H_2O$ + 0.1% formic acid (A) and acetonitrile + 0.1% formic acid (B). The peptide was eluted using a gradient of 0-70% mobile phase B over 4 min at a 0.5 mL/min flow rate. Mobile phase B was held at 70% for 1 min. The column was equilibrated with 0% mobile phase for 1 minute before and 2 minutes after the gradient. The peptide purity was determined to be 95%.

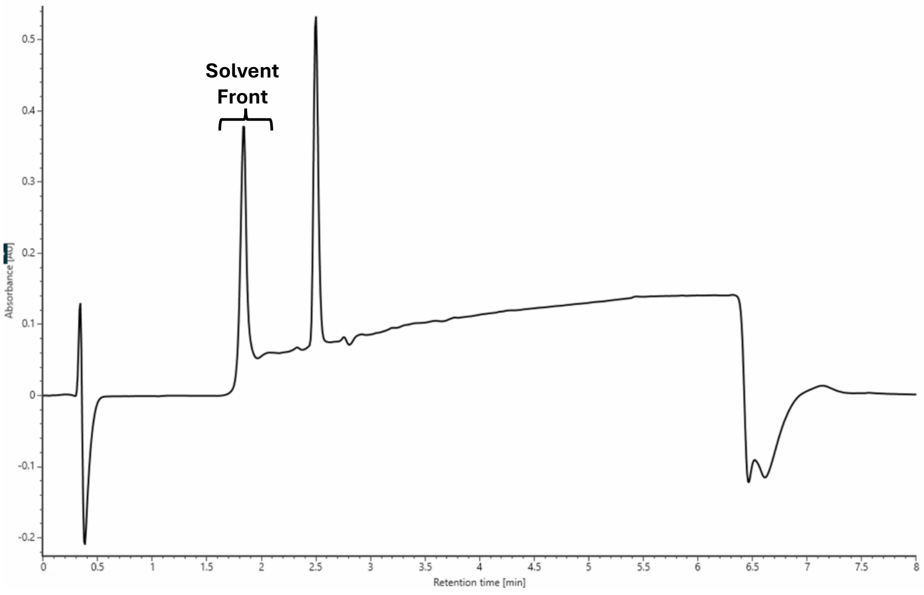

SMMP-dY-R-dF-Y

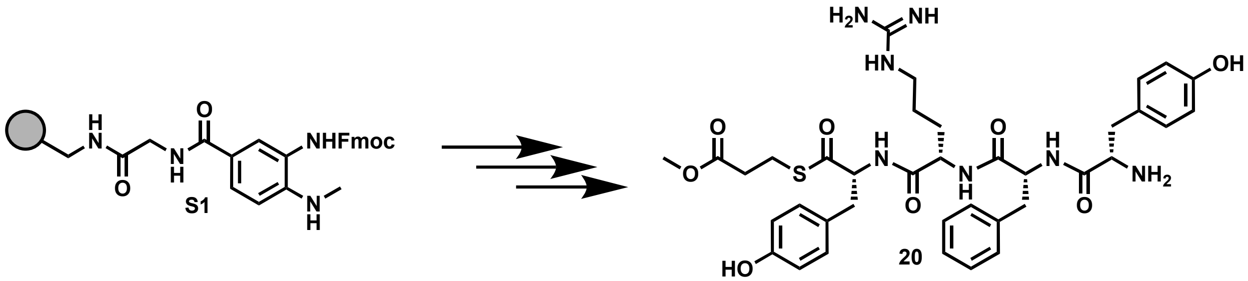

Peptide **20** was synthesized following the peptide synthesis protocol outlined above. Starting with Fmoc-Me-Dbz-OH loaded resin **S1** (0.1 mmol), amino acids Fmoc-D-Tyr(tBu)-OH, Fmoc-L-Arg(Pbf)-OH, Fmoc-D-Phe-OH, and Boc-L-Tyr(tBu)-OH were used with methyl 3-mercaptopropionate as the cleaving thiol. The crude peptide was purified by reverse-phase semi-preparative HPLC with mobile phases of $H_2O$ + 0.05% TFA (A) and acetonitrile + 0.05% TFA (B). The peptide was eluted using a gradient of 0-55% mobile phase B over 25 minutes at a 20 mL/min flow rate. The column was equilibrated with 0% mobile phase B for 1 minute before and 5 minutes after the gradient, yielding an off-white solid (33 mg, 34% yield).

**$^1$H NMR** (800 MHz, DMSO) δ 8.84 (d, *J* = 8.4 Hz, 1H), 8.68 (d, *J* = 8.4 Hz, 1H), 8.45 (d, *J* = 8.2 Hz, 1H), 7.97-7.88 (m, 3H), 7.73 (t, *J* = 5.1 Hz, 1H), 7.56-6.98 (m, 11H), 6.76 (d, *J* = 8.3 Hz, 2H), 6.67 (d, *J* = 7.7 Hz, 2H), 6.62 (d, *J* = 7.7 Hz, 2H), 4.85 (dt, *J* = 5.7, 8.6 Hz, 1H), 4.51 (dt, *J* = 4.8, 8.4 Hz, 1H), 4.37 ( dd, *J* = 7.7, 12.3 Hz, 1H), 3.96-3.89 (m, 1H), 3.03-2.90 (m, 7H), 2.69 (t, *J* = 12.4, 3H), 2.60-2.53 (m, 2H), 2.43 (dd, *J* = 7.9, 13.8 Hz, 1H), 1.40 (dt, *J* = 7.7, 12.8 Hz, 1H), 1.27 (dt, *J* = 7.7, 15.3 Hz, 1H), 1.17 (quintet, *J* = 7.6 Hz, 2H).

**$^{13}$C NMR** (201 MHz, DMSO) δ 200.46, 172.10, 171.93, 170.82, 168.21, 157.33, 157.01, 156.58, 137.70, 130.95, 130.63, 129.88, 128.46, 127.16, 126.93, 124.99, 115.69, 115.50, 61.23, 54.06, 53.97, 52.08, 40.72, 39.39, 36.88, 36.60, 33.83, 30.11, 25.02, 23.86.

**Mass spec:** expected neutral mass for $C_{37}H_{47}N_7O_8S$ (Da): 749.3107, observed neutral mass (Da): 749.3235, mass error (ppm): 3.8.

**UPLC Trace** Obtained with mobile phases of $H_2O$ + 0.1% formic acid (A) and acetonitrile + 0.1% formic acid (B). The peptide was eluted using a gradient of 0-70% mobile phase B over 4 min at a 0.5 mL/min flow rate. Mobile phase B was held at 70% for 1 min. The column was equilibrated with 0% mobile phase for 1 minute before and 2 minutes after the gradient. The peptide purity was determined to be 97%.

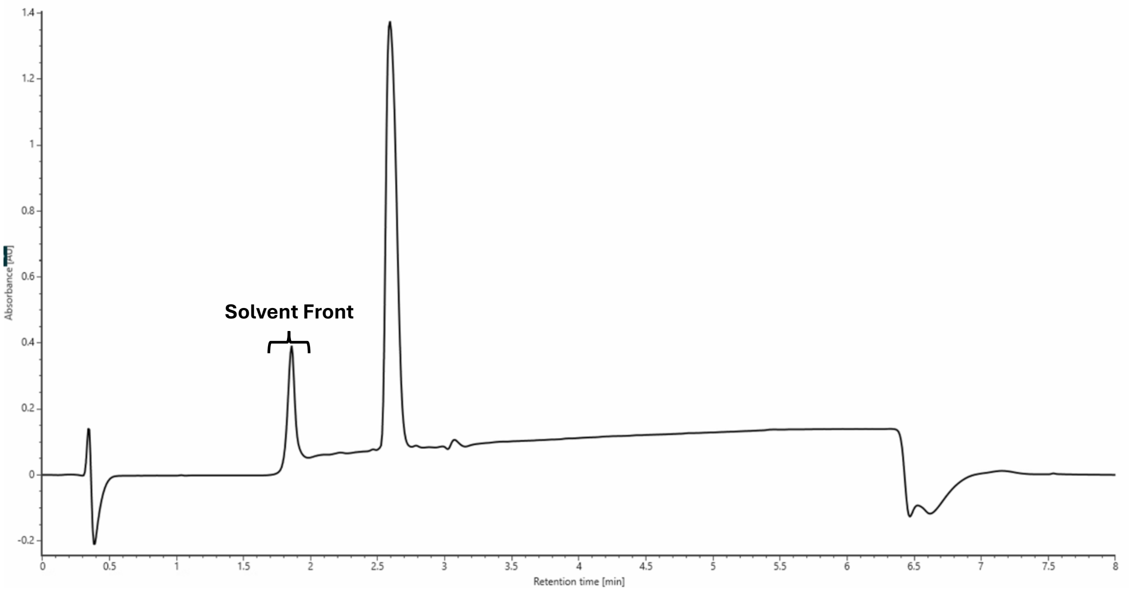

SMMP-dY-R-dF-T

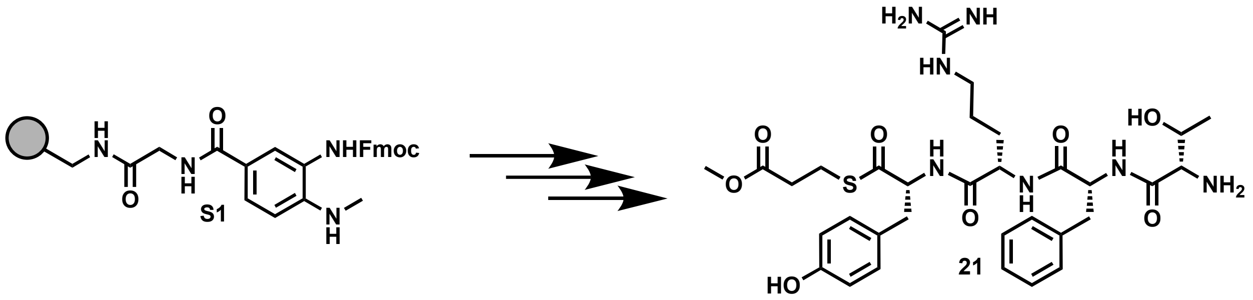

Peptide **21** was synthesized following the peptide synthesis protocol outlined above. Starting with Fmoc-Me-Dbz-OH loaded resin **S1** (0.1 mmol), amino acids Fmoc-D-Tyr(tBu)-OH, Fmoc-L-Arg(Pbf)-OH, Fmoc-D-Phe-OH, and Boc-L-Thr(tBu)-OH were used with methyl 3-mercaptopropionate as the cleaving thiol. The crude peptide was purified by reverse-phase semi-preparative HPLC with mobile phases of $H_2O$ + 0.05% TFA (A) and acetonitrile + 0.05% TFA (B). The peptide was eluted using a gradient of 0-55% mobile phase B over 25 minutes at a 20 mL/min flow rate. The column was equilibrated with 0% mobile phase B for 1 minute before and 5 minutes after the gradient, yielding an off-white solid (18 mg, 20% yield).

**$^{1}$H NMR** (800 MHz, DMSO) δ 9.31 (s, 1H), 8.75 (d, $J$ = 6.6 Hz, 1H), 8.66 (d, $J$ = 6.6 Hz, 1H), 8.38 (d, $J$ = 5.0 Hz, 1H), 8.09-7.99 (m, 3H), 7.72-7.61 (m, 1H), 7.55-6.94 (m, 10H), 6.66 (d, $J$ = 7.6 Hz, 2H), 5.52 (s, 1H), 4.92-4.83 (m, 1H), 4.51 (dd, $J$ = 7.8, 12.5 Hz, 1H), 4.36 (dd, $J$ = 8.7, 12.5 Hz, 1H), 3.55 (s, 1H), 3.49-3.32 (m, 6H), 3.07-2.87 (m, 6H), 2.78 (t, $J$ = 11.6 Hz, 1H), 2.70 (t, $J$ = 11.6 Hz, 1H), 2.63-2.54 (m, 2H), 1.36 (dt, $J$ = 7.8, 14.6 Hz, 1H), 1.23 (dd, $J$ = 7.0, 12.9 Hz, 1H), 1.16-1.06 (m, 2H), 0.74 (d, $J$ = 5.5 Hz, 3H).

**$^{13}$C NMR** (201 MHz, DMSO) δ 200.44, 172.11, 171.86, 170.67, 167.11, 157.27, 156.57, 137.46, 130.63, 129.73, 128.50, 127.15, 126.88, 115.50, 66.26, 61.22, 58.55, 54.05, 52.10, 52.01, 40.71, 39.20, 36.88, 33.84, 30.09, 24.94, 23.87, 19.89.

**Mass spec:** expected neutral mass for $C_{32}H_{45}N_7O_8S$ (Da): 687.3050, observed neutral mass (Da): 687.3083, mass error (ppm): 4.8.

**UPLC Trace** Obtained with mobile phases of $H_2O$ + 0.1% formic acid (A) and acetonitrile + 0.1% formic acid (B). The peptide was eluted using a gradient of 0-70% mobile phase B over 4 min at a 0.5 mL/min flow rate. Mobile phase B was held at 70% for 1 min. The column was equilibrated with 0% mobile phase for 1 minute before and 2 minutes after the gradient. The peptide purity was determined to be 96%.

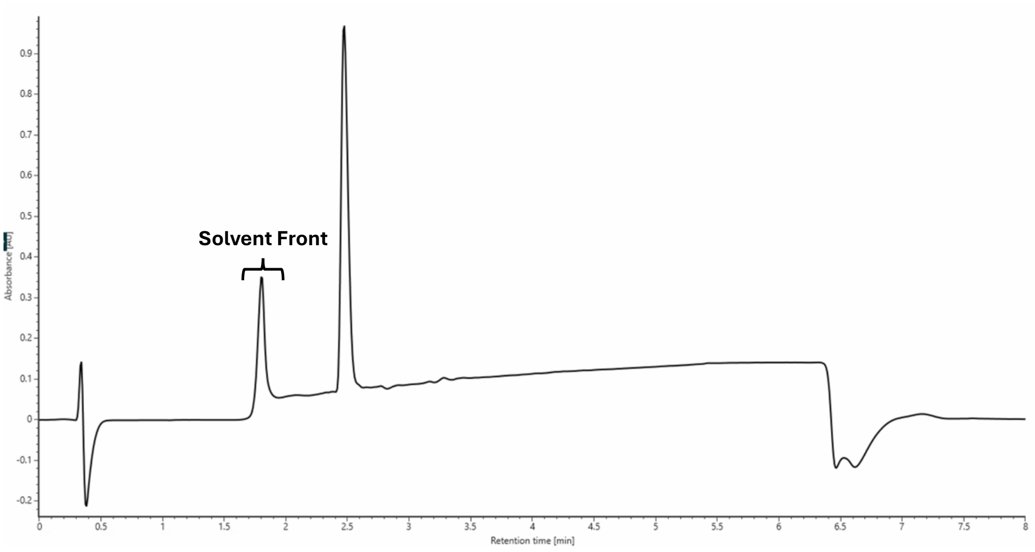

SMMP-dY-dR-dF-G

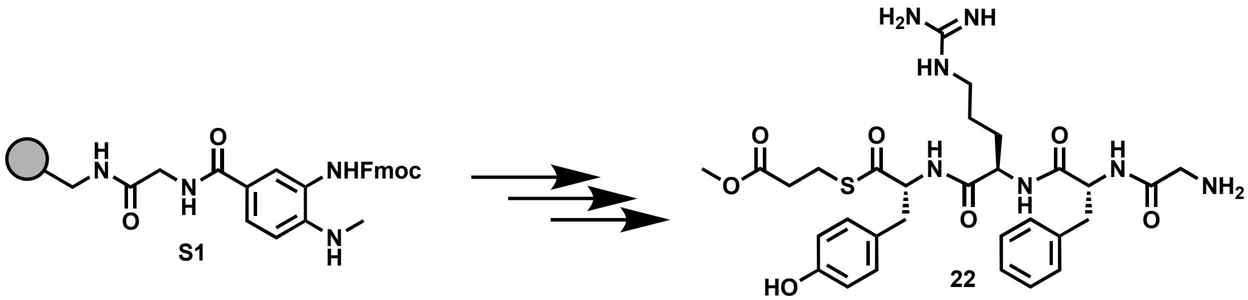

Peptide **22** was synthesized following the peptide synthesis protocol outlined above. Starting with Fmoc-Me-Dbz-OH loaded resin **S1** (0.1 mmol), amino acids Fmoc-D-Tyr(tBu)-OH, Fmoc-D-Arg(Pbf)-OH, Fmoc-D-Phe-OH, and Boc-Gly-OH were used with methyl 3-mercaptopropionate as the cleaving thiol. The crude peptide was purified by reverse-phase semi-preparative HPLC with mobile phases of $H_2O$ + 0.05% TFA (A) and acetonitrile + 0.05% TFA (B). The peptide was eluted using a gradient of 0-55% mobile phase B over 25 minutes at a 20 mL/min flow rate. The column was equilibrated with 0% mobile phase B for 1 minute before and 5 minutes after the gradient, yielding an off-white solid (27 mg, 31% yield).

**$^{1}$H NMR** (800 MHz, DMSO) δ 8.69-8.58 (m, 2H), 8.43 (d, *J* = 8.3 Hz, 1H), 7.99 (s, 3H), 7.86 (t, *J* = 6.2 Hz, 1H), 7.61-6.96 (m, 11H), 6.66 (d, *J* = 7.9 Hz, 2H), 4.67 (t, *J* = 8.9 Hz, 1H), 4.50 (dd, *J* = 7.1, 14.2 Hz, 1H), 4.36 (dd, *J* = 7.1, 13.5 Hz, 1H), 3.56-3.51 (m, 1H), 3.43 (d, *J* = 16.2 Hz, 1H), 3.11 (nonet, *J* = 6.8 Hz, 2H), 3.04-2.93 (m, 5H), 2.81 (dd, *J* = 9.5, 13.7 Hz, 1H), 2.72 (dd, *J* = 10.5, 13.3 Hz, 1H), 2.55 (t, *J* = 6.7 Hz, 2H), 1.78-1.68 (m, 1H), 1.60-1.42 (m, 3H).

**$^{13}$C NMR** (201 MHz, DMSO) δ 200.64, 172.04, 171.97, 171.07, 166.10, 157.37, 156.55, 61.31, 54.29, 52.50, 52.03, 40.86, 38.39, 36.53, 33.88, 29.59, 25.58, 23.78.

**Mass spec:** expected neutral mass for $C_{30}H_{41}N_7O_7S$ (Da): 643.2788, observed neutral mass (Da): 643.2791, mass error (ppm): 0.5.

**UPLC Trace** Obtained with mobile phases of $H_2O$ + 0.1% formic acid (A) and acetonitrile + 0.1% formic acid (B). The peptide was eluted using a gradient of 0-70% mobile phase B over 4 min at a 0.5 mL/min flow rate. Mobile phase B was held at 70% for 1 min. The column was equilibrated with 0% mobile phase for 1 minute before and 2 minutes after the gradient. The peptide purity was determined to be 95%.

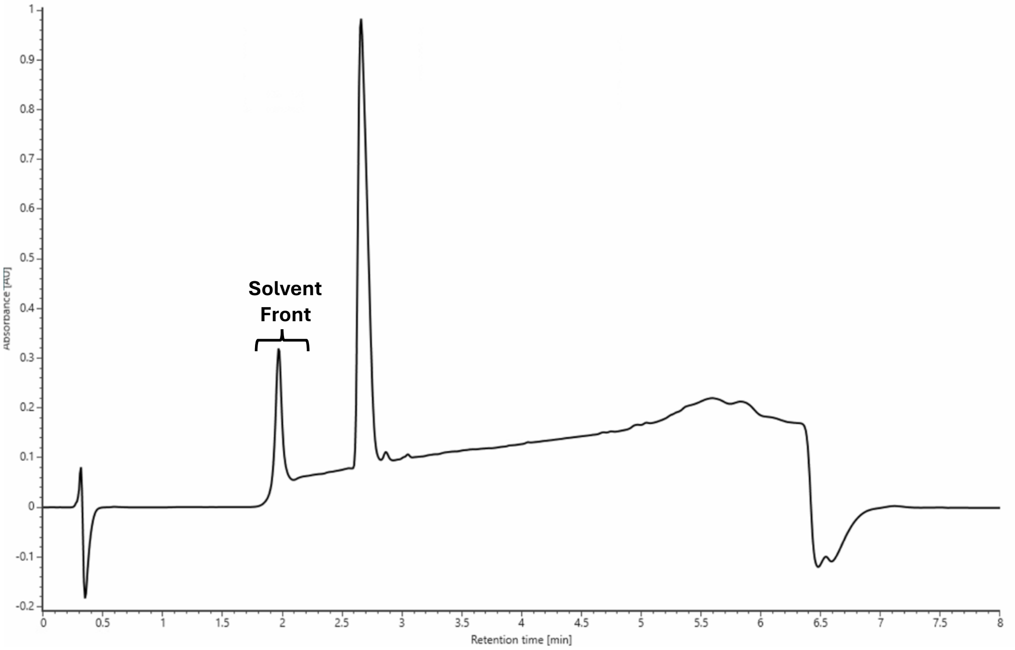

SMMP-dY-R-dF-G

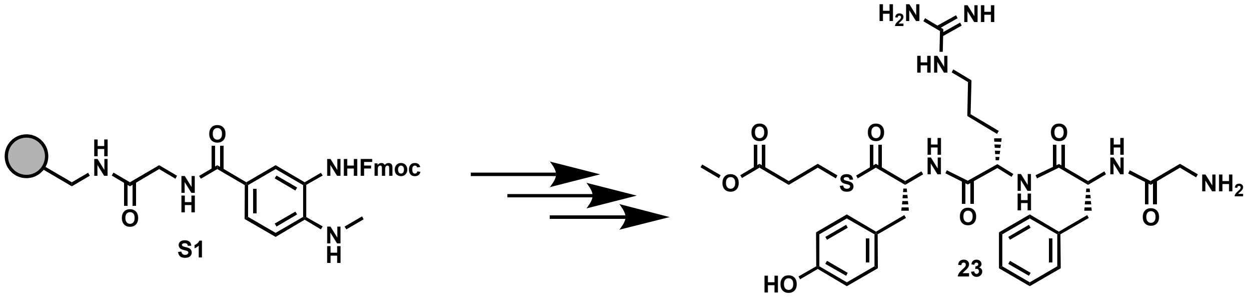

Peptide **23** was synthesized following the peptide synthesis protocol outlined above. Starting with Fmoc-Me-Dbz-OH loaded resin **S1** (0.1 mmol), amino acids Fmoc-D-Tyr(tBu)-OH, Fmoc-L-Arg(Pbf)-OH, Fmoc-D-Phe-OH, and Boc-Gly-OH were used with methyl 3-mercaptopropionate as the cleaving thiol. The crude peptide was purified by reverse-phase semi-preparative HPLC with mobile phases of $H_2O$ + 0.05% TFA (A) and acetonitrile + 0.05% TFA (B). The peptide was eluted using a gradient of 0-55% mobile phase B over 25 minutes at a 20 mL/min flow rate. The column was equilibrated with 0% mobile phase B for 1 minute before and 5 minutes after the gradient, yielding an off-white solid (27 mg, 31% yield).

**$^1$H NMR** (800 MHz, DMSO) δ 9.33 (s, 1H), 8.70 (d, *J* = 8.2 Hz, 1H), 8.61 (d, *J* = 8.1 Hz, 1H), 8.38 (d, *J* = 8.5 Hz, 1H), 7.99 (s, 3H), 7.72 (t, *J* = 5.1 Hz, 1H), 7.63-6.94 (m, 11H), 6.66 (d, *J* = 8.1 Hz, 2H), 4.76 (dd, *J* = 7.7, 14.2 Hz, 1H), 4.49 (dd, *J* = 8.5, 13.6 Hz, 1H), 4.32 (dd, *J* = 7.4, 13.6 Hz, 1H), 3.56 (d, *J* = 16.7 Hz, 1H), 3.44 (d, *J* = 16.5 Hz, 1H), 3.03-2.89 (m, 7H), 2.77 (dd, *J* = 9.5, 13.4 Hz, 1H), 2.71 (dd, *J* = 10.1, 12.5 Hz, 1H), 2.57 (dd, *J* = 5.6, 10.2 Hz, 2H), 1.39 (dt, *J* = 6.6, 12.7 Hz, 1H), 1.26 (sextet, *J* = 7.3 Hz, 1H), 1.12 (quintet, *J* = 7.3 Hz, 2H).

**$^{13}$C NMR** (201 MHz, DMSO) δ 200.51, 172.11, 171.88, 170.75, 166.00, 157.33, 156.56, 137.66, 130.61, 129.72, 128.52, 127.21, 126.86, 115.50, 61.27, 54.30, 52.12, 40.71, 39.05, 36.76, 33.84, 29.84, 25.00, 23.85.

**Mass spec:** expected neutral mass for $C_{30}H_{41}N_7O_7S$ (Da): 643.2788, observed neutral mass (Da): 643.2810, mass error (ppm): 3.4.

**UPLC Trace** Obtained with mobile phases of $H_2O$ + 0.1% formic acid (A) and acetonitrile + 0.1% formic acid (B). The peptide was eluted using a gradient of 0-70% mobile phase B over 4 min at a 0.5 mL/min flow rate. Mobile phase B was held at 70% for 1 min. The column was equilibrated with 0% mobile phase for 1 minute before and 2 minutes after the gradient. The peptide purity was determined to be 96%.

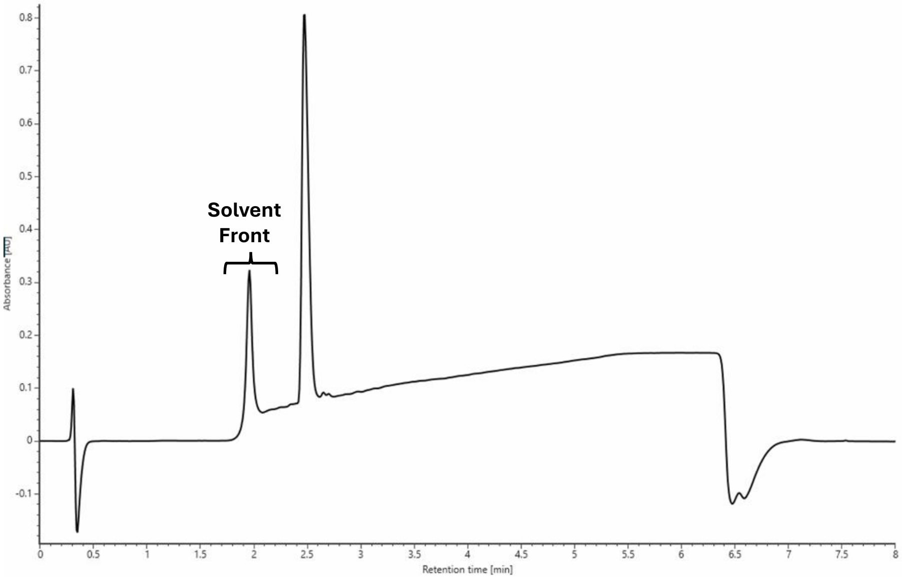

SMMP-dY-A-dF-V

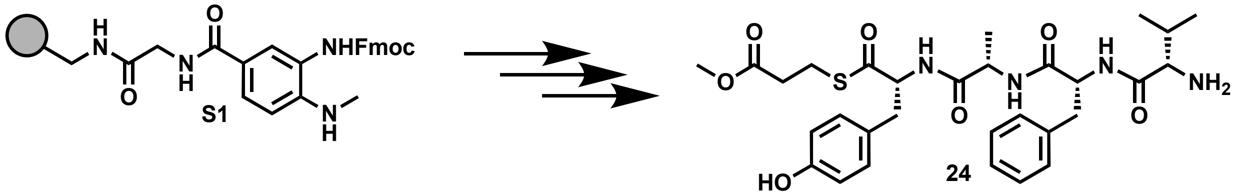

Peptide **24** was synthesized following the peptide synthesis protocol outlined above. Starting with Fmoc-Me-Dbz-OH loaded resin **S1** (0.1 mmol), amino acids Fmoc-D-Tyr(tBu)-OH, Fmoc-L-Ala-OH, Fmoc-D-Phe-OH, and Boc-L-Val-OH were used with methyl 3-mercaptopropionate as the cleaving thiol. The crude peptide was purified by reverse-phase semi-preparative HPLC with mobile phases of $H_2O$ + 0.05% TFA (A) and acetonitrile + 0.05% TFA (B). The peptide was eluted using a gradient of 0-55% mobile phase B over 25 minutes at a 20 mL/min flow rate. The column was equilibrated with 0% mobile phase B for 1 minute before and 5 minutes after the gradient, yielding an off-white solid (31mg, 43% yield).

**$^1$H NMR** (800 MHz, DMSO) δ 8.69 (d, *J* = 8.8 Hz, 1H), 8.65 (d, *J* = 8.2 Hz, 1H), 8.36 (d, *J* = 8.1 Hz, 1H), 7.93 (d, *J* = 4.5 Hz, 3H), 7.23 (q, *J* = 7.7 Hz, 4H), 7.16 (t, *J* = 7.2 Hz, 1H), 7.01 (d, *J* = 8.0 Hz, 2H), 6.64 (d, *J* = 8.2 Hz, 2H), 4.84 (dt, *J* = 4.7, 9.6 Hz, 1H), 4.51 (dt, *J* = 4.4, 8.6 Hz, 1H), 4.36 (quintet, *J* = 7.1 Hz, 1H), 3.61 (s, 4H), 3.05-2.95 (m, 4H), 2.69 (ddd, *J* = 11.0, 12.9, 21.2 Hz, 2H), 2.59 (dt, *J* = 4.0, 6.5 Hz, 2H), 1.83 (dq, *J* = 6.8, 12.4 Hz, 1H), 0.91 (d, *J* = 6.8 Hz, 3H), 0.69 (d, *J* = 6.8 Hz, 3H), 0.44 (d, *J* = 6.8 Hz, 3H).

**$^{13}$C NMR** (201 MHz, DMSO) δ 200.65, 172.82, 172.13, 170.45, 167.96, 156.51, 137.73, 130.60, 129.73, 128.45, 127.25, 126.81, 115.42, 61.09, 57.66, 53.99, 52.09, 48.16, 39.23, 36.75, 33.85, 30.06, 23.88, 19.20, 18.76, 16.95.

**Mass spec:** expected neutral mass for $C_{30}H_{40}N_4O_7S$ (Da): 600.2618, observed neutral mass (Da): 600.2621, mass error (ppm): 0.6.

**UPLC Trace:** Obtained with mobile phases of $H_2O$ + 0.1% formic acid (A) and acetonitrile + 0.1% formic acid (B). The peptide was eluted using a gradient of 0-70% mobile phase B over 4 min at a 0.5 mL/min flow rate. Mobile phase B was held at 70% for 1 min. The column was equilibrated with 0% mobile phase for 1 minute before and 2 minutes after the gradient. The peptide purity was determined to be 98%.

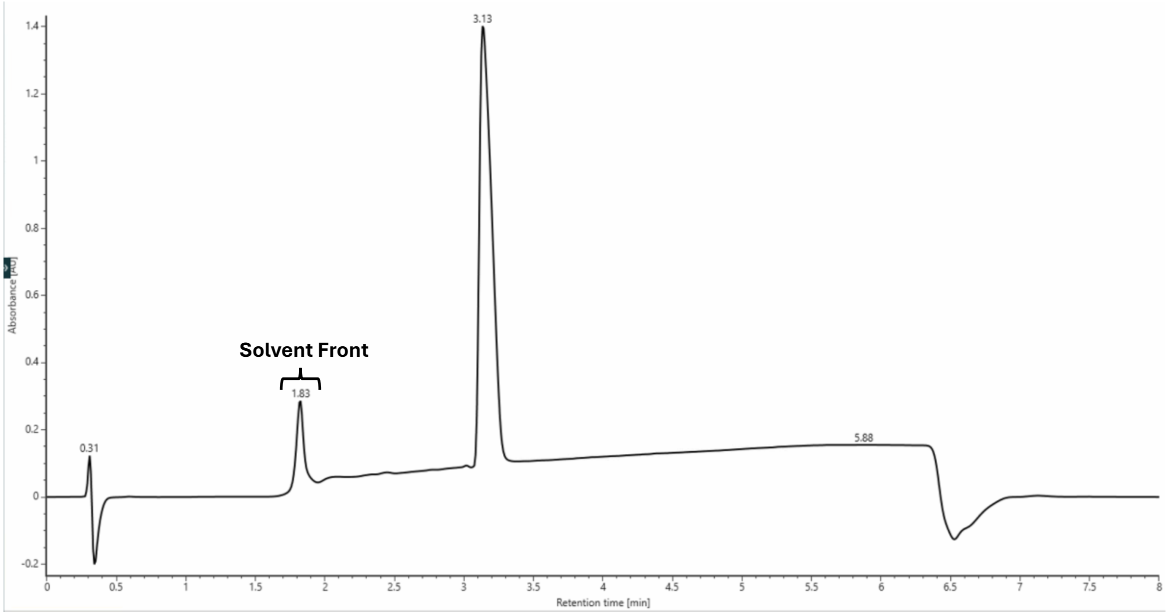

dY-W-dF-V

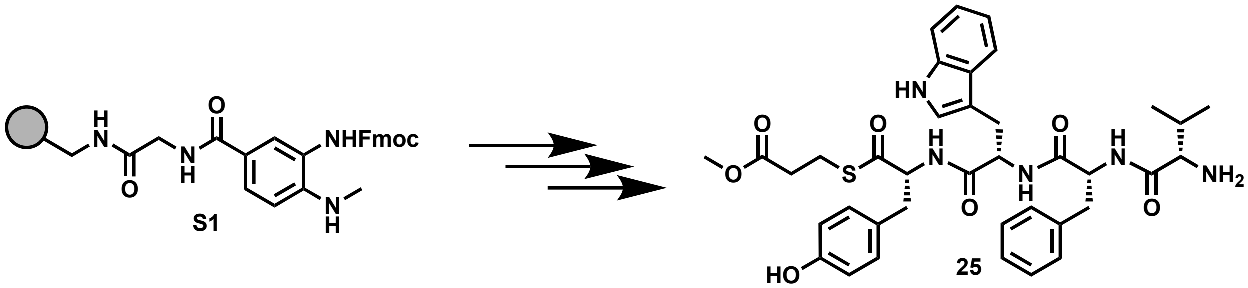

Peptide **25** was synthesized following the peptide synthesis protocol outlined above. Starting with Fmoc-Me-Dbz-OH loaded resin **S1** (0.1 mmol), amino acids Fmoc-D-Tyr(tBu)-OH, Fmoc-L-Trp(Boc)-OH, Fmoc-D-Phe-OH, and Boc-L-Val-OH were used with methyl 3-mercaptopropionate as the cleaving thiol. The crude peptide was purified by reverse-phase semi-preparative HPLC with mobile phases of $H_2O$ + 0.05% TFA (A) and acetonitrile + 0.05% TFA (B). The peptide was eluted using a gradient of 0-55% mobile phase B over 25 minutes at a 20 mL/min flow rate. The column was equilibrated with 0% mobile phase B for 1 minute before and 5 minutes after the gradient, yielding an off-white solid (30mg, 36% yield).
**$^1$H NMR** (800 MHz, DMSO) δ 10.80 (s, 1H), 9.01 (d, *J* = 8.8 Hz, 1H), 8.53 (q, *J* = 8.5 Hz, 2H), 7.89 (s, 3H), 7.77 (d, *J* = 7.3 Hz, 1H), 7.29 (d, *J* = 7.8 Hz, 1H), 7.13 (d, *J* = 8.9 Hz, 3H), 7.09-7.00 (m, 5H), 6.92 (d, *J* = 7.6 Hz, 2H), 6.65 (d, *J* = 7.7 Hz, 2H), 4.84 (dt, *J* = 3.0, 9.5 Hz, 1H), 4.61 (septet, *J* = 3.2 Hz, 1H), 4.57 (sextet, *J* = 5.1 Hz, 1H), 3.62 (s, 3H), 3.09-2.96 (m, 3H), 2.72 (t, *J* = 12.3 Hz, 2H), 2.65-2.53 (m, 4H), 2.28 (t, *J* = 12.2 Hz, 1H), 1.79 (octet, *J* = 6.6 Hz, 1H), 0.67 (d, *J* = 6.4 Hz, 3H), 0.40 (d, *J* = 6.6 Hz, 3H).
**$^{13}$C NMR** (201 MHz, DMSO) δ 200.64, 172.75, 172.06, 170.48, 167.87, 156.48, 137.61, 136.50, 130.72, 129.60, 128.16, 127.52, 127.26, 126.52, 124.67, 121.24, 118.51, 115.39, 111.60, 110.22, 61.26, 57.47, 53.69, 53.38, 52.03, 38.99, 36.91, 33.79, 29.96, 28.83, 23.83, 18.77, 16.68.
**Mass spec:** expected neutral mass for $C_{38}H_{45}N_5O_7S$ (Da): 715.3040, observed neutral mass (Da): 715.3029, mass error (ppm): -1.5.
**UPLC Trace:** Obtained with mobile phases of $H_2O$ + 0.1% formic acid (A) and acetonitrile + 0.1% formic acid (B). The peptide was eluted using a gradient of 0-70% mobile phase B over 4 min at a 0.5 mL/min flow rate. Mobile phase B was held at 70% for 1 min. The column was equilibrated with 0% mobile phase for 1 minute before and 2 minutes after the gradient. The peptide purity was determined to be 98%.

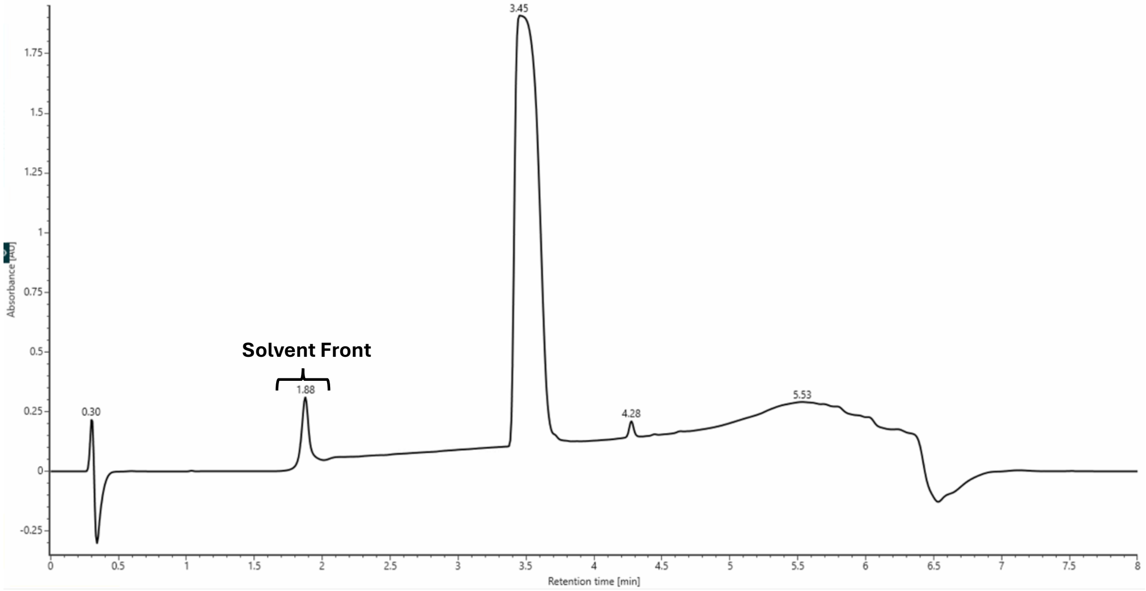

SMMP-dY-S-dF-V

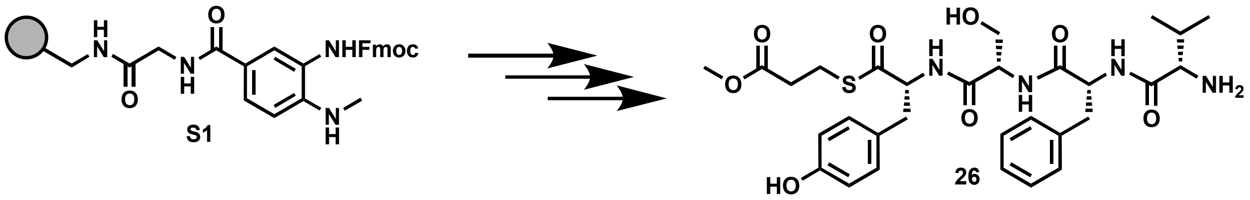

Peptide **26** was synthesized following the peptide synthesis protocol outlined above. Starting with Fmoc-Me-Dbz-OH loaded resin **S1** (0.1 mmol), amino acids Fmoc-D-Tyr(tBu)-OH, Fmoc-L-Ser(tBu)-OH, Fmoc-D-Phe-OH, and Boc-L-Val-OH were used with methyl 3-mercaptopropionate as the cleaving thiol. The crude peptide was purified by reverse-phase semi-preparative HPLC with mobile phases of $H_2O$ + 0.05% TFA (A) and acetonitrile + 0.05% TFA (B). The peptide was eluted using a gradient of 0-55% mobile phase B over 25 minutes at a 20 mL/min flow rate. The column was equilibrated with 0% mobile phase B for 1 minute before and 5 minutes after the gradient, yielding an off-white solid (28mg, 38% yield).

**$^1$H NMR** (800 MHz, DMSO) δ 8.66 (d, *J* = 8.3 Hz, 2H), 8.39 (d, *J* = 8.3 Hz, 1H), 7.92 (s, 3H), 7.27 (d, *J* = 7.8 Hz, 2H), 7.23 (t, *J* = 7.2 Hz, 2H), 7.16 (t, *J* = 7.2 Hz, 1H), 7.00 (d, *J* = 7.9 Hz, 2H), 6.65 (d, *J* = 7.7 Hz, 2H), 4.92 (dt, *J* = 3.7, 9.6 Hz, 1H), 4.53-4.43 (m, 2H), 3.61 (s, 3H), 3.38 (dd, *J* = 4.9, 10.5 Hz, 1H), 3.30 (dd, *J* = 7.3, 10.2 Hz, 1H), 3.09 (dd, *J* = 2.7, 13.3 Hz, 1H), 3.02-2.90 (m, 3H), 2.76 (dd, *J* = 9.6, 13.6 Hz, 1H), 2.68 (t, *J* = 12.3 Hz, 1H), 2.56 (t, *J* = 6.7 Hz, 2H), 1.81 (sextet, *J* = 6.1 Hz, 1H), 0.67 (d, *J* = 6.5 Hz, 3H), 0.40 (d, *J* = 6.5 Hz, 3H).

**$^{13}$C NMR** (201 MHz, DMSO) δ 200.55, 172.11, 171.25, 170.56, 167.96, 156.53, 137.93, 130.59, 129.75, 128.42, 127.10, 126.76, 115.49, 62.32, 61.25, 57.67, 55.27, 54.04, 52.09, 39.27, 36.82, 33.84, 30.05, 23.84, 18.76, 16.83.

**Mass spec:** expected neutral mass for $C_{30}H_{40}N_4O_8S$ (Da): 616.2567, observed neutral mass (Da): 616.2577, mass error (ppm): 1.7.

**UPLC Trace:** Obtained with mobile phases of $H_2O$ + 0.1% formic acid (A) and acetonitrile + 0.1% formic acid (B). The peptide was eluted using a gradient of 0-70% mobile phase B over 4 min at a 0.5 mL/min flow rate. Mobile phase B was held at 70% for 1 min. The column was equilibrated with 0% mobile phase for 1 minute before and 2 minutes after the gradient. The peptide purity was determined to be 98%.

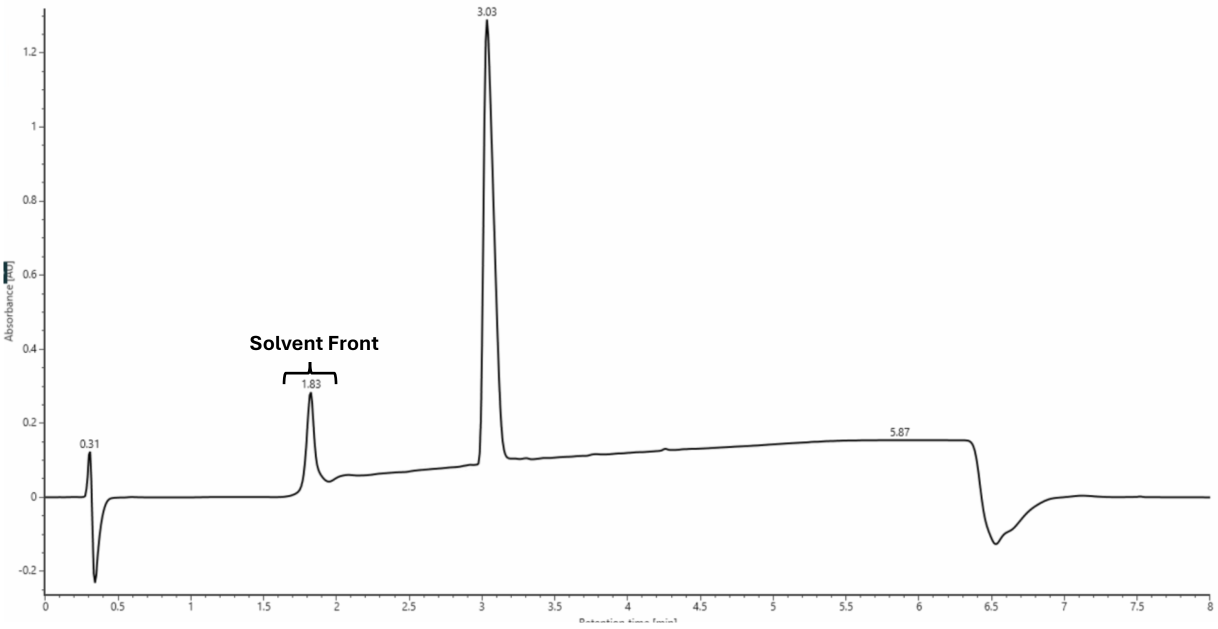

SBMP-dY-Q-dF-V

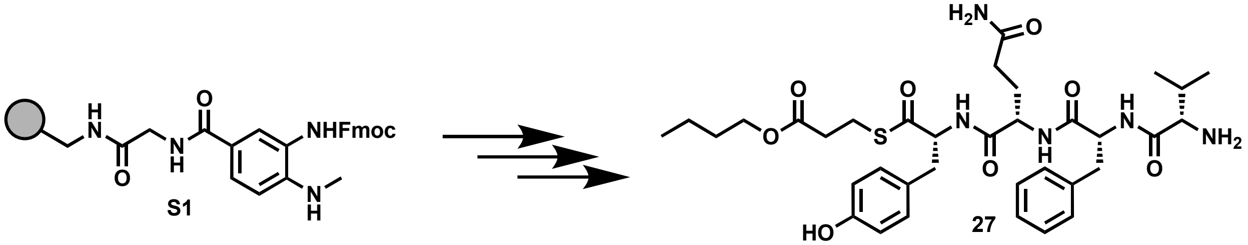

Peptide **27** was synthesized following the peptide synthesis protocol outlined above. Starting with Fmoc-Me-Dbz-OH loaded resin **S1** (0.1 mmol), amino acids Fmoc-D-Tyr(tBu)-OH, Fmoc-L-Gln(Trt)-OH, Fmoc-D-Phe-OH, and Boc-L-Val-OH were used with butyl 3-mercaptopropionate as the cleaving thiol. The crude peptide was purified by reverse-phase semi-preparative HPLC with mobile phases of $H_2O$ + 0.05% TFA (A) and acetonitrile + 0.05% TFA (B). The peptide was eluted using a gradient of 0-55% mobile phase B over 25 minutes at a 20 mL/min flow rate. The column was equilibrated with 0% mobile phase B for 1 minute before and 5 minutes after the gradient, yielding an off-white solid (30 mg, 37% yield).

**$^1$H NMR** (800 MHz, DMSO) δ 8.71 (t, *J* = 8.9 Hz, 2H), 8.42 (d, *J* = 8.1 Hz, 1H), 7.94 (s, 3H), 7.26 (d, *J* = 7.5 Hz, 2H), 7.24 (t, *J* = 7.3 Hz, 2H), 7.20 (s, 1H), 7.16 (t, *J* = 7.3 Hz, 1H), 7.01 (d, *J* = 7.5 Hz, 2H), 6.77 (s, 1H), 6.64 (d, *J* = 7.6 Hz, 2H), 4.85 (septet, *J* = 4.1 Hz, 1H), 4.48 (dt, *J* = 8.2, 5.3 Hz, 1H), 4.38 (dt, *J* = 8.2, 4.5 Hz, 1H), 4.03 (t, *J* = 6.4 Hz, 2H), 3.58 (t, *J* = 4.5 Hz, 1H), 3.09 (dd, *J* = 3.5, 13.7 Hz, 1H), 3.02 – 2.92 (m, 3H), 2.76 (dd, J = 10.0, 13.8 Hz, 1H), 2.69 (t, *J* = 11.9 Hz, 1H), 2.55 (dt, *J* = 4.5, 6.5 Hz, 2H), 1.93 (octet, *J* = 7.8 Hz, 2H), 1.81 (sextet, 6.4 Hz, 1H), 1.76 (sextet, *J* = 6.3 Hz, 1H), 1.62 – 1.5 (m, 3H), 1.32 (sextet, *J* = 7.3 Hz, 2H), 0.88 (t, *J* = 7.1, 3H), 0.65 (d, *J* = 6.6 Hz, 3H), 0.41 (d, *J* = 6.6 Hz, 3H).

**$^{13}$C NMR** (201 MHz, DMSO) δ 200.40, 174.06, 171.82, 171.58, 170.89, 167.98, 156.47, 137.81, 130.48, 129.61, 128.40, 127.10, 126.74, 115.45, 64.34, 61.29, 57.66, 54.16, 52.30, 39.12, 36.73, 33.96, 31.55, 30.52, 29.92, 28.61, 23.81, 18.98, 18.62, 16.84, 13.94.

**Mass spec:** expected neutral mass for $C_{35}H_{49}N_5O_8S$ (Da): 699.33008, observed neutral mass (Da): 699.3294, mass error (ppm): -1.1.

**UPLC Trace:** Obtained with mobile phases of $H_2O$ + 0.1% formic acid (A) and acetonitrile + 0.1% formic acid (B). The peptide was eluted using a gradient of 0-70% mobile phase B over 4 min at a 0.5 mL/min flow rate. Mobile phase B was held at 70% for 1 min. The column was equilibrated with 0% mobile phase for 1 minute before and 2 minutes after the gradient. The peptide purity was determined to be 98%.

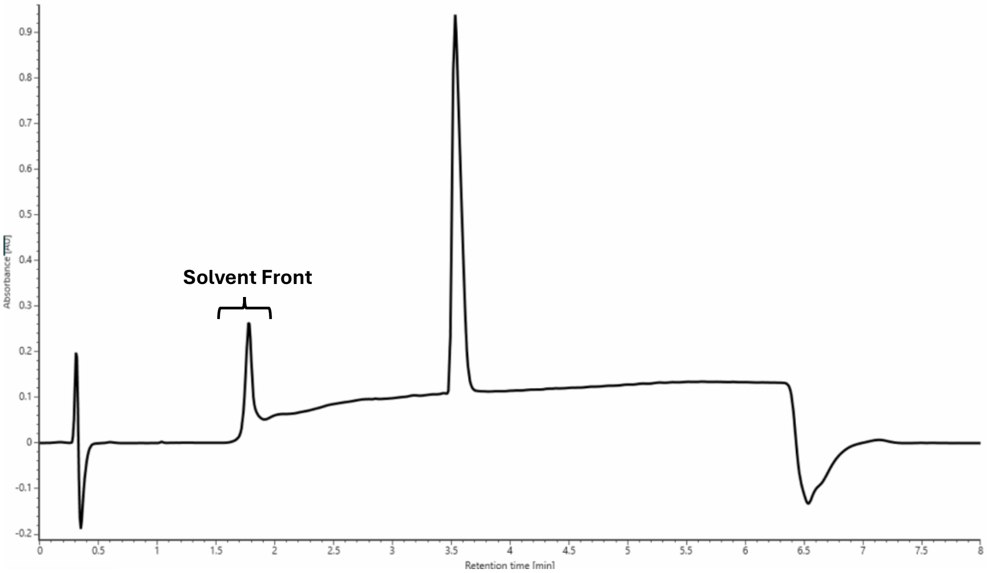

SMMP-dY-E-dF-V

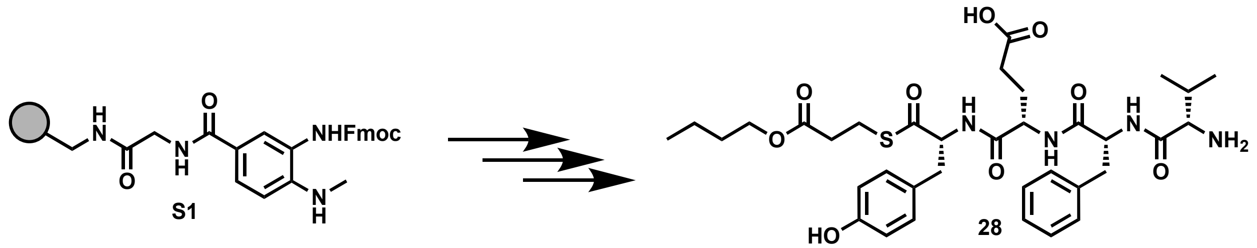

Peptide **28** was synthesized following the peptide synthesis protocol outlined above. Starting with Fmoc-Me-Dbz-OH loaded resin **S1** (0.1 mmol), amino acids Fmoc-D-Tyr(tBu)-OH, Fmoc-L-Glu(tBu)-OH, Fmoc-D-Phe-OH, and Boc-L-Val-OH were used with methyl 3-mercaptopropionate as the cleaving thiol. The crude peptide was purified by reverse-phase semi-preparative HPLC with mobile phases of $H_2O$ + 0.05% TFA (A) and acetonitrile + 0.05% TFA (B). The peptide was eluted using a gradient of 0-55% mobile phase B over 25 minutes at a 20 mL/min flow rate. The column was equilibrated with 0% mobile phase B for 1 minute before and 5 minutes after the gradient, yielding an off-white solid (29 mg, 37% yield).
**$^1$H NMR** (800 MHz, DMSO) δ 8.76-8.69 (m, 2H), 8.43 (d, *J* = 8.9 Hz, 1H), 7.93 (s, 3H), 7.29-7.21 (m, 5H), 7.16 (t, *J* = 7.4 Hz, 1H), 7.01 (d, *J* = 7.7 Hz, 2H), 6.65 (d, *J* = 7.7 Hz, 2H), 4.89 (dt, *J* = 4.2, 9.4 Hz, 1H), 4.50 (dt, *J* = 5.2, 8.5 Hz, 1H), 4.42 (dd, *J* = 7.3, 12.5 Hz, 1H), 3.61-3.56 (m, 1H), 3.06-2.94 (m, 5H), 2.71 (quintet, *J* = 11.5 Hz, 2H), 2.58 (dd, *J* = 6.8, 8.8 Hz, 2H), 2.05-1.97 (m, 1H), 1.96-1.89 (m, 1H), 1.83 (sextet, *J* = 6.6 Hz, 1H), 1.74 (dq, *J* = 4.5, 9.7 Hz, 1H), 1.61-1.53 (m, 1H), 0.68 (d, *J* = 6.5 Hz, 3H), 0.43 (d, *J* = 6.5 Hz, 3H).
**$^{13}$C NMR** (201 MHz, DMSO) δ 200.48, 174.11, 172.11, 171.64, 170.94, 167.99, 156.57, 137.72, 130.53, 129.70, 128.46, 127.16, 126.85, 115.50, 61.32, 57.67, 54.08, 52.09, 51.85, 39.31, 36.75, 33.84, 30.20, 30.06, 28.30, 23.87, 18.74, 16.92.
**Mass spec:** expected neutral mass for $C_{32}H_{42}N_4O_9S$ (Da): 658.2672, observed neutral mass (Da): 658.2700, mass error (ppm): 4.1.
**UPLC Trace:** Obtained with mobile phases of $H_2O$ + 0.1% formic acid (A) and acetonitrile + 0.1% formic acid (B). The peptide was eluted using a gradient of 0-70% mobile phase B over 4 min at a 0.5 mL/min flow rate. Mobile phase B was held at 70% for 1 min. The column was equilibrated with 0% mobile phase for 1 minute before and 2 minutes after the gradient. The peptide purity was determined to be 97%.

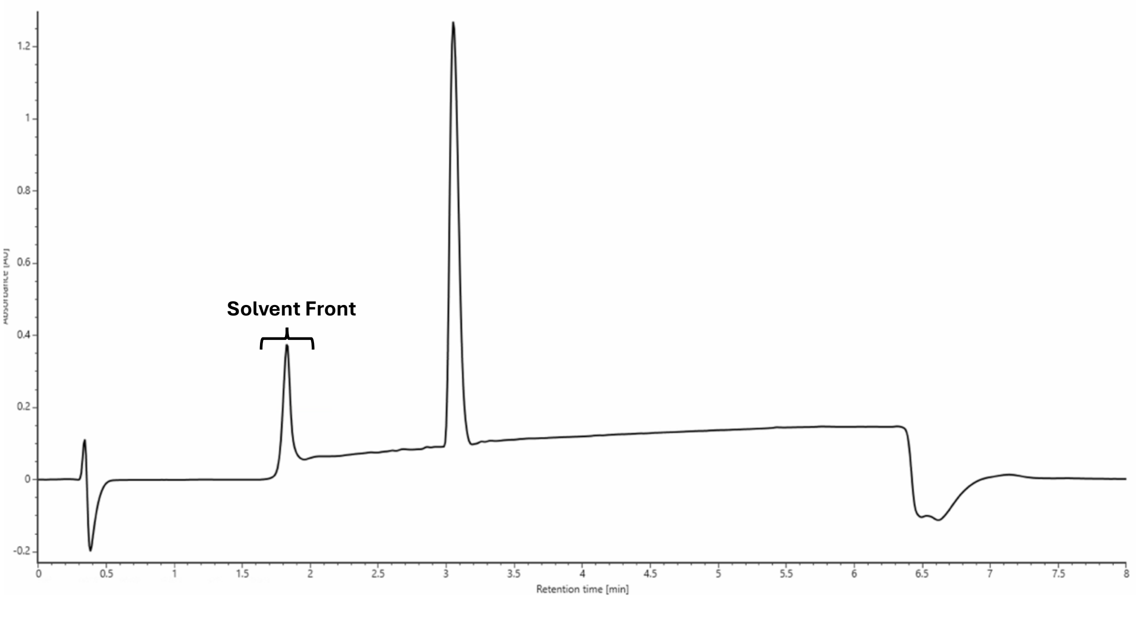

SMMP-dY-R-dT-V

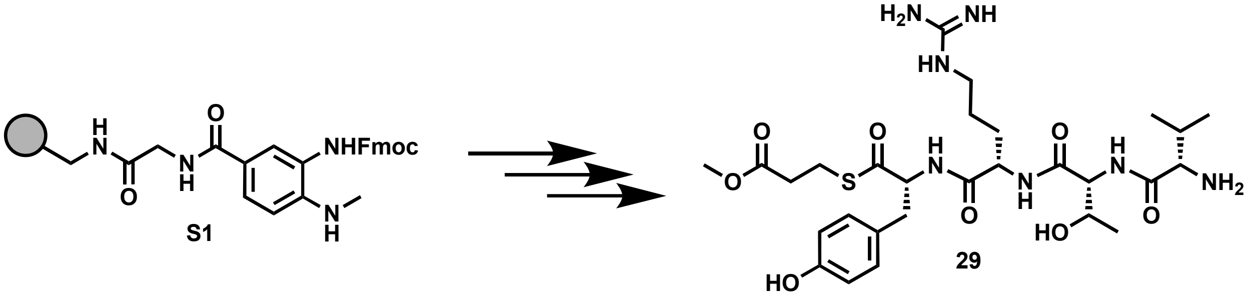

Peptide **29** was synthesized following the peptide synthesis protocol outlined above. Starting with Fmoc-Me-Dbz-OH loaded resin **S1** (0.1 mmol), amino acids Fmoc-D-Tyr(tBu)-OH, Fmoc-L-Arg(Pbf)-OH, Fmoc-D-Thr(tBu)-OH, and Boc-L-Val-OH were used with methyl 3-mercaptopropionate as the cleaving thiol. The crude peptide was purified by reverse-phase semi-preparative HPLC with mobile phases of $H_2O$ + 0.05% TFA (A) and acetonitrile + 0.05% TFA (B). The peptide was eluted using a gradient of 0-55% mobile phase B over 25 minutes at a 20 mL/min flow rate. The column was equilibrated with 0% mobile phase B for 1 minute before and 5 minutes after the gradient, yielding an off-white solid (25 mg, 29% yield).

**$^{1}$H NMR** (800 MHz, DMSO) δ 9.32 (s, 1H), 8.65 (d, *J* = 8.2 Hz, 1H), 8.51 (d, *J* = 8.4 Hz, 1H), 8.08 (s, 3H), 8.02 (d, *J* = 8.3 Hz, 1H), 7.71 (t, *J* = 5.4 Hz, 1H), 7.57-6.92 (m, 5H), 6.65 (d, *J* = 8.2 Hz, 2H), 5.07 (s, 1H), 4.49 (dt, *J* = 5.6, 8.6 Hz, 1H), 4.44 (q, *J* = 4.3 Hz, 1H), 4.39 (dd, *J* = 7.9, 13.2 Hz, 1H), 3.91 (t, *J* = 4.9 Hz, 1H), 3.85 (s, 1H), 3.61 (s, 3H), 3.03-2.93 (m, 5H), 2.69 (dd, *J* = 10.0, 13.9 Hz, 1H), 2.56 (dt, *J* = 2.7, 6.8 Hz, 2H), 2.10 (octet, *J* = 6.5 Hz, 1H), 1.48 (dt, *J* = 7.2, 12.2 Hz, 1H), 1.38-1.21 (m, 3H), 1.02 (d, *J* = 6.2 Hz, 3H), 0.94 (dd, *J* = 7.0, 23.0 Hz, 6H).

**$^{13}$C NMR** (201 MHz, DMSO) δ 200.37, 172.02, 171.85, 169.34, 168.49, 157.24, 156.49, 130.53, 127.09, 115.43, 67.40, 61.20, 58.27, 57.54, 52.00, 40.60, 36.72, 33.75, 30.32, 29.89, 24.94, 23.77, 19.96, 18.76, 17.70.

**Mass spec:** expected neutral mass for $C_{28}H_{45}N_7O_8S$ (Da): 639.3050, observed neutral mass (Da): 639.3063, mass error (ppm): 1.9.

**UPLC Trace:** Obtained with mobile phases of $H_2O$ + 0.1% formic acid (A) and acetonitrile + 0.1% formic acid (B). The peptide was eluted using a gradient of 0-70% mobile phase B over 4 min at a 0.5 mL/min flow rate. Mobile phase B was held at 70% for 1 min. The column was equilibrated with 0% mobile phase for 1 minute before and 2 minutes after the gradient. The peptide purity was determined to be 97%.

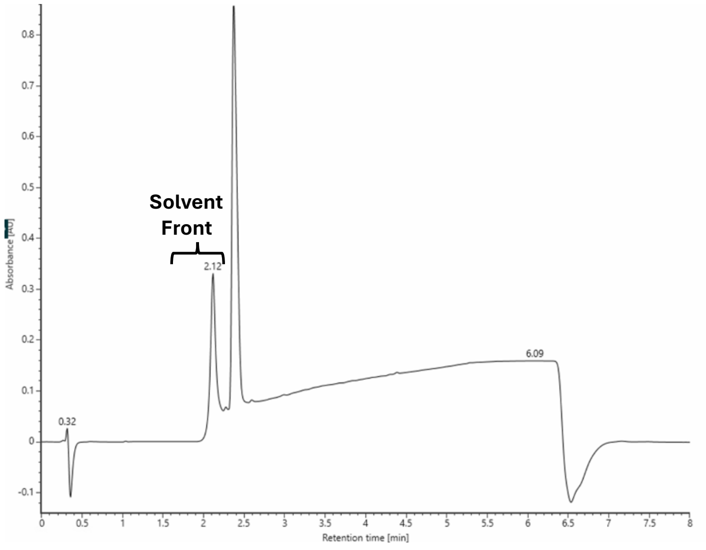

SMMP-dY-R-dW-V

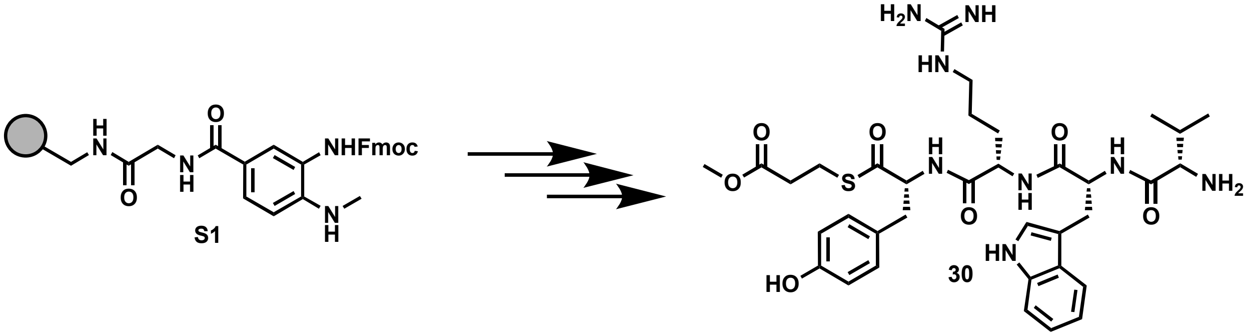

Peptide **30** was synthesized following the peptide synthesis protocol outlined above. Starting with Fmoc-Me-Dbz-OH loaded resin **S1** (0.1 mmol), amino acids Fmoc-D-Tyr(tBu)-OH, Fmoc-L-Arg(Pbf)-OH, Fmoc-D-Gln(Trt)-OH, and Boc-L-Val-OH were used with methyl 3-mercaptopropionate as the cleaving thiol. The crude peptide was purified by reverse-phase semi-preparative HPLC with mobile phases of $H_2O$ + 0.05% TFA (A) and acetonitrile + 0.05% TFA (B). The peptide was eluted using a gradient of 0-55% mobile phase B over 25 minutes at a 20 mL/min flow rate. The column was equilibrated with 0% mobile phase B for 1 minute before and 5 minutes after the gradient, yielding an off-white solid (26 mg, 27% yield).

**$^1$H NMR** (800 MHz, DMSO) δ 10.82 (s, 1H), 9.34 (s, 1H), 8.68 (d, *J* = 7.6 Hz, 1H), 8.61 (d, 8.1 Hz, 1H), 8.48 (d, 7.6 Hz, 1H), 7.90 (s, 3H), 7.70 (d, *J* = 7.3 Hz, 2H), 7.58-7.06 (m, 5H), 7.02 (q, *J* = 8.1 Hz, 3H), 6.94 (t, *J* = 7.3 Hz, 1H), 6.66 (d, *J* = 8.3 Hz, 2H), 4.93 (dt, *J* = 4.6, 9.2 Hz, 1H), 4.51 (dt, *J* = 5.4, 8.5 Hz, 1H), 4.40 (dt, *J* = 5.4, 8.1 Hz, 1H), 3.64-3.60 (m, 3H), 3.09 (dd, *J* = 4.6, 13.9 Hz, 1H), 3.04-2.85 (m, 6H), 2.70 (dd, *J* = 9.4, 13.7 Hz, 1H), 2.57 (dt, *J* = 2.2, 6.5 Hz, 2H), 1.76 (sextet, *J* = 6.9 Hz, 1H), 1.47-1.36 (m, 1H), 1.35-1.13 (m, 3H), 0.63 (d, *J* = 7.2 Hz, 3H), 0.33 (d, *J* = 6.6 Hz, 3H).

**$^{13}$C NMR** (201 MHz, DMSO) δ 200.40, 172.02, 171.45, 167.87, 157.25, 156.51, 136.59, 130.56, 127.50, 127.08, 124.55, 121.17, 118.27, 115.44, 111.50, 109.79, 61.18, 57.51, 53.25, 52.00, 40.62, 36.81, 33.76, 30.01, 29.61, 24.99, 23.78, 18.66, 16.57.

**Mass spec:** expected neutral mass for $C_{35}H_{48}N_8O_7S$ (Da): 724.3367, observed neutral mass (Da): 724.3376, mass error (ppm): -0.3.

**UPLC Trace:** Obtained with mobile phases of $H_2O$ + 0.1% formic acid (A) and acetonitrile + 0.1% formic acid (B). The peptide was eluted using a gradient of 0-70% mobile phase B over 4 min at a 0.5 mL/min flow rate. Mobile phase B was held at 70% for 1 min. The column was equilibrated with 0% mobile phase for 1 minute before and 2 minutes after the gradient. The peptide purity was determined to be >99%.

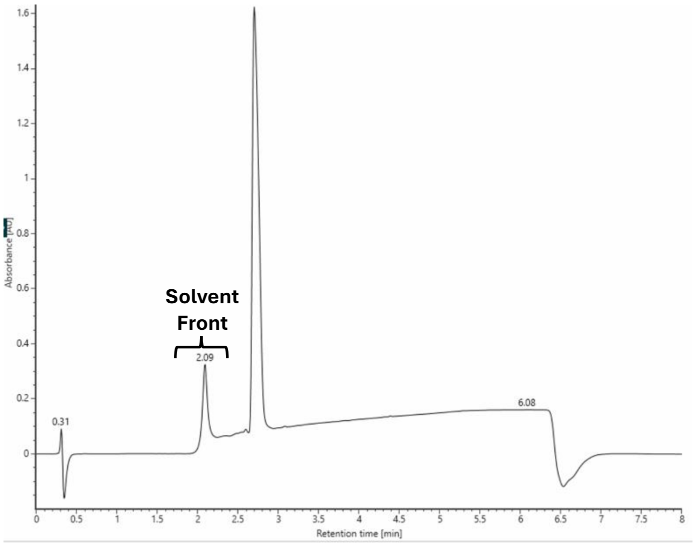

SMMP-dY-R-dA-V

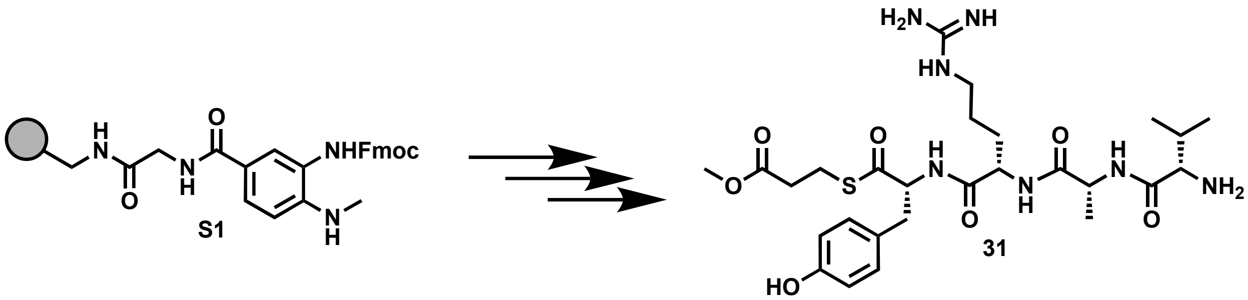

Peptide **31** was synthesized following the peptide synthesis protocol outlined above. Starting with Fmoc-Me-Dbz-OH loaded resin **S1** (0.1 mmol), amino acids Fmoc-D-Tyr(tBu)-OH, Fmoc-L-Arg(Pbf)-OH, Fmoc-D-Ala-OH, and Boc-L-Val-OH were used with methyl 3-mercaptopropionate as the cleaving thiol. The crude peptide was purified by reverse-phase semi-preparative HPLC with mobile phases of $H_2O$ + 0.05% TFA (A) and acetonitrile + 0.05% TFA (B). The peptide was eluted using a gradient of 0-55% mobile phase B over 25 minutes at a 20 mL/min flow rate. The column was equilibrated with 0% mobile phase B for 1 minute before and 5 minutes after the gradient, yielding an off-white solid (19 mg, 23% yield).

**$^{1}$H NMR** (800 MHz, DMSO) δ 8.66 (dd, *J* = 8.1, 11.3 Hz, 1H), 8.24 (d, *J* = 8.4 Hz, 1H), 8.10 (s, 2H), 7.78 (t, *J* = 4.6 Hz, 1H), 7.63-6.90 (m, 3H), 6.66 (d, *J* = 8.2 Hz, 1H), 4.60-4.21 (m, 3H), 3.69 (s, 1H), 3.61 (s, 2H), 3.43 (q, *J* = 7.1 Hz, 4H), 2.98 (octet, *J* = 6.8 Hz, 3H), 2.69 (dd, *J* = 10.2, 13.8 Hz, 1H), 2.57 (dt, *J* = 3.4, 6.7 Hz, 1H), 2.09-1.99 (m, 3H), 1.45 (dt, *J* = 5.5, 12.8 Hz, 1H), 1.34-1.08 (m, 4H), 1.05 (t, *J* = 7.0 Hz, 6H), 0.92 (d, *J* = 6.7 Hz, 4H).

**$^{13}$C NMR** (201 MHz, DMSO) δ 206.95, 200.48, 172.10, 171.94, 171.85, 167.77, 157.34, 156.57, 130.61, 127.20, 115.50, 61.24, 57.55, 56.49, 52.07, 51.93, 48.55, 40.65, 36.78, 33.84, 31.12, 30.30, 29.98, 25.12, 23.85, 19.89, 18.99, 18.69, 18.20.

**Mass spec:** expected neutral mass for $C_{27}H_{43}N_7O_7S$ (Da): 609.2945, observed neutral mass (Da): 609.2959, mass error (ppm): 2.4.

**UPLC Trace:** Obtained with mobile phases of $H_2O$ + 0.1% formic acid (A) and acetonitrile + 0.1% formic acid (B). The peptide was eluted using a gradient of 0-70% mobile phase B over 4 min at a 0.5 mL/min flow rate. Mobile phase B was held at 70% for 1 min. The column was equilibrated with 0% mobile phase for 1 minute before and 2 minutes after the gradient. The peptide purity was determined to be 97%.

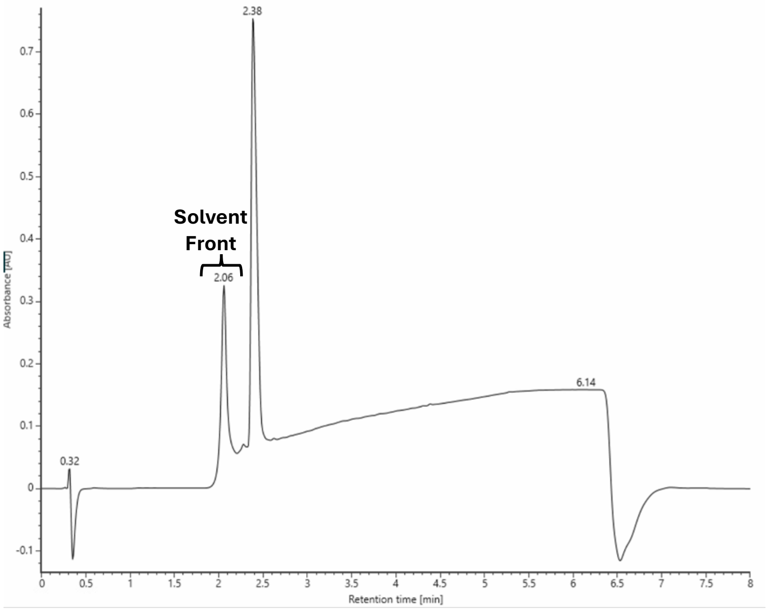

SMMP-dY-R-dDap-V

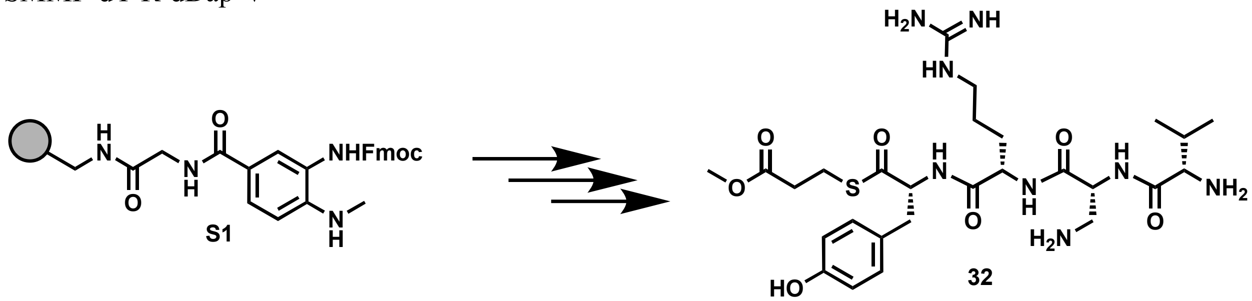

Peptide **32** was synthesized following the peptide synthesis protocol outlined above. Starting with Fmoc-Me-Dbz-OH loaded resin **S1** (0.1 mmol), amino acids Fmoc-D-Tyr(tBu)-OH, Fmoc-L-Arg(Pbf)-OH, Fmoc-D-Dap(Boc)-OH, and Boc-L-Val-OH were used with methyl 3-mercaptopropionate as the cleaving thiol. The crude peptide was purified by reverse-phase semi-preparative HPLC with mobile phases of $H_2O$ + 0.05% TFA (A) and acetonitrile + 0.05% TFA (B). The peptide was eluted using a gradient of 0-55% mobile phase B over 25 minutes at a 20 mL/min flow rate. The column was equilibrated with 0% mobile phase B for 1 minute before and 5 minutes after the gradient, yielding an off-white solid (28 mg, 29% yield).

**$^1$H NMR** (800 MHz, DMSO) δ 9.34 (s, 1H), 9.09 (d, *J* = 7.0 Hz, 1H), 8.68 (d, *J* = 8.1 Hz, 1H), 8.24-8.09 (m, 6H), 7.74 (t, *J* = 4.9 Hz, 1H), 7.63-6.87 (m, 5H), 6.66 (d, *J* = 7.7 Hz, 2H), 4.60 (q, *J* = 6.0 Hz, 1H), 4.51 (dt, *J* = 5.1, 8.4 Hz, 1H), 4.36 (dd, *J* = 7.5, 13.0 Hz, 1H), 3.68 (d, 4.6 Hz, 1H), 3.61 (s, 3H), 3.44 (q, *J* = 6.9 Hz, 1H), 3.19 (dd, *J* = 4.4, 12.6 Hz, 1H), 3.06-2.88 (m, 6H), 2.70 (dd, *J* = 10.4, 13.1 Hz, 1H), 2.56 (dd, *J* = 6.6, 104 Hz, 2H), 2.19 (sextet, *J* = 6.2 Hz, 1H), 1.50 (dd, *J* = 9.5, 12.7 Hz, 1H), 1.39-1.30 (m, 1H), 1.23 (octet, *J* = 7.0 Hz, 2H), 1.05 (t, *J* = 7.0 Hz, 1H), 0.95 (d, *J* = 6.8 Hz, 3H), 0.89 (d, *J* = 6.7 Hz, 3H).

**$^{13}$C NMR** (201 MHz, DMSO) δ 200.30, 172.01, 171.46, 169.36, 168.01, 157.29, 156.50, 130.52, 127.11, 115.43, 61.23, 58.08, 56.41, 52.33, 51.99, 51.53, 40.64, 36.71, 33.75, 29.80, 29.66, 24.87, 23.77, 19.00, 18.93, 17.48.

**Mass spec:** expected neutral mass for $C_{27}H_{44}N_8O_7S$ (Da): 624.3054, observed neutral mass (Da): 624.3066, mass error (ppm): 2.0.

**UPLC Trace:** Obtained with mobile phases of $H_2O$ + 0.1% formic acid (A) and acetonitrile + 0.1% formic acid (B). The peptide was eluted using a gradient of 0-70% mobile phase B over 4 min at a 0.5 mL/min flow rate. Mobile phase B was held at 70% for 1 min. The column was equilibrated with 0% mobile phase for 1 minute before and 2 minutes after the gradient. The peptide purity was determined to be 94%.

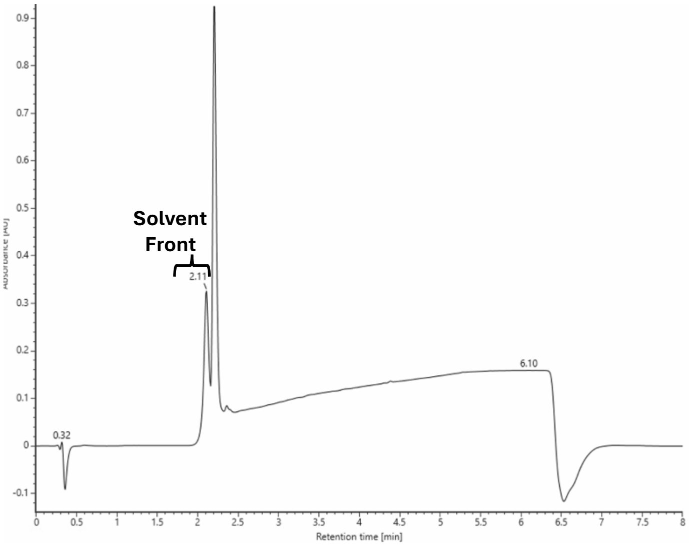

SMMP-dY-R-dQ-V

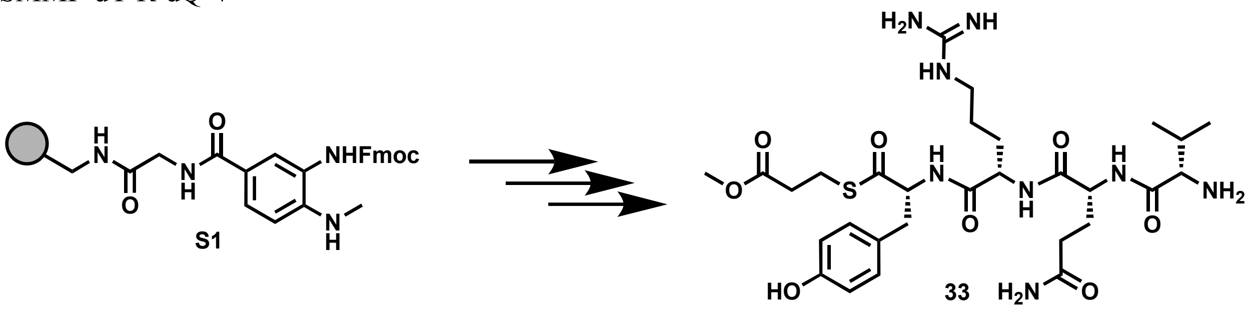

Peptide **33** was synthesized following the peptide synthesis protocol outlined above. Starting with Fmoc-Me-Dbz-OH loaded resin **S1** (0.1 mmol), amino acids Fmoc-D-Tyr(tBu)-OH, Fmoc-L-Arg(Pbf)-OH, Fmoc-D-Gln(Trt)-OH, and Boc-L-Val-OH were used with methyl 3-mercaptopropionate as the cleaving thiol. The crude peptide was purified by reverse-phase semi-preparative HPLC with mobile phases of $H_2O$ + 0.05% TFA (A) and acetonitrile + 0.05% TFA (B). The peptide was eluted using a gradient of 0-55% mobile phase B over 25 minutes at a 20 mL/min flow rate. The column was equilibrated with 0% mobile phase B for 1 minute before and 5 minutes after the gradient, yielding an off-white solid (22 mg, 25% yield).

**$^1$H NMR** (800 MHz, DMSO) δ 8.67 (t, $J$ = 6.3 Hz, 2H), 8.23 (d, $J$ = 7.8 Hz, 1H), 8.08 (s, 1H), 7.67 (t, $J$ = 5.2 Hz, 1H), 7.58-6.91 (m, 6H), 6.79 (s, 1H), 6.65 (d, $J$ = 7.9 Hz, 2H), 4.54-4.46 (m, 2H), 4.38 (dd, $J$ = 6.6, 12.1 Hz, 1H), 3.71 (t, $J$ = 5.0 Hz, 1H), 3.61 (s, 3H), 3.05-2.94 (m, 5H), 2.70 (t, $J$ = 11.7 Hz, 1H), 2.56 (dd, $J$ = 6.2, 8.8 Hz, 2H), 2.13-2.02 (m, 3H), 1.83 (sextet, $J$ = 5.8 Hz, 1H), 1.75 (sextet, $J$ = 7.2 Hz, 1H), 1.46 (sextet, $J$ = 7.3, 1H), 1.36-1.21 (m, 3H), 0.93 (dd, $J$ = 6.5, 20.1 Hz, 6H).

**$^{13}$C NMR** (201 MHz, DMSO) δ 200.38, 173.84, 172.02, 171.79, 170.81, 168.20, 157.24, 156.49, 130.54, 127.09, 115.43, 70.17, 61.19, 57.62, 52.53, 52.00, 51.90, 40.58, 36.75, 33.76, 31.64, 30.21, 29.90, 29.23, 24.97, 23.77, 18.82, 17.83.

**Mass spec:** expected neutral mass for $C_{29}H_{46}N_8O_8S$ (Da): 666.3159, observed neutral mass (Da): 666.3184, mass error (ppm): 3.7.

**UPLC Trace:** Obtained with mobile phases of $H_2O$ + 0.1% formic acid (A) and acetonitrile + 0.1% formic acid (B). The peptide was eluted using a gradient of 0-70% mobile phase B over 4 min at a 0.5 mL/min flow rate. Mobile phase B was held at 70% for 1 min. The column was equilibrated with 0% mobile phase for 1 minute before and 2 minutes after the gradient. The peptide purity was determined to be 97%.

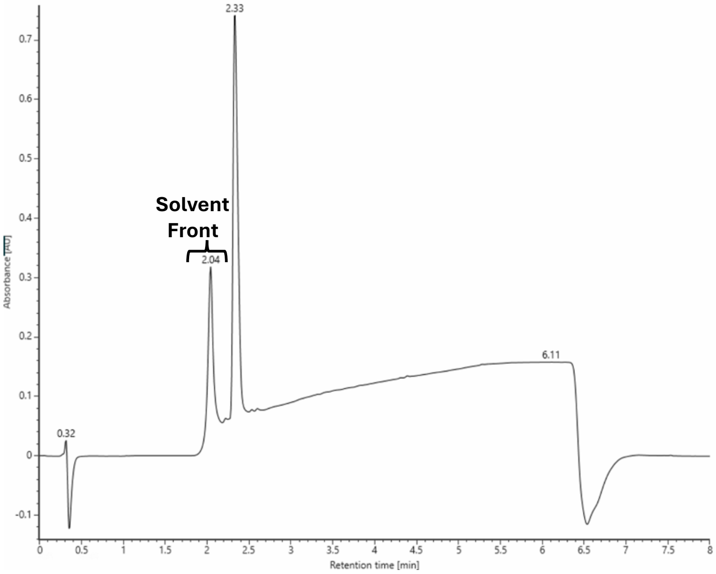

SMMP-dY-R-dE-V

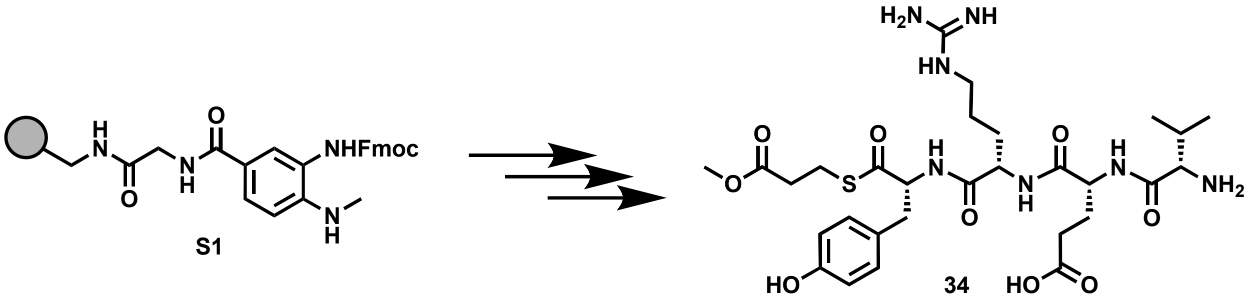

Peptide **34** was synthesized following the peptide synthesis protocol outlined above. Starting with Fmoc-Me-Dbz-OH loaded resin **S1** (0.1 mmol), amino acids Fmoc-D-Tyr(tBu)-OH, Fmoc-L-Arg(Pbf)-OH, Fmoc-D-Glu(tBu)-OH, and Boc-L-Val-OH were used with methyl 3-mercaptopropionate as the cleaving thiol. The crude peptide was purified by reverse-phase semi-preparative HPLC with mobile phases of $H_2O$ + 0.05% TFA (A) and acetonitrile + 0.05% TFA (B). The peptide was eluted using a gradient of 0-55% mobile phase B over 25 minutes at a 20 mL/min flow rate. The column was equilibrated with 0% mobile phase B for 1 minute before and 5 minutes after the gradient, yielding an off-white solid (63 mg, 70%).
**$^1$H NMR** (800 MHz, DMSO) δ 9.32 (s, 1H), 8.70 (dd, *J* = 8.2, 14.6 Hz, 2H), 8.28-7.93 (m, 3H), 7.66 (t, *J* = 4.8 Hz, 1H), 7.56-6.87 (m, 5H), 6.67 (d, *J* = 7.9 Hz, 2H), 4.56-4.47 (m, 2H), 4.38 (dd, *J* = 7.0, 13.4 Hz, 1H), 3.71 (d, *J* = 5.2 Hz, 1H), 3.36 (s, 4H), 3.04-2.94 (m, 5H), 2.70 (dd, *J* = 10.5, 13.5 Hz, 1H), 2.60-2.53 (m, 2H), 2.21 (quintet, *J* = 4.9 Hz, 2H), 2.08 (sextet, *J* = 6.4 Hz, 1H), 1.88 (dt, *J* = 5.8, 13.9 Hz, 1H), 1.78 (dt, *J* = 7.9, 13.8 Hz, 1H), 1.46 (sextet, *J* = 6.5 Hz, 1H), 1.37-1.20 (m, 3H), 0.92 (dd, *J* = 7.0, 18.5 Hz, 6H).
**$^{13}$C NMR** (201 MHz, DMSO) δ 200.44, 174.10, 172.09, 171.87, 170.68, 168.37, 157.29, 156.57, 130.63, 127.17, 115.51, 70.25, 61.28, 57.72, 52.26, 52.09, 52.02, 40.66, 36.83, 33.84, 30.48, 30.29, 29.92, 28.77, 25.07, 23.85, 18.84, 17.89.
**Mass spec:** expected neutral mass for $C_{29}H_{45}N_7O_9S$ (Da): 667.2999, observed neutral mass (Da): 667.3025, mass error (ppm): 3.9.
**UPLC Trace:** Obtained with mobile phases of $H_2O$ + 0.1% formic acid (A) and acetonitrile + 0.1% formic acid (B). The peptide was eluted using a gradient of 0-70% mobile phase B over 4 min at a 0.5 mL/min flow rate. Mobile phase B was held at 70% for 1 min. The column was equilibrated with 0% mobile phase for 1 minute before and 2 minutes after the gradient. The peptide purity was determined to be 96%.

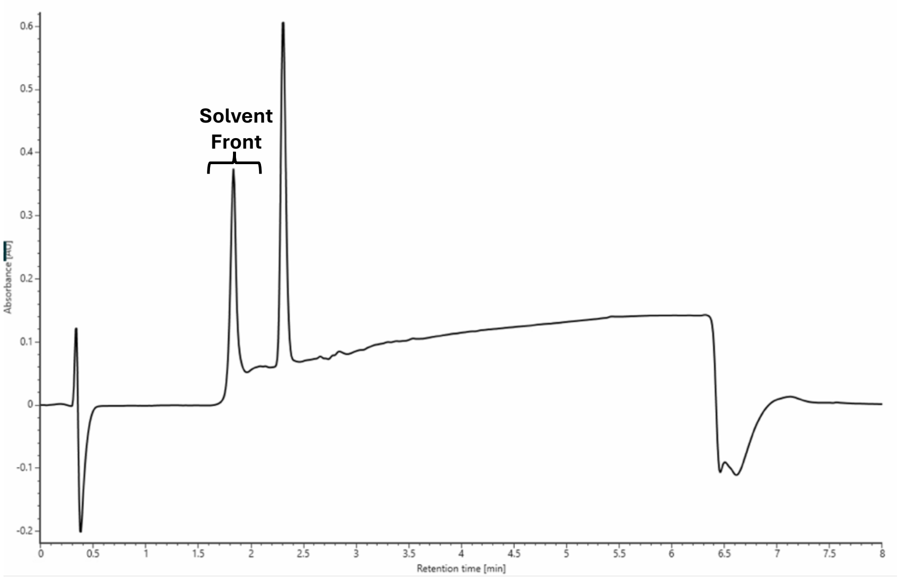

SMMP-dY-dR-F

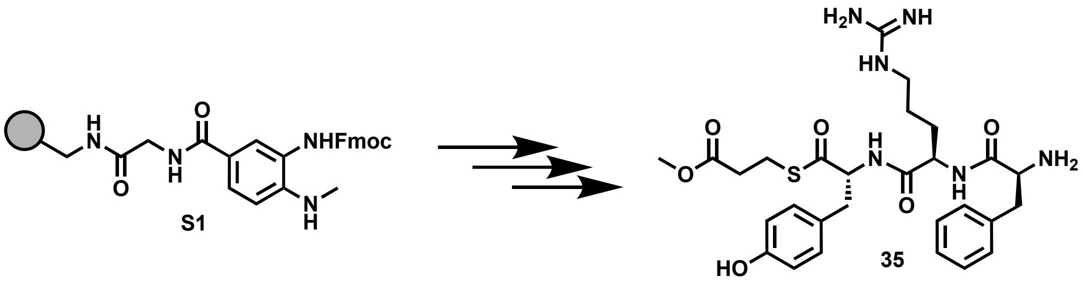

Peptide **35** was synthesized following the peptide synthesis protocol outlined above. Starting with Fmoc-Me-Dbz-OH loaded resin **S1** (0.1 mmol), amino acids Fmoc-D-Tyr(tBu)-OH, Fmoc-D-Arg(Pbf)-OH, Boc-L-Phe-OH were used with methyl 3-mercaptopropionate as the cleaving thiol. The crude peptide was purified by reverse-phase semi-preparative HPLC with mobile phases of $H_2O$ + 0.05% TFA (A) and acetonitrile + 0.05% TFA (B). The peptide was eluted using a gradient of 0-55% mobile phase B over 25 minutes at a 20 mL/min flow rate. The column was equilibrated with 0% mobile phase B for 1 minute before and 5 minutes after the gradient, yielding an off-white solid (13 mg, 16% yield).

**$^1$H NMR** (800 MHz, DMSO) δ 9.36 (s, 1H), 8.78 (d, $J$ = 7.3 Hz, 1H), 8.62 (d, $J$ = 8.0 Haz, 1H), 8.20 (s, 3H), 7.80 (t, $J$ = 5.2 Hz, 1H), 7.59-6.92 (m, 11H), 6.66 (d, $J$ = 8.3 Hz, 2H), 4.45 (dd, $J$ = 7.6, 13.8 Hz, 1H), 4.38 (dd, $J$ = 7.7, 12.5 Hz, 1H), 4.14 (t, $J$ = 6.2 Hz, 1H), 3.60 (s, 3H), 3.10-2.89 (m, 7H), 2.75 (dd, $J$ = 9.6, 14.0 Hz, 1H), 2.54 (t, $J$ = 6.9 Hz, 2H), 1.61 (q, $J$ = 6.5 Hz, 1H), 1.50-1.39 (m, 1H), 1.34-1.12 (m, 2H),

**$^{13}$C NMR** (201 MHz, DMSO) δ 200.52, 171.97, 171.36, 168.16, 157.25, 156.50, 135.23, 130.42, 129.85, 128.85, 127.50, 127.10, 115.52, 61.45, 53.53, 52.09, 51.97, 40.78, 37.67, 36.55, 33.80, 29.71, 24.88, 23.69.

**Mass spec:** expected neutral mass for $C_{28}H_{38}N_6O_6S$ (Da): 586.2574, observed neutral mass (Da): 586.2568, mass error (ppm): -0.9.

**UPLC Trace:** Obtained with mobile phases of $H_2O$ + 0.1% formic acid (A) and acetonitrile + 0.1% formic acid (B). The peptide was eluted using a gradient of 0-70% mobile phase B over 4 min at a 0.5 mL/min flow rate. Mobile phase B was held at 70% for 1 min. The column was equilibrated with 0% mobile phase for 1 minute before and 2 minutes after the gradient. The peptide purity was determined to be 99%.

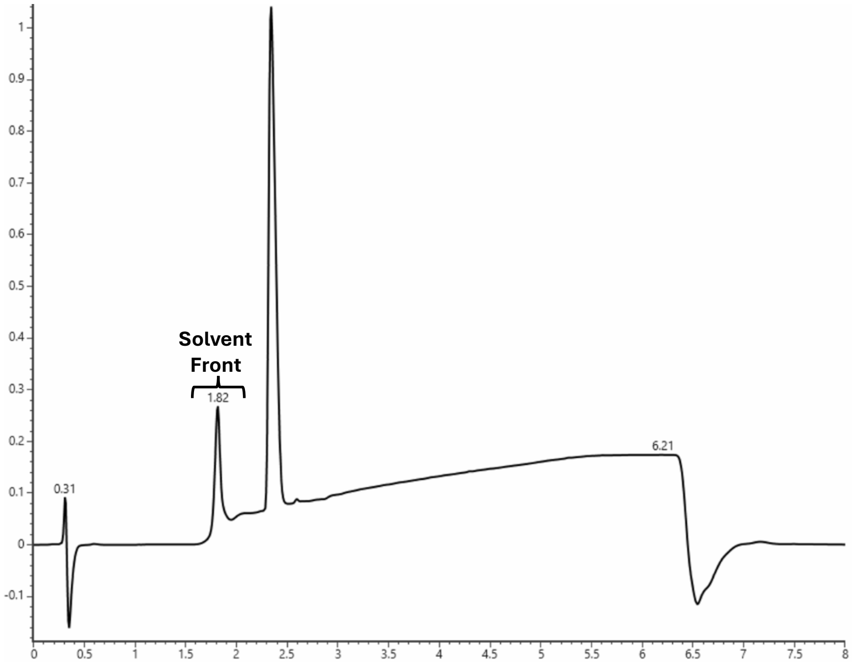

SMMP-dY-dR-dV

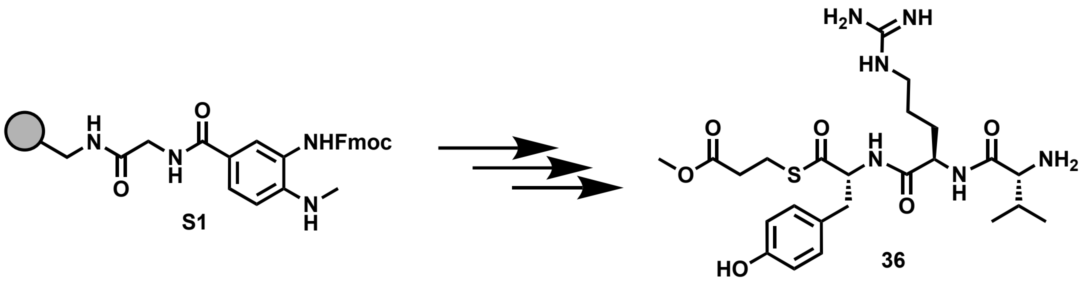

Peptide **36** was synthesized following the peptide synthesis protocol outlined above. Starting with Fmoc-Me-Dbz-OH loaded resin **S1** (0.1 mmol), amino acids Fmoc-D-Tyr(tBu)-OH, Fmoc-D-Arg(Pbf)-OH, Boc-D-Val-OH were used with methyl 3-mercaptopropionate as the cleaving thiol. The crude peptide was purified by reverse-phase semi-preparative HPLC with mobile phases of $H_2O$ + 0.05% TFA (A) and acetonitrile + 0.05% TFA (B). The peptide was eluted using a gradient of 0-55% mobile phase B over 25 minutes at a 20 mL/min flow rate. The column was equilibrated with 0% mobile phase B for 1 minute before and 5 minutes after the gradient, yielding an off-white solid (27mg, 35% yield).

**$^1$H NMR** (800 MHz, DMSO) δ 9.28 (d, 1H), 8.76 (d, *J* = 7.7 Hz, 1H), 8.51 (d, *J* = 7.7 Hz, 1H), 8.10 (s, 3H), 7.89 (t, *J* = 5.6 Hz, 1H), 7.61-6.94 (m, 5H), 6.62 (d, *J* = 8.4 Hz, 2H), 4.51 (ddd, *J* = 5.0, 7.9, 12.9 Hz, 1H), 4.41 (dd, *J* = 7.5, 13.7 Hz, 1H), 3.64-3.57 (m, 4H), 3.10 (dd, *J* = 7.6, 13.7 Hz, 2H), 3.00-2.93 (m, 3H), 2.77 (dd, *J* = 9.6, 14.5 Hz, 1H), 2.55 (t, *J* = 7.0 Hz, 2H), 1.98 (sextet, *J* = 6.6 Hz, 1H), 1.76-1.68 (m, 1H), 1.59-1.42 (m, 3H), 0.85 (t, *J* = 7.5 Hz, 6H).

**$^{13}$C NMR** (201 MHz, DMSO) δ 200.48, 171.97, 171.50, 168.03, 157.31, 156.43, 130.19, 127.05, 115.41, 61.02, 57.54, 52.40, 51.96, 40.79, 36.33, 33.78, 30.28, 29.60, 25.39, 23.73, 18.61, 17.97.

**Mass spec:** expected neutral mass for $C_{24}H_{38}N_6O_6S$ (Da): 538.2574, observed neutral mass (Da): 538.2579, mass error (ppm): 1.0.

**UPLC Trace:** Obtained with mobile phases of $H_2O$ + 0.1% formic acid (A) and acetonitrile + 0.1% formic acid (B). The peptide was eluted using a gradient of 0-70% mobile phase B over 4 min at a 0.5 mL/min flow rate. Mobile phase B was held at 70% for 1 min. The column was equilibrated with 0% mobile phase for 1 minute before and 2 minutes after the gradient. The peptide purity was determined to be 99%.

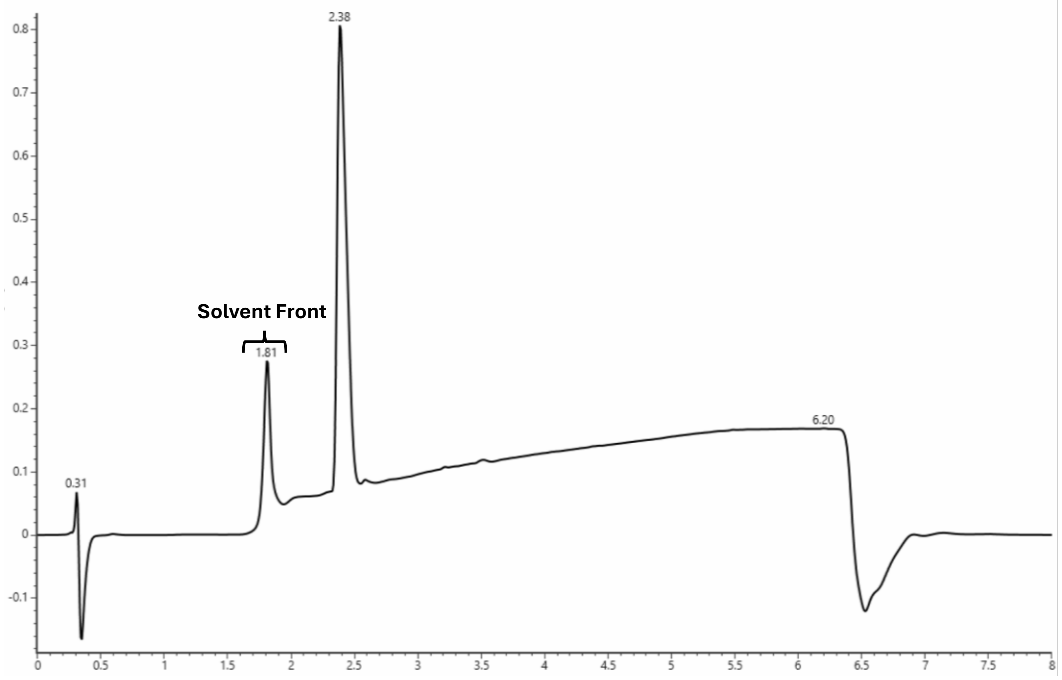

SMMP-dY-R-F

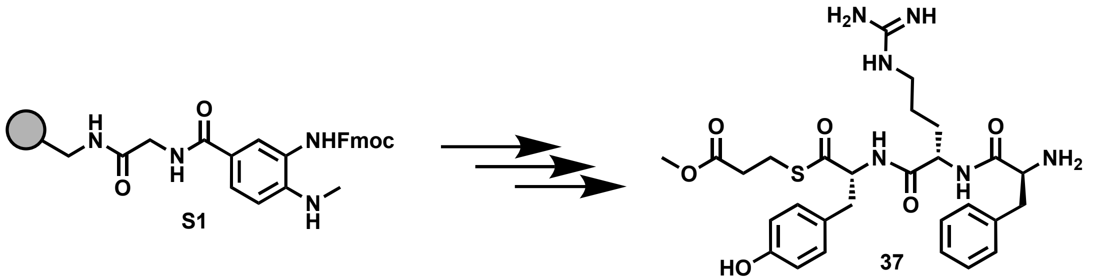

Peptide **37** was synthesized following the peptide synthesis protocol outlined above. Starting with Fmoc-Me-Dbz-OH loaded resin **S1** (0.1 mmol), amino acids Fmoc-D-Tyr(tBu)-OH, Fmoc-L-Arg(Pbf)-OH, Boc-L-Phe-OH were used with methyl 3-mercaptopropionate as the cleaving thiol. The crude peptide was purified by reverse-phase semi-preparative HPLC with mobile phases of $H_2O$ + 0.05% TFA (A) and acetonitrile + 0.05% TFA (B). The peptide was eluted using a gradient of 0-55% mobile phase B over 25 minutes at a 20 mL/min flow rate. The column was equilibrated with 0% mobile phase B for 1 minute before and 5 minutes after the gradient, yielding an off-white solid (32 mg, 39% yield)
**$^1$H NMR** (800 MHz, DMSO) δ 9.32 (s, 1H), 8.84 (d, *J* = 8.2 Hz, 1H), 8.75 (d, *J* = 8.5 Hz, 1H), 8.13 (s, 3H), 7.75 (t, *J* = 5.1 Hz, 1H), 7.60-7.00 (m, 10H), 6.65 (dd, *J* = 2.6, 8.6 Hz, 2H), 4.51 (sextet, 5.0 Hz, 1H), 4.42 (dd, *J* = 14.0, 7.5 Hz, 1H), 4.08 (t, 5.9 Hz, 1H), 3.56 (s, 3H), 3.36 (s, 2H), 3.07 (dd, *J* = 4.4, 14.1 Hz, 1H), 3.05-2.96 (m, 5H), 2.90 (dd, *J* = 7.7, 14.0 Hz, 1H), 2.71 (dd, *J* = 10.7, 13.3 Hz, 1H), 2.57 (t, *J* = 6.5 Hz, 2H), 1.47 (septet, *J* = 5.2 Hz, 1H), 1.42-1.27 (m, 2H), 1.22 (quintet, *J* = 5.8 Hz, 1H).
**$^{13}$C NMR** (201 MHz, DMSO) δ 200.45, 171.96, 171.45, 168.19, 157.29, 156.49, 135.13, 130.52, 129.94, 128.89, 127.49, 127.23, 115.42, 61.32, 53.56, 52.41, 51.94, 40.63, 37.38, 36.45, 33.80, 29.89, 25.11, 23.81.
**Mass spec:** expected neutral mass for $C_{28}H_{38}N_6O_6S$ (Da): 586.2574, observed neutral mass (Da): 586.2571, mass error (ppm): -0.4.
**UPLC Trace:** Obtained with mobile phases of $H_2O$ + 0.1% formic acid (A) and acetonitrile + 0.1% formic acid (B). The peptide was eluted using a gradient of 0-70% mobile phase B over 4 min at a 0.5 mL/min flow rate. Mobile phase B was held at 70% for 1 min. The column was equilibrated with 0% mobile phase for 1 minute before and 2 minutes after the gradient. The peptide purity was determined to be 97%.

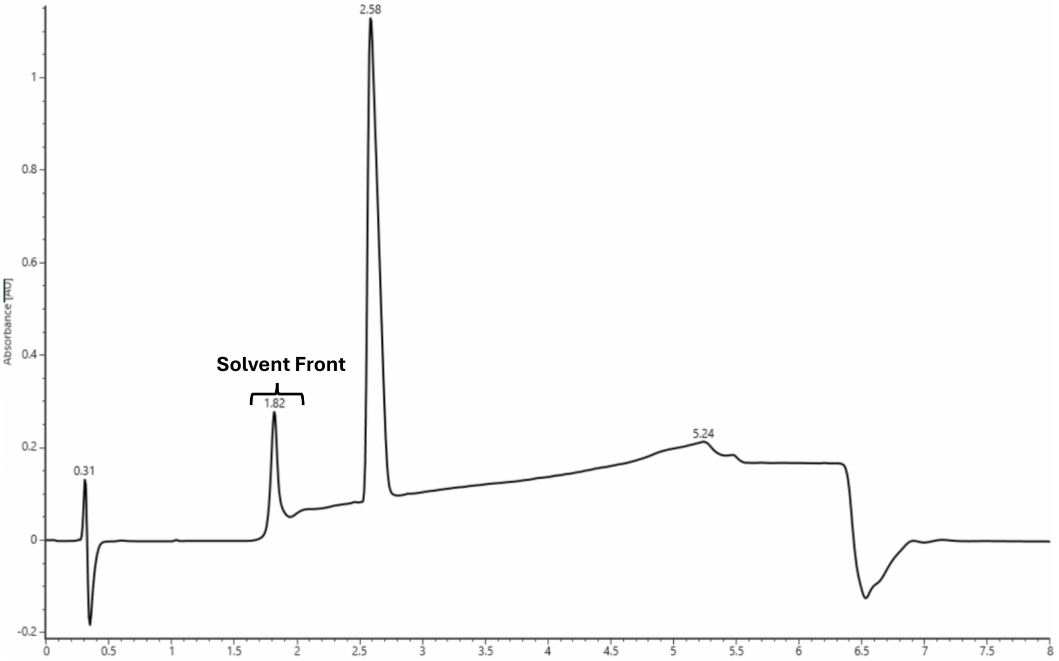

SMMP-dY-R-W-dV-V

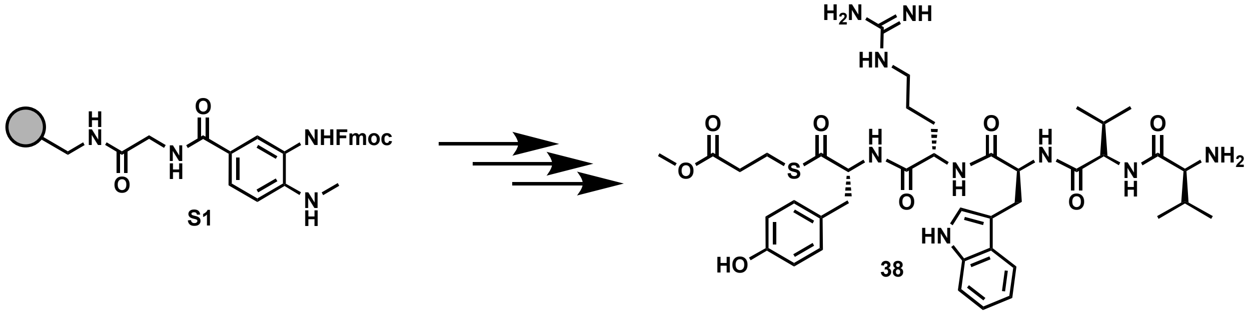

Peptide **38** was synthesized following the peptide synthesis protocol outlined above. Starting with Fmoc-Me-Dbz-OH loaded resin **S1** (0.1 mmol), amino acids Fmoc-D-Tyr(tBu)-OH, Fmoc-L-Arg(Pbf)-OH, Fmoc-L-Trp(Boc)-OH, Fmoc-D-Val-OH, Boc-L-Val-OH were used with methyl 3-mercaptopropionate as the cleaving thiol. The crude peptide was purified by reverse-phase semi-preparative HPLC with mobile phases of $H_2O$ + 0.05% TFA (A) and acetonitrile + 0.05% TFA (B). The peptide was eluted using a gradient of 0-55% mobile phase B over 25 minutes at a 20 mL/min flow rate. The column was equilibrated with 0% mobile phase B for 1 minute before and 5 minutes after the gradient, yielding an off-white solid (63 mg, 60% yield).
**$^1$H NMR** (800 MHz, DMSO) δ 10.79 (s, 1H), 8.71 (d, *J* = 8.2 Hz, 1H), 8.39 (d, *J* = 8.3 Hz, 1H), 8.29 ( d, *J* = 8.3 Hz, 1H), 8.25 (d, *J* = 7.4 Hz, 1H), 8.07-8.00 (m, 3H), 7.79 (t, *J* = 5.1 Hz, 1H), 7.68 (d, *J* = 8.1 Hz, 1H), 7.55-6.99 (m, 8H), 6.96 (t, *J* = 7.6 Hz, 1H), 6.67 (d, *J* = 7.5 Hz, 2H), 4.75-4.66 (m, 1H), 4.50 (dd, *J* = 4.9, 8.9, 1H), 4.37 (dd, *J* = 5.9, 11.7 Hz, 3H), 3.77 (t, *J* = 4.8 Hz, 1H), 3.16 (d, *J* = 14.3 Hz, 1H), 3.07-2.95 (m, 5H), 2.86 (t, *J* = 12.2 Hz, 1H), 2.73 (t, *J* = 12.2 Hz, 1H), 2.57 (t, *J* = 7.1 Hz, 2H), 2.07 (dt, *J* = 6.7, 12.3 Hz, 1H), 1.66 (sextet, *J* = 6.1 Hz, 1H), 1.59-1.51 (m, 1H), 1.45-1.22 (m, 3H), 0.94 (d, *J* = 6.9 Hz, 3H), 0.89 (d, *J* = 6.9 Hz, 3H), 0.55 (d, *J* = 7.0 Hz, 3H), 0.37 (d, *J* = 6.3 Hz, 3H).
**$^{13}$C NMR** (201 MHz, DMSO) δ 200.62, 172.20, 172.14, 172.04, 170.39, 168.48, 157.38, 156.54, 136.69, 130.59, 127.57, 127.35, 124.56, 121.21, 115.52, 111.60, 110.32, 61.41, 57.64, 57.53, 53.41, 52.56, 52.02, 40.77, 36.51, 33.85, 31.57, 30.41, 29.61, 28.87, 25.38, 23.87, 19.56, 18.95, 17.50, 17.38.
**Mass spec:** expected neutral mass for $C_{40}H_{57}N_9O_8S$ (Da): 823.4051, observed neutral mass (Da): 823.4091, mass error (ppm): 4.9.
**UPLC Trace:** Obtained with mobile phases of $H_2O$ + 0.1% formic acid (A) and acetonitrile + 0.1% formic acid (B). The peptide was eluted using a gradient of 0-70% mobile phase B over 4 min at a 0.5 mL/min flow rate. Mobile phase B was held at 70% for 1 min. The column was equilibrated with 0% mobile phase for 1 minute before and 2 minutes after the gradient. The peptide purity was determined to be 96%.

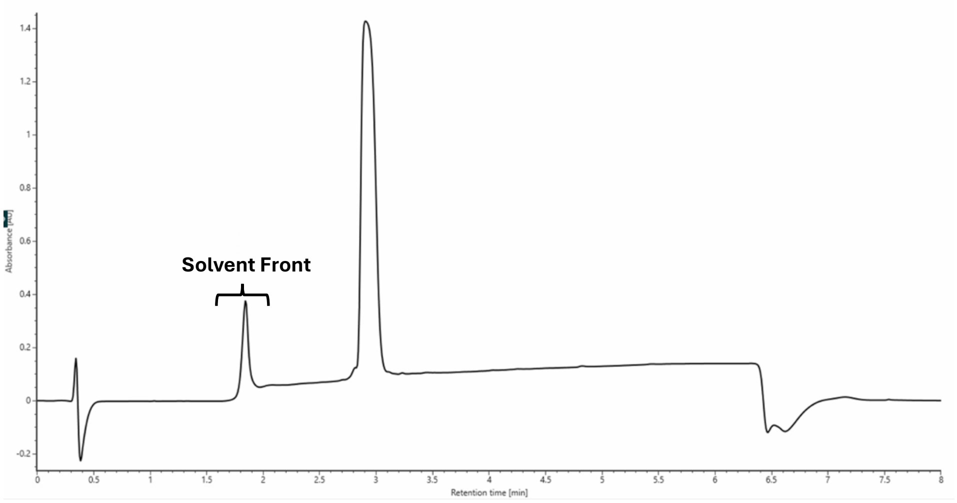

| | Sequence | | | | | | | | | | |
|---|---|---|---|---|---|---|---|---|---|---|---|
| Substrate | 1 | 2 | 3 | 4 | 5 | 6 | Thiol | [Peptide] | [WP516] | TTN | Cyc:Hyd |
| 1 | DY | LR | DF | LV | - | - | SBMP | 400 μM | 33 nM | 11219 | 47.4 |
| 2 | DY | DR | DF | LV | - | - | SBMP | 400 μM | 53 nM | 3873 | 38.7 |
| 3 | DY | DR | LF | LV | - | - | SBMP | 400 μM | 221 nM | 1753 | 477 |
| 4 | DY | LR | LF | LV | - | - | SBMP | 400 μM | 155 nM | 1332 | 2.6 |
| 5 | DY | LDap | DF | LV | - | - | SBMP | 400 μM | 125 nM | 2901 | 26.4 |
| 6 | DY | DDap | DF | LV | - | - | SBMP | 400 μM | 125 nM | 2292 | 29.1 |
| 7 | DY | DDap | LF | LV | - | - | SBMP | 400 μM | 125 nM | 1031 | 34 |
| 8 | DY | LDap | LF | LV | - | - | SBMP | 600 μM | 177 nM | 674 | 0.9 |
| 9 | DQ | LR | DF | LV | - | - | SMMP | 400 μM | 221 nM | 1145 | 16.7 |
| 10 | DS | LR | DF | LV | - | - | SMMP | 400 μM | 221 nM | 1501 | 15.4 |
| 11 | DK | LR | DF | LV | - | - | SMMP | 400 μM | 221 nM | 259 | 3.3 |
| 12 | DF | LR | DF | LV | - | - | SBMP | 400 μM | 53 nM | 7183 | 51.4 |
| 13 | DV | LR | DF | LV | - | - | SMMP | 400 μM | 221 nM | ND | ND |
| 14 | DE | LR | DF | LV | - | - | SMMP | 400 μM | 250 nM | ND | ND |
| 15 | G | LR | DF | LV | - | - | SMMP | 400 μM | 250 nM | 714 | 7.0 |
| 16 | G | LR | LF | LV | - | - | SMMP | 400 μM | 4 μM | ND | ND |
| 17 | DY | LR | DF | LP | - | - | SMMP | 400 μM | 250 nM | ND | ND |
| 18 | DY | LR | DF | LH | - | - | SMMP | 400 μM | 250 nM | ND | ND |
| 19 | DY | LR | DF | LE | - | - | SMMP | 400 μM | 250 nM | ND | ND |
| 20 | DY | LR | DF | LY | - | - | SMMP | 400 μM | 101 nM | 3123 | 6.6 |
| 21 | DY | LR | DF | LT | - | - | SMMP | 400 μM | 53 nM | 6554 | 24.5 |
| 22 | DY | DR | DF | G | - | - | SMMP | 400 μM | 315 nM | 383 | 0.8 |
| 23 | DY | LR | DF | G | - | - | SMMP | 400 μM | 315 nM | 266 | 0.4 |
| 24 | DY | LA | DF | LV | - | - | SMMP | 400 μM | 1.4 μM | 142 | 1.5 |
| 25 | DY | LW | DF | LV | - | - | SMMP | 600 μM | 1.4 μM | 24 | 1.6 |
| 26 | DY | LS | DF | LV | - | - | SMMP | 400 μM | 1.4 μM | 218 | 5.5 |
| 27 | DY | LQ | DF | LV | - | - | SMMP | 400 μM | 500 nM | 120 | ND |
| 28 | DY | LE | DF | LV | - | - | SMMP | 400 μM | 53 nM | 6330 | 19.6 |
| 29 | DY | LR | DT | LV | - | - | SMMP | 400 μM | 53 nM | 6346 | 28.7 |
| 30 | DY | LR | DW | LV | - | - | SMMP | 400 μM | 53 nM | 7268 | 47.8 |
| 31 | DY | LR | DA | LV | - | - | SMMP | 400 μM | 53 nM | 7281 | 15 |
| 32 | DY | LR | DDap | LV | - | - | SMMP | 400 μM | 53 nM | 4075 | ND |
| 33 | DY | LR | DQ | LV | - | - | SMMP | 400 μM | 30 nM | 10307 | 119 |
| 34 | DY | LR | DE | LV | - | - | SMMP | 400 μM | 250 nM | ND | 48.4 |
| 35 | DY | DR | LF | - | - | - | SMMP | 400 μM | 221 nM | ND | ND |
| 36 | DY | DR | DV | - | - | - | SMMP | 400 μM | 1.4 μM | ND | ND |
| 37 | DY | LR | LF | - | - | - | SMMP | 400 μM | 1.4 μM | ND | ND |
| 38 | DY | LR | LW | DV | LV | - | SMMP | 400 μM | 250 nM | 1526 | 45.2 |
| 39 | DQ | LOrn | LI | LL | DV | LW | SBMP | 400 μM | 250 nM | ND | ND |
| 40 | G | LN | LA | DL | LL | LW | SMMP | 400 μM | 250 nM | ND | ND |

**SI Table 1 Total Turnover Numbers (TTNs) of WP516 with peptide thioesters:** Reactions were conducted in 20 mM Tris buffer at pH 8.0 with 5% DMSO for 4 hours at 20 °C for 4 hours. The concentrations of peptides and WP516 for each reaction are indicated. SBMP: butyl 3- mercaptopropionate; SMMP: methyl 3- mercaptopropionate. ND = not detected. All reactions were performed in triplicate.

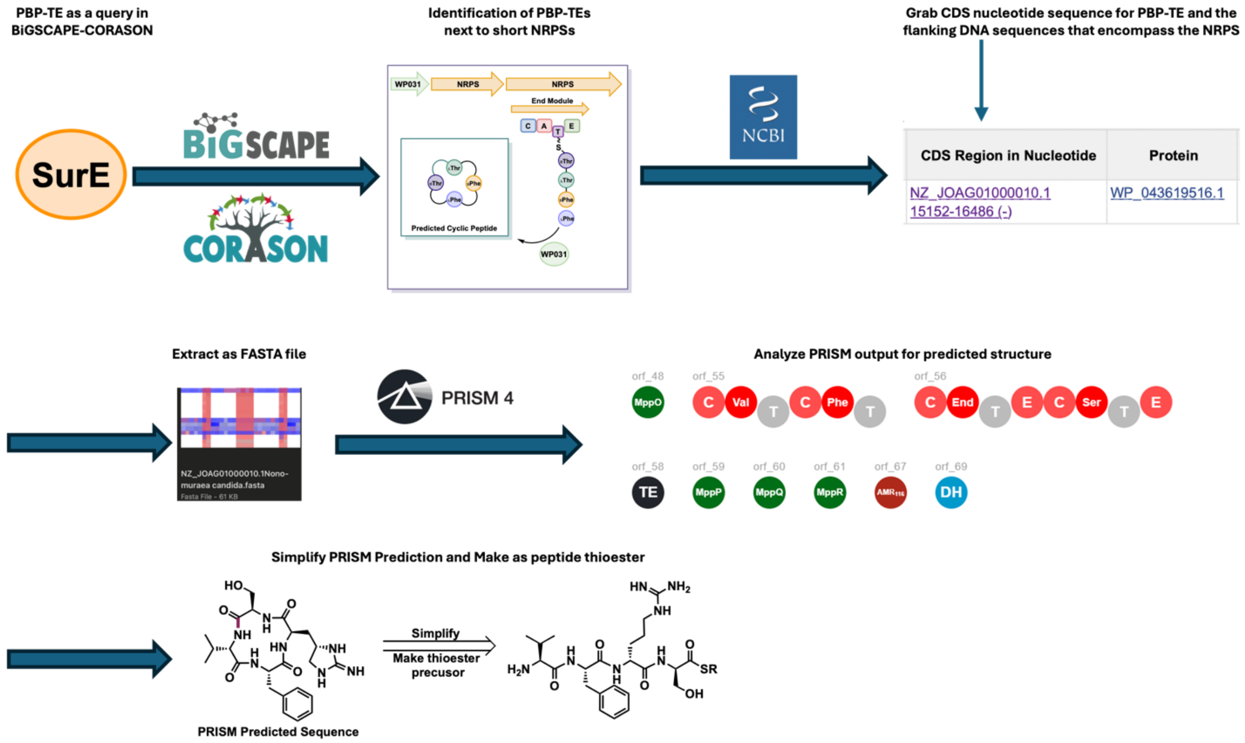

**Figure S1. Bioinformatic Workflow for PBP-TE identification and Substrate Prediction.** Graphical representation of workflow outlined in methods section.

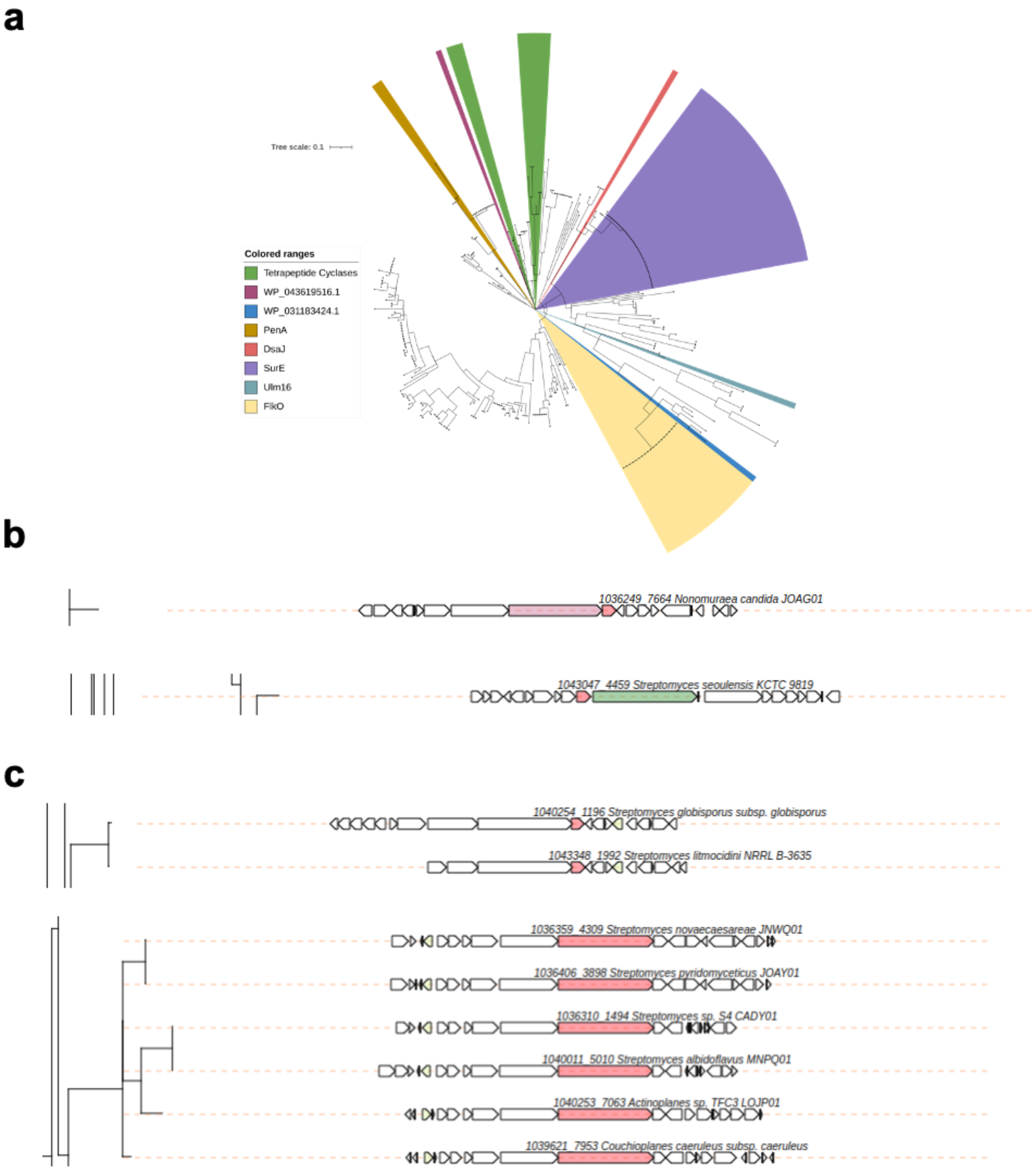

**Figure S2. Bioinformatics of PBP-TEs. A.)** Tree of BiG-SCAPE CORASON output highlighting WP_043619516.1 and WP_031183424.1 and their relationship with currently validated PBP-TEs (PenA, DsaJ, SurE, Ulm16, and FlkO). **B.)** Close up of BiG-SCAPE CORASON output for tetrapeptide PBP-TE BGCs for WP_043619516.1 (top) and WP_031183424.1 (bottom) **C.)** Close up of BiG-SCAPE CORASON of other tetrapeptide PBP-TE BGC's.

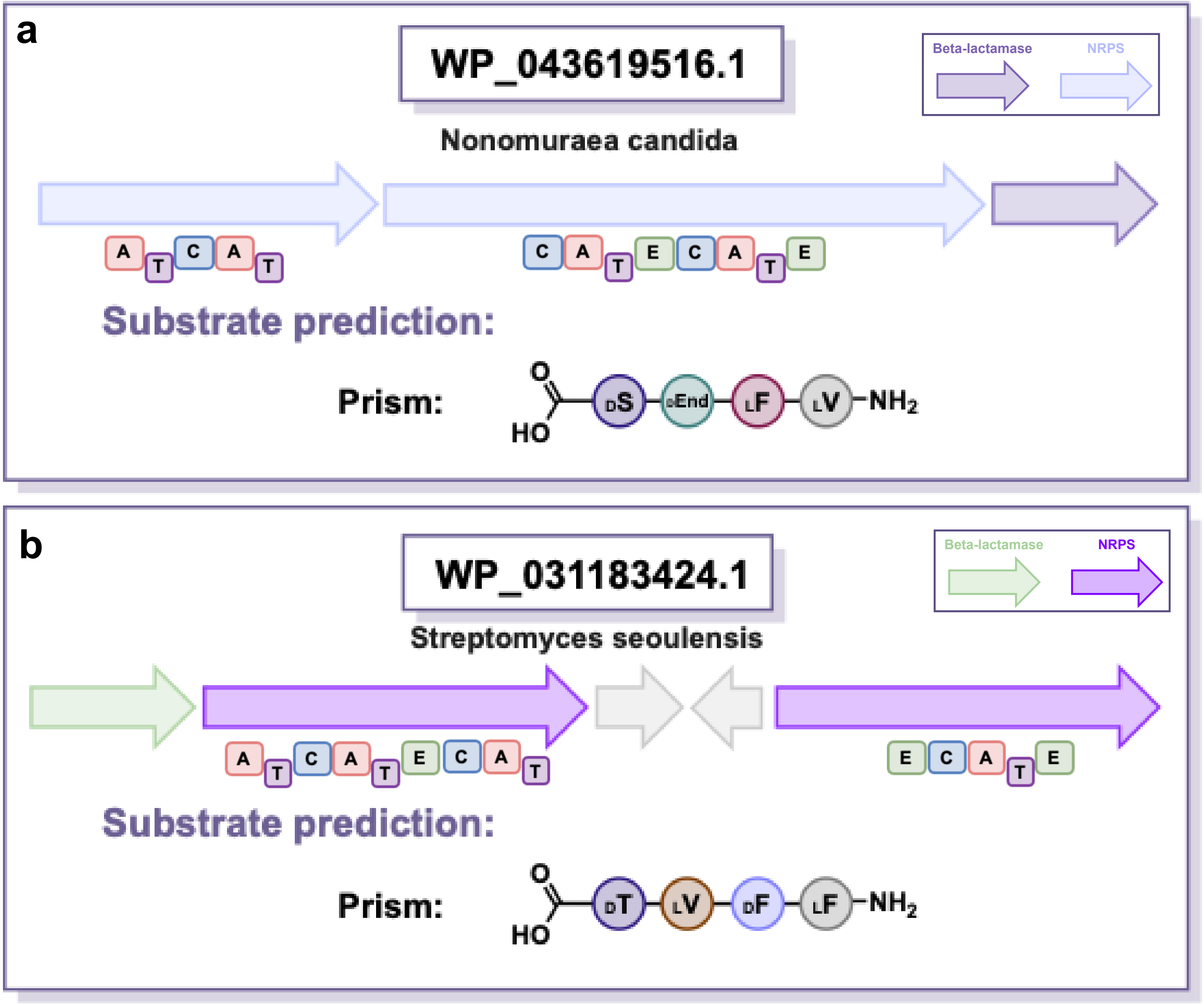

**Figure S3. WP_043619516.1 and WP_031183424 PRISM Predictions. A.)** PRISM prediction for WP_043619516.1 (WP516) from C to N-terminus the prediction consisted of D-Serine (DS), D-Enduracididine (a noncanonical amino acid resulting from a cyclized arginine, DEnd), L-Phenylalanine (LF), and L-Valine (LV) **B.)** PRISM Prediction for WP_031183424.1 from C to N-terminus the prediction consisted of D-Threonine (DT), L-Valine (LV), D-Phenylalanine (DF), and L-Phenylalanine (LF).

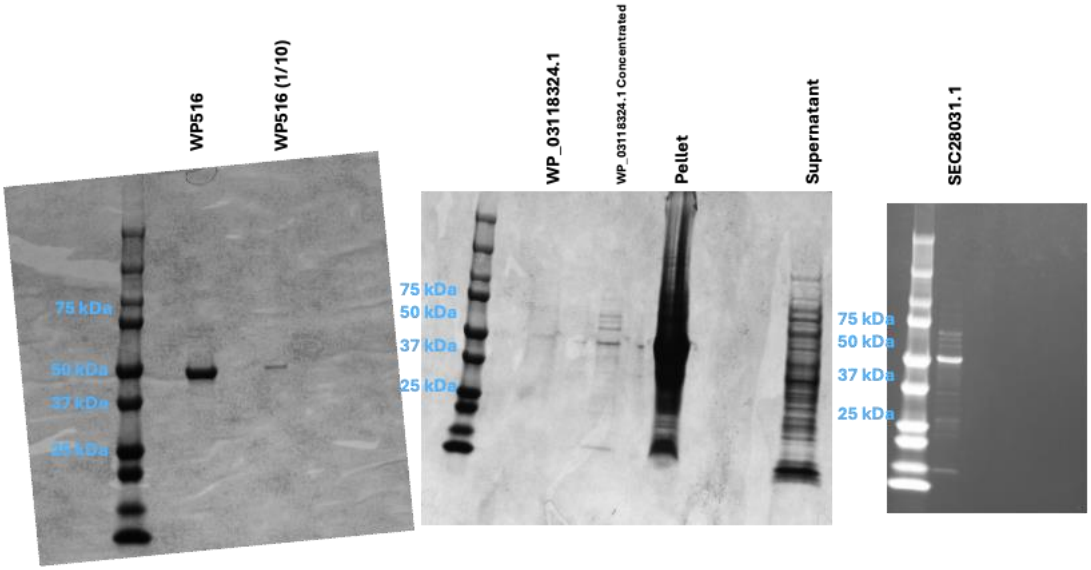

**Figure S4. Expression gel of WP516, WP_03118324.1, and SEC28031.1.** 1/10 indicates a 1/10 dilution of the initial protein stock used in the "WP516" lane. Expression of the proteins was performed once, and gels were run once. For the WP_03118324.1 gel, pellet refers to pelleted material after centrifugation. The expected molecular weight of all His-tagged proteins is 47 kDa.

**a**

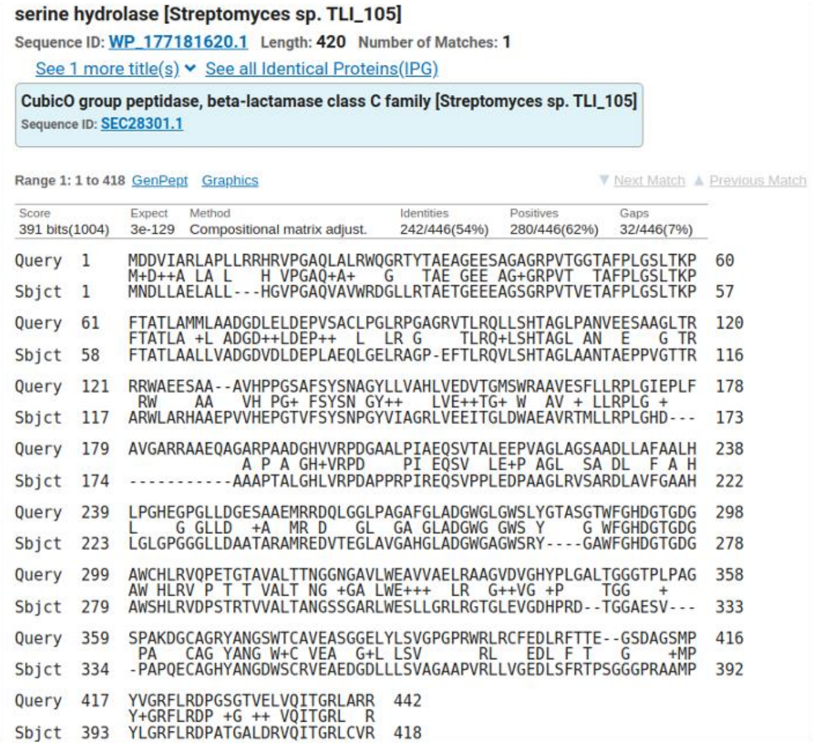
serine hydrolase [Streptomyces sp. TLI_105]
Sequence ID: WP_177181620.1 Length: 420 Number of Matches: 1
See 1 more title(s) See all Identical Proteins(IPG)
CubicO group peptidase, beta-lactamase class C family [Streptomyces sp. TLI_105]
Sequence ID: SEC28301.1
Range 1: 1 to 418 GenPept Graphics ▼ Next Match ▲ Previous Match

| Score | Expect | Method | Identities | Positives | Gaps |
|---|---|---|---|---|---|
| 391 bits(1004) | 3e-129 | Compositional matrix adjust. | 242/446(54%) | 280/446(62%) | 32/446(7%) |

```
Query  1    MDDVIARLAPLLRRHRVPGAQLALRWQGRTYTAEAGEESAGAGRPVTGGTAFPLGSLTKP  60
            M+D++A LA L    H VPGAQ+A+    G    TAE GEE AG+GRPVT   TAFPLGSLTKP
Sbjct  1    MNDLLAELALL---HGVPGAQVAVWRDGLLRTAETGEEEAGSGRPVTVETAFPLGSLTKP  57

Query  61   FTATLAMMLAADGDLELDEPVSACLPGLRPGAGRVTLRQLLSHTAGLPANVEESAAGLTR  120
            FTATLA +L ADGD++LDEP++   L  LR G    TLRQ+LSHTAGL AN  E   G TR
Sbjct  58   FTATLAALLVADGDVDLDEPLAEQLGELRAGP-EFTLRQVLSHTAGLAANTAEPPVGTTR  116

Query  121  RRWAEESAA--AVHPPGSAFSYSNAGYLLVAHLVEDVTGMSWRAAVESFLLRPLGIEPLF  178
             RW    AA   VH PG+ FSYSN GY++    LVE++TG+ W  AV + LLRPLG +
Sbjct  117  ARWLARHAAEPVVHEPGTVFSYSNPGYVIAGRLVEEITGLDWAEAVRTMLLRPLGHD---  173

Query  179  AVGARRAAEQAGARPAADGHVVRPDGAALPIAEQSVTALEEPVAGLAGSAADLLAFAALH  238
                        A P A GH+VRPD    PI EQSV  LE+P AGL  SA DL  F A H
Sbjct  174  -----------AAAPTALGHLVRPDAPPRPIREQSVPPLEDPAAGLRVSARDLAVFGAAH  222

Query  239  LPGHEGPGLLDGESAAEMRRDQLGGLPAGAFGLADGWGLGWSLYGTASGTWFGHDGTGDG  298
            L     G GLLD  +A  MR D   GL  GA GLADGWG GWS Y     G WFGHDGTGDG
Sbjct  223  LGLGPGGGLLDAATARAMREDVTEGLAVGAHGLADGWGAGWSRY----GAWFGHDGTGDG  278

Query  299  AWCHLRVQPETGTAVALTTNGGNGAVLWEAVVAELRAAGVDVGHYPLGALTGGGTPLPAG  358
            AW HLRV P T T VALT NG +GA LWE+++   LR  G++VG +P     TGG     +
Sbjct  279  AWSHLRVDPSTRTVVALTANGSSGARLWESLLGRLRGTGLEVGDHPRD--TGGAESV---  333

Query  359  SPAKDGCAGRYANGSWTCAVEASGGELYLSVGPGPRWRLRCFEDLRFTTE--GSDAGSMP  416
             PA   CAG YANG W+C VEA   G+L LSV          RL   EDL F T   G    +MP
Sbjct  334  -PAPQECAGHYANGDWSCRVEAEDGDLLLSVAGAAPVRLLVGEDLSFRTPSGGGPRAAMP  392

Query  417  YVGRFLRDPGSGTVELVQITGRLARR  442
            Y+GRFLRDP +G ++ VQITGRL  R
Sbjct  393  YLGRFLRDPATGALDRVQITGRLCVR  418
```

**b**

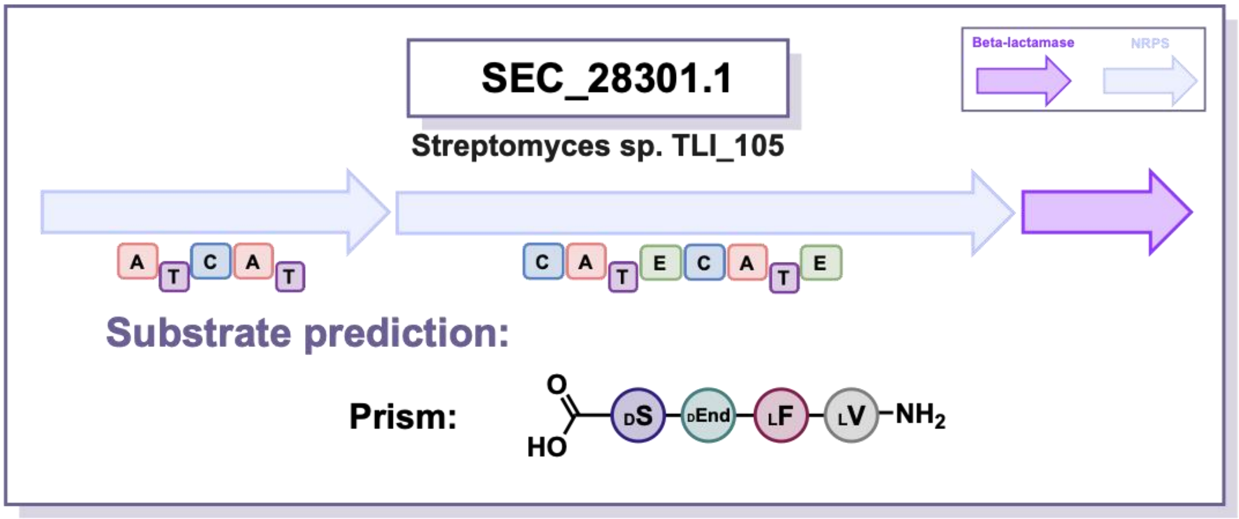

**Figure S5. Identification of PBP-TE SEC28301.1 A.)** BlastP Multiple sequence alignment of WP_043619516.1 (top) and SEC28301.1 (bottom) highlighting their similarity. **B.)** PRISM prediction for

SEC28301.1 from C to N-terminus the prediction consisted of D-Serine (DS), D-Enduracididine (DEnd), L-Phenylalanine (LF), and L-Valine (LV).

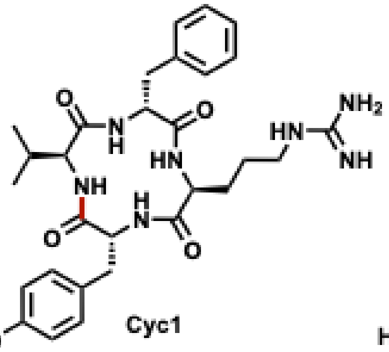
Cyc1

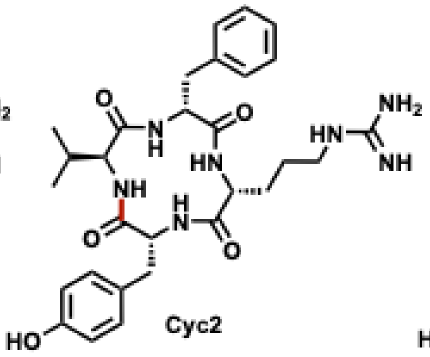
Cyc2

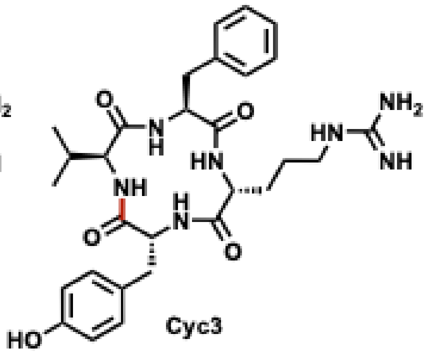
Cyc3

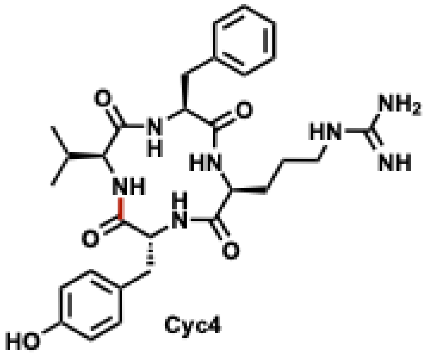
Cyc4

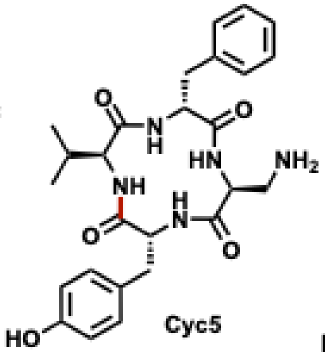
Cyc5

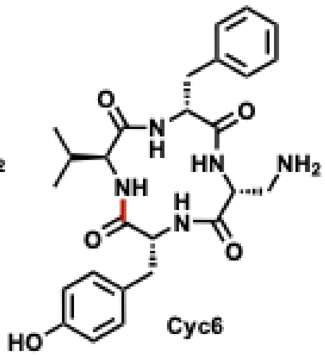
Cyc6

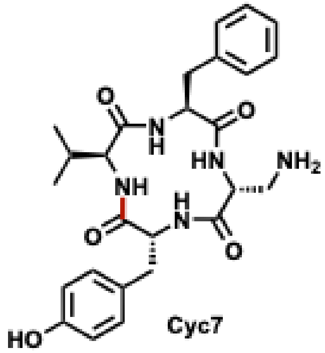
Cyc7

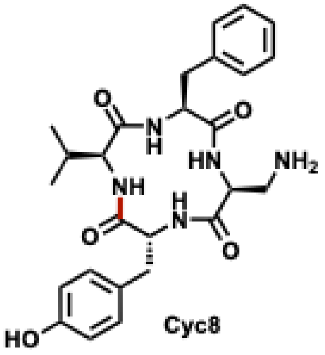
Cyc8

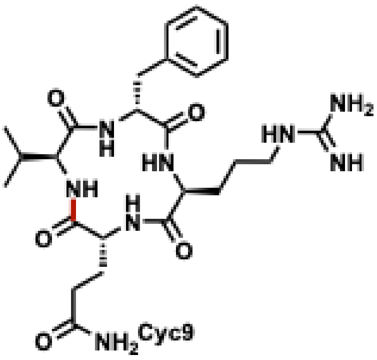
Cyc9

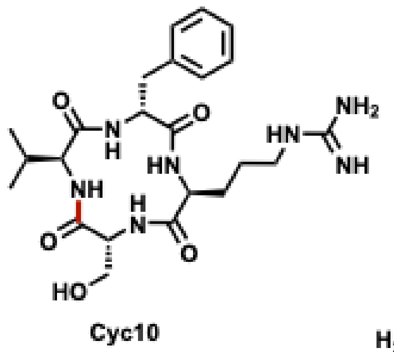
Cyc10

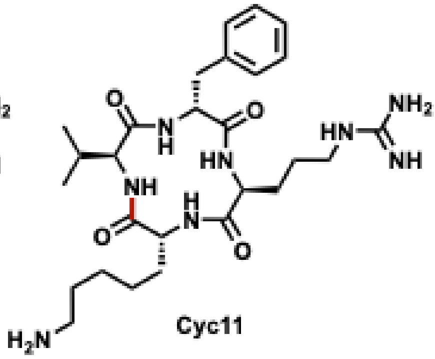
Cyc11

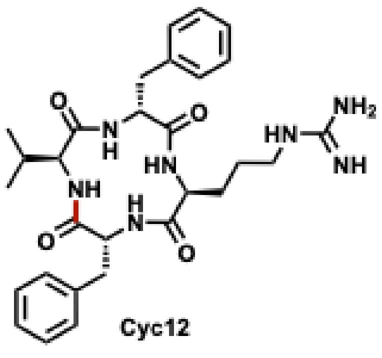
Cyc12

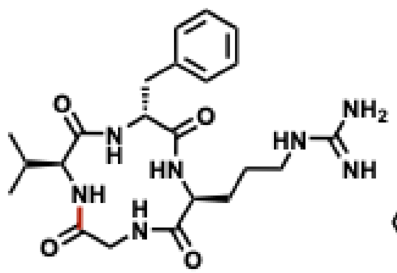

Cyc15

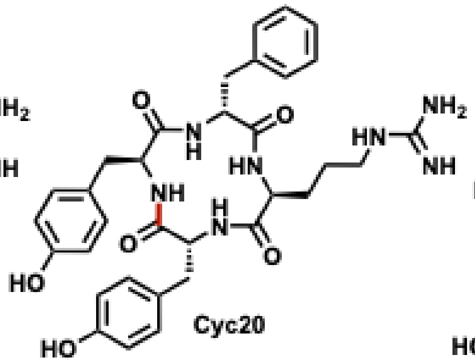

Cyc20

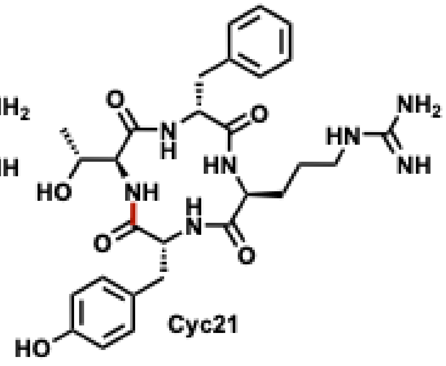

Cyc21

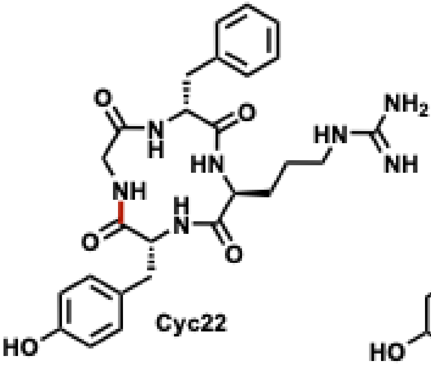

Cyc22

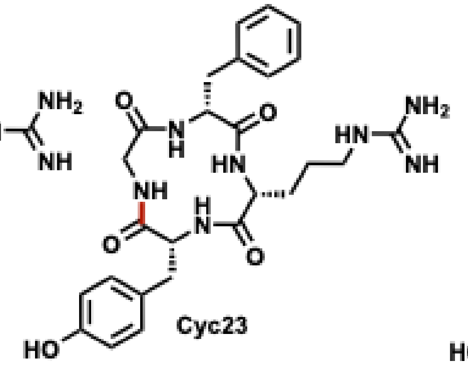

Cyc23

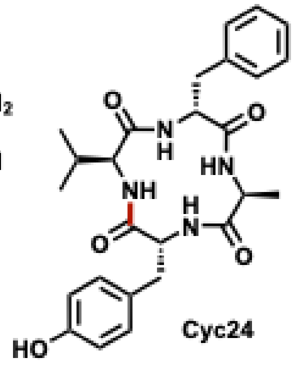

Cyc24

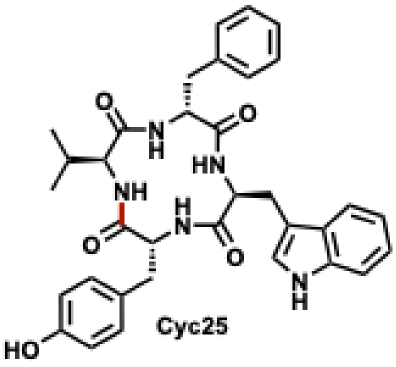

Cyc25

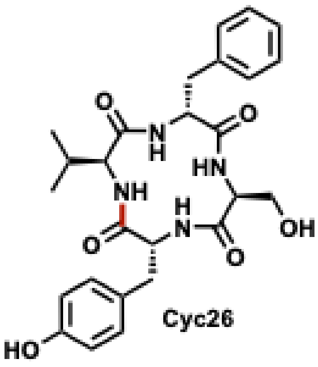

Cyc26

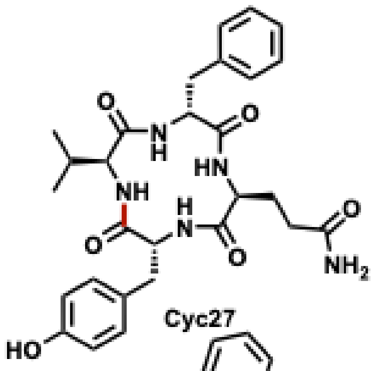

Cyc27

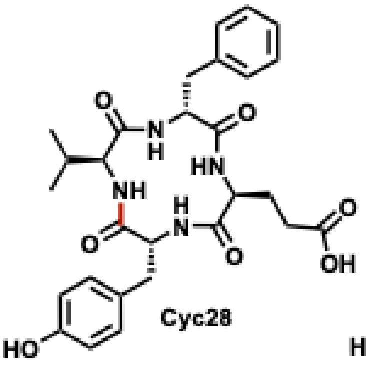

Cyc28

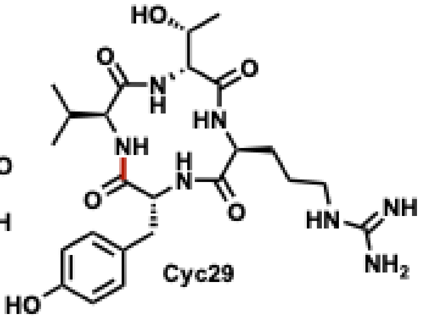

Cyc29

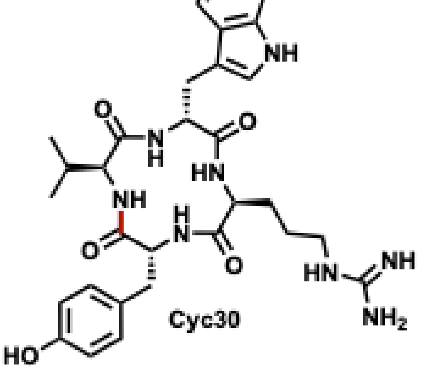

Cyc30

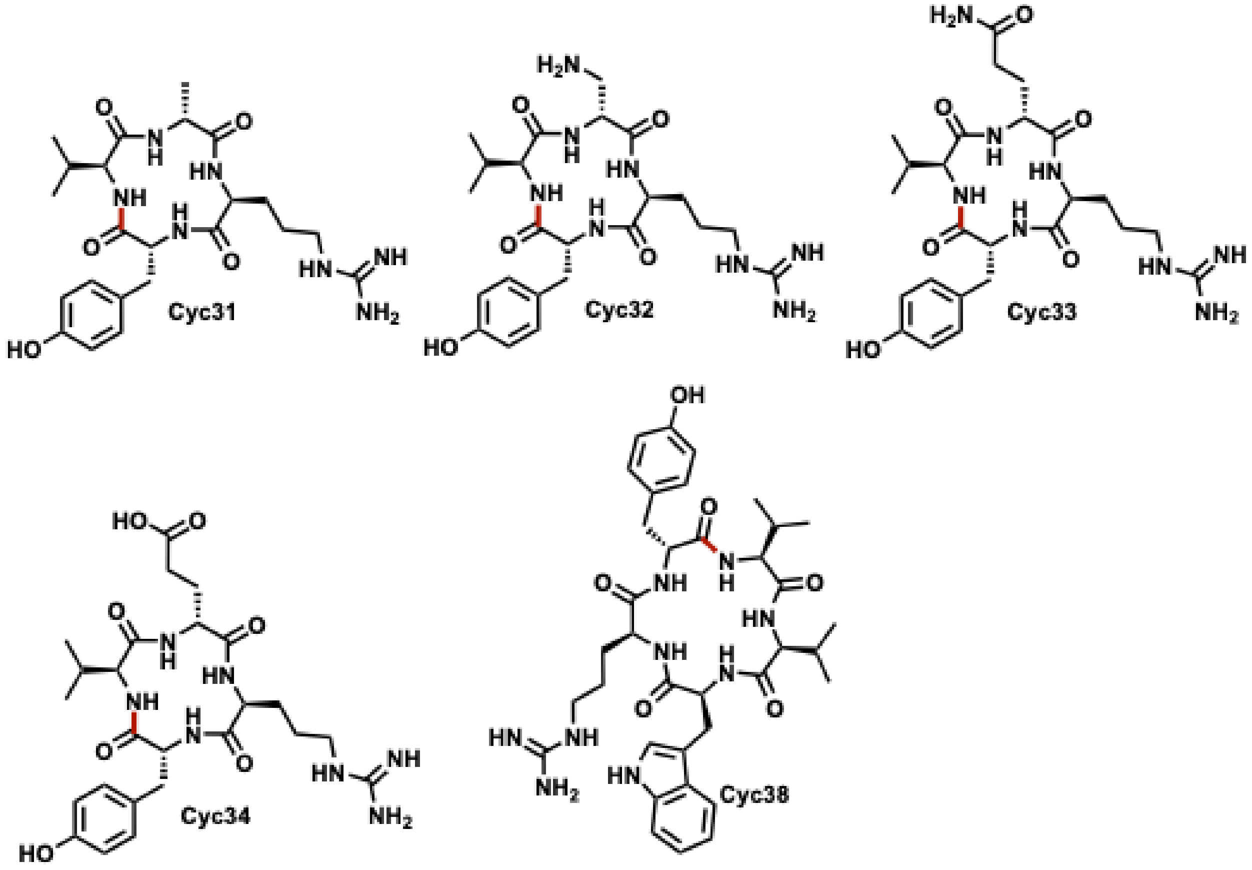

**Figure S6: Cyclic peptides identified by MS/MS analysis.** For all peptides point of cyclization is colored in maroon. Actual MS/MS data can be found in figures S85-S112.

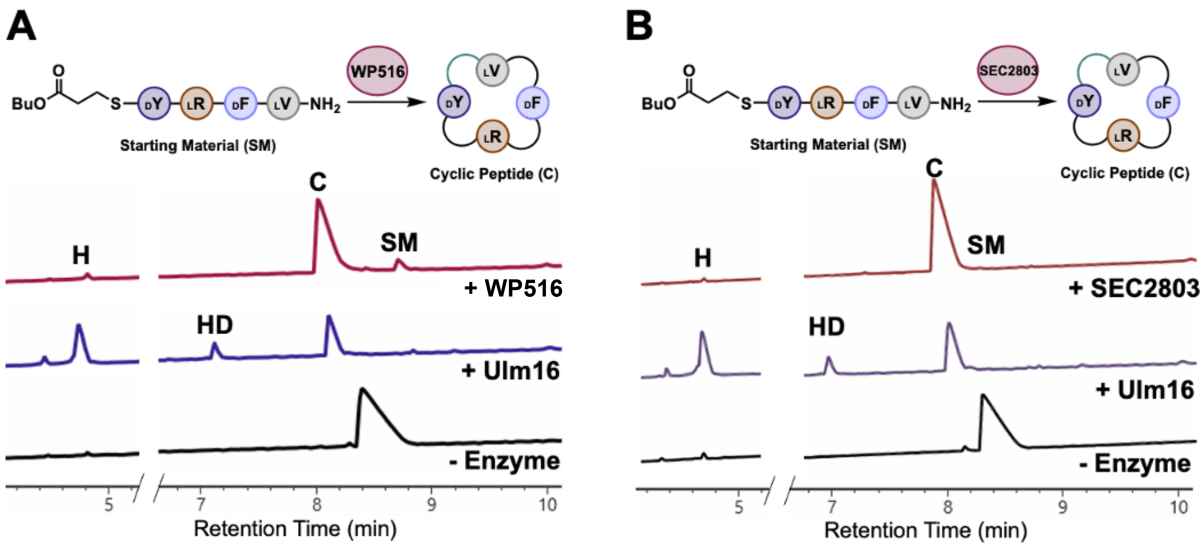

**Figure S7. Initial testing of PBP-TEs (A) WP516 and (B) SEC28301.1 with tetrapeptide substrate 1.** And their comparison with Ulm16 UPLC UV trace (214 nm). The substrate is labeled with its corresponding number from the paper where 'C', 'H', and 'HD' represent head-to-tail cyclic, hydrolyzed peptide, and hydrolyzed dimer, respectively. The shown UPLC trace is representative of three replicates.

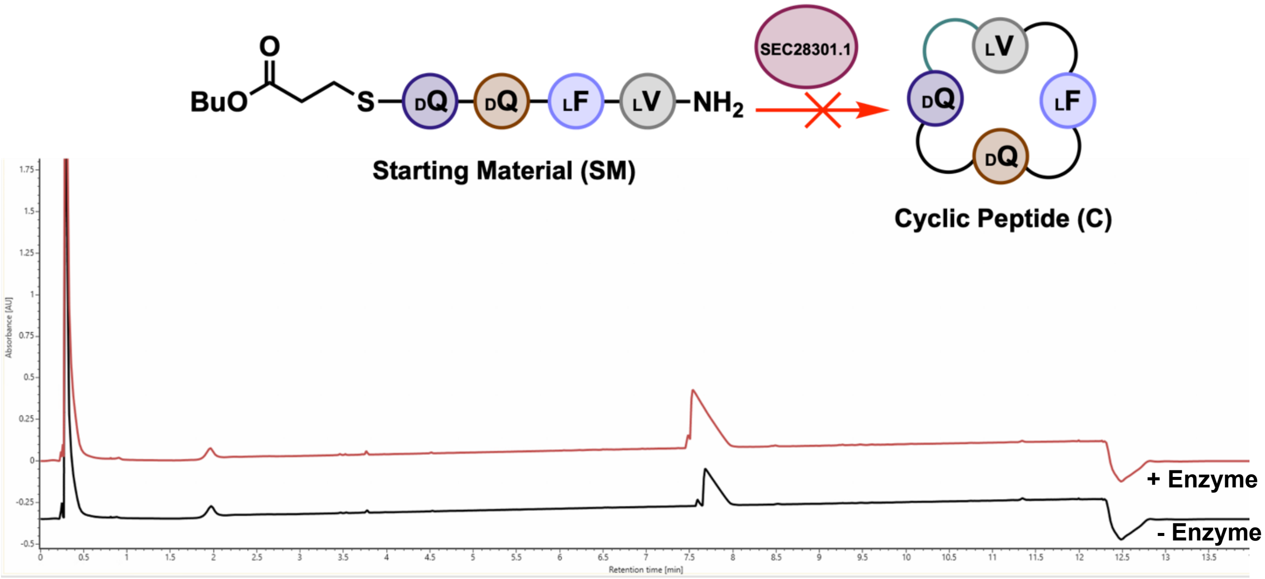

**Figure S8. UPLC UV trace (214 nm) is shown for the total turnover number (TTN) assay of SEC28301.1 with tetrapeptide substrate dQ-dQ-F-V.** A final concentration 250 nM SEC28301.1 and 320 µM substrate was used. The substrate and its synthesis was previously reported in a previous paper[6] No reaction was observed when compared to the no enzyme control. Similar results were observed when tested with other substrates bearing a C-Terminal glutamine or glutamate. For this reason, we chose not to fully explore its catalytic potential. The shown UPLC trace (214 nm) is representative of three replicates.

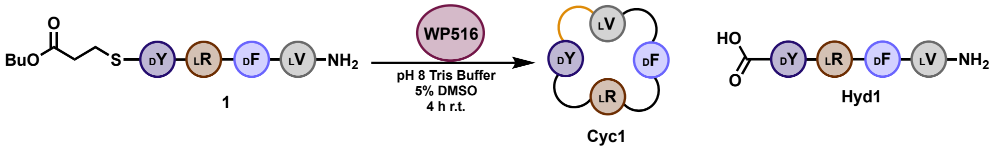

| Enzyme | Peptide | Area Peptide Thioester | Area Cyclic Peptide | Area Hydrolyzed | Sum | TTN | Product/Starting Material | Product/Hydrolysis |
|---|---|---|---|---|---|---|---|---|
| 33 nM | 400 uM | 83752 | 1411911 | 29804 | 1525468 | 11219 | 16.9 | 47.4 |

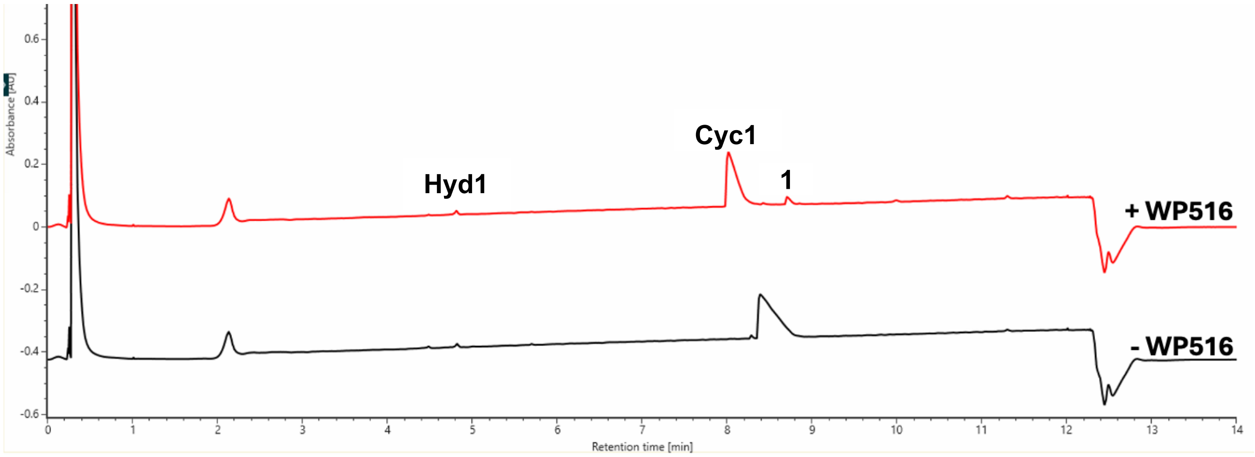

**Figure S9. UPLC UV trace (214 nm) is shown for the total turnover number (TTN) assay of WP516 with tetrapeptide substrate 1.** Enzyme and substrate concentrations are indicated above the trace. The substrate is labeled with its corresponding number from the paper where 'Cyc' and 'Hyd' represent head-to-tail cyclic and hydrolyzed peptide, respectively. The shown UPLC trace is representative of three replicates while the TTN, product/starting material, and product/hydrolysis are averages of three replicates.

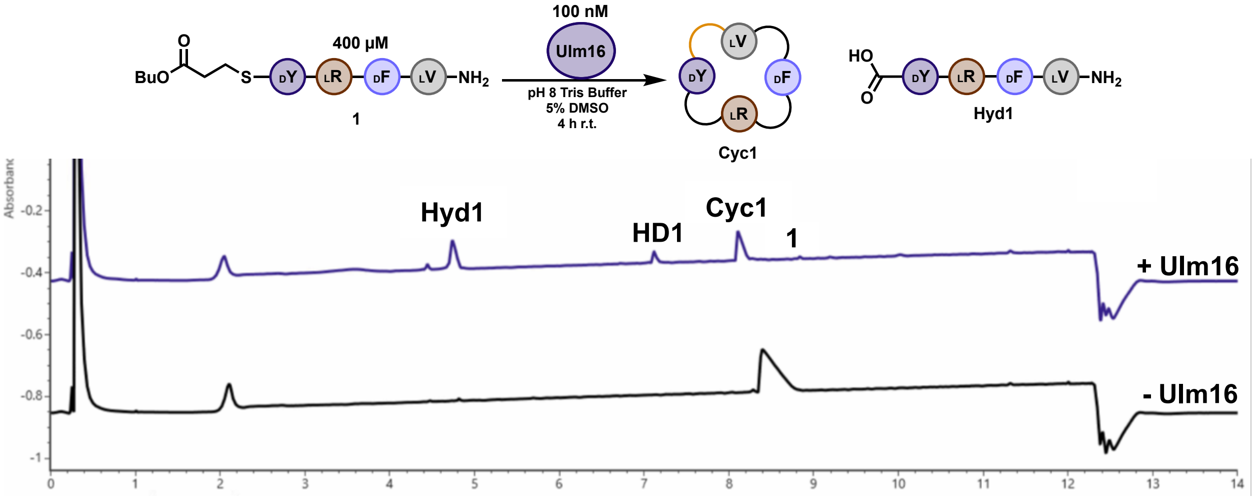

**Figure S10. UPLC UV trace (214 nm) is shown for the reaction of Ulm16 with tetrapeptide substrate 1.** Enzyme and substrate concentrations are indicated above the trace. The substrate is labeled with its corresponding number from the paper where 'Cyc' and 'Hyd' represent head-to-tail cyclic and hydrolyzed peptide, respectively. The shown UPLC trace is representative of three replicates.

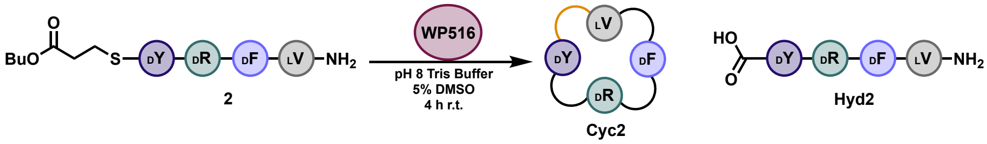

| Enzyme | Peptide | Area Peptide Thioester | Area Cyclic Peptide | Area Hydrolyzed | Sum | TTN | Product/Starting Material | Product/Hydrolysis |
|---|---|---|---|---|---|---|---|---|
| 53 nM | 400 µM | 495327 | 536766 | 13875 | 1045968 | 3873 | 1.1 | 38.7 |

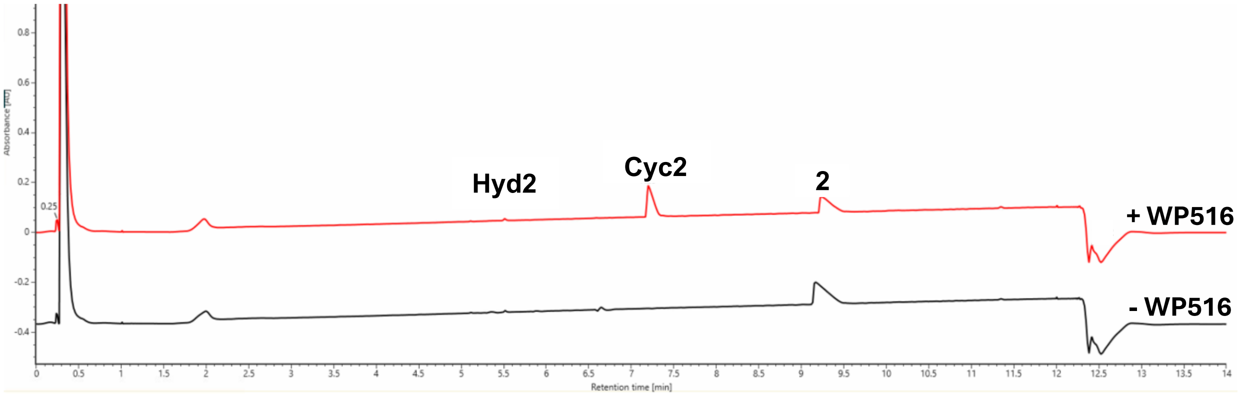

**Figure S11. UPLC UV trace (214 nm) is shown for the total turnover number (TTN) assay of WP516 with tetrapeptide substrate 2.** Enzyme and substrate concentrations are indicated above the trace. The substrate is labeled with its corresponding number from the paper where ‘Cyc’ and ‘Hyd’ represent head-to-tail cyclic and hydrolyzed peptide, respectively. The shown UPLC trace is representative of three replicates while the TTN, product/starting material, and product/hydrolysis are averages of three replicates. Note: In the +WP516 sample, there is less hydrolysis than in the no enzyme control.

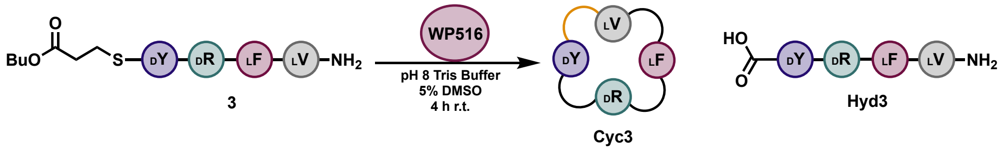

| Enzyme | Peptide | Area Peptide Thioester | Area Cyclic Peptide | Area Hydrolyzed | Sum | TTN | Product/Starting Material | Product/Hydrolysis |
|---|---|---|---|---|---|---|---|---|
| 221 nM | 400 µM | 50227 | 1645290 | 3447 | 1698964 | 1753 | 32.8 | 477.3 |

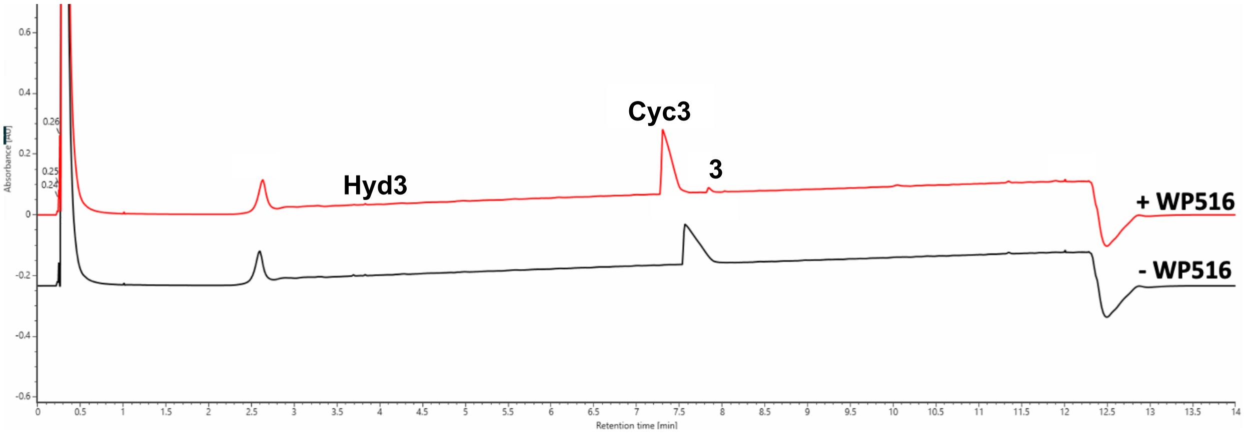

**Figure S12. UPLC UV trace (214 nm) is shown for the total turnover number (TTN) assay of WP516 with tetrapeptide substrate 3.** Enzyme and substrate concentrations are indicated above the trace. The substrate is labeled with its corresponding number from the paper where 'Cyc' and 'Hyd' represent head-to-tail cyclic and hydrolyzed peptide, respectively. The shown UPLC trace is representative of three replicates while the TTN, product/starting material, and product/hydrolysis are averages of three replicates.

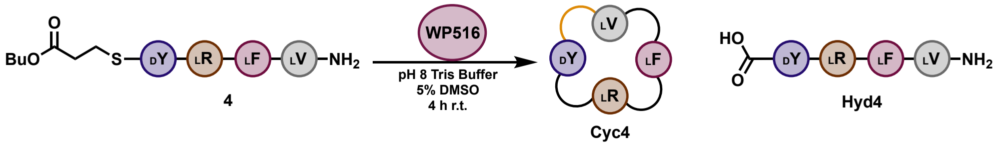

| Enzyme | Peptide | Area Peptide Thioester | Area Cyclic Peptide | Area Hydrolyzed | Sum | TTN | Product/Starting Material | Product/Hydrolysis |
|---|---|---|---|---|---|---|---|---|
| 155 nM | 400 uM | 385098 | 702006 | 273476 | 1360580 | 1332 | 1.8 | 2.6 |

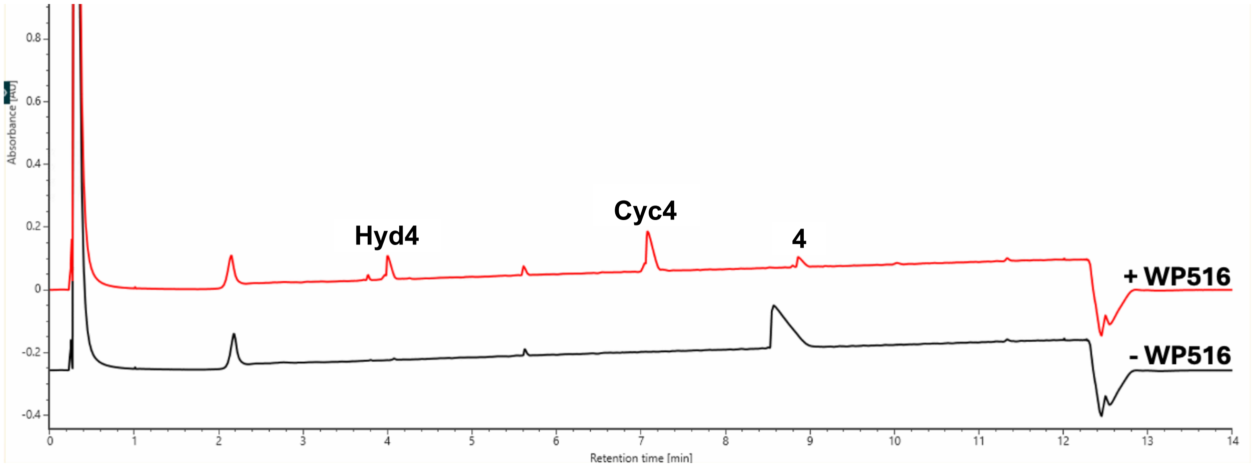

**Figure S13. UPLC UV trace (214 nm) is shown for the total turnover number (TTN) assay of WP516 with tetrapeptide substrate 4.** Enzyme and substrate concentrations are indicated above the trace. The substrate is labeled with its corresponding number from the paper where 'Cyc' and 'Hyd' represent head-to-tail cyclic and hydrolyzed peptide, respectively. The shown UPLC trace is representative of three replicates while the TTN, product/starting material, and product/hydrolysis are averages of three replicates.

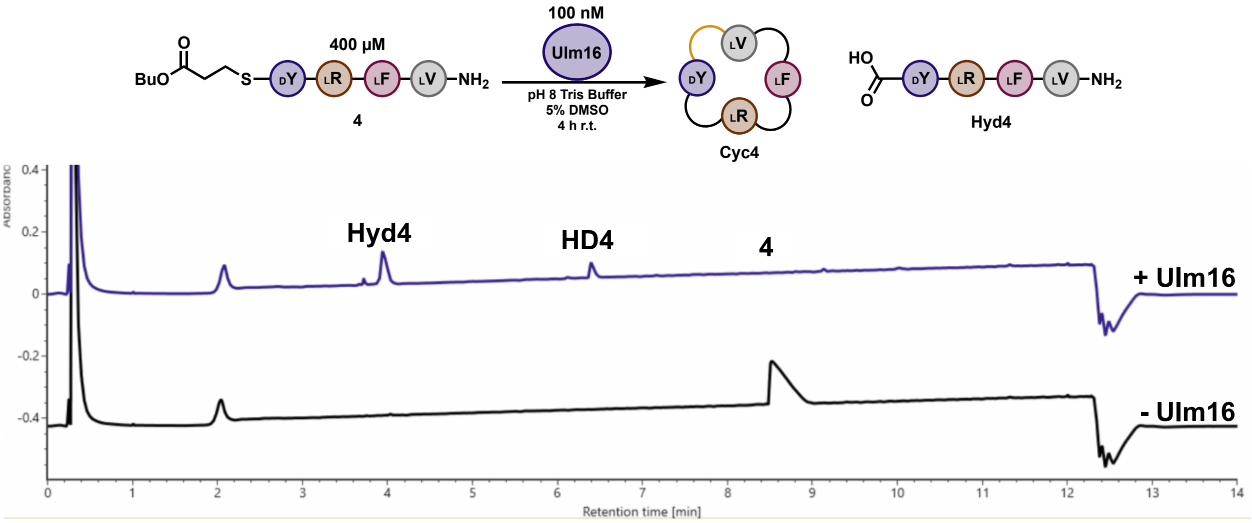

**Figure S14. UPLC UV trace (214 nm) is shown for the reaction of Ulm16 with tetrapeptide substrate 4.** Enzyme and substrate concentrations are indicated above the trace. The substrate is labeled with its corresponding number from the paper where 'Cyc', 'Hyd', and 'HD' represent head-to-tail cyclic peptide, hydrolyzed peptide, and hydrolyzed dimer, respectively. The shown UPLC trace is representative of three replicates.

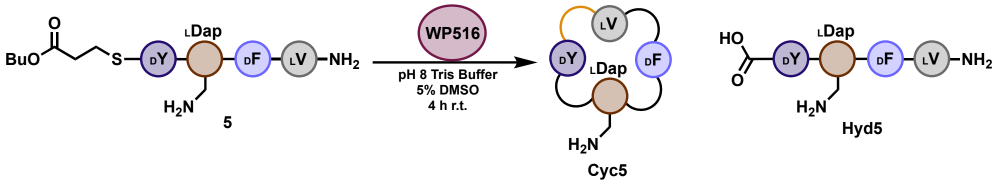

| Enzyme | Peptide | Area Peptide Thioester | Area Cyclic Peptide | Area Hydrolyzed | Sum | TTN | Product/Starting Material | Product/Hydrolysis |
|---|---|---|---|---|---|---|---|---|
| 125 nM | 400 µM | 15344 | 235613 | 8934 | 259891 | 2901 | 15.4 | 26.4 |

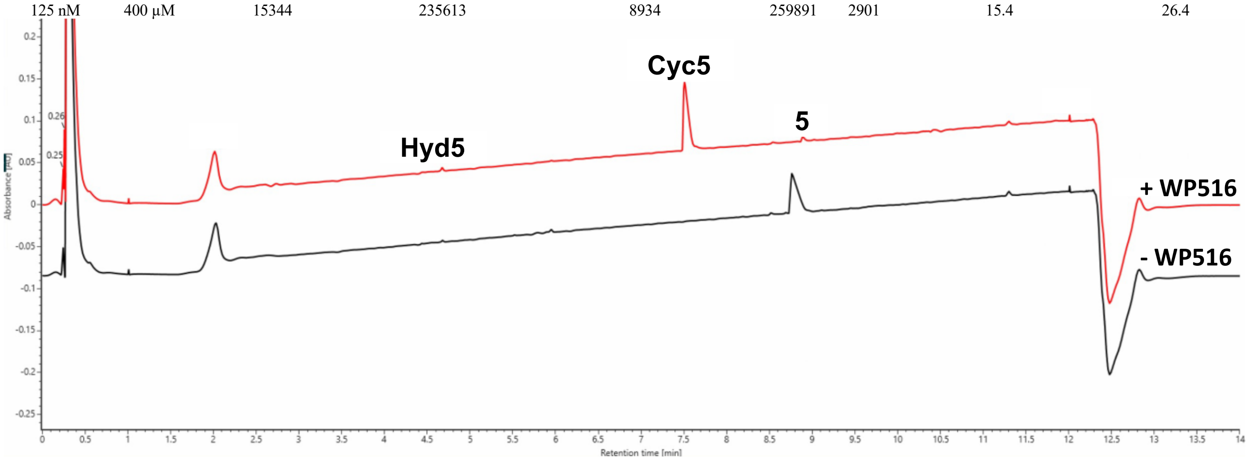

**Figure S15. UPLC UV trace (214 nm) is shown for the total turnover number (TTN) assay of WP516 with tetrapeptide substrate 5.** Enzyme and substrate concentrations are indicated above the trace. The substrate is labeled with its corresponding number from the paper where 'Cyc' and 'Hyd' represent head-to-tail cyclic and hydrolyzed peptide, respectively. The shown UPLC trace is representative of three replicates while the TTN, product/starting material, and product/hydrolysis are averages of three replicates.

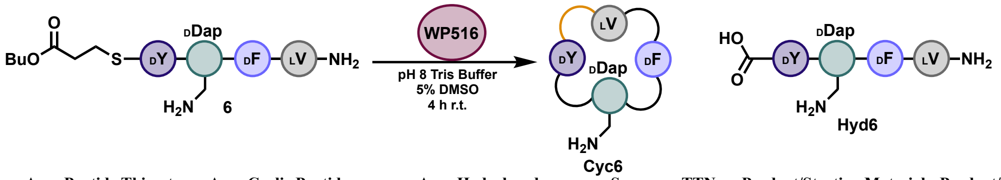

| Enzyme | Peptide | Area Peptide Thioester | Area Cyclic Peptide | Area Hydrolyzed | Sum | TTN | Product/Starting Material | Product/Hydrolysis |
|---|---|---|---|---|---|---|---|---|
| 125 nM | 400 µM | 139509 | 385275 | 13224 | 538009 | 2292 | 2.8 | 29.1 |

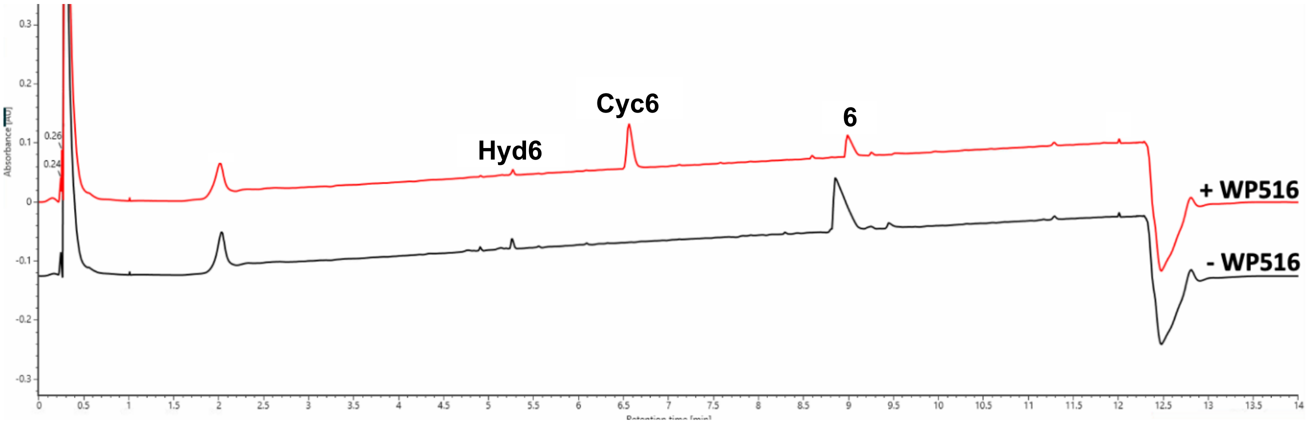

**Figure S16. UPLC UV trace (214 nm) is shown for the total turnover number (TTN) assay of WP516 with tetrapeptide substrate 6.** Enzyme and substrate concentrations are indicated above the trace. The substrate is labeled with its corresponding number from the paper where 'Cyc' and 'Hyd' represent head-to-tail cyclic and hydrolyzed peptide, respectively. The shown UPLC trace is representative of three replicates while the TTN, product/starting material, and product/hydrolysis are averages of three replicates. Note: In the +WP516 sample, there is less hydrolysis than in the no enzyme control (area hydrolyzed in no enzyme control: 21459).

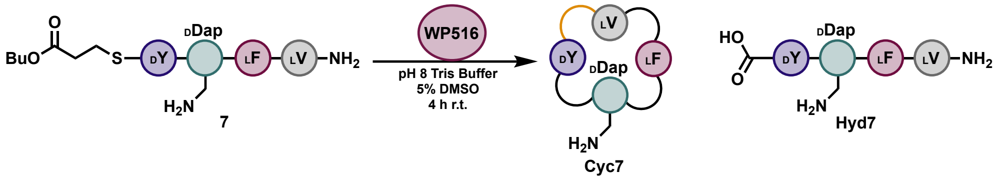

| Enzyme | Peptide | Area Peptide Thioester | Area Cyclic Peptide | Area Hydrolyzed | Sum | TTN | Product/Starting Material | Product/Hydrolysis |
|---|---|---|---|---|---|---|---|---|
| 125 nM | 400 µM | 580312 | 279687 | 8221 | 868220 | 1031 | 0.5 | 34.0 |

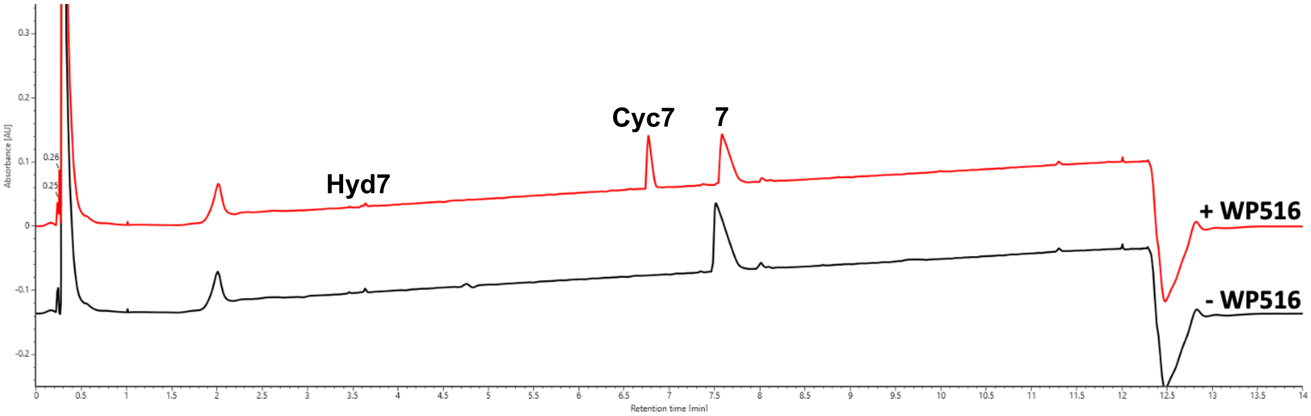

**Figure S17. UPLC UV trace (214 nm) is shown for the total turnover number (TTN) assay of WP516 with tetrapeptide substrate 7.** Enzyme and substrate concentrations are indicated above the trace. The substrate is labeled with its corresponding number from the paper where ‘Cyc’ and ‘Hyd’ represent head-to-tail cyclic and hydrolyzed peptide, respectively. The shown UPLC trace is representative of three replicates while the TTN, product/starting material, and product/hydrolysis are averages of three replicates. Note: In the +WP516 sample, there is less hydrolysis than in the no enzyme control (area hydrolyzed in no enzyme control: 12675).

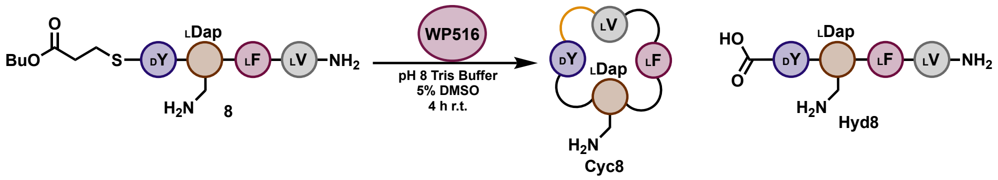

| Enzyme | Peptide | Area Peptide Thioester | Area Cyclic Peptide | Area Hydrolyzed | Sum | TTN | Product/Starting Material | Product/Hydrolysis |
|---|---|---|---|---|---|---|---|---|
| 177 nM | 600 uM | 1052977 | 358351 | 390590 | 1801918 | 674 | 0.3 | 0.9 |

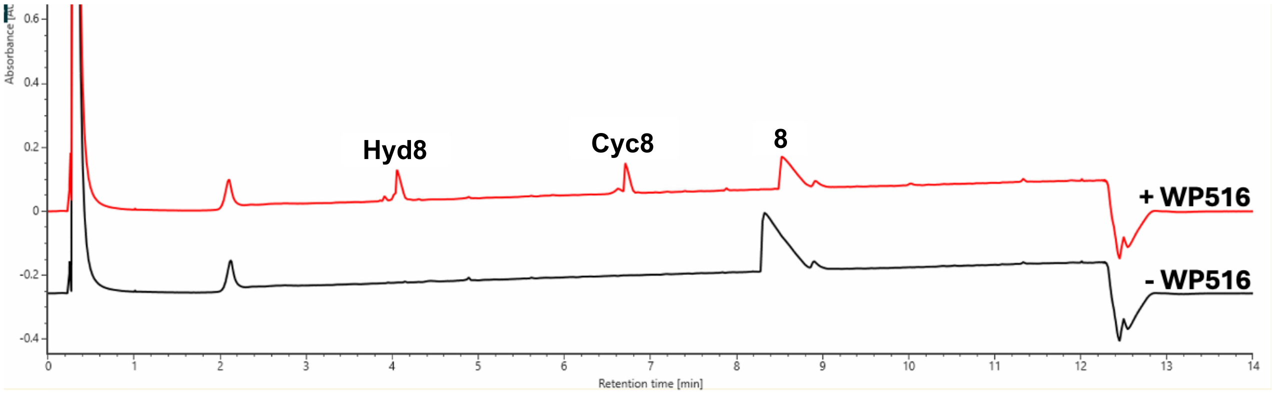

**Figure S18. UPLC UV trace (214 nm) is shown for the total turnover number (TTN) assay of WP516 with tetrapeptide substrate 8.** Enzyme and substrate concentrations are indicated above the trace. The substrate is labeled with its corresponding number from the paper where 'Cyc' and 'Hyd' represent head-to-tail cyclic and hydrolyzed peptide, respectively. The shown UPLC trace is representative of three replicates while the TTN, product/starting material, and product/hydrolysis are averages of three replicates.

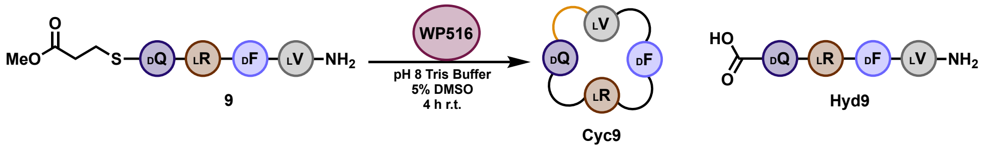

| Enzyme | Peptide | Area Peptide Thioester | Area Cyclic Peptide | Area Hydrolyzed | Sum | TTN | Product/Starting Material | Product/Hydrolysis |
|---|---|---|---|---|---|---|---|---|
| 221 nM | 400 µM | 246642 | 473951 | 28393 | 748986 | 1145 | 1.9 | 16.7 |

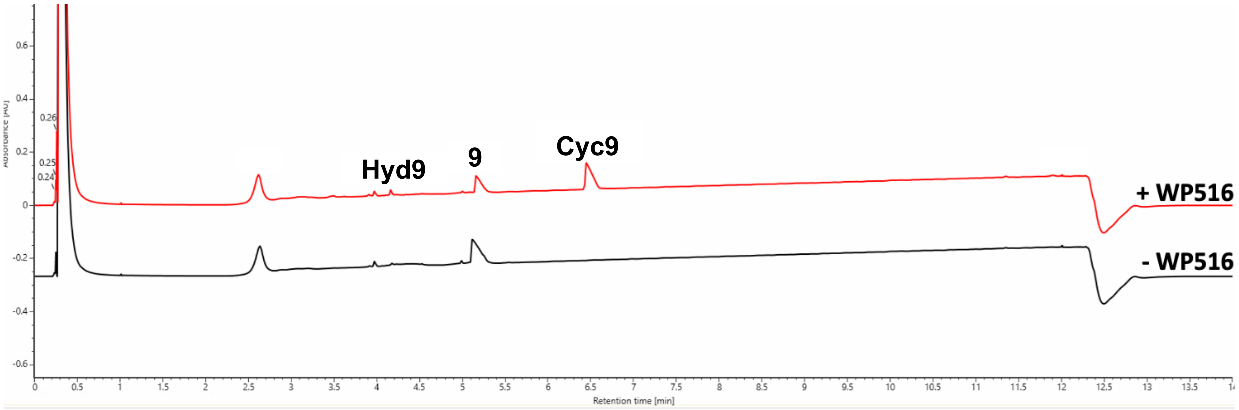

**Figure S19. UPLC UV trace (214 nm) is shown for the total turnover number (TTN) assay of WP516 with tetrapeptide substrate 9.** Enzyme and substrate concentrations are indicated above the trace. The substrate is labeled with its corresponding number from the paper where 'Cyc' and 'Hyd' represent head-to-tail cyclic and hydrolyzed peptide, respectively. The shown UPLC trace is representative of three replicates while the TTN, product/starting material, and product/hydrolysis are averages of three replicates.

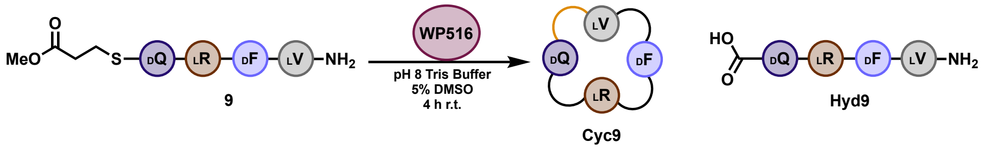

| Enzyme | Peptide | Area Peptide Thioester | Area Cyclic Peptide | Area Hydrolyzed | Sum | TTN | Product/Starting Material | Product/Hydrolysis |
|---|---|---|---|---|---|---|---|---|
| 221 nM | 400 µM | 246642 | 473951 | 28393 | 748986 | 1145 | 1.9 | 16.7 |

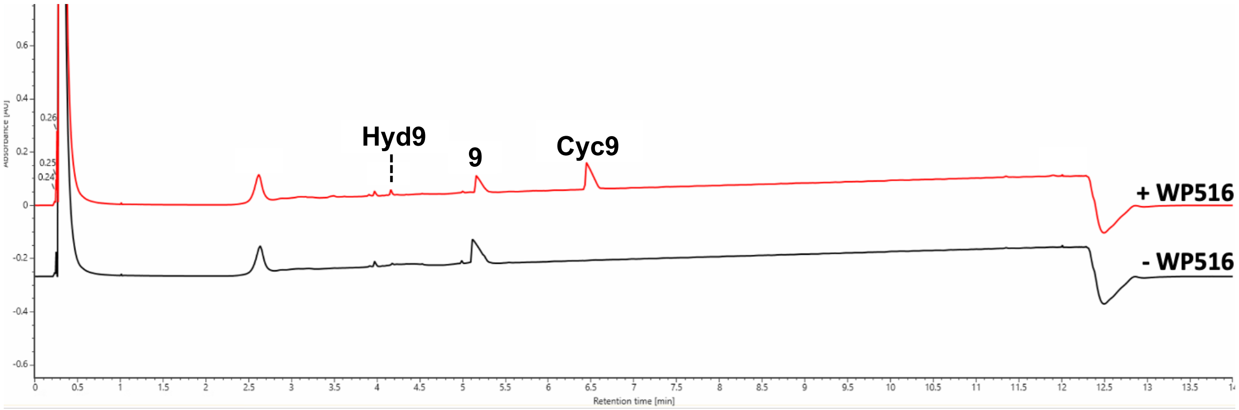

**Figure S20. UPLC UV trace (214 nm) is shown for the total turnover number (TTN) assay of WP516 with tetrapeptide substrate 10.** Enzyme and substrate concentrations are indicated above the trace. The substrate is labeled with its corresponding number from the paper where 'Cyc' and 'Hyd' represent head-to-tail cyclic and hydrolyzed peptide, respectively. The shown UPLC trace is representative of three replicates while the TTN, product/starting material, and product/hydrolysis are averages of three replicates.

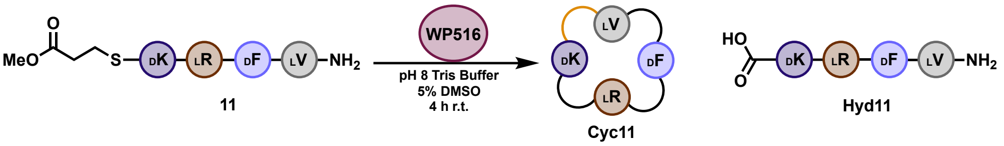

| Enzyme | Peptide | Area Peptide Thioester | Area Cyclic Peptide | Area Hydrolyzed | Sum | TTN | Product/Starting Material | Product/Hydrolysis |
|---|---|---|---|---|---|---|---|---|
| 221 nM | 400 µM | 429411 | 75676 | 22890 | 527978 | 259 | 0.2 | 3.3 |

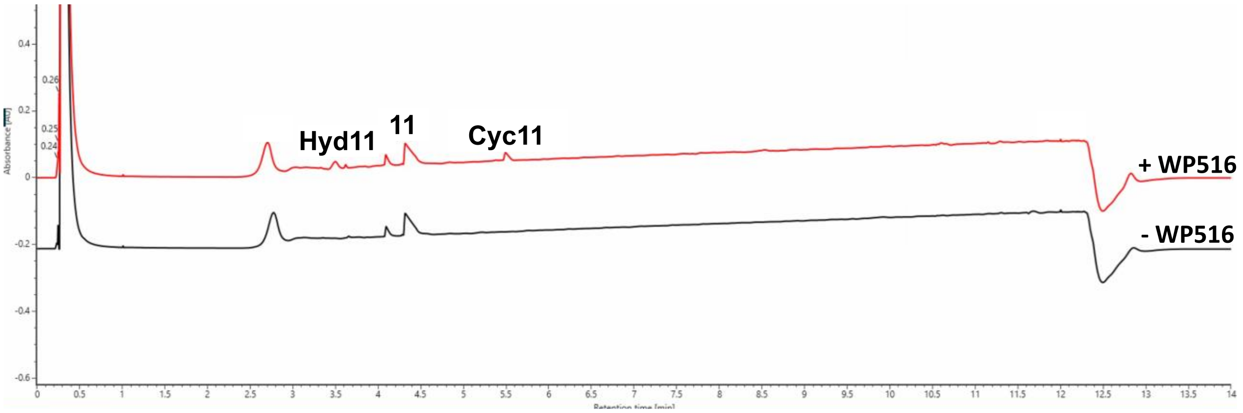

**Figure S21. UPLC UV trace (214 nm) is shown for the total turnover number (TTN) assay of WP516 with tetrapeptide substrate 11.** Enzyme and substrate concentrations are indicated above the trace. The substrate is labeled with its corresponding number from the paper where 'Cyc' and 'Hyd' represent head-to-tail cyclic and hydrolyzed peptide, respectively. The shown UPLC trace is representative of three replicates while the TTN, product/starting material, and product/hydrolysis are averages of three replicates.

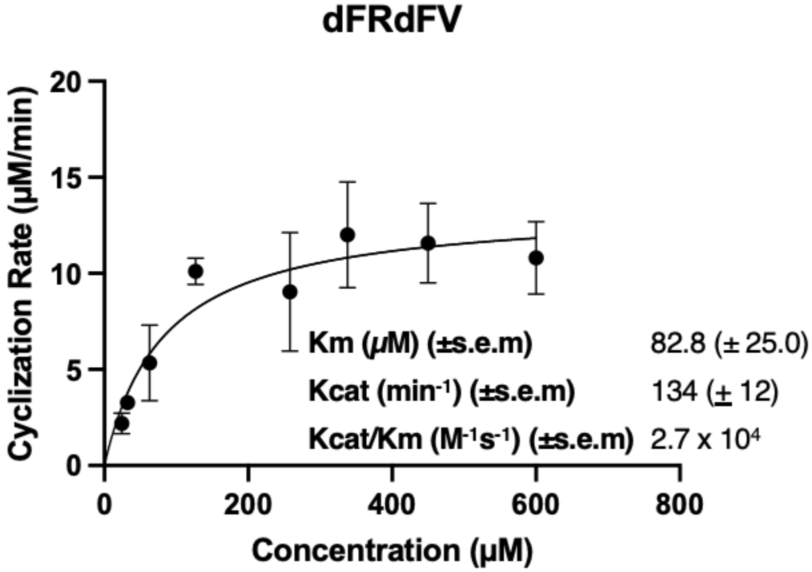

**Figure S22. Michaelis-Menten plots of WP516 incubated with tetrapeptide 12.** The enzyme concentration employed are provided in the methods section. The plots represent the mean of triplicate experiments, and the error bars indicate the standard error of the mean (S.E.M).

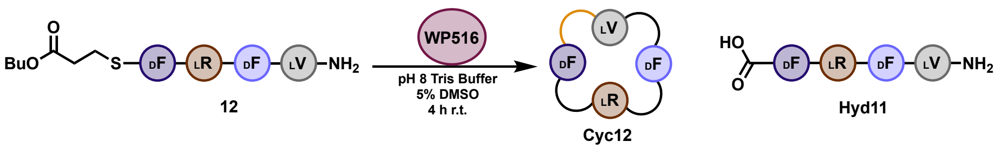

| Enzyme | Peptide | Area Peptide Thioester | Area Cyclic Peptide | Area Hydrolyzed | Sum | TTN | Product/Starting Material | Product/Hydrolysis |
|---|---|---|---|---|---|---|---|---|
| 53 nM | 400 µM | 36661 | 1175553 | 22860 | 1235074 | 7183 | 32.1 | 51.4 |

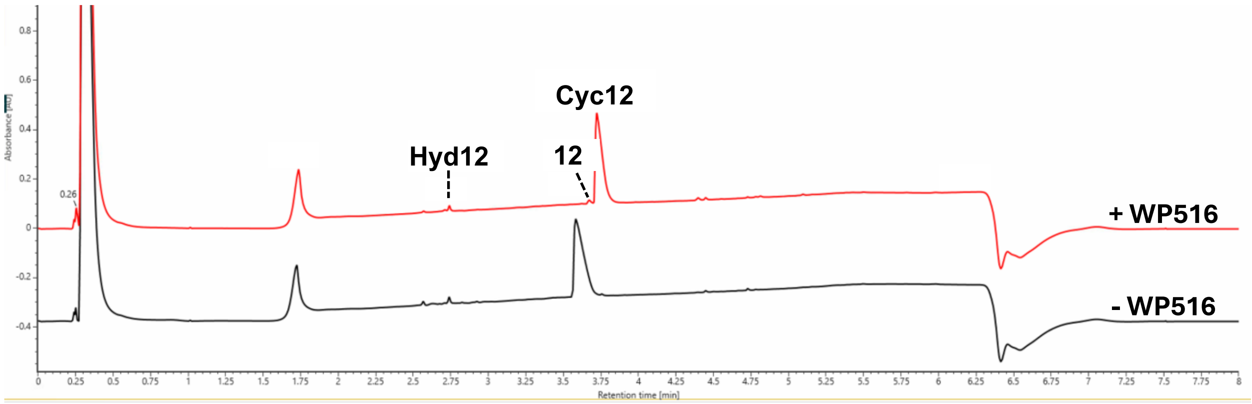

**Figure S23. UPLC UV trace (214 nm) is shown for the total turnover number (TTN) assay of WP516 with tetrapeptide substrate 12.** Enzyme and substrate concentrations are indicated above the trace. The substrate is labeled with its corresponding number from the paper where 'Cyc' and 'Hyd' represent head-to-tail cyclic and hydrolyzed peptide, respectively. The shown UPLC trace is representative of three replicates while the TTN, product/starting material, and product/hydrolysis are averages of three replicates. The reaction was analyzed using a gradient of 0-60% (acetonitrile + 0.1% formic acid) over 4 min instead of the typical 0-40%. Note: In the +WP516 sample, there is less hydrolysis than in the no enzyme control (area hydrolyzed in no enzyme control: 26571).

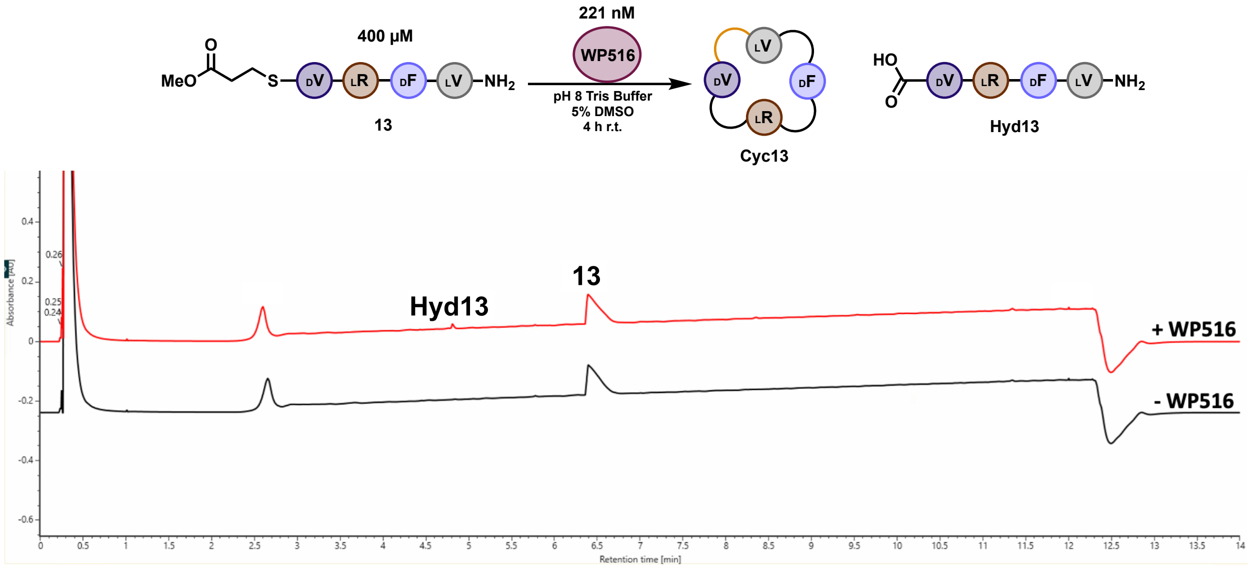

**Figure S24. UPLC UV trace (214 nm) is shown for the total turnover number (TTN) assay of WP516 with tetrapeptide substrate 13.** Enzyme and substrate concentrations are indicated above the trace. The substrate is labeled with its corresponding number from the paper where ‘Cyc’ and ‘Hyd’ represent head-to-tail cyclic and hydrolyzed peptide, respectively. The shown UPLC trace is representative of three replicates while the TTN, product/starting material, and product/hydrolysis are averages of three replicates.

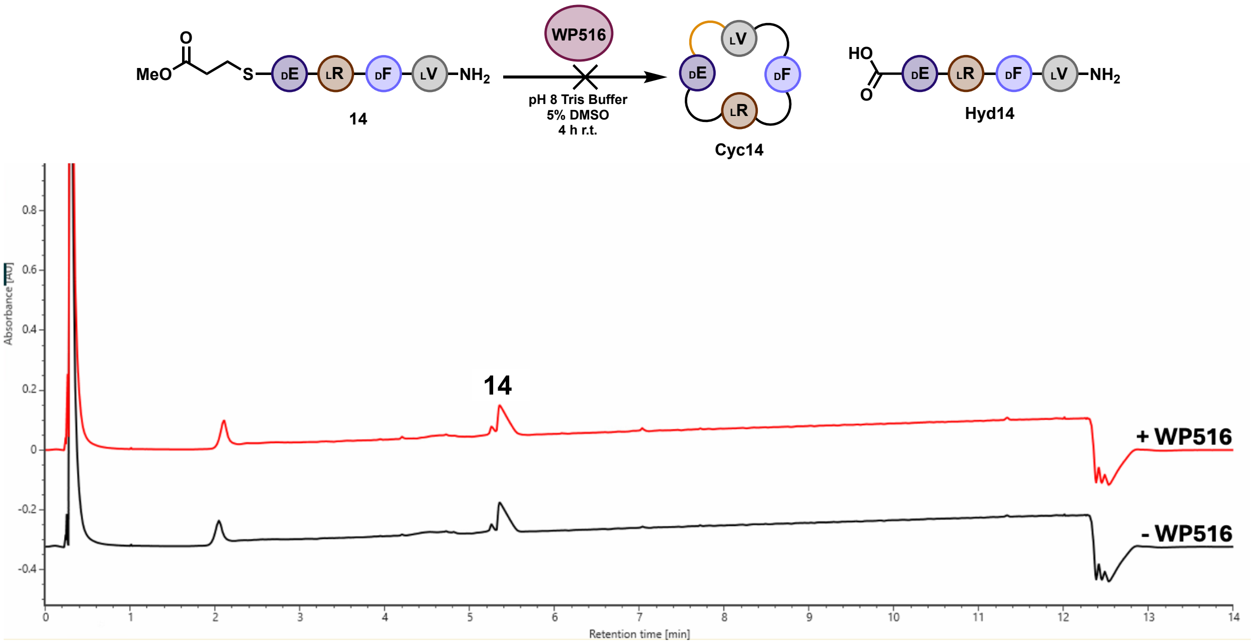

**Figure S25. UPLC UV trace (214 nm) is shown for the total turnover number (TTN) assay of WP516 with tetrapeptide substrate 14.** Enzyme and substrate concentrations are indicated above the trace. The substrate is labeled with its corresponding number from the paper where 'Cyc' and 'Hyd' represent head-to-tail cyclic and hydrolyzed peptide, respectively. The shown UPLC trace is representative of three replicates while the TTN, product/starting material, and product/hydrolysis are averages of three replicates.

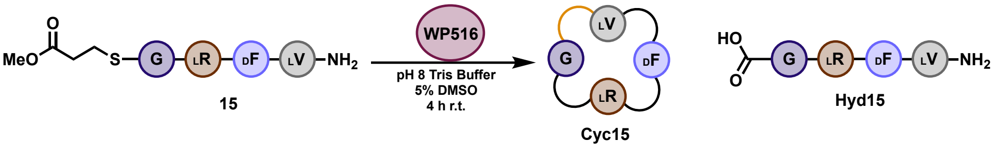

| Enzyme | Peptide | Area Peptide Thioester | Area Cyclic Peptide | Area Hydrolyzed | Sum | TTN | Product/Starting Material | Product/Hydrolysis |
|---|---|---|---|---|---|---|---|---|
| 250 nM | 400 uM | 418326 | 381628 | 54850 | 854804 | 714 | 0.9 | 7.0 |

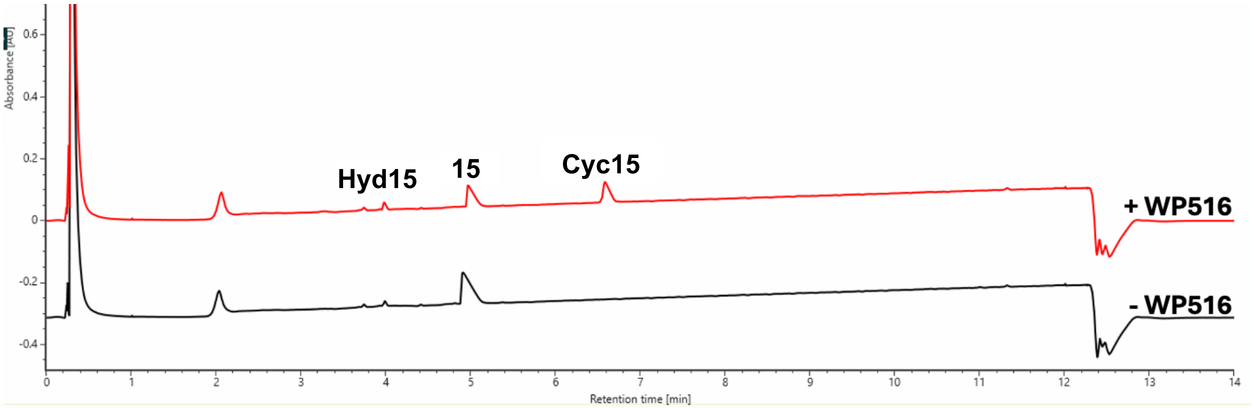

**Figure S26. UPLC UV trace (214 nm) is shown for the total turnover number (TTN) assay of WP516 with tetrapeptide substrate 15.** Enzyme and substrate concentrations are indicated above the trace. The substrate is labeled with its corresponding number from the paper where 'Cyc' and 'Hyd' represent head-to-tail cyclic and hydrolyzed peptide, respectively. The shown UPLC trace is representative of three replicates while the TTN, product/starting material, and product/hydrolysis are averages of three replicates.

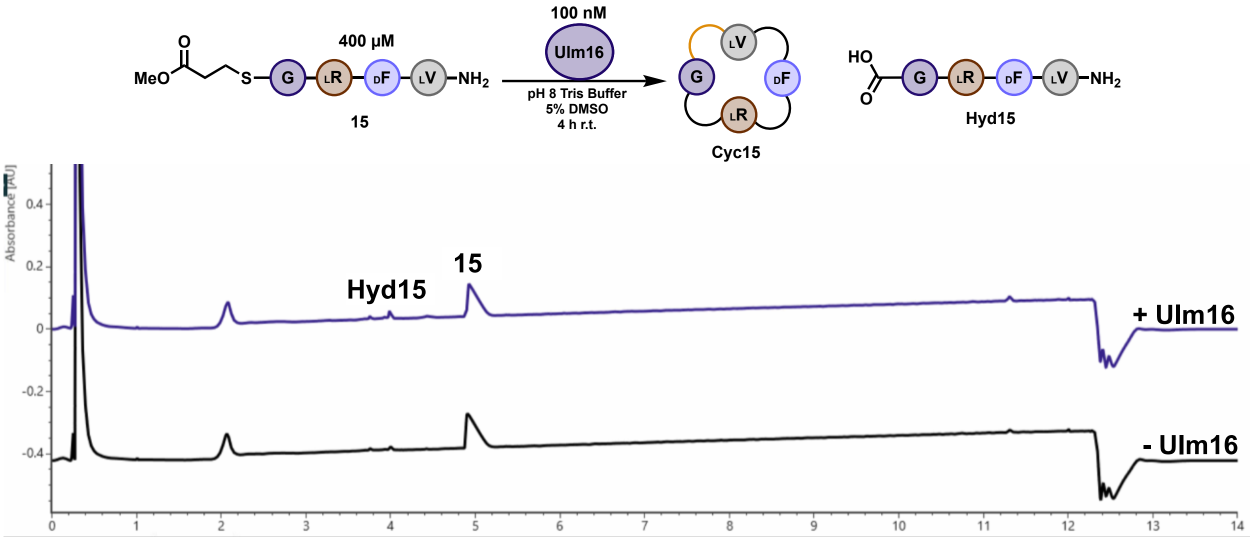

**Figure S27. UPLC UV trace (214 nm) is shown for the reaction of Ulm16 with tetrapeptide substrate 15.** Enzyme and substrate concentrations are indicated above the trace. The substrate is labeled with its corresponding number from the paper where 'Cyc' and 'Hyd' represent head-to-tail cyclic and hydrolyzed peptide, respectively. The shown UPLC trace is representative of three replicates.

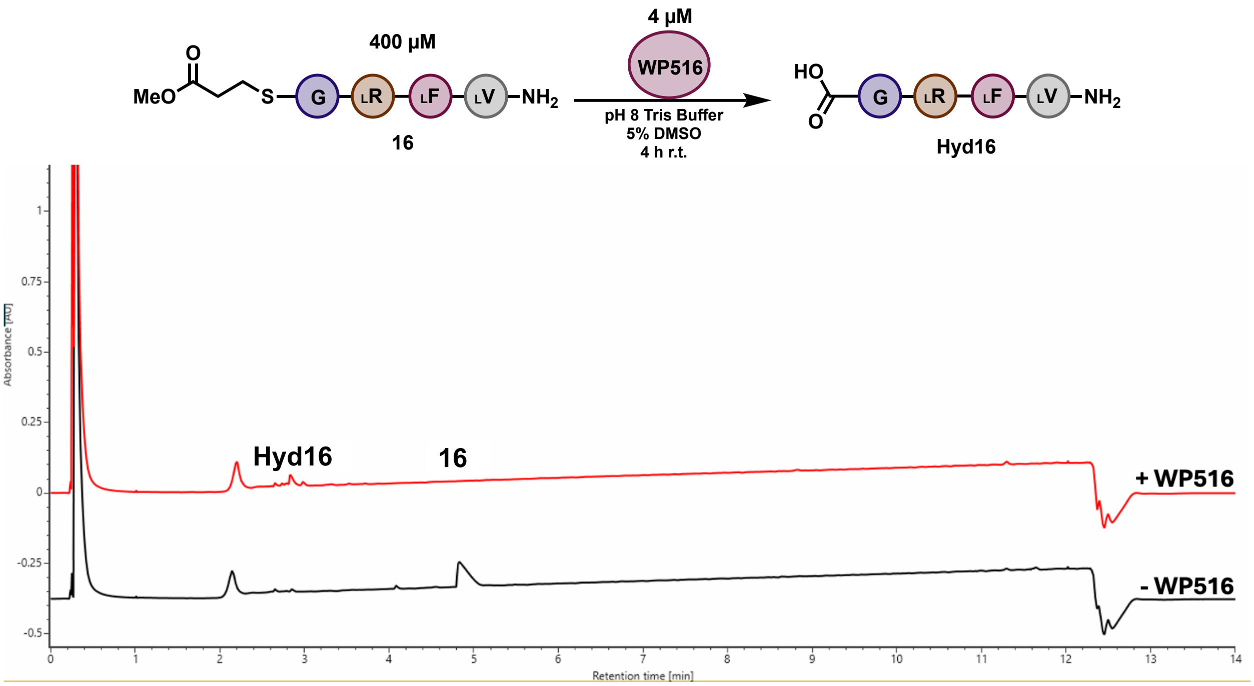

**Figure S28. UPLC UV trace (214 nm) is shown for the total turnover number (TTN) assay of WP516 with tetrapeptide substrate 16.** Enzyme and substrate concentrations are indicated above the trace. The substrate is labeled with its corresponding number from the paper where 'Cyc' and 'Hyd' represent head-to-tail cyclic and hydrolyzed peptide, respectively. The shown UPLC trace is representative of three replicates while the TTN, product/starting material, and product/hydrolysis are averages of three replicates.

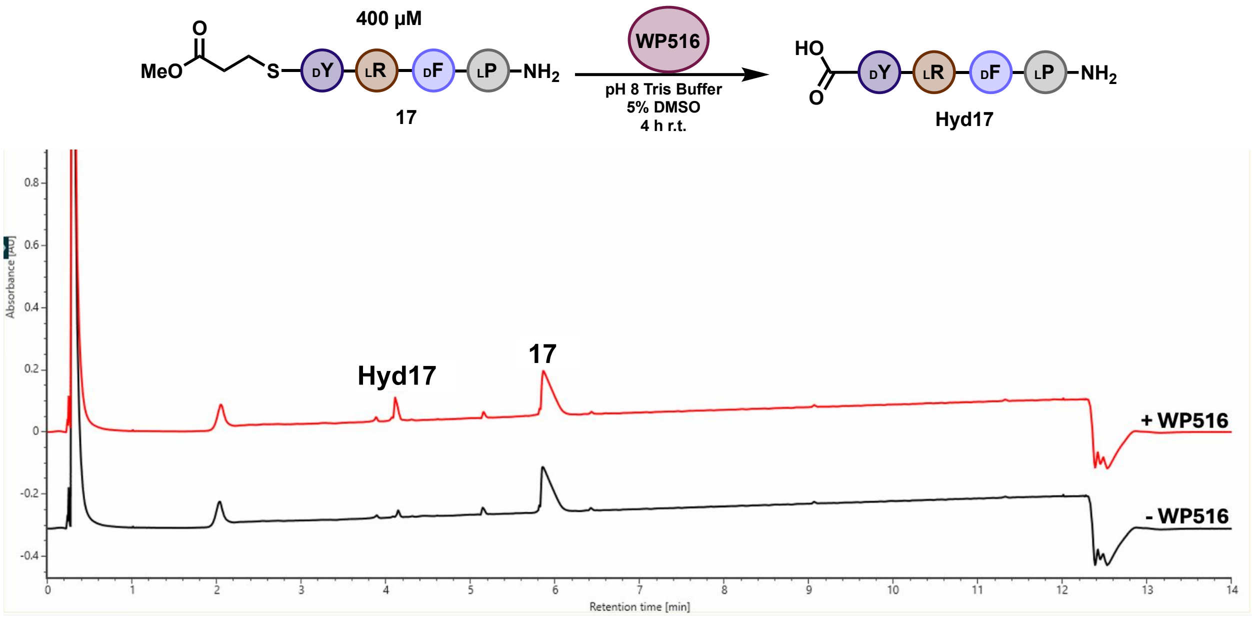

**Figure S29. UPLC UV trace (214 nm) is shown for the total turnover number (TTN) assay of WP516 with tetrapeptide substrate 17.** Enzyme and substrate concentrations are indicated above the trace. The substrate is labeled with its corresponding number from the paper where 'Cyc' and 'Hyd' represent head-to-tail cyclic and hydrolyzed peptide, respectively. The shown UPLC trace is representative of three replicates while the TTN, product/starting material, and product/hydrolysis are averages of three replicates.

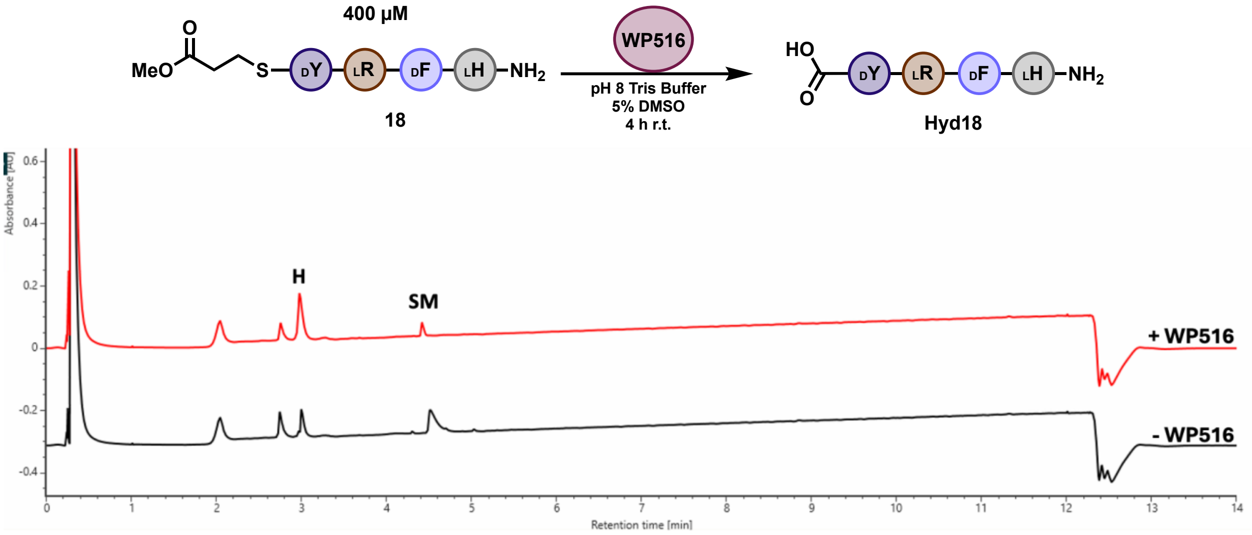

**Figure S30. UPLC UV trace (214 nm) is shown for the total turnover number (TTN) assay of WP516 with tetrapeptide substrate 18.** Enzyme and substrate concentrations are indicated above the trace. The substrate is labeled with its corresponding number from the paper where 'Cyc' and 'Hyd' represent head-to-tail cyclic and hydrolyzed peptide, respectively. The shown UPLC trace is representative of three replicates while the TTN, product/starting material, and product/hydrolysis are averages of three replicates.

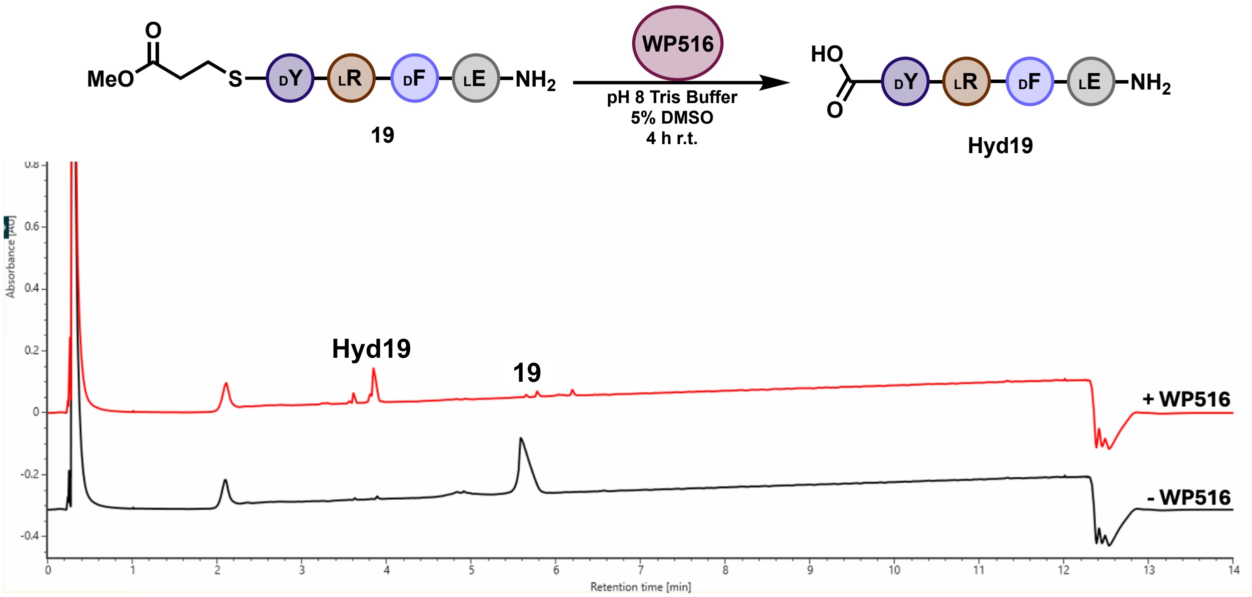

**Figure S31. UPLC UV trace (214 nm) is shown for the total turnover number (TTN) assay of WP516 with tetrapeptide substrate 19.** Enzyme and substrate concentrations are indicated above the trace. The substrate is labeled with its corresponding number from the paper where 'Cyc' and 'Hyd' represent head-to-tail cyclic and hydrolyzed peptide, respectively. The shown UPLC trace is representative of three replicates while the TTN, product/starting material, and product/hydrolysis are averages of three replicates.

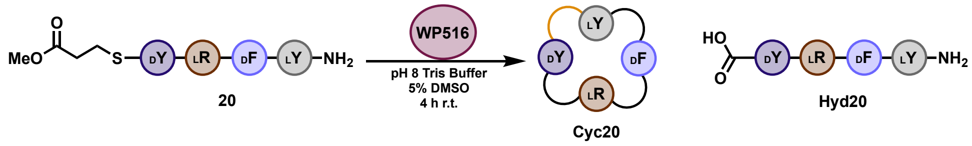

| Enzyme | Peptide | Area Peptide Thioester | Area Cyclic Peptide | Area Hydrolyzed | Sum | TTN | Product/Starting Material | Product/Hydrolysis |
|---|---|---|---|---|---|---|---|---|
| 101 nM | 400 uM | 155865 | 1345446 | 204870 | 1706181 | 3123 | 8.6 | 6.6 |

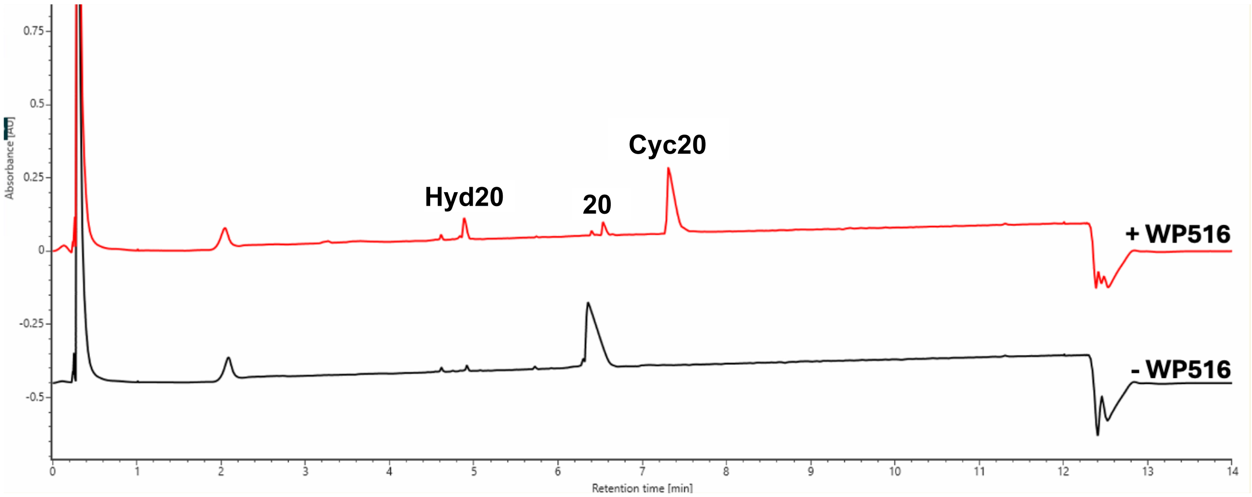

**Figure S32. UPLC UV trace (214 nm) is shown for the total turnover number (TTN) assay of WP516 with tetrapeptide substrate 20.** Enzyme and substrate concentrations are indicated above the trace. The substrate is labeled with its corresponding number from the paper where 'Cyc' and 'Hyd' represent head-to-tail cyclic and hydrolyzed peptide, respectively. The shown UPLC trace is representative of three replicates while the TTN, product/starting material, and product/hydrolysis are averages of three replicates.

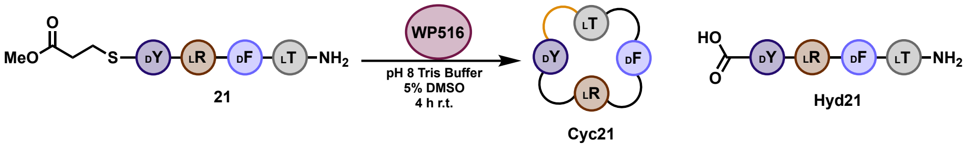

| Enzyme | Peptide | Area Peptide Thioester | Area Cyclic Peptide | Area Hydrolyzed | Sum | TTN | Product/Starting Material | Product/Hydrolysis |
|---|---|---|---|---|---|---|---|---|
| 53 nM | 400 uM | 63195 | 570822 | 23319 | 657336 | 6554 | 9.0 | 24.5 |

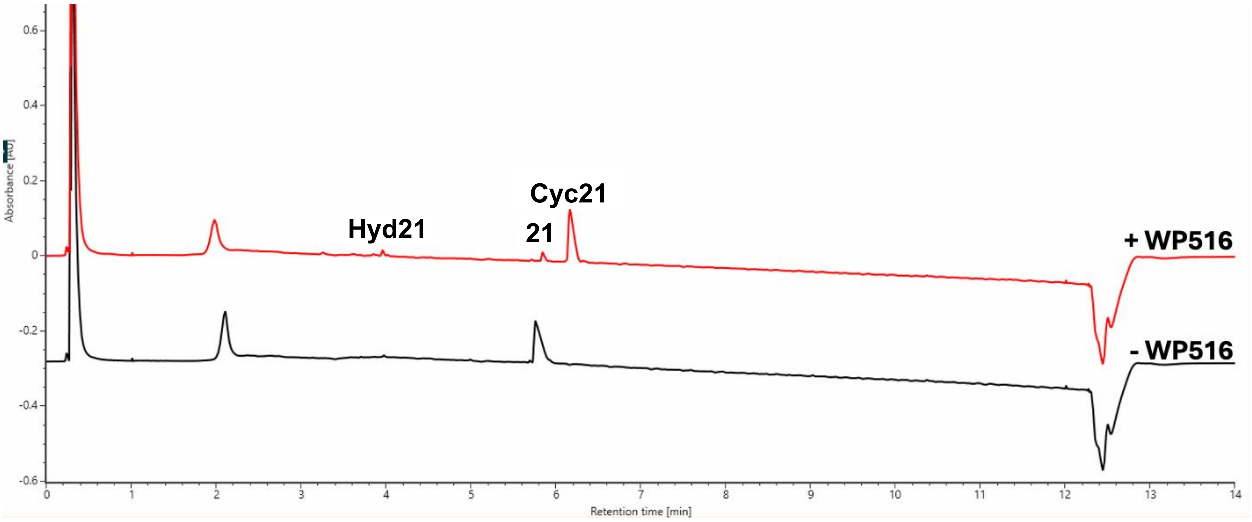

**Figure S33. UPLC UV trace (214 nm) is shown for the total turnover number (TTN) assay of WP516 with tetrapeptide substrate 21.** Enzyme and substrate concentrations are indicated above the trace. The substrate is labeled with its corresponding number from the paper where 'Cyc' and 'Hyd' represent head-to-tail cyclic and hydrolyzed peptide, respectively. The shown UPLC trace is representative of three replicates while the TTN, product/starting material, and product/hydrolysis are averages of three replicates. This reaction was analyzed by loading 2 μL onto a CORTECS T3 Column instead of the typical 5 μL.

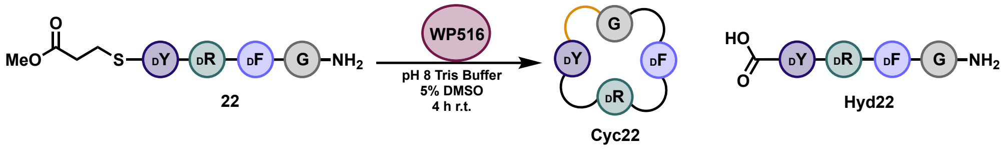

| Enzyme | Peptide | Area Peptide Thioester | Area Cyclic Peptide | Area Hydrolyzed | Sum | TTN | Product/Starting Material | Product/Hydrolysis |
|---|---|---|---|---|---|---|---|---|
| 315 nM | 400 uM | 296800 | 273298 | 336965 | 907063 | 383 | 0.9 | 0.8 |

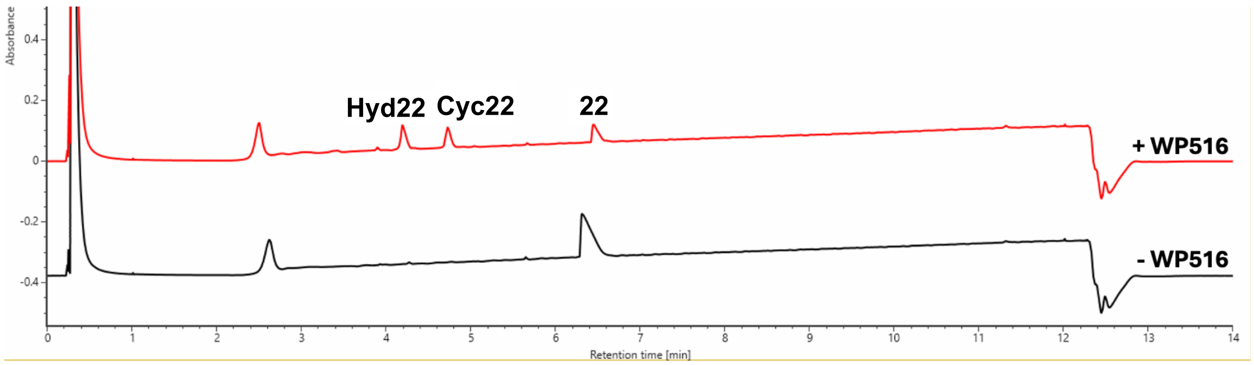

**Figure S34. UPLC UV trace (214 nm) is shown for the total turnover number (TTN) assay of WP516 with tetrapeptide substrate 22.** Enzyme and substrate concentrations are indicated above the trace. The substrate is labeled with its corresponding number from the paper where 'Cyc' and 'Hyd' represent head-to-tail cyclic and hydrolyzed peptide, respectively. The shown UPLC trace is representative of three replicates while the TTN, product/starting material, and product/hydrolysis are averages of three replicates.

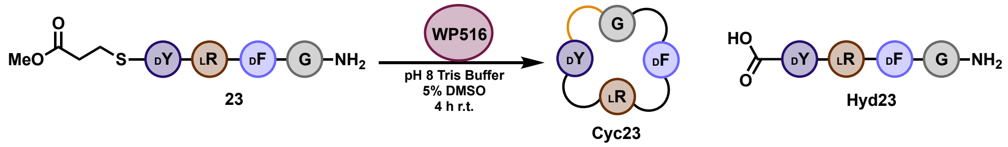

| Enzyme | Peptide | Area Peptide Thioester | Area Cyclic Peptide | Area Hydrolyzed | Sum | TTN | Product/Starting Material | Product/Hydrolysis |
|---|---|---|---|---|---|---|---|---|
| 315 nM | 400 uM | 114963 | 106148 | 286204 | 507315 | 266 | 0.9 | 0.4 |

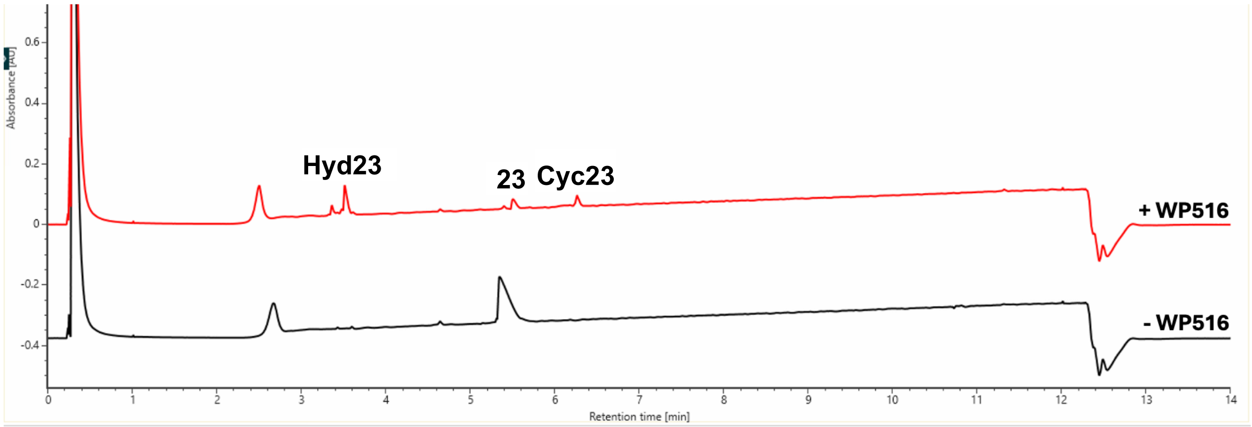

**Figure S35. UPLC UV trace (214 nm) is shown for the total turnover number (TTN) assay of WP516 with tetrapeptide substrate 23.** Enzyme and substrate concentrations are indicated above the trace. The substrate is labeled with its corresponding number from the paper where 'Cyc' and 'Hyd' represent head-to-tail cyclic and hydrolyzed peptide, respectively. The shown UPLC trace is representative of three replicates while the TTN, product/starting material, and product/hydrolysis are averages of three replicates.

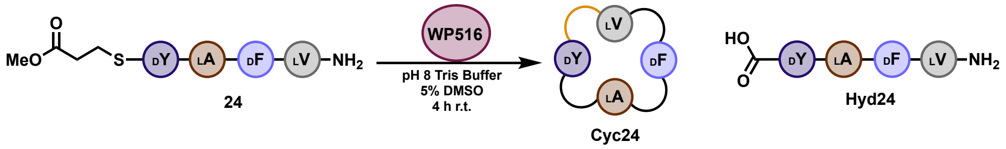

| Enzyme | Peptide | Area Peptide Thioester | Area Cyclic Peptide | Area Hydrolyzed | Sum | TTN | Product/Starting Material | Product/Hydrolysis |
|---|---|---|---|---|---|---|---|---|
| 1.4 µM | 600 µM | 3506220 | 215114 | 133426 | 3854761 | 24 | 0.1 | 1.6 |

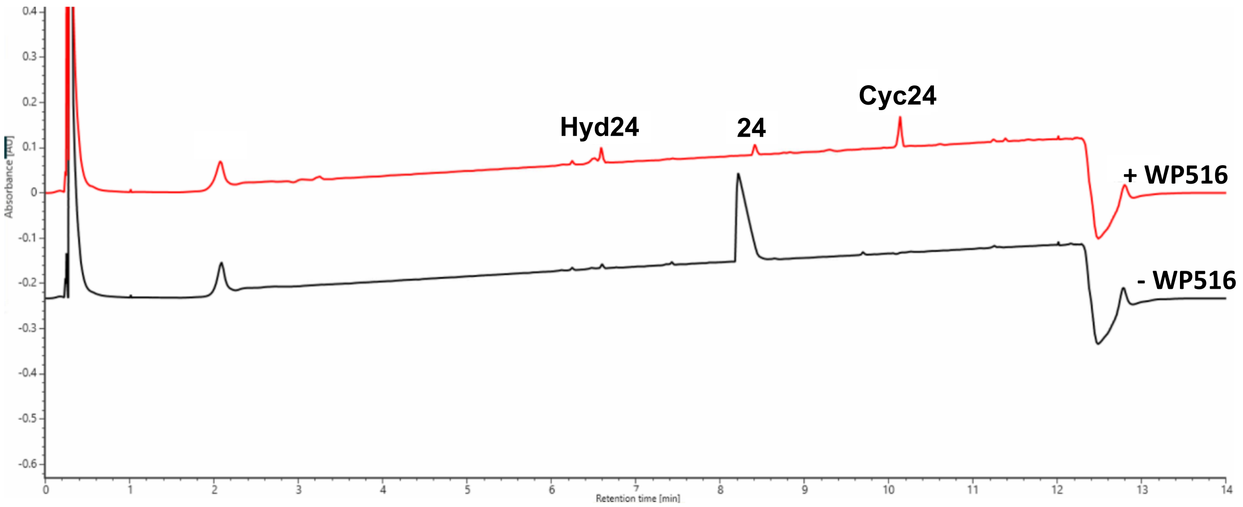

**Figure S36. UPLC UV trace (214 nm) is shown for the total turnover number (TTN) assay of WP516 with tetrapeptide substrate 24.** Enzyme and substrate concentrations are indicated above the trace. The substrate is labeled with its corresponding number from the paper where 'Cyc' and 'Hyd' represent head-to-tail cyclic and hydrolyzed peptide, respectively. The shown UPLC trace is representative of three replicates while the TTN, product/starting material, and product/hydrolysis are averages of three replicates.

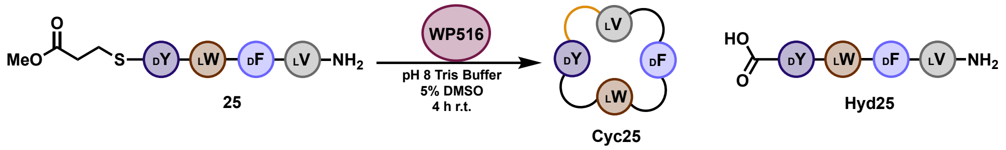

| Enzyme | Peptide | Area Peptide Thioester | Area Cyclic Peptide | Area Hydrolyzed | Sum | TTN | Product/Starting Material | Product/Hydrolysis |
|---|---|---|---|---|---|---|---|---|
| 1.4 µM | 600 µM | 3506220 | 215114 | 133426 | 3854761 | 24 | 0.1 | 1.6 |

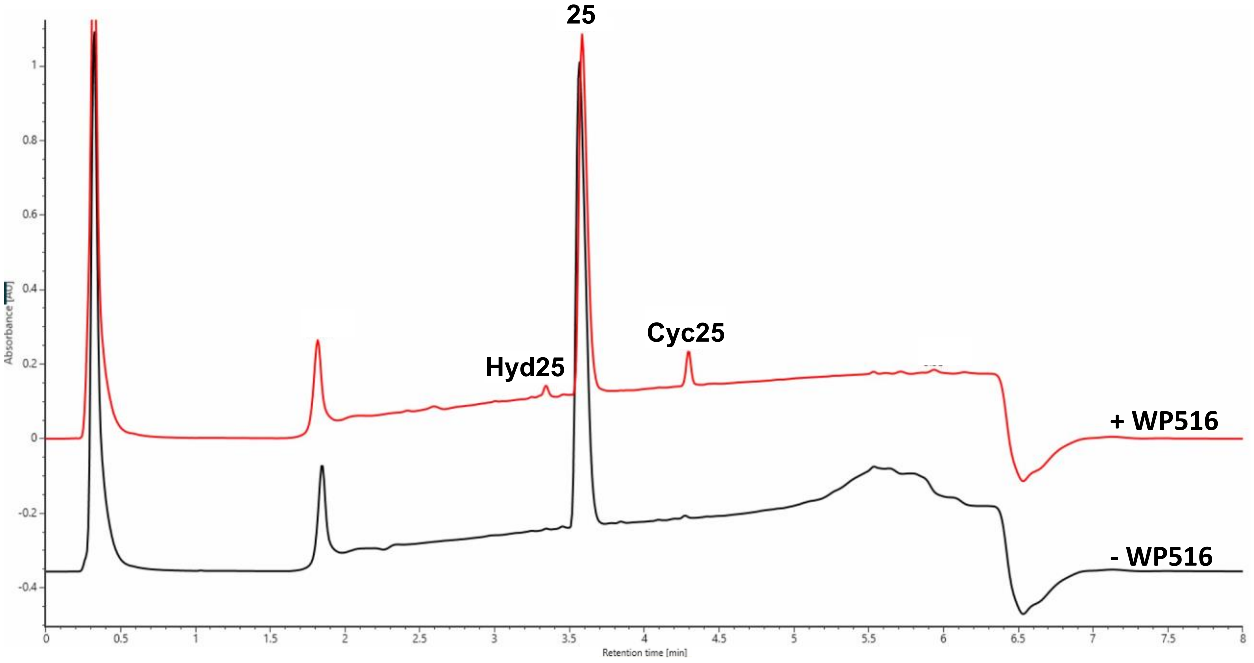

**Figure S37. UPLC UV trace (214 nm) is shown for the total turnover number (TTN) assay of WP516 with tetrapeptide substrate 25.** Enzyme and substrate concentrations are indicated above the trace. The substrate is labeled with its corresponding number from the paper where 'Cyc' and 'Hyd' represent head-to-tail cyclic and hydrolyzed peptide, respectively. The shown UPLC trace is representative of three replicates while the TTN, product/starting material, and product/hydrolysis are averages of three replicates. The reaction was analyzed using a gradient of 0-70% (acetonitrile + 0.1% formic acid) over 4 min instead of the typical 0-40%.

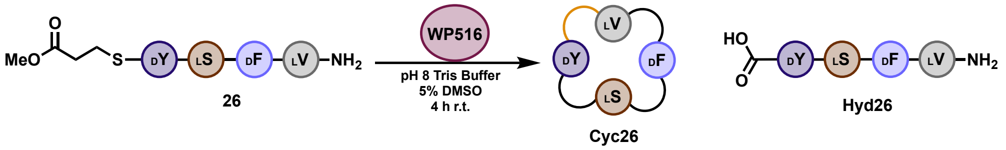

| Enzyme | Peptide | Area Peptide Thioester | Area Cyclic Peptide | Area Hydrolyzed | Sum | TTN | Product/Starting Material | Product/Hydrolysis |
| --- | --- | --- | --- | --- | --- | --- | --- | --- |
| 1.4 µM | 400 µM | 29211 | 232254 | 42396 | 303861 | 218 | 8.0 | 5.5 |

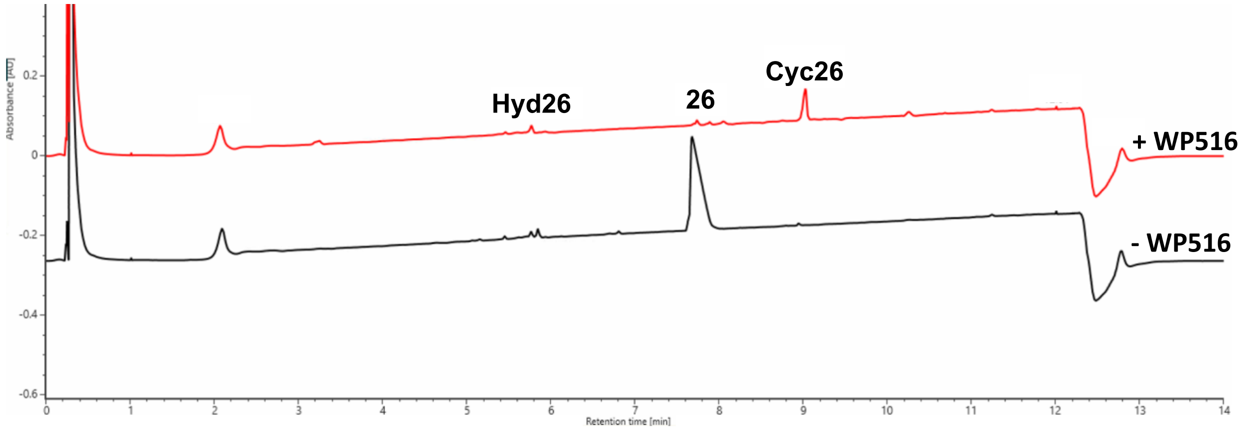

**Figure S38. UPLC UV trace (214 nm) is shown for the total turnover number (TTN) assay of WP516 with tetrapeptide substrate 26.** Enzyme and substrate concentrations are indicated above the trace. The substrate is labeled with its corresponding number from the paper where 'Cyc' and 'Hyd' represent head-to-tail cyclic and hydrolyzed peptide, respectively. The shown UPLC trace is representative of three replicates while the TTN, product/starting material, and product/hydrolysis are averages of three replicates.

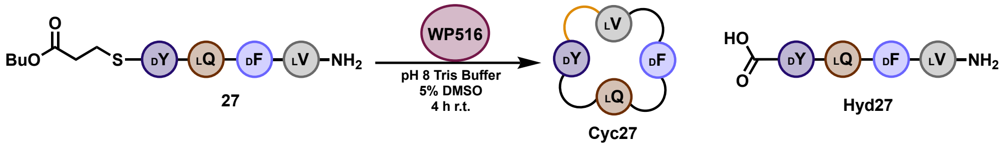

| Enzyme | Peptide | Area Peptide Thioester | Area Cyclic Peptide | Area Hydrolyzed | Sum | TTN | Product/Starting Material | Product/Hydrolysis |
|---|---|---|---|---|---|---|---|---|
| 500 nM | 400 µM | 653366 | 115590 | - | 768956 | 120 | 0.2 | - |

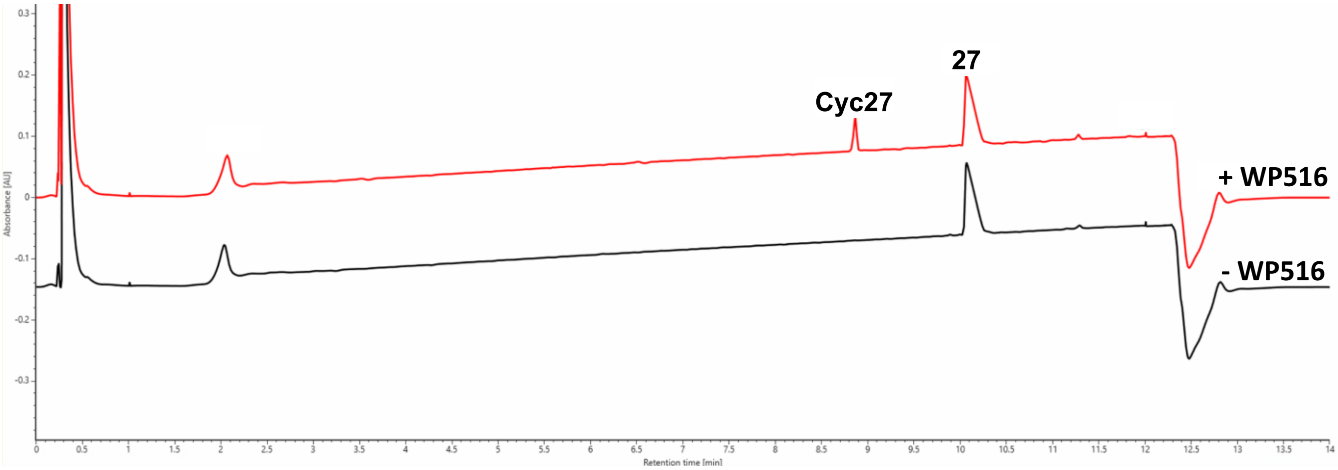

**Figure S39. UPLC UV trace (214 nm) is shown for the total turnover number (TTN) assay of WP516 with tetrapeptide substrate 27.** Enzyme and substrate concentrations are indicated above the trace. The substrate is labeled with its corresponding number from the paper where ‘Cyc’ and ‘Hyd’ represent head-to-tail cyclic and hydrolyzed peptide, respectively. The shown UPLC trace is representative of three replicates while the TTN, product/starting material, and product/hydrolysis are averages of three replicates.

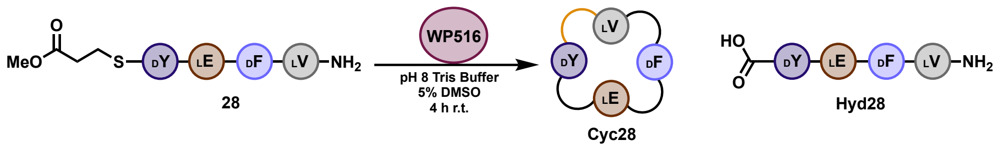

| Enzyme | Peptide | Area Peptide Thioester | Area Cyclic Peptide | Area Hydrolyzed | Sum | TTN | Product/Starting Material | Product/Hydrolysis |
|---|---|---|---|---|---|---|---|---|
| 53 nM | 400 uM | 157390 | 1113732 | 56823 | 1327945 | 6330 | 7.1 | 19.6 |

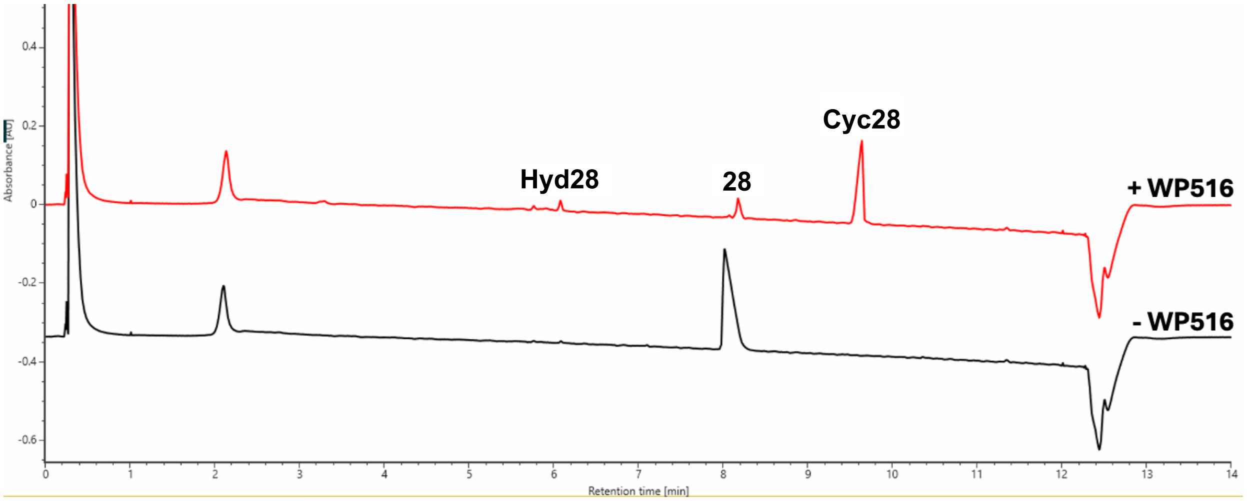

**Figure S40. UPLC UV trace (214 nm) is shown for the total turnover number (TTN) assay of WP516 with tetrapeptide substrate 28.** Enzyme and substrate concentrations are indicated above the trace. The substrate is labeled with its corresponding number from the paper where 'Cyc' and 'Hyd' represent head-to-tail cyclic and hydrolyzed peptide, respectively. The shown UPLC trace is representative of three replicates while the TTN, product/starting material, and product/hydrolysis are averages of three replicates.

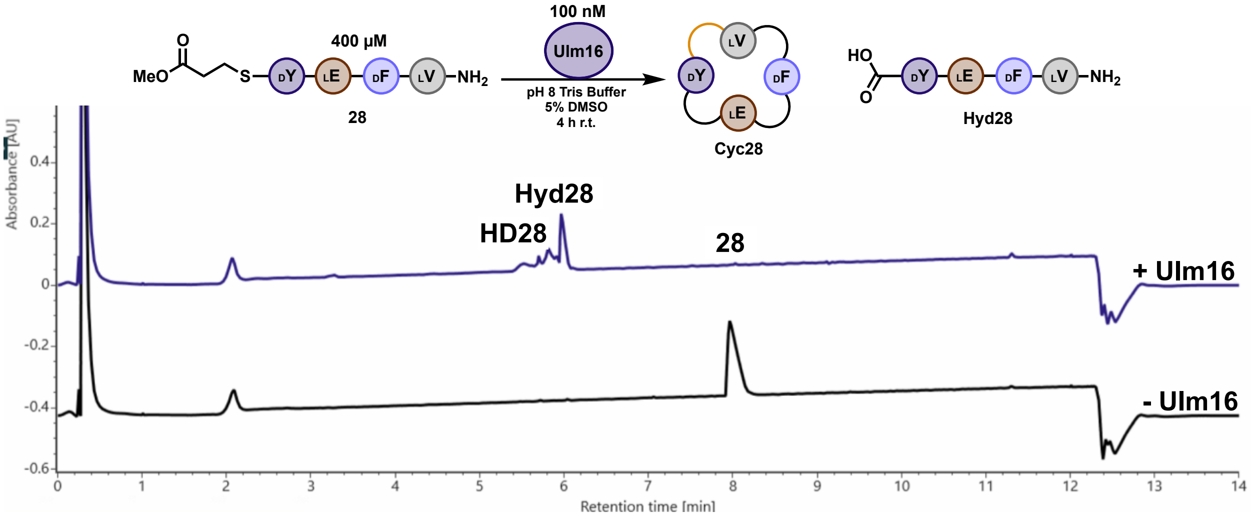

**Figure S41. UPLC UV trace (214 nm) is shown for the reaction of Ulm16 with tetrapeptide substrate 28.** Enzyme and substrate concentrations are indicated above the trace. The substrate is labeled with its corresponding number from the paper where 'Cyc', 'Hyd', and 'HD' represent head-to-tail cyclic, hydrolyzed peptide, and hydrolyzed dimer, respectively. The shown UPLC trace is representative of three replicates.

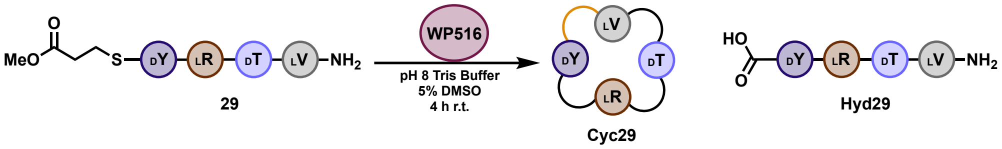

| Enzyme | Peptide | Area Peptide Thioester | Area Cyclic Peptide | Area Hydrolyzed | Sum | TTN | Product/Starting Material | Product/Hydrolysis |
|---|---|---|---|---|---|---|---|---|
| 53 nM | 400 uM | 52162 | 337778 | 11788 | 401728 | 6346 | 6.5 | 28.7 |

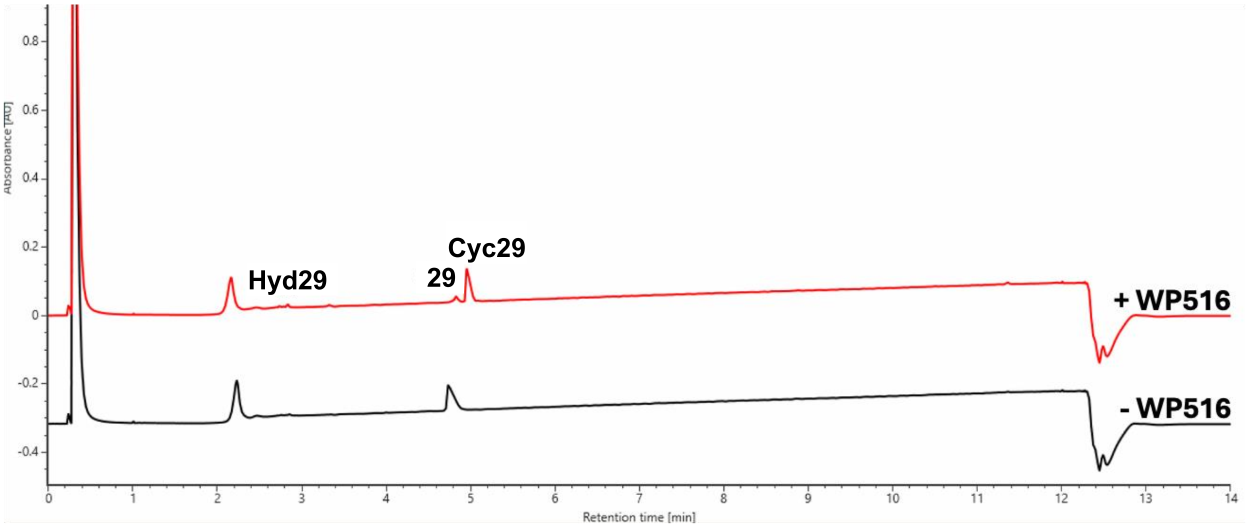

**Figure S42. UPLC UV trace (214 nm) is shown for the total turnover number (TTN) assay of WP516 with tetrapeptide substrate 29.** Enzyme and substrate concentrations are indicated above the trace. The substrate is labeled with its corresponding number from the paper where 'Cyc' and 'Hyd' represent head-to-tail cyclic and hydrolyzed peptide, respectively. The shown UPLC trace is representative of three replicates while the TTN, product/starting material, and product/hydrolysis are averages of three replicates. This reaction was analyzed by loading 2 μL onto a CORTECS T3 Column instead of the typical 5 μL.

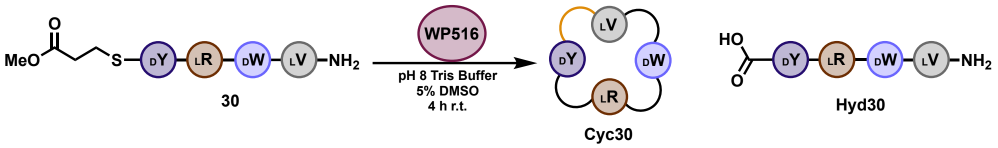

| Enzyme | Peptide | Area Peptide Thioester | Area Cyclic Peptide | Area Hydrolyzed | Sum | TTN | Product/Starting Material | Product/Hydrolysis |
|---|---|---|---|---|---|---|---|---|
| 53 nM | 400 µM | 40304 | 2300622 | 48118 | 2389045 | 7268 | 57.1 | 47.8 |

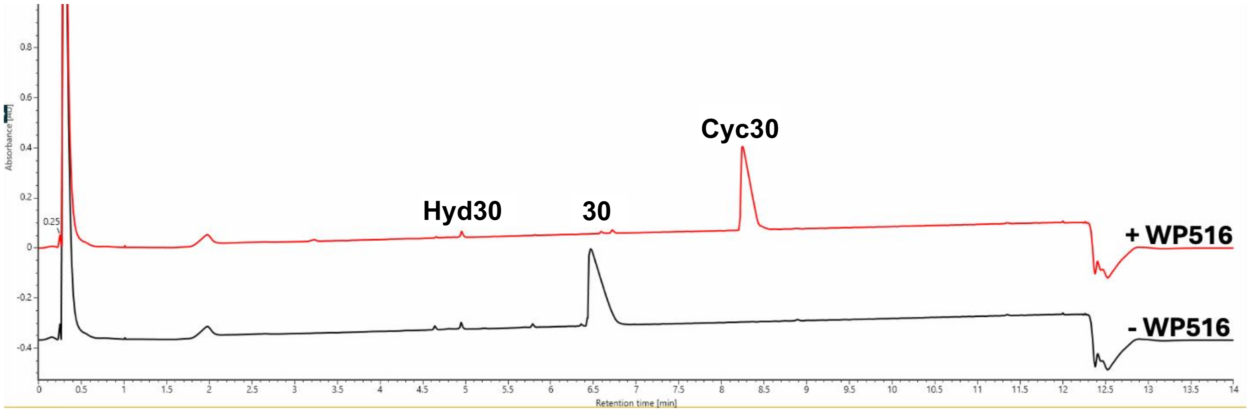

**Figure S43. UPLC UV trace (214 nm) is shown for the total turnover number (TTN) assay of WP516 with tetrapeptide substrate 30.** Enzyme and substrate concentrations are indicated above the trace. The substrate is labeled with its corresponding number from the paper where ‘Cyc’ and ‘Hyd’ represent head-to-tail cyclic and hydrolyzed peptide, respectively. The shown UPLC trace is representative of three replicates while the TTN, product/starting material, and product/hydrolysis are averages of three replicates.

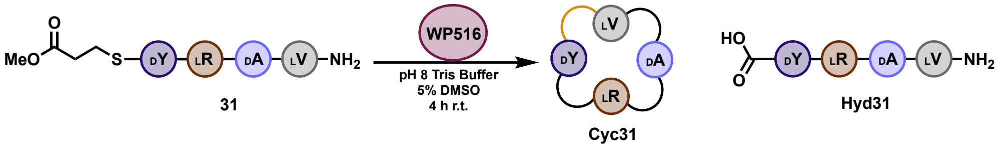

| Enzyme | Peptide | Area Peptide Thioester | Area Cyclic Peptide | Area Hydrolyzed | Sum | TTN | Product/Starting Material | Product/Hydrolysis |
|---|---|---|---|---|---|---|---|---|
| 53 nM | 400 µM | 22407 | 740959 | 4666 | 768032 | 7281 | 33.1 | 158.8 |

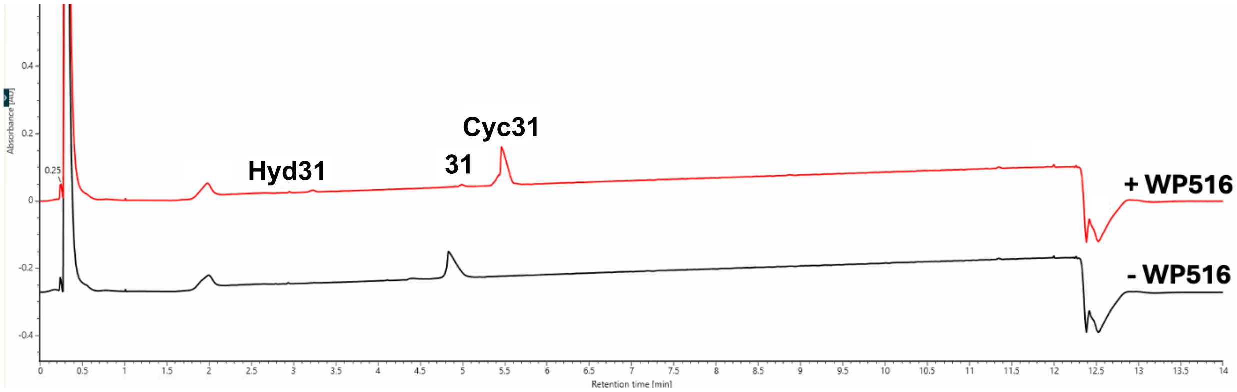

**Figure S44. UPLC UV trace (214 nm) is shown for the total turnover number (TTN) assay of WP516 with tetrapeptide substrate 31.** Enzyme and substrate concentrations are indicated above the trace. The substrate is labeled with its corresponding number from the paper where 'Cyc' and 'Hyd' represent head-to-tail cyclic and hydrolyzed peptide, respectively. The shown UPLC trace is representative of three replicates while the TTN, product/starting material, and product/hydrolysis are averages of three replicates.

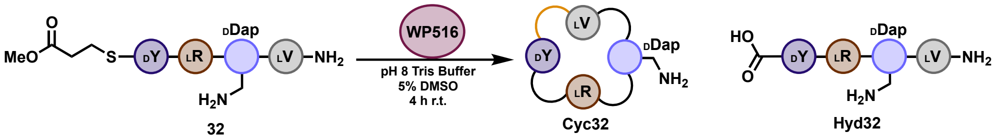

| Enzyme | Peptide | Area Peptide Thioester | Area Cyclic Peptide | Area Hydrolyzed | Sum | TTN | Product/Starting Material | Product/Hydrolysis |
|---|---|---|---|---|---|---|---|---|
| 53 nM | 400 µM | 281057 | 329889 | - | 610946 | 4075 | 1.2 | - |

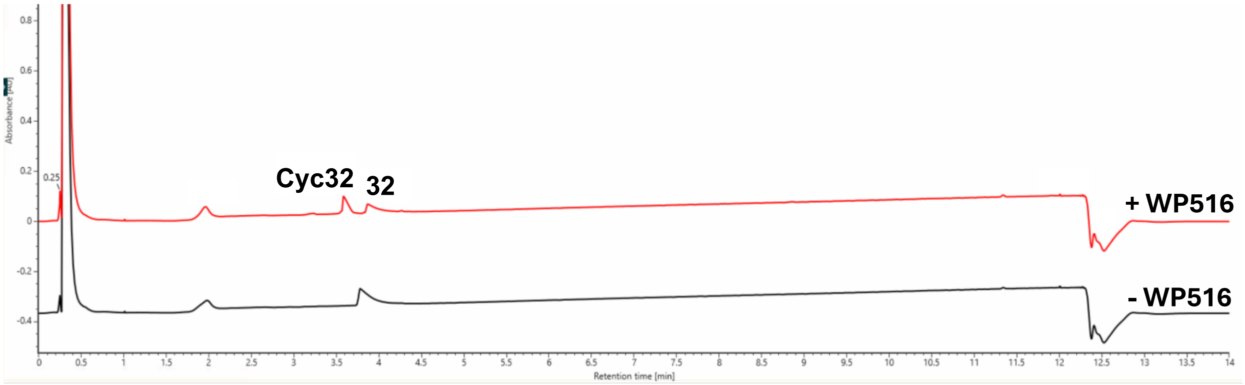

**Figure S45. UPLC UV trace (214 nm) is shown for the total turnover number (TTN) assay of WP516 with tetrapeptide substrate 32.** Enzyme and substrate concentrations are indicated above the trace. The substrate is labeled with its corresponding number from the paper where 'Cyc' and 'Hyd' represent head-to-tail cyclic and hydrolyzed peptide, respectively. The shown UPLC trace is representative of three replicates while the TTN, product/starting material, and product/hydrolysis are averages of three replicates.

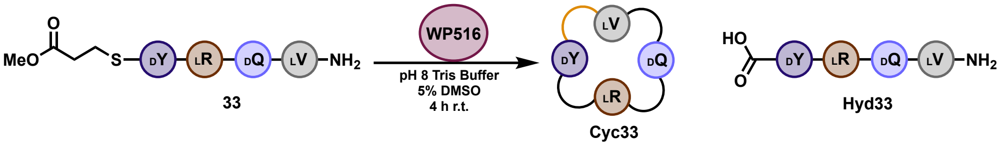

| Enzyme | Peptide | Area Peptide Thioester | Area Cyclic Peptide | Area Hydrolyzed | Sum | TTN | Product/Starting Material | Product/Hydrolysis |
|---|---|---|---|---|---|---|---|---|
| 30 nM | 400 uM | 94021 | 329604 | 2760 | 426385 | 10307 | 3.5 | 119.4 |

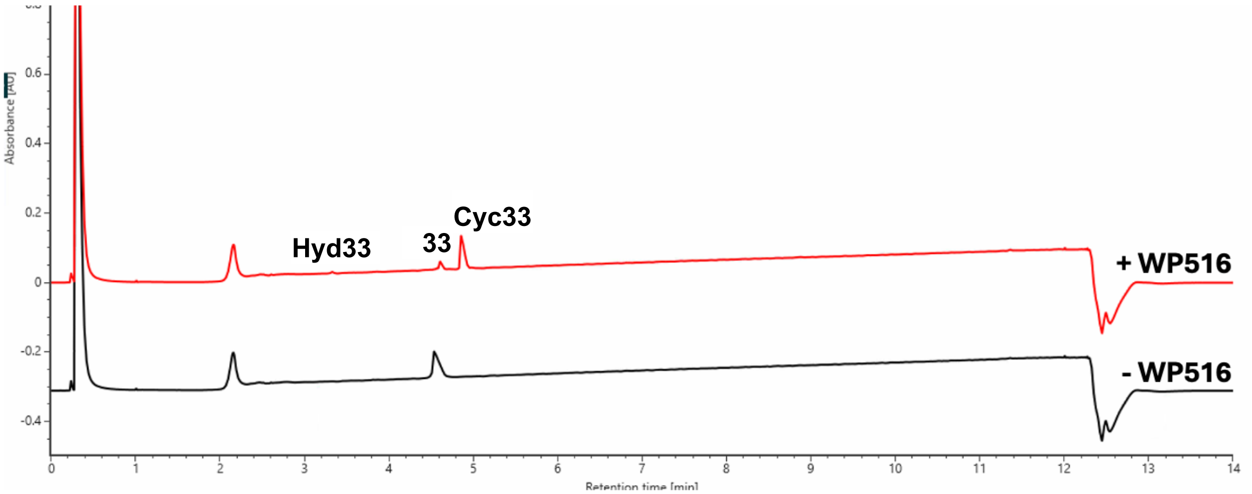

**Figure S46. UPLC UV trace (214 nm) is shown for the total turnover number (TTN) assay of WP516 with tetrapeptide substrate 33.** Enzyme and substrate concentrations are indicated above the trace. The substrate is labeled with its corresponding number from the paper where 'Cyc' and 'Hyd' represent head-to-tail cyclic and hydrolyzed peptide, respectively. The shown UPLC trace is representative of three replicates while the TTN, product/starting material, and product/hydrolysis are averages of three replicates. This reaction was analyzed by loading 2 µL onto a CORTECS T3 Column instead of the typical 5 µL.

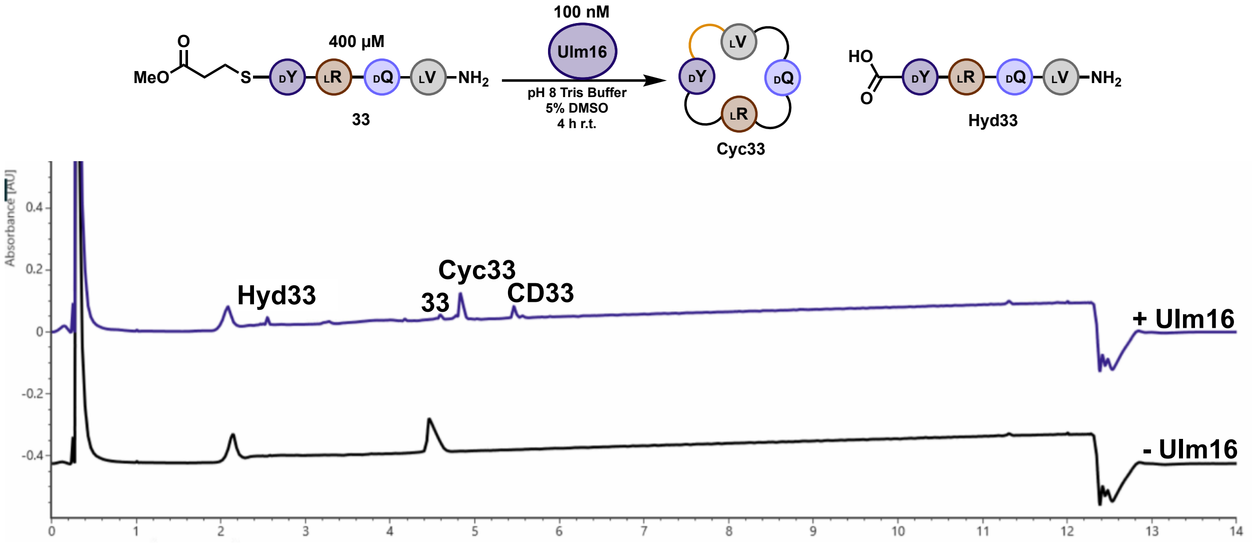

**Figure S47. UPLC UV trace (214 nm) is shown for the reaction of Ulm16 with tetrapeptide substrate 33.** Enzyme and substrate concentrations are indicated above the trace. The substrate is labeled with its corresponding number from the paper where 'Cyc' and 'Hyd' represent head-to-tail cyclic and hydrolyzed peptide, respectively. The shown UPLC trace is representative of three replicates.

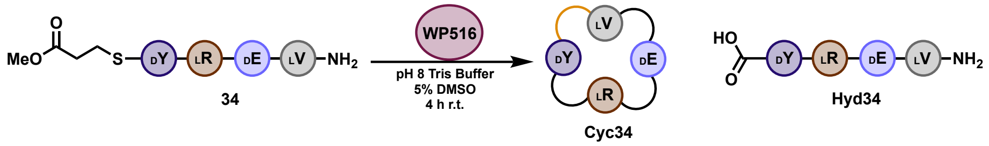

| Enzyme | Peptide | Area Peptide Thioester | Area Cyclic Peptide | Area Hydrolyzed | Sum | TTN | Product/Starting Material | Product/Hydrolysis |
|---|---|---|---|---|---|---|---|---|
| 250 nM | 400 uM | - | 389438 | 8054 | 397492 | - | - | 48.4 |

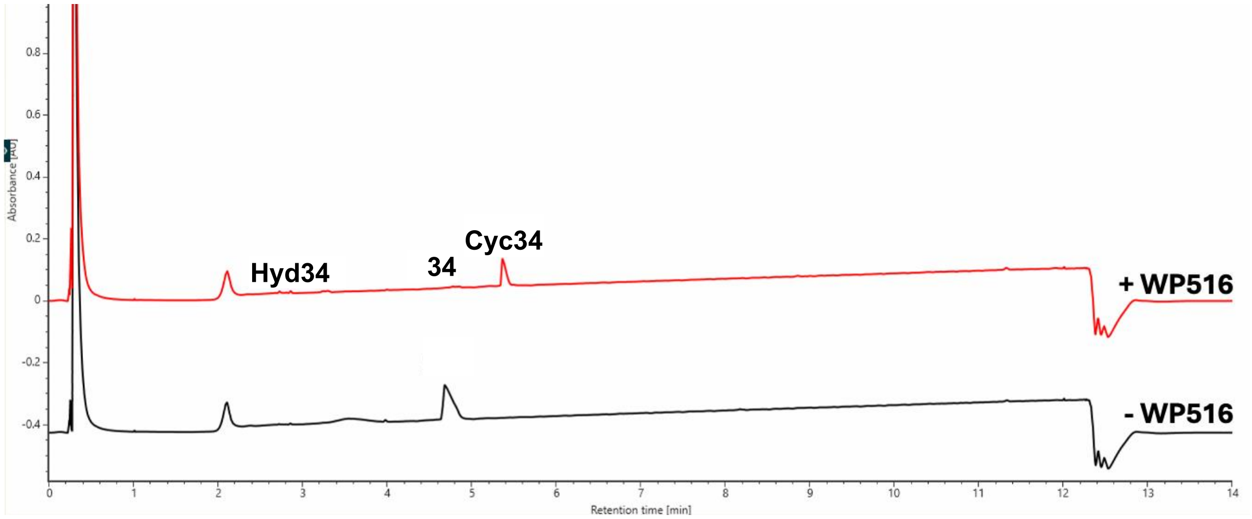

**Figure S48. UPLC UV trace (214 nm) is shown for the total turnover number (TTN) assay of WP516 with tetrapeptide substrate 34.** Enzyme and substrate concentrations are indicated above the trace. The substrate is labeled with its corresponding number from the paper where ‘Cyc’ and ‘Hyd’ represent head-to-tail cyclic and hydrolyzed peptide, respectively. The shown UPLC trace is representative of three replicates while the TTN, product/starting material, and product/hydrolysis are averages of three replicates. The cyclic peptide was too insoluble to determine TTN.

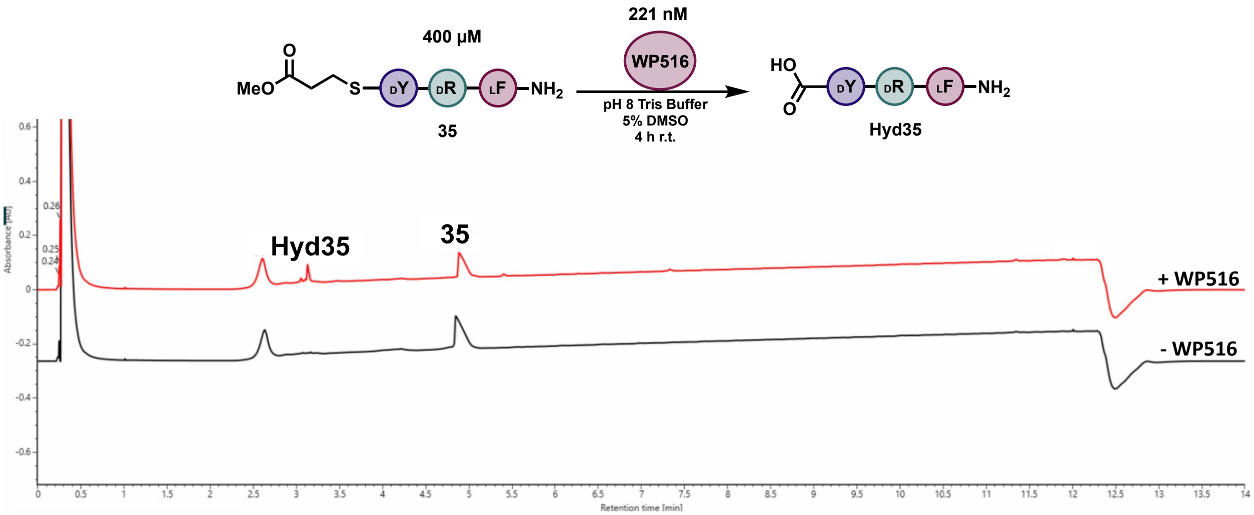

**Figure S49. UPLC UV trace (214 nm) is shown for the total turnover number (TTN) assay of WP516 with tetrapeptide substrate 35.** Enzyme and substrate concentrations are indicated above the trace. The substrate is labeled with its corresponding number from the paper where 'Cyc' and 'Hyd' represent head-to-tail cyclic and hydrolyzed peptide, respectively. The shown UPLC trace is representative of three replicates while the TTN, product/starting material, and product/hydrolysis are averages of three replicates.

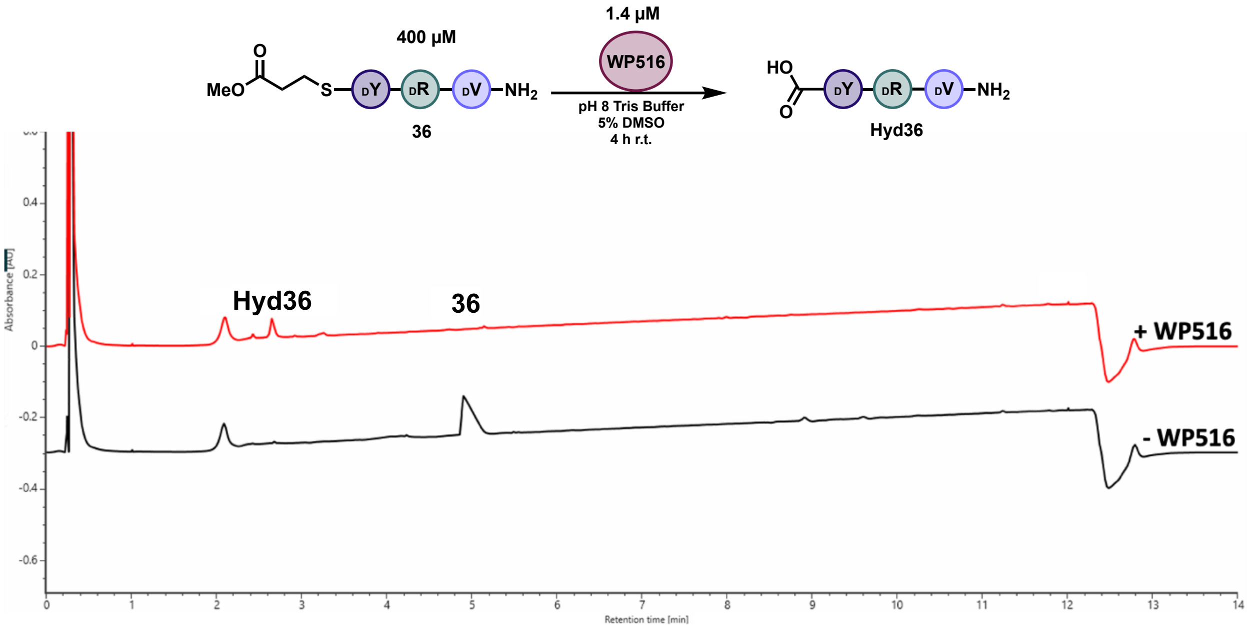

**Figure S50. UPLC UV trace (214 nm) is shown for the total turnover number (TTN) assay of WP516 with tetrapeptide substrate 36.** Enzyme and substrate concentrations are indicated above the trace. The substrate is labeled with its corresponding number from the paper where 'Cyc' and 'Hyd' represent head-to-tail cyclic and hydrolyzed peptide, respectively. The shown UPLC trace is representative of three replicates while the TTN, product/starting material, and product/hydrolysis are averages of three replicates.

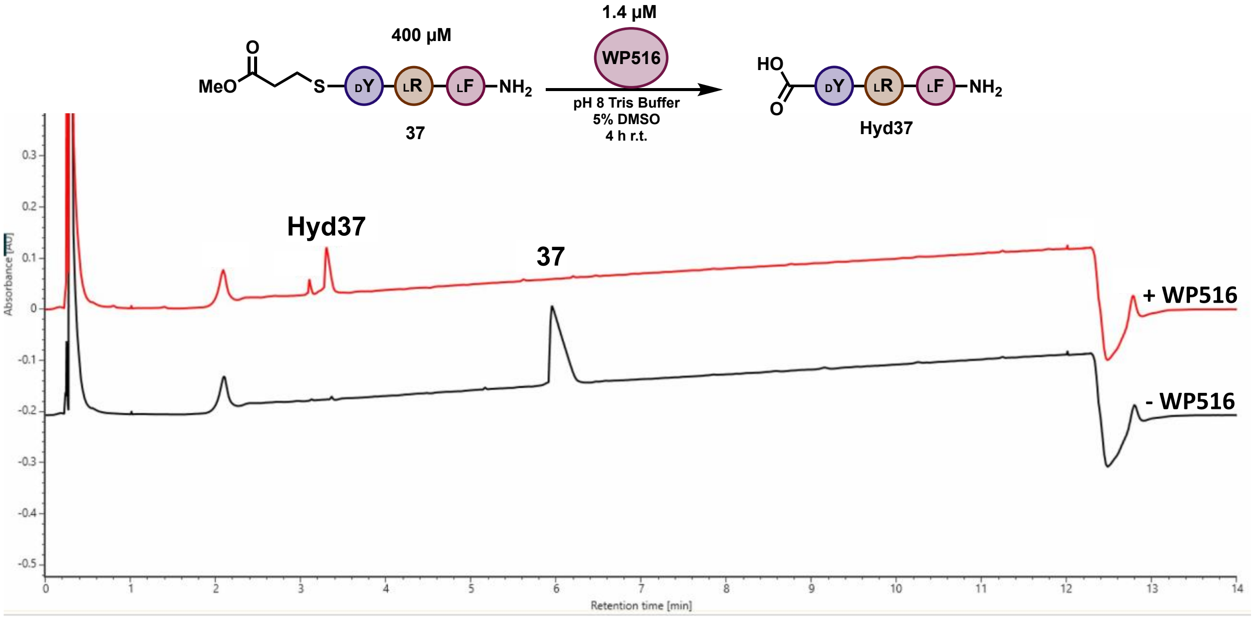

**Figure S51. UPLC UV trace (214 nm) is shown for the total turnover number (TTN) assay of WP516 with tetrapeptide substrate 37.** Enzyme and substrate concentrations are indicated above the trace. The substrate is labeled with its corresponding number from the paper where 'Cyc' and 'Hyd' represent head-to-tail cyclic and hydrolyzed peptide, respectively. The shown UPLC trace is representative of three replicates while the TTN, product/starting material, and product/hydrolysis are averages of three replicates.

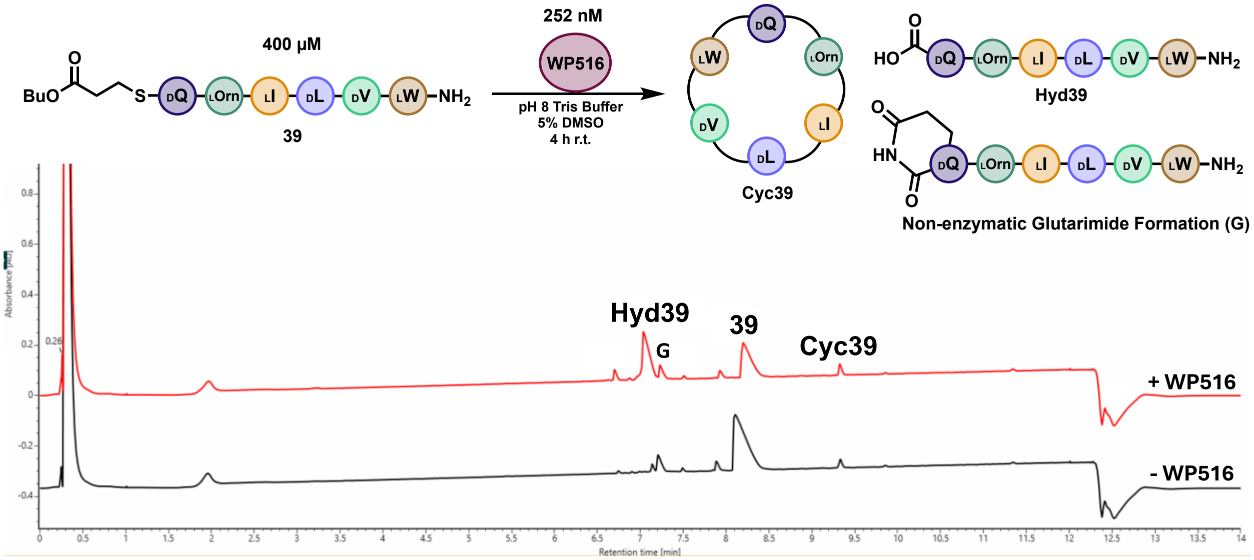

**Figure S52. UPLC UV trace (214 nm) is shown for the total turnover number (TTN) assay of WP516 with peptide substrate 39.** Enzyme and substrate concentrations are indicated above the trace. The substrate is labeled with its corresponding number from the paper where ‘Cyc’ and ‘Hyd’ represent head-to-tail cyclic and hydrolyzed peptide, respectively. The shown UPLC trace is representative of three replicates while the TTN, product/starting material, and product/hydrolysis are averages of three replicates.

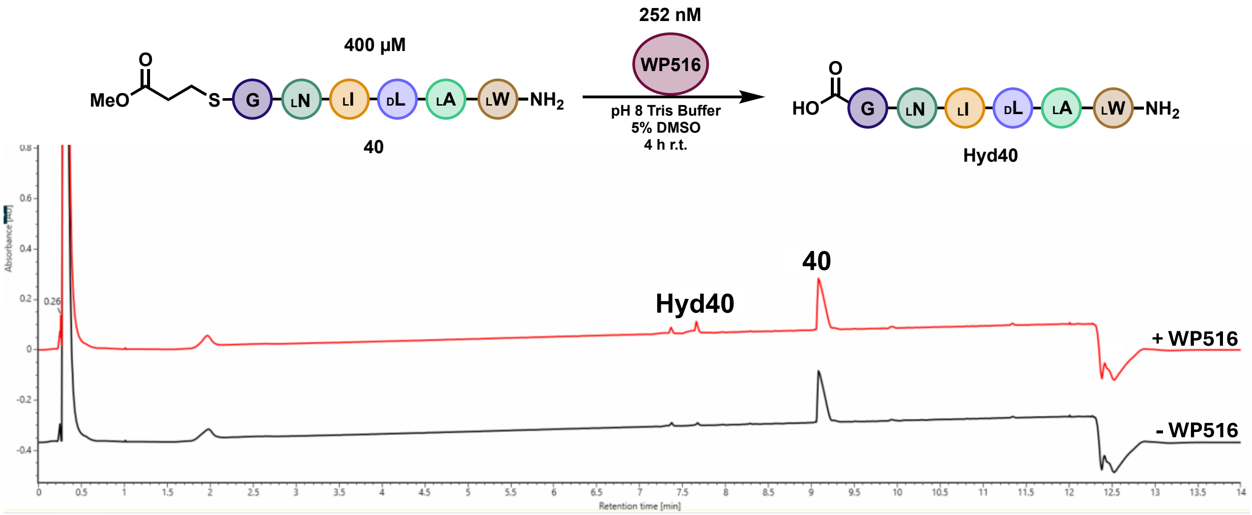

**Figure S53. UPLC UV trace (214 nm) is shown for the total turnover number (TTN) assay of WP516 with peptide substrate 40.** Enzyme and substrate concentrations are indicated above the trace. The substrate is labeled with its corresponding number from the paper where 'Cyc' and 'Hyd' represent head-to-tail cyclic and hydrolyzed peptide, respectively. The shown UPLC trace is representative of three replicates while the TTN, product/starting material, and product/hydrolysis are averages of three replicates.

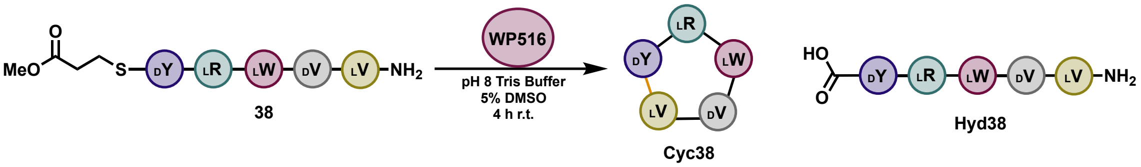

| Enzyme | Peptide | Area Peptide Thioester | Area Cyclic Peptide | Area Hydrolyzed | Sum | TTN | Product/Starting Material | Product/Hydrolysis |
|---|---|---|---|---|---|---|---|---|
| 250 nM | 400 uM | 69426 | 2617223 | 57869 | 2744517 | 1526 | 37.7 | 45.2 |

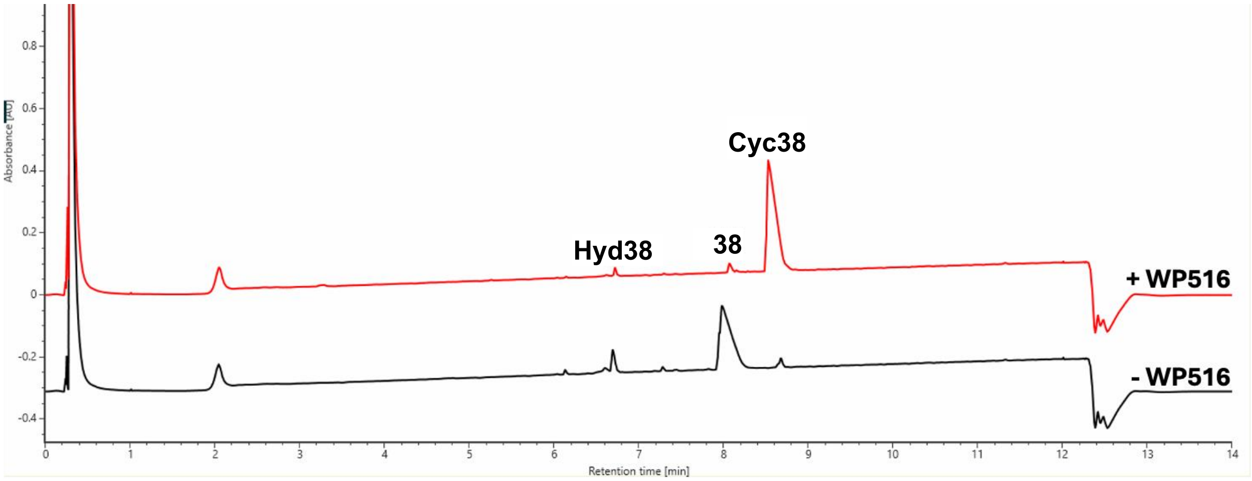

**Figure S54. UPLC UV trace (214 nm) is shown for the total turnover number (TTN) assay of WP516 with peptide substrate 38.** Enzyme and substrate concentrations are indicated above the trace. The substrate is labeled with its corresponding number from the paper where 'Cyc' and 'Hyd' represent head-to-tail cyclic and hydrolyzed peptide, respectively. The shown UPLC trace is representative of three replicates while the TTN, product/starting material, and product/hydrolysis are averages of three replicates. Note: In the +WP516 sample, there is less hydrolysis than in the no enzyme control.

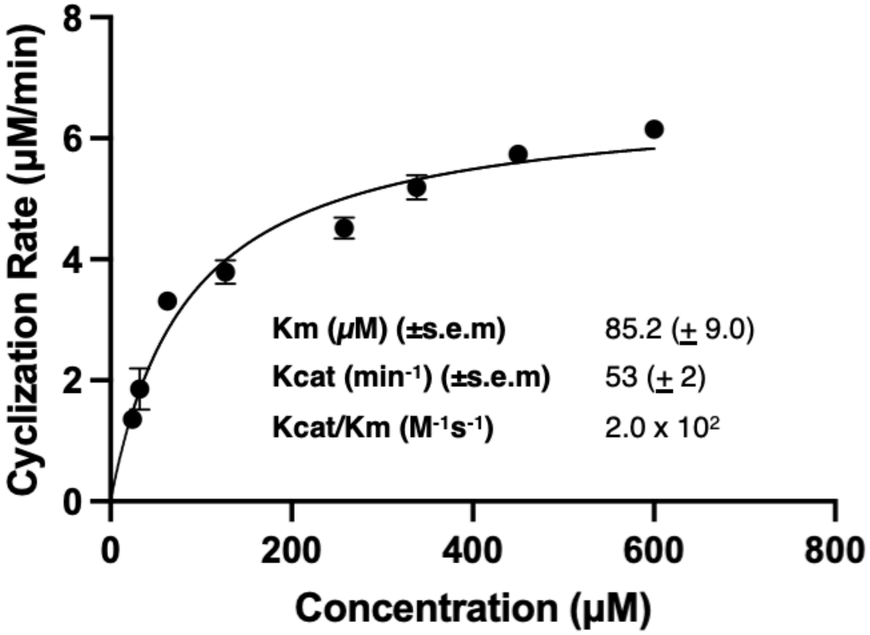

**Figure S55. Michaelis-Menten plots of WP516 incubated with pentapeptide 38.** The enzyme concentration employed is provided in the methods section. The plots represent the mean of triplicate experiments, and the error bars indicate the standard error of the mean (S.E.M).

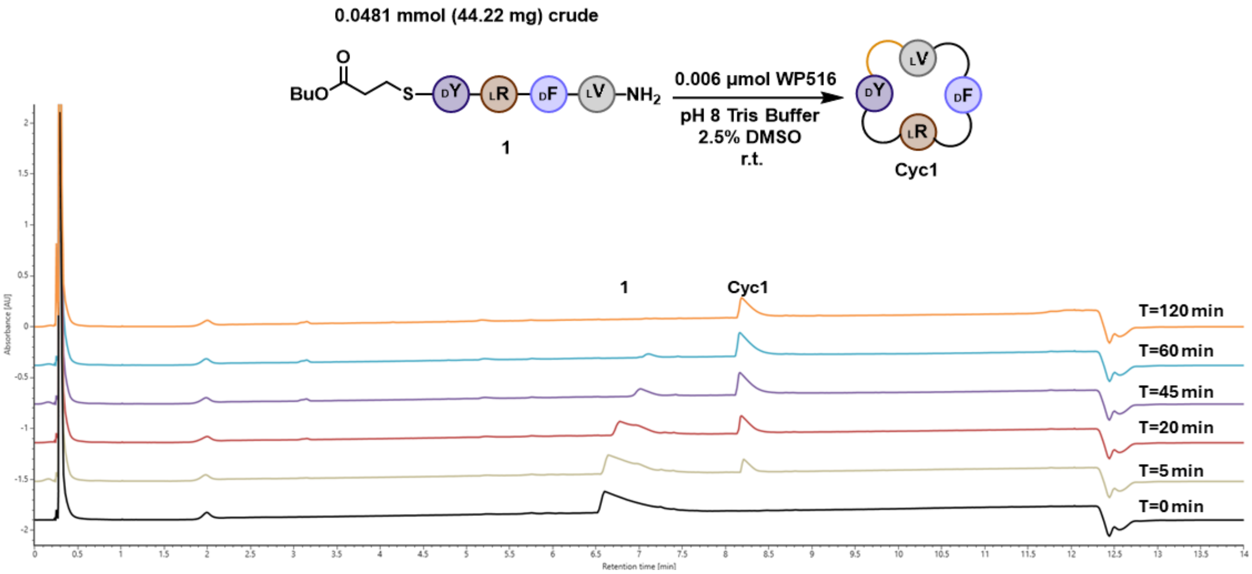

**Figure S56. Reaction time course of scale up with peptide 1.** 50 µL Aliquots taken from reaction scale up. The substrate is labeled with its corresponding number from the paper where 'Cyc' represent head-to-tail cyclic peptide. All substrate was consumed after 2 hours (120 minutes).

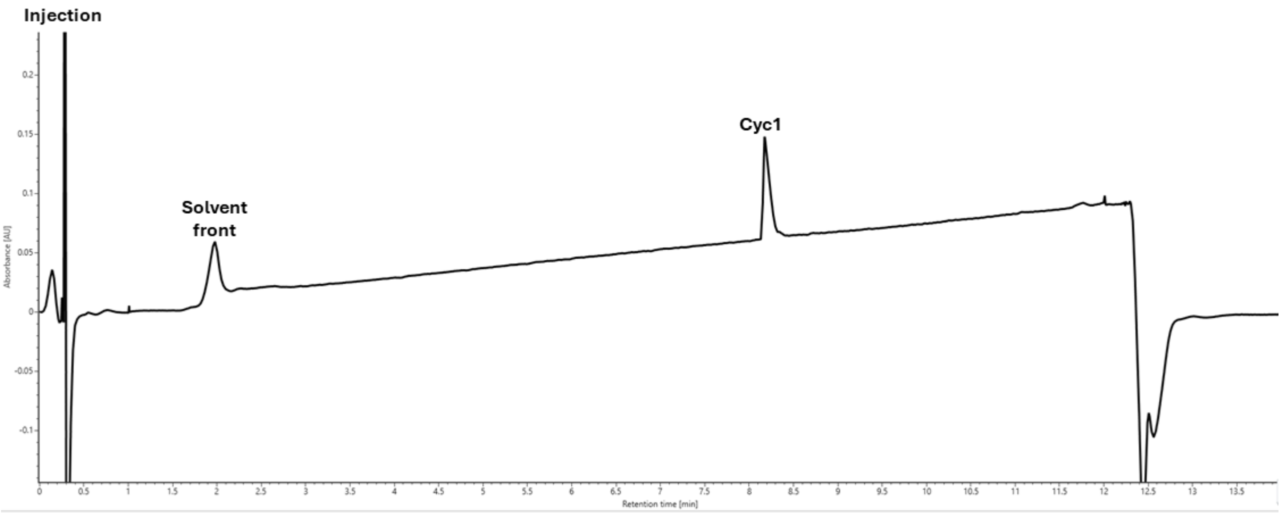

**Figure S57. UPLC UV trace of Cyclic peptide 1 (Cyc1) after HPLC purification, and lyophilization.** Product was >99% pure by measurement at 214 nm wavelength. See Figure S85 for MS/MS and Figure S58 for NMR.

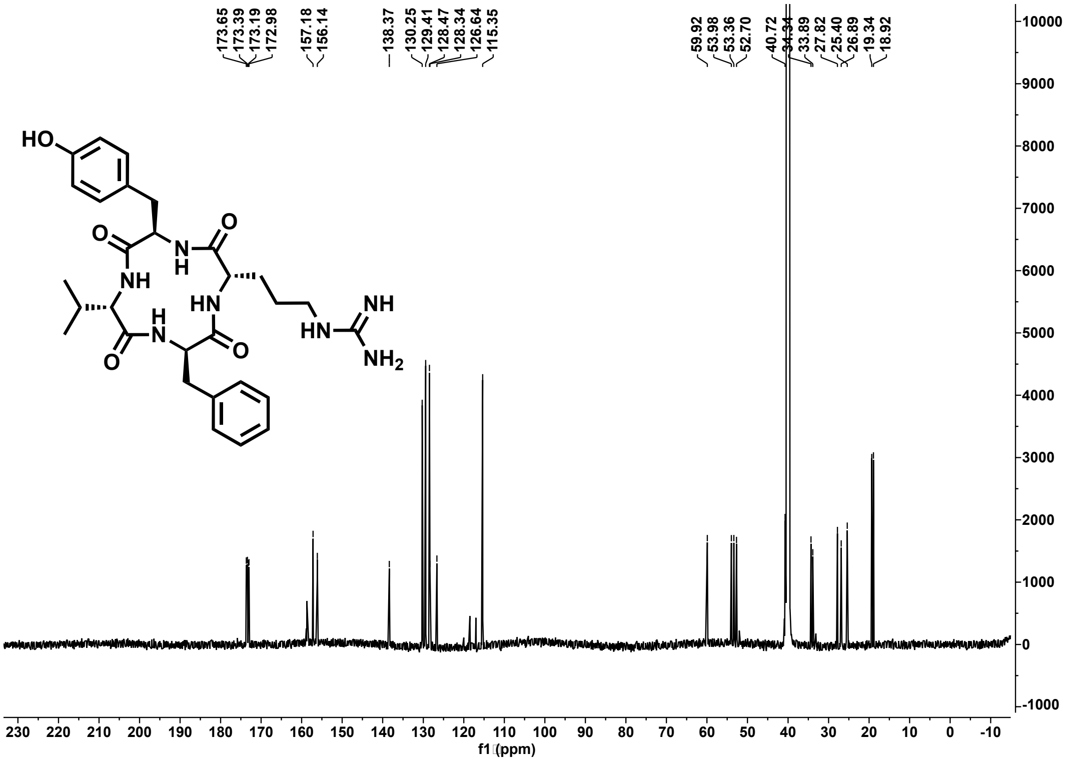

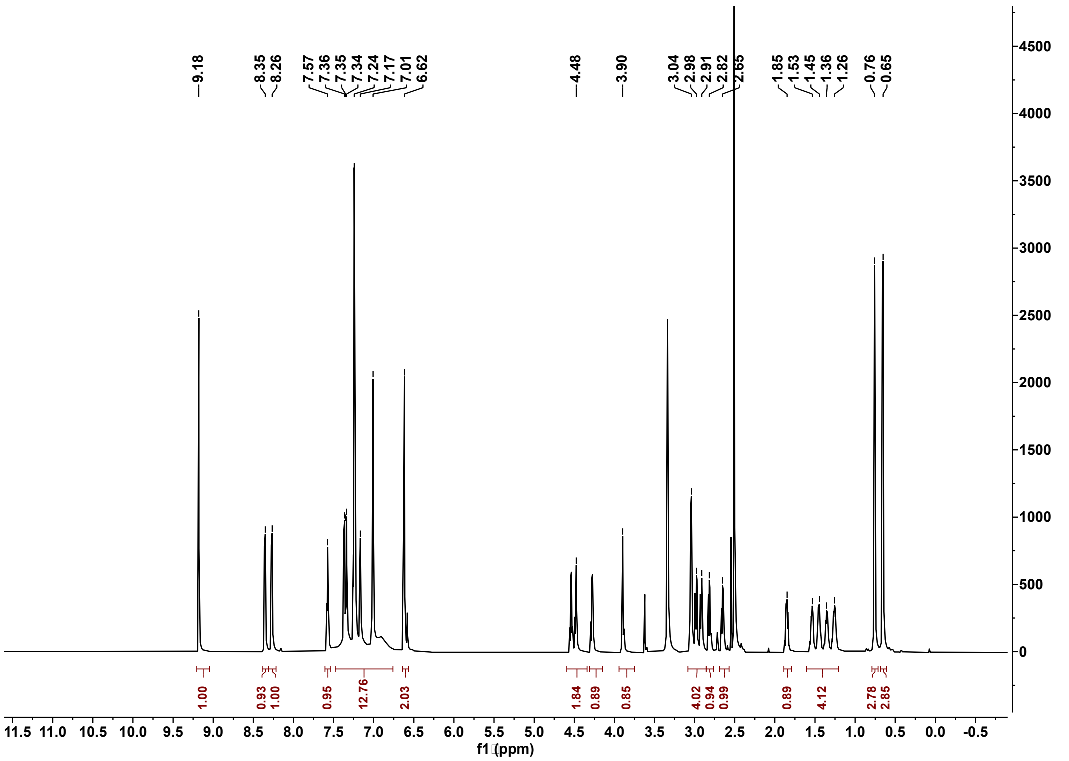

Figure S58. $^{13}C$ (201 MHz) and $^{1}H$ (800 MHz) NMR spectra of cyclic peptide 1 in DMSO-$d_6$.

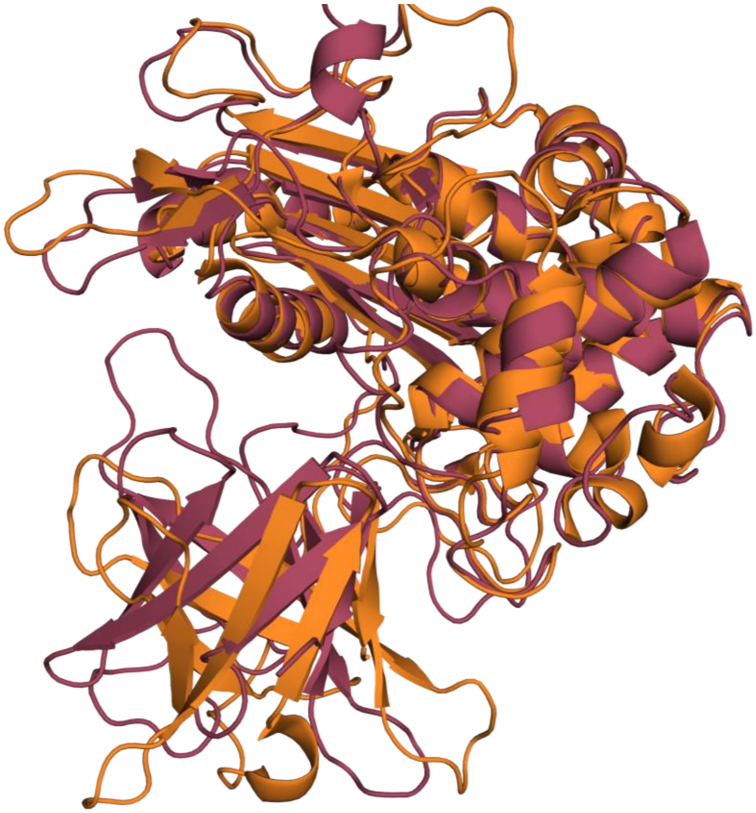

**Figure S59. AF3 Model comparison to SurE.** WP516 AlphaFold3 model (red) overlayed with SurE crystal structure (PDB: 6KSU, orange) highlighting the differences in lipocalin domain angle. RMSD: 1.292

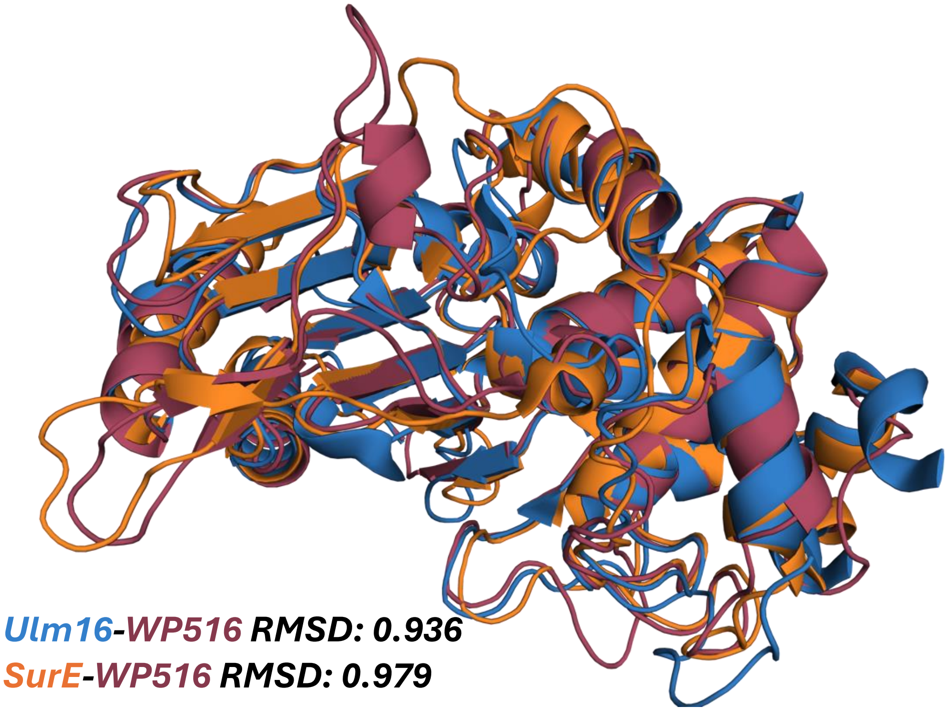

**Figure S60. Overlay of WP516, Ulm16, and SurE α-β-hydrolase domains.** All three proteins have highly similar α-β-hydrolase domains despite the fact they differ widely in peptide substrate scope (SurE 5-22 amino acids, Ulm16 4-6 amino acids, and WP516 4-5 amino acids).

**A**

```
SurE_XDVG       FGLADGWGLGLMRHGTGDGAWYGHDGAVGGASCNLRIHPDRSLALALTANSTAGPKLWEA    60
Ulm16_XDVG      FGLADGWGPGLGRFGPAGNRWLGHDGTLDGATCHLRIHPGRGTVVALTTNSPTGQALWDA    60
WP516_XDVG      FGLADGWGLGWSLYGTASGTWFGHDGTGDGAWCHLRVQPETGTAVALTTNGGNGAVLWEA    60
                ******** *   .* ... * ****: .** *:**::*  . .:***:*.  *  **:*

SurE_XDVG       LVARLPEAGLDVG   73
Ulm16_XDVG      VVDALRDADIDVG   73
WP516_XDVG      VVAELRAAGVDVG   73
                :*  *  *.:***
```

**B**

```
Ulm16_Lipocalin         -------VHRPAPP-PAIAAAAFADCTGTYRNGDLAVTVGIDG-PYLVLELPGGARELAQ
SurE_Lipocalin          HYAL--------PVPDSAPLAPDAGHLGTYANGDLELMVTHDAAGDLFLTRESYSDYRLS
WP516_Lipocalin         HYPLGALTGGGTPLPAGS--PAKDGCAGRYANGSWTCAV-EASGGELYLSVGPGPRWRLR
                                    *   .       .  * * **.    *   .   * *

Ulm16_Lipocalin         PLAHRTFS-----SRGAGFLGRFVTDARSDAVHALQYSGRTLLREAGES  95
SurE_Lipocalin          LHEDDLFVARSGEPGALPITGRFVREHPAGPVALLQYGGRAMHRL----  97
WP516_Lipocalin         CFEDLRFTTEGSDAGSMPYVGRFLRDPGSGTVELVQITGRLARRHG---  103
                           .  *        .    ***: :  :. *  :*  **   *
```

**C**

| % AA ID | SurE | Ulm16 | WP516 |
|---|---|---|---|
| SurE | - | 56.16 | 57.53 |
| Ulm16 | 56.16 | - | 61.64 |
| WP516 | 57.53 | 61.64 | - |

**D**

| % AA ID | SurE | Ulm16 | WP516 |
|---|---|---|---|
| SurE | - | 30.23 | 35.11 |
| Ulm16 | 30.23 | - | 25.84 |
| WP516 | 35.11 | 25.84 | - |

**Figure S61. Clustal Omega multiple sequence alignment (MSA) results. A)** MSA output for highly homologous 73 amino acid region leading into the lipocalin domain for SurE, Ulm16, and WP516. **B)** MSA output for the unstructured loop and lipocalin domains of Ulm16, SurE, and WP516. **C)** Percent identity matrix for 73 amino acid region leading into the lipocalin domain. **D)** Percent identity matrix for unstructured loop and lipocalin domain

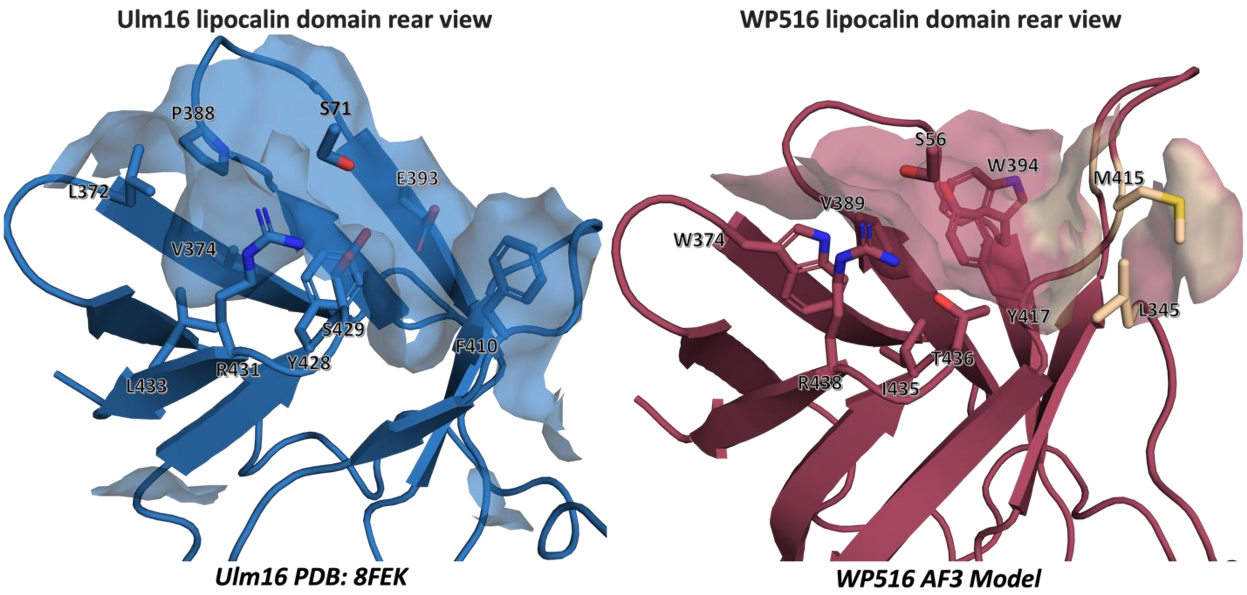

```
Ulm16_Lipocalin      ------VHRPAPPPAIAAAAFADCTGTYRNGDLAVTVGIDGPYLVLELPGGARELAQPLA      54
WP516_Lipocalin      HYPLGALTGGGTPLPAGSPAKDGCAGRYANGSWTCAVEASGGELYLSVGPGPRWRLRCFE      60

Ulm16_Lipocalin      HRTFSS-----RGAGFLGRFVTDARSDAVHALQYSGRTLLREAGES      95
WP516_Lipocalin      DLRFTTEGSDAGSMPYVGRFLRDPGSGTVELVQITGRLARRHG---      103
```

**Figure S62. Lipocalin domain comparison between Ulm16 and WP516. A.)** Lipocalin domain of Ulm16 crystal structure (blue) and WP516 Alpha Fold3 model (red). The selected residues have been shown as a surface model to reflect how WP516's lipocalin domain results in only 1 pocket for the peptide to travel into (between Y417 and M415;L345) while Ulm16 displays multiple cavities. **B.)** MSA output of Ulm16 and WP516 lipocalin domain and loop. Residues that make the lipocalin domain surface are highlighted.

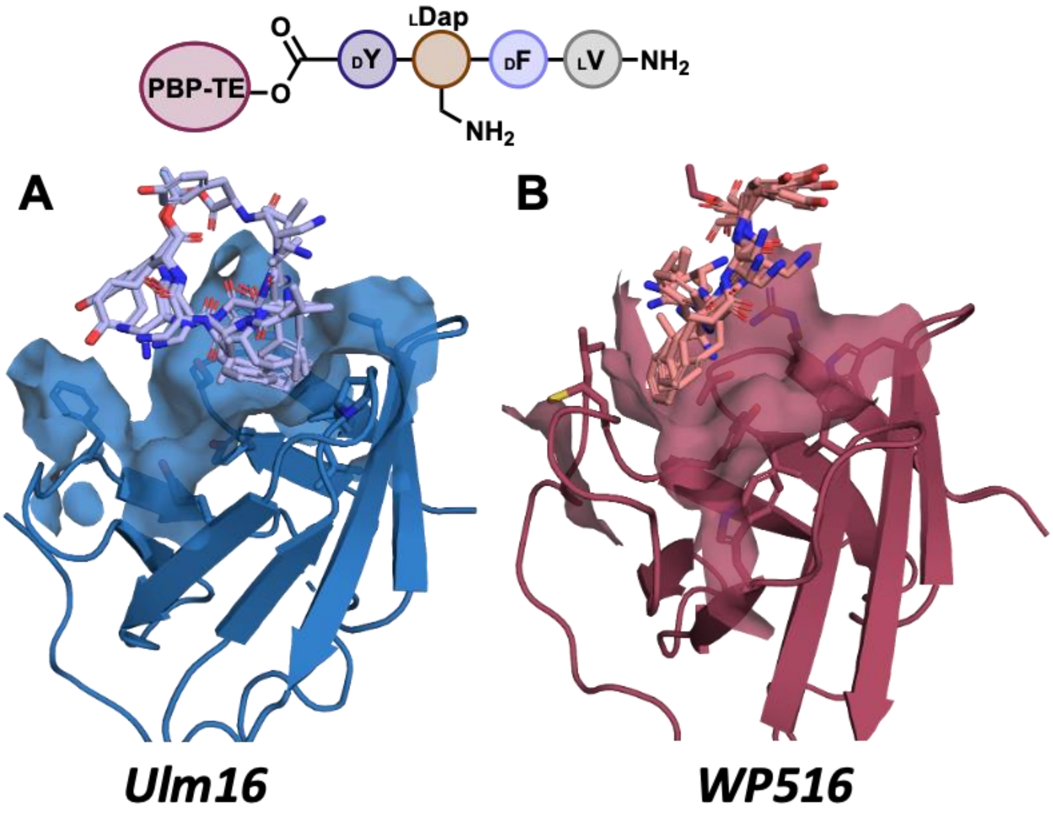

**Figure S63. Top 5 Molecular Mechanics with Generalized Born and Surface Area Solvation (MM/GBSA) scoring outputs from Schrodinger peptide 5 covalent docking. A.)** Peptide 5 covalently docked to Ulm16 crystal structure (PDB: 8FEK) showing the peptide traveling into a deep binding pocket made up of residues L372, L386, P387, Y428, S429, R431 preventing the peptide from adopting a cyclic conformation. **B.)** Peptide 5 covalently docked to the WP516 Alpha Fold3 model showing how the addition of Trp 374 fills that pocket and forces the peptide into a shallower binding pocket made up of L345, W394 M415, Y418, and T436, resulting in the peptide adopting a cyclic conformation.

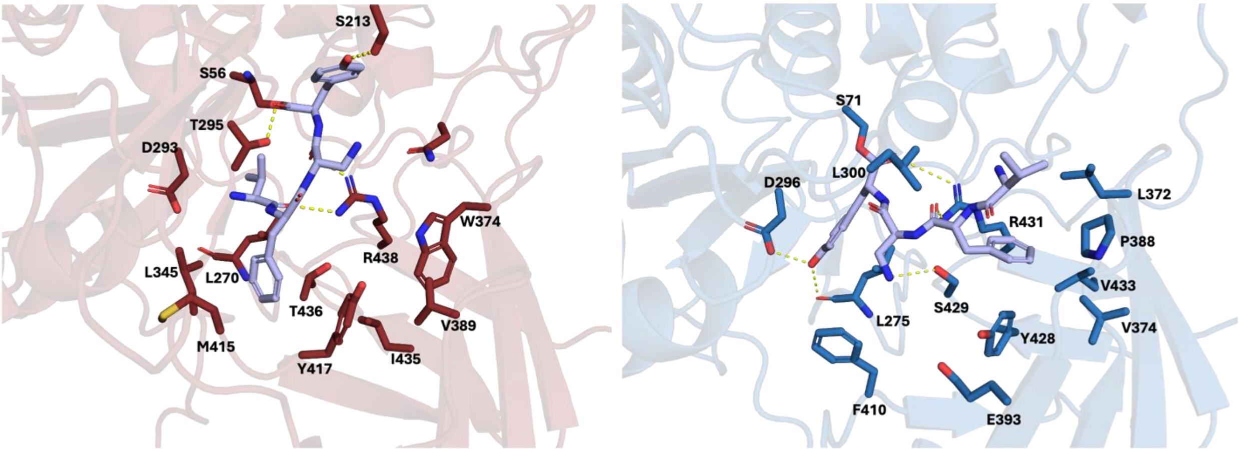

**Figure S64. Covalent docking of peptide 5.** with A.) WP516; W394 of the lipocalin domain has been omitted for clarity B.) Ulm16. Residues within hydrogen bonding distance or those that play a role in the lipocalin surface are highlighted.

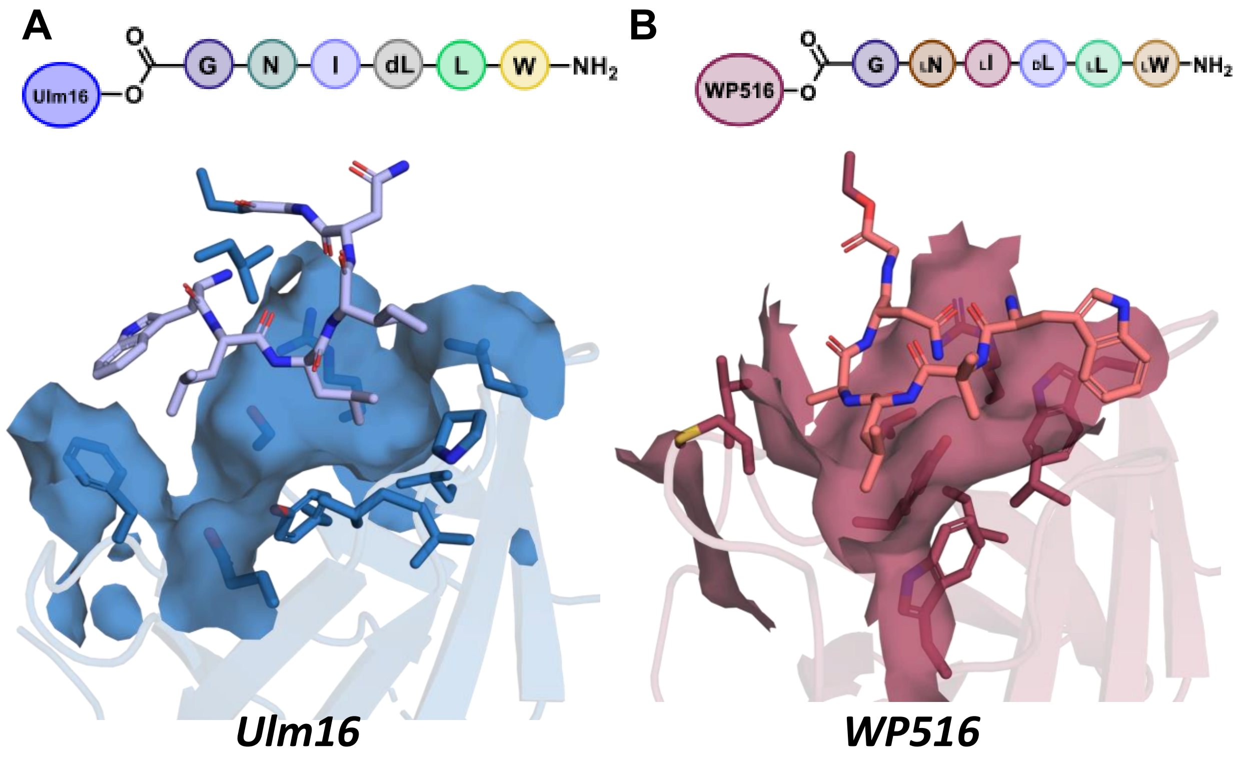

**Figure S65. Top MMGBSA scoring output from Schrodinger peptide 40 covalent docking. A.)** Peptide **40** covalently docked to Ulm16 crystal structure (PDB: 8FEK) showing the peptide traveling into the binding pocket made up of residues L372, L386, P387, Y428, S429, R431. Unlike with the tetrapeptides, the additional 2 amino acids of the hexapeptide **40** are forced to turn back allowing the peptide to take a cyclic conformation **B.)** Peptide **40** covalently docked to the WP516 AF3 model. The increased bulk of the WP516 lipocalin domain does not force the peptide to turn back on itself . Instead, the peptide leaves the lipocalin domain, shedding light on why longer peptides are not cyclized by this enzyme.

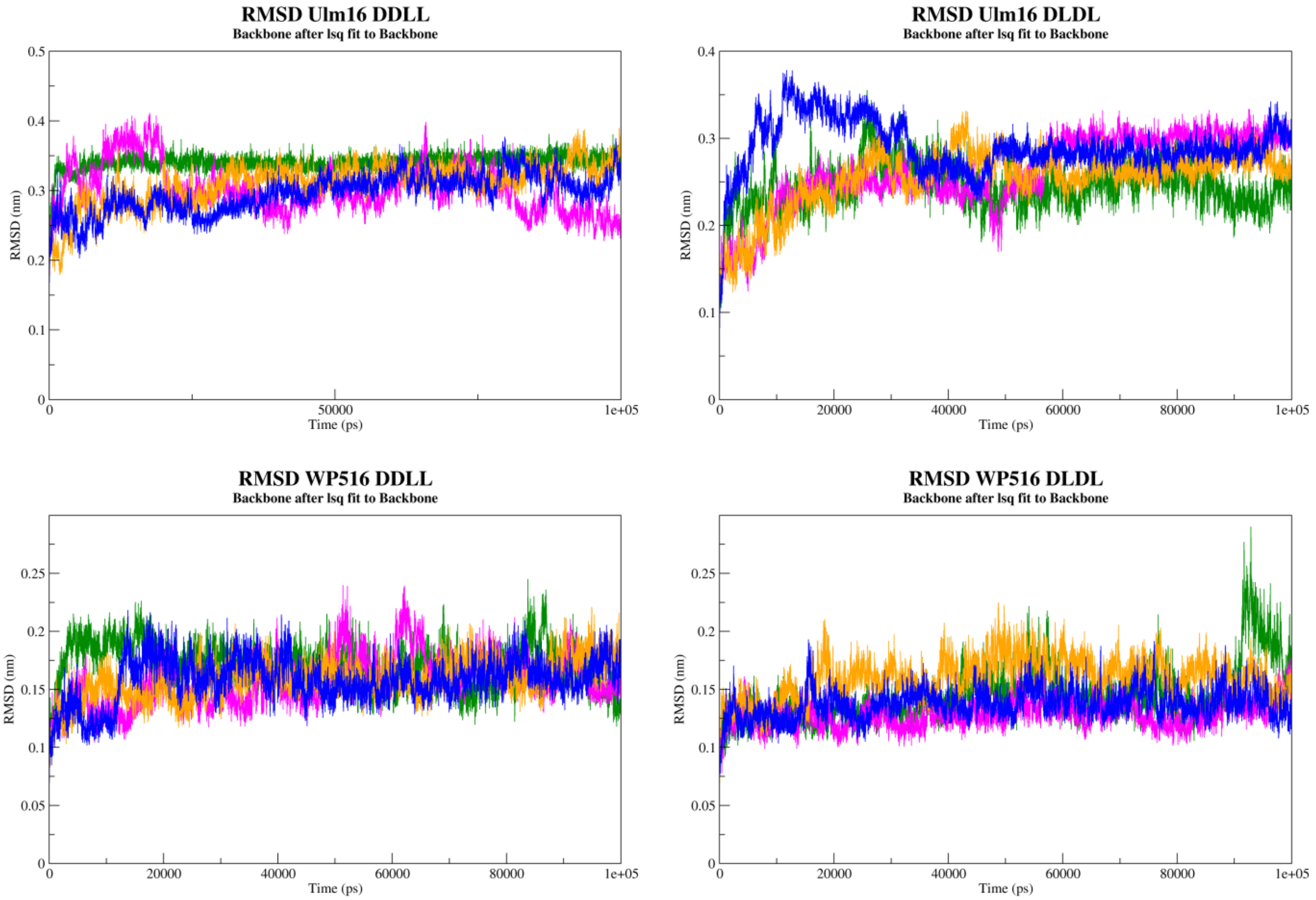

**Figure S66. RMSD of protein backbone over the course of MD simulations.** Each independent replicate of each system was plotted. Peptides that were examined were 3 (DDLL) and 1 (DLDL).

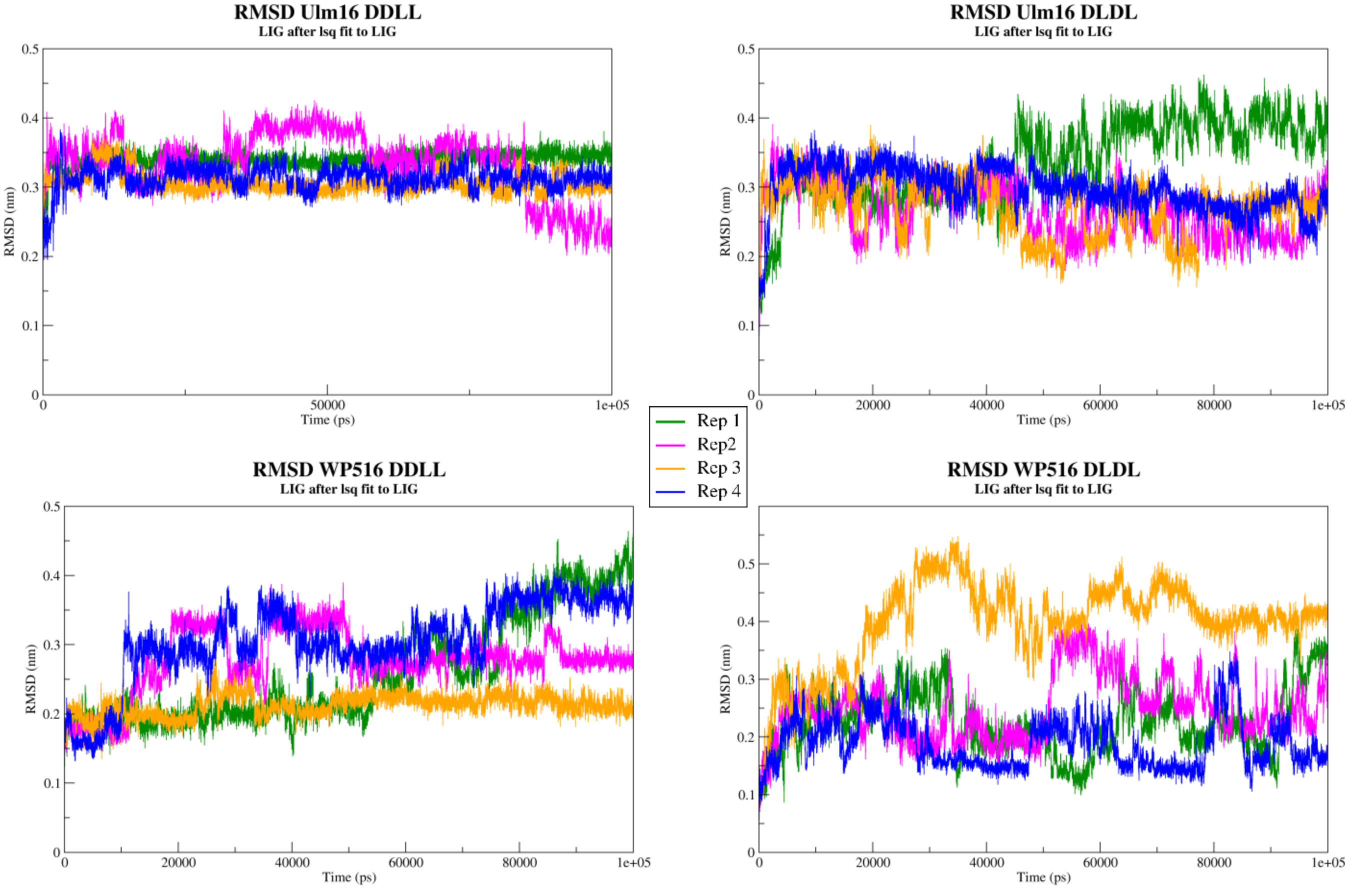

**Figure S67. RMSD of peptide substrate over the course of MD simulations.** Each independent replicate of each system was plotted. Peptides that were examined were 3 (DDLL) and 1 (DLDL).

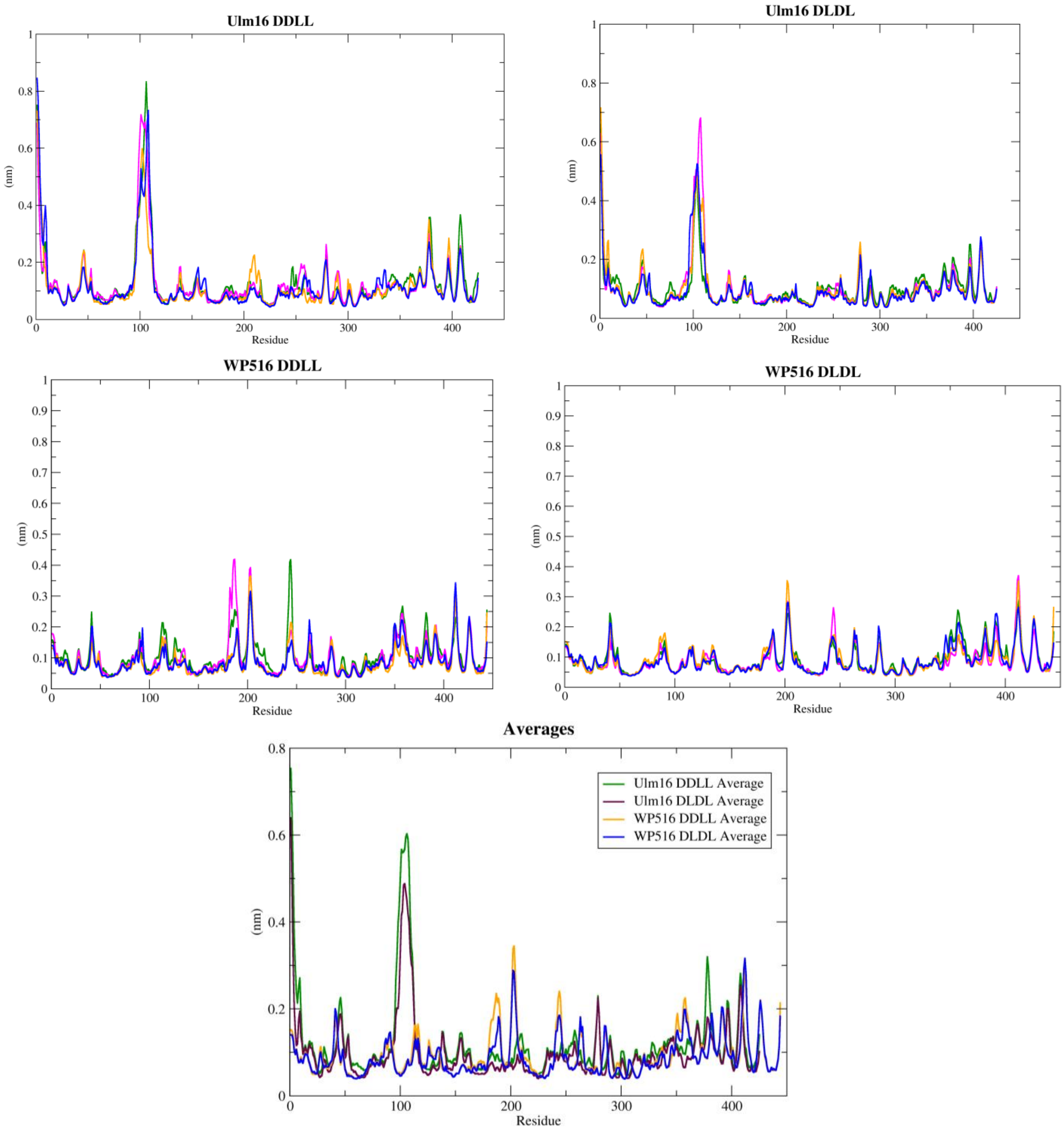

**Figure S68. RMS fluctuation of protein backbone by residue over the course of MD simulations.** Each independent replicate of each system was plotted in addition to the average of the four replicates. Peptides that were examined were 3 (DDLL) and 1 (DLDL).

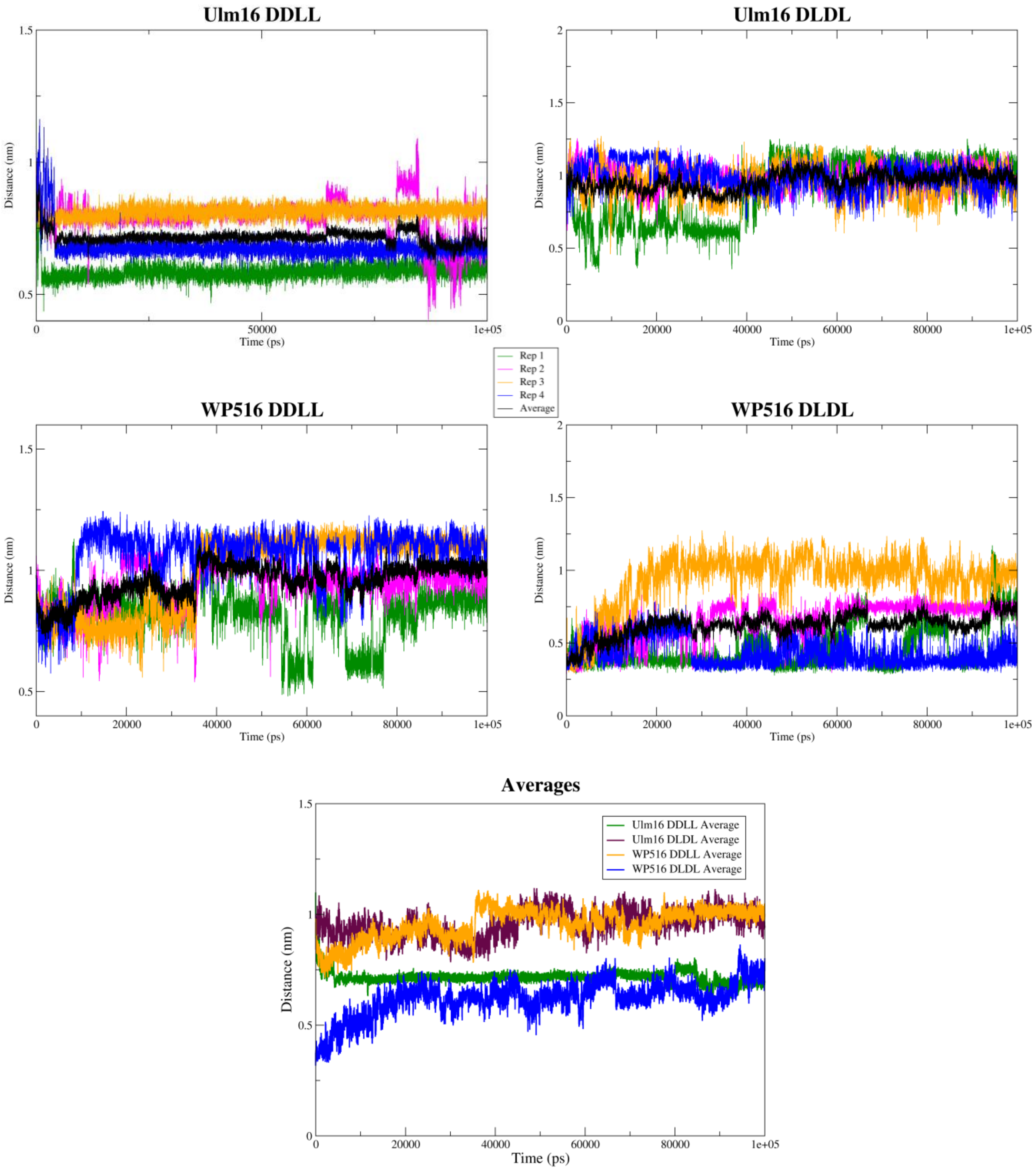

**Figure S69. The distance between the N-terminal N and C-terminal carbonyl C of the peptide substrate over the course of the MD simulations.** Each independent replicate of each system was plotted in addition to the average of the four replicates. Peptides that were examined were 3 (DDLL) and 1 (DLDL).

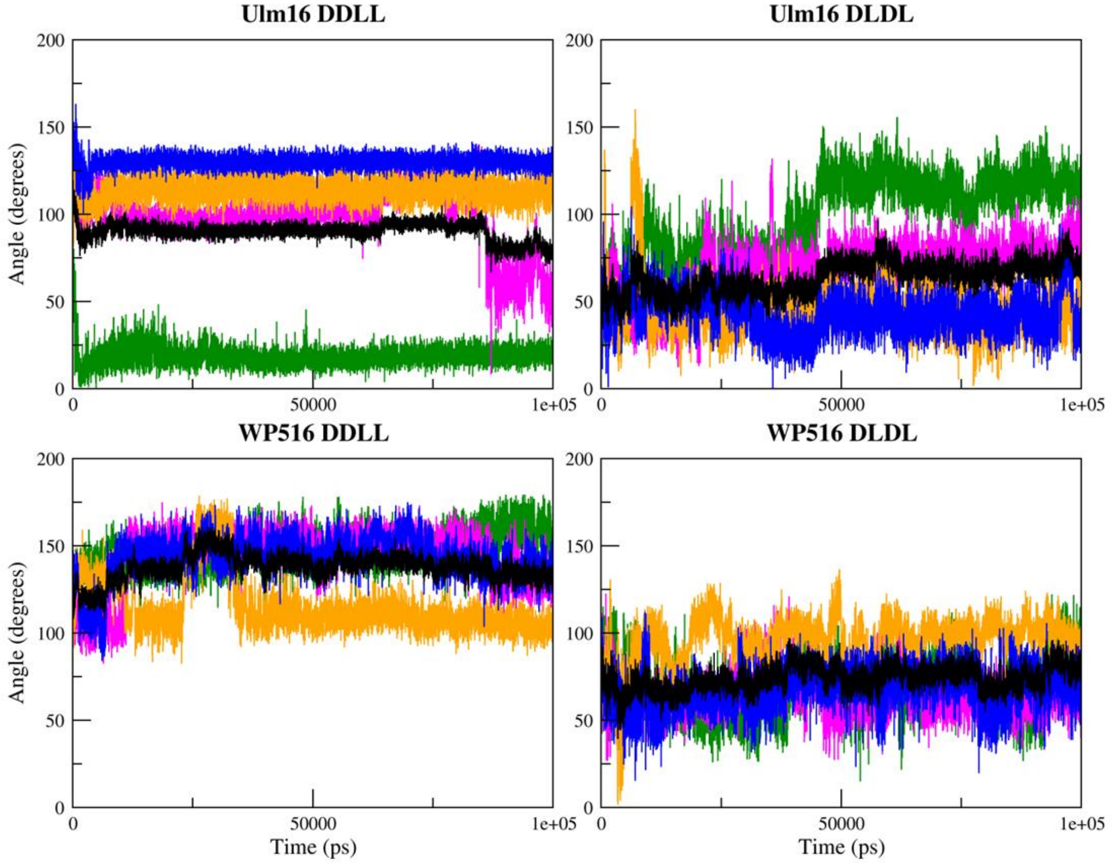

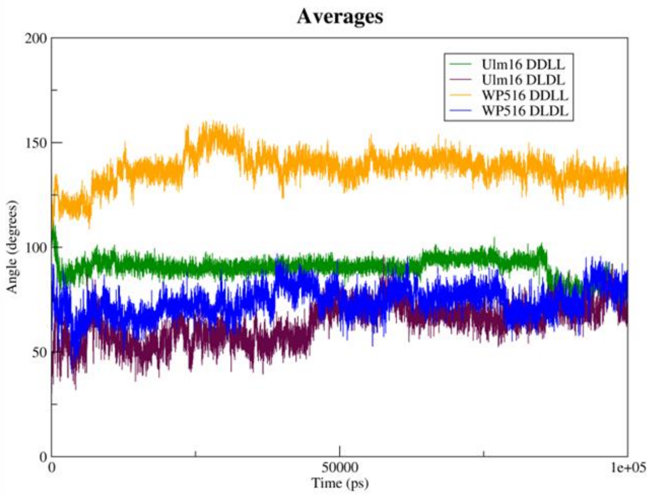

**Figure S70. The angle between the N-terminal N, C-terminal carbonyl C, and C-terminal carbonyl O of the peptide substrate over the course of the MD simulations.** Each independent replicate of each system was plotted in addition to the average of the four replicates. Peptides that were examined were 3 (DDLL) and 1 (DLDL).

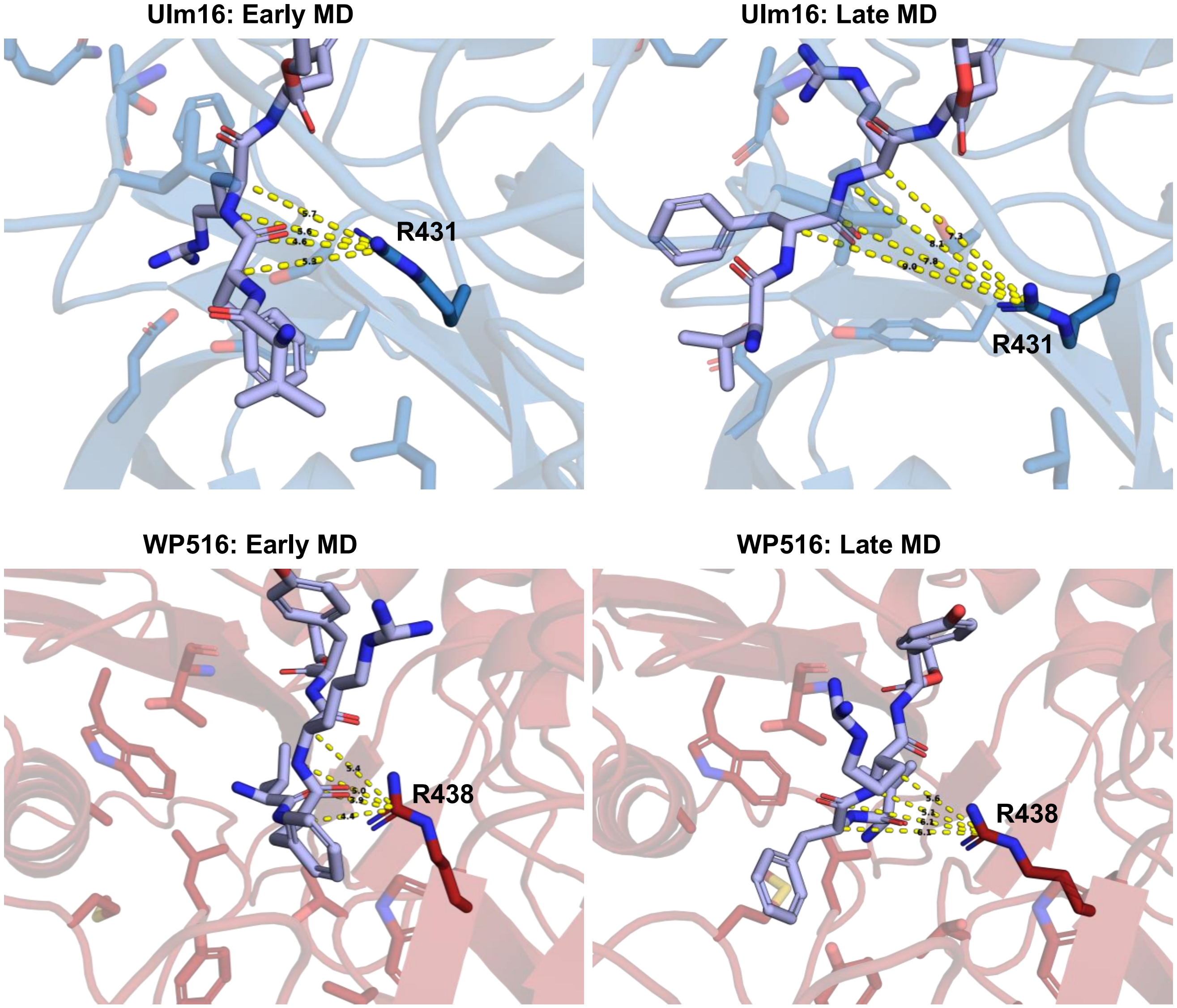

**Figure S71. Key frames from MD simulations with peptide 1 docked to Ulm16 (top, blue) and WP516 (bottom, red) showing distance between substrate and R431/R438.** The distance between atoms of the substrate backbone and the central carbon of the arginine were measured. For Ulm16 the average early frame distance was 5.45 Å, and late frame distance was 8.3 Å. For. WP516 average early frame distance was 4.7 Å, and late frame average was distance was 5.73 Å. The distances get larger over the course of the simulation as the substrate shifts toward β-strands 6 and 7.

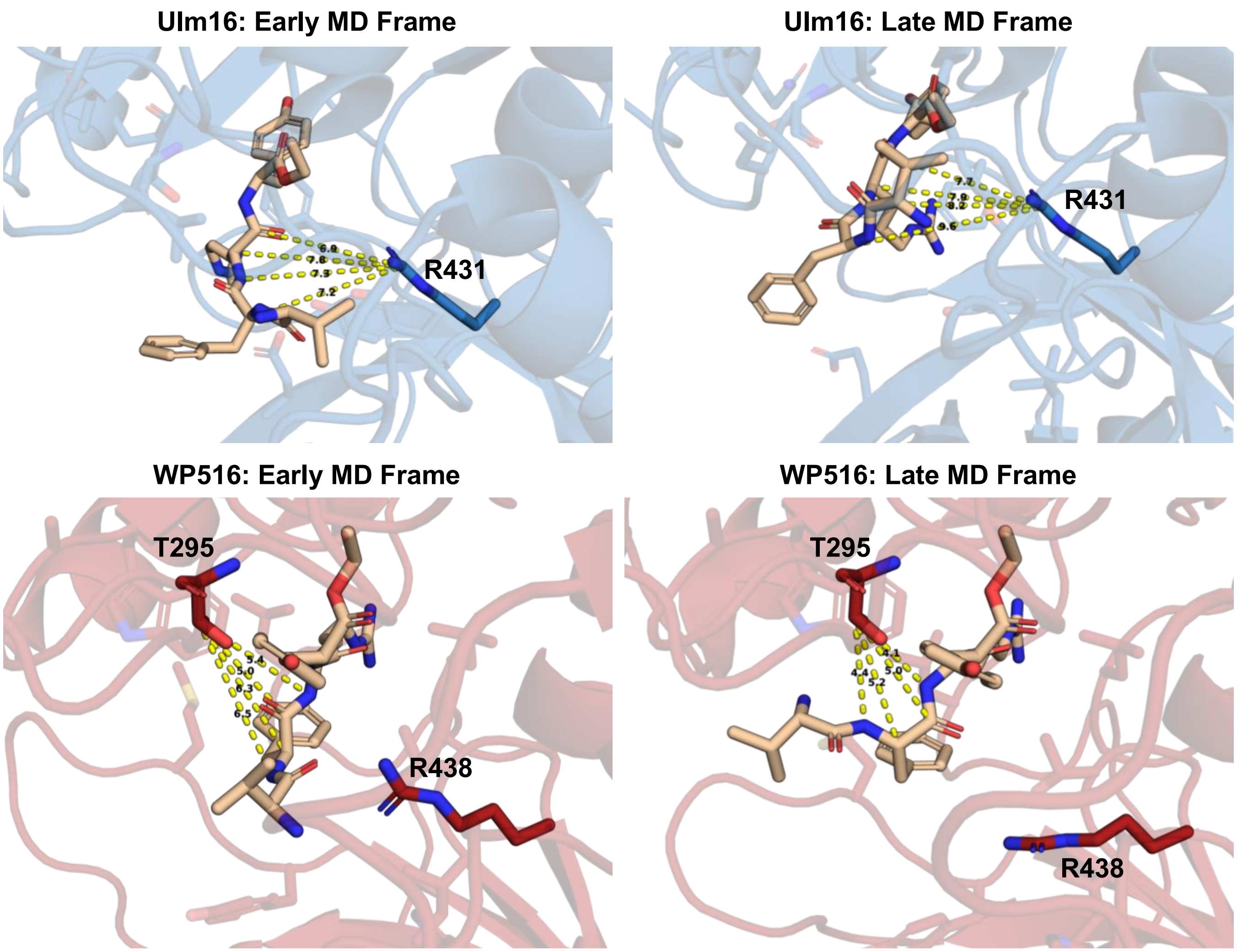

**Figure S72. Key frames from MD simulations with peptide 3 docked to Ulm16 (top, blue) and WP516 (bottom, red) showing substrate migration toward the α/β-hydrolase domain.** For Ulm16, the distance between atoms of the substrate backbone and the central carbon of R431 were measured (early frame average = 7.25 Å, late frame average = 8.35 Å). Due to the movement of R438 in WP516, the distance to T295 from the substrate was measured instead (early frame average = 5.8 Å, late frame average = 4.43 Å). The distances over the course of the simulation reflect the substrate shifting toward β-strands 6 and 7.

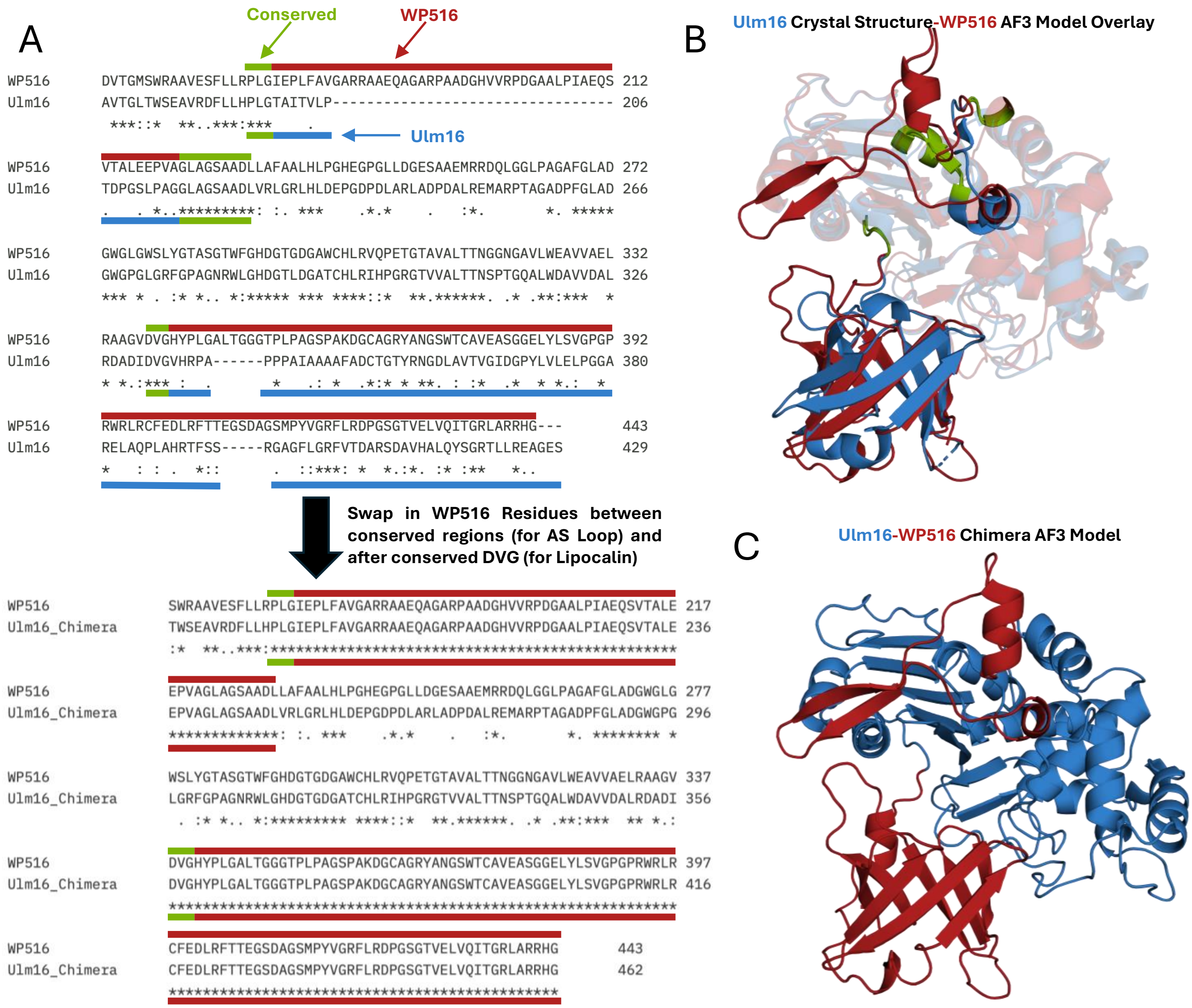

**Figure S73. Making of Ulm16 Chimera (Ulm16 C).** A) Partial multiple sequence alignment of WP516, and Ulm16 Chimera, with green representing either conserved regions leading into the active site loop, or lipocalin. Red lines represent amino acids native to WP516, and blue represent Ulm16 amino acids native to Ulm16. Between conserved regions Ulm16 amino acids were replaced for WP516 leading to creation of the Ulm16 chimera. B) comparison of WP516 AF3 model (red), and Ulm16 crystal structure (8FEK, blue) with active site loop, and lipocalins highlighted. C) AF3 model of Ulm16 Chimera with WP516s portions in red, and Ulm16 portions in Blue.

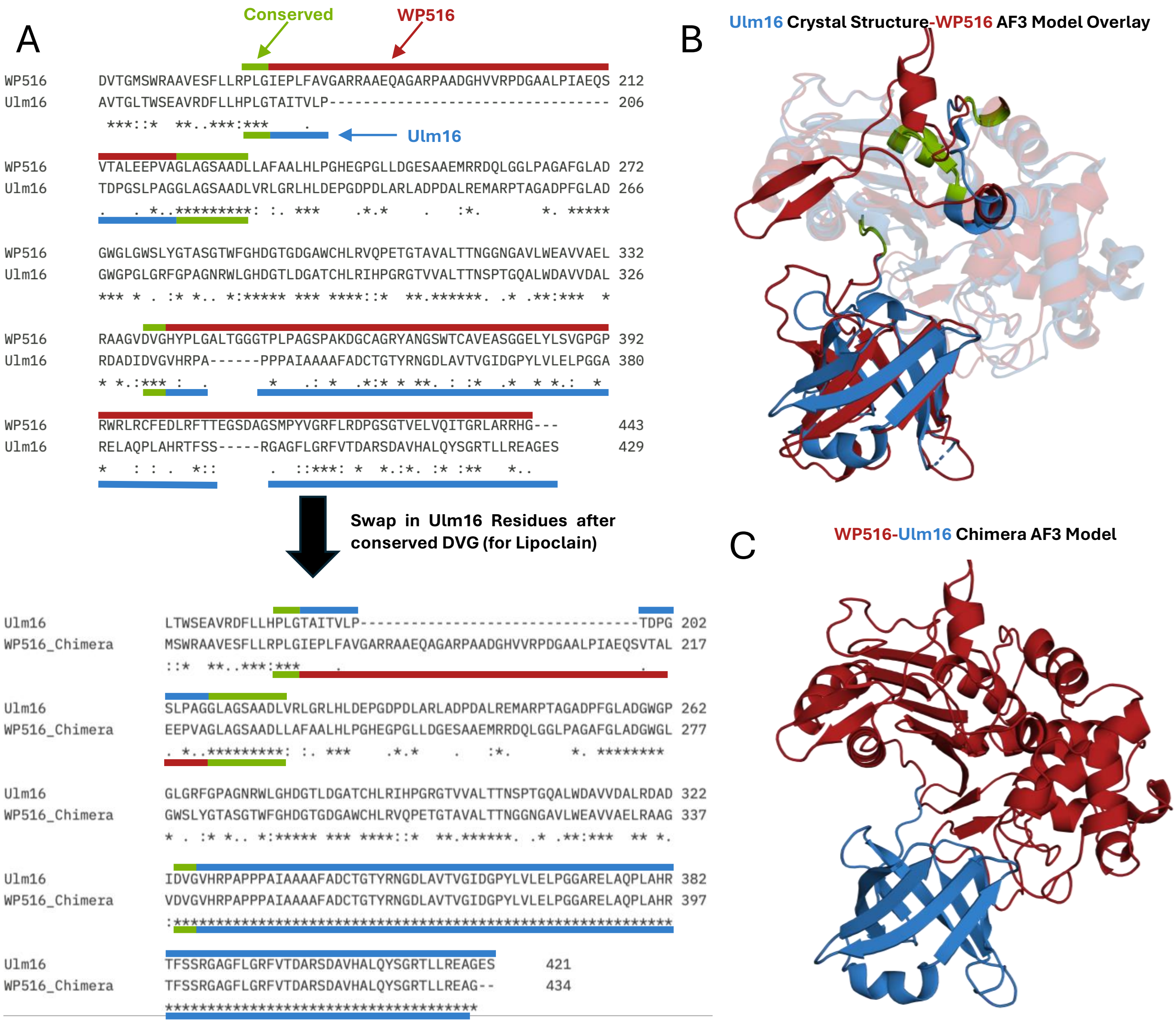

**Figure S74. Making of WP516 Chimera (WP516 C).** A) Partial multiple sequence alignment of WP516, and Ulm16 Chimera, with green representing either conserved regions leading into the active site loop, or lipocalin. Red lines represent amino acids native to WP516, and blue represent Ulm16 amino acids native to Ulm16. The active site loop of WP516 was left intact, however WP516s lipocalin domain was swapped after the DVG motif. B) comparison of WP516 AF3 model (red), and Ulm16 crystal structure (8FEK, blue) with active site loop, and lipocalins highlighted. C) AF3 model of WP516 Chimera with WP516s portions in red, and Ulm16 portions in Blue.

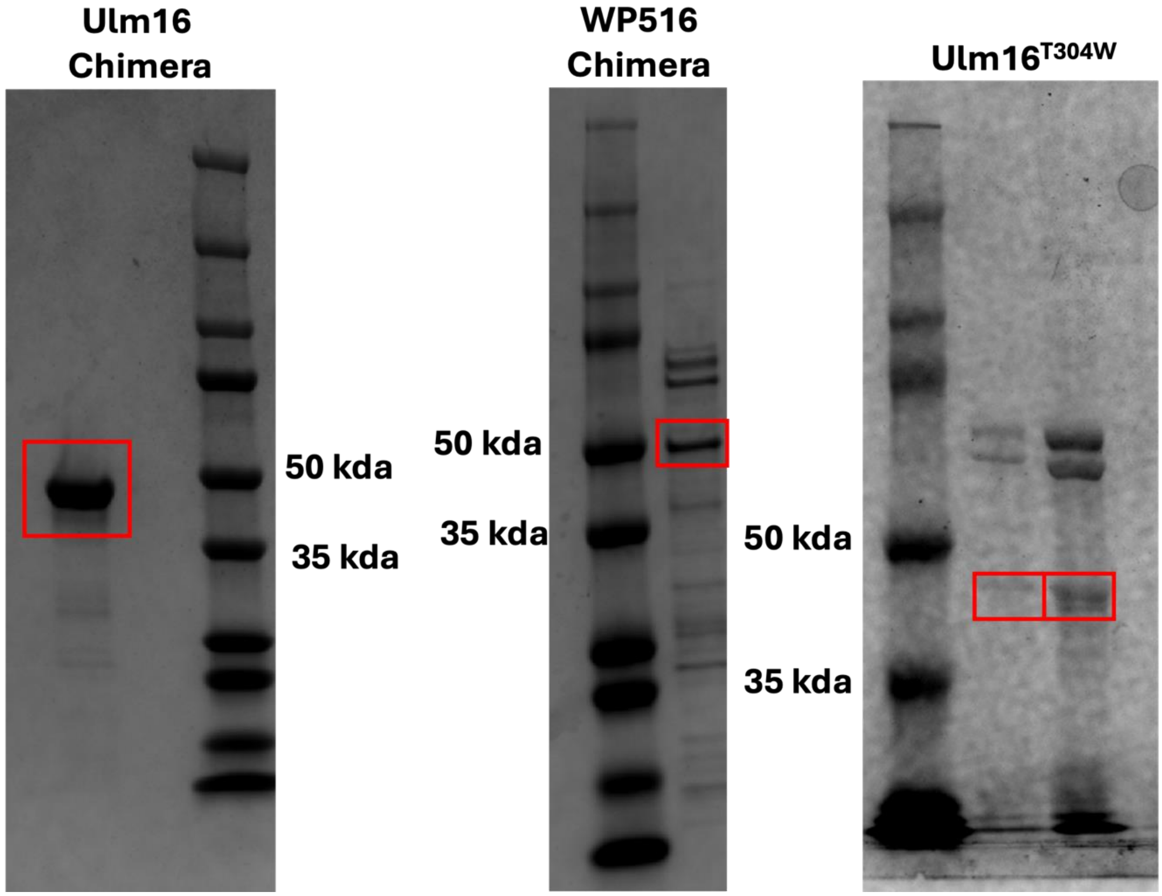

**Figure S75. Expression gels of Ulm16 Chimera, WP516 Chimera, and Ulm16$^{T304W}$.**
Protein at the expected molecular weights are boxed in red.

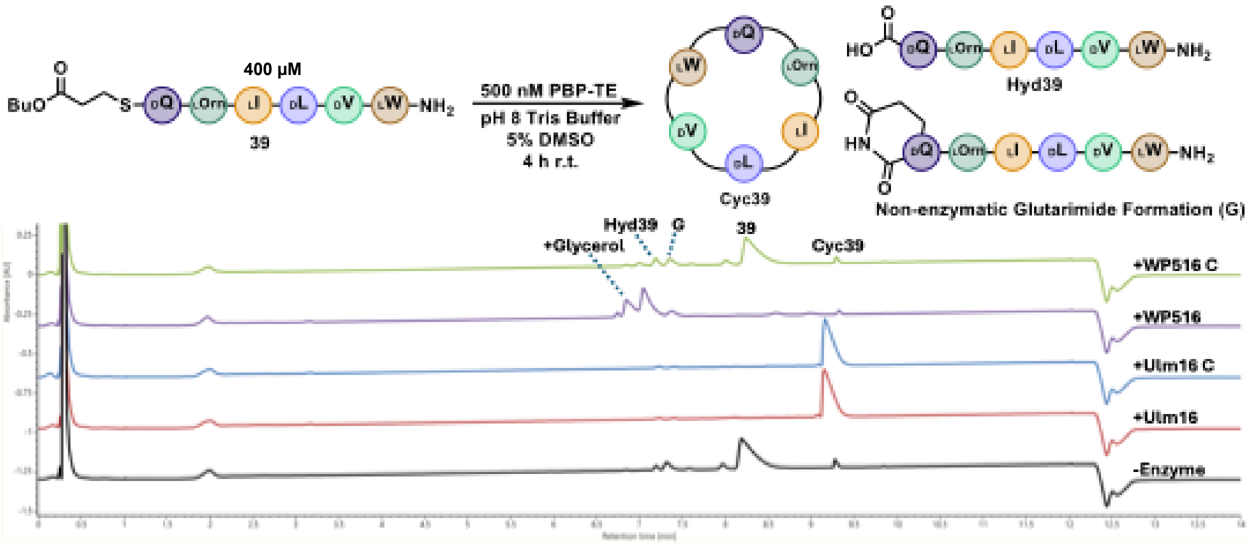

**Figure S76. UPLC UV trace (214 nm) is shown for the assay of WT enzymes and chimeric mutants with peptide substrate 39.** Enzyme and substrate concentration are indicated in the reaction scheme. The substrate is labeled with its corresponding number from the paper where 'Cyc' and 'Hyd' represent head-to-tail cyclic and hydrolyzed peptide, respectively. Non-enzymatic glutarimide formation is indicated by 'G' and the addition of glycerol to the peptide substrate by '+ Glycerol'. The shown UPLC traces are representative of three replicates.

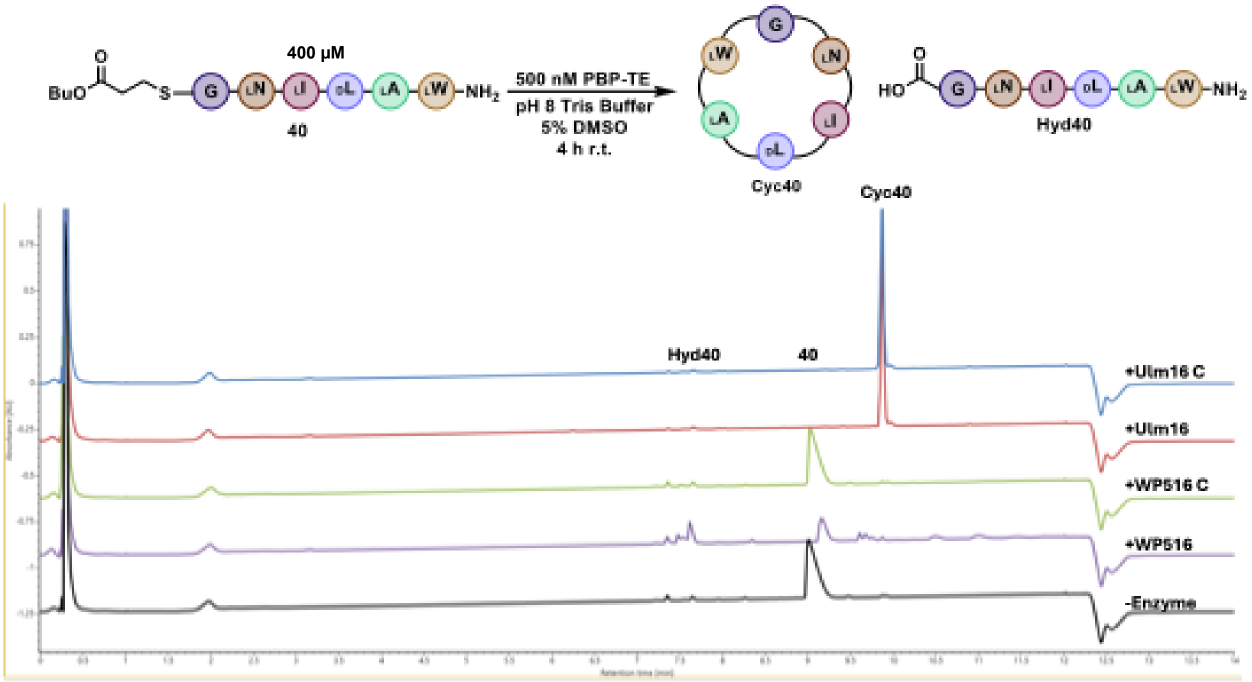

**Figure S77. UPLC UV trace (214 nm) is shown for the assay of WT enzymes and chimeric mutants with peptide substrate 40.** Enzyme and substrate concentration are indicated in the reaction scheme. The substrate is labeled with its corresponding number from the paper where 'Cyc' and 'Hyd' represent head-to-tail cyclic and hydrolyzed peptide, respectively. The shown UPLC traces are representative of three replicates.

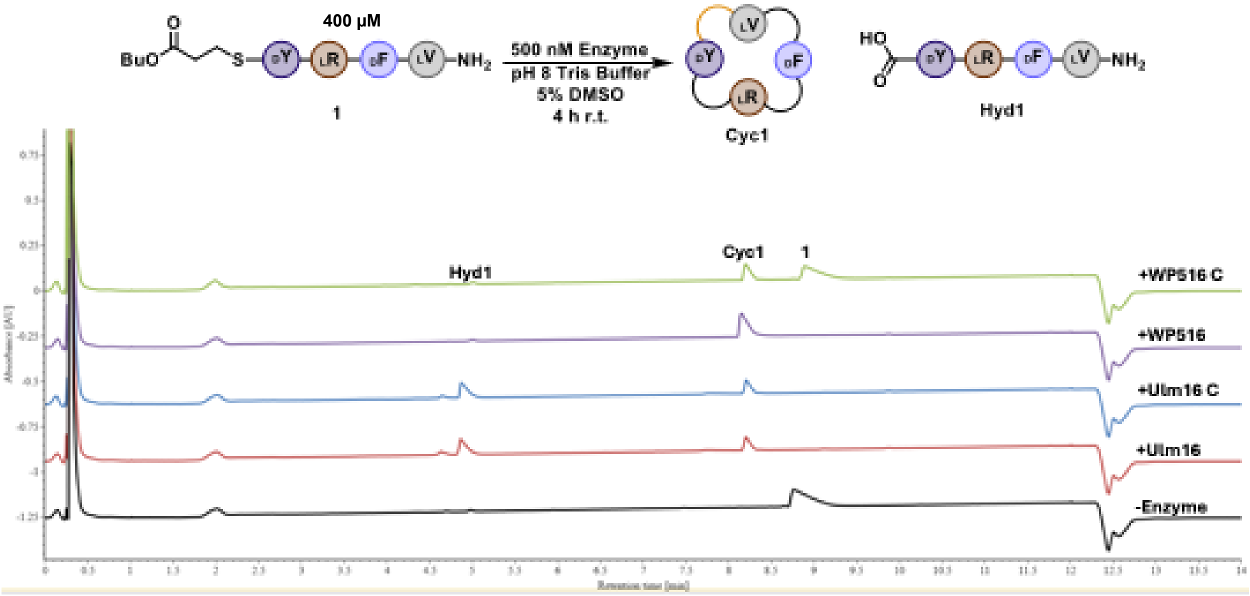

**Figure S78. UPLC UV trace (214 nm) is shown for the assay of WT enzymes and chimeric mutants with peptide substrate 1.** Enzyme and substrate concentration are indicated in the reaction scheme. The substrate is labeled with its corresponding number from the paper where 'Cyc' and 'Hyd' represent head-to-tail cyclic and hydrolyzed peptide, respectively. The shown UPLC traces are representative of three replicates.

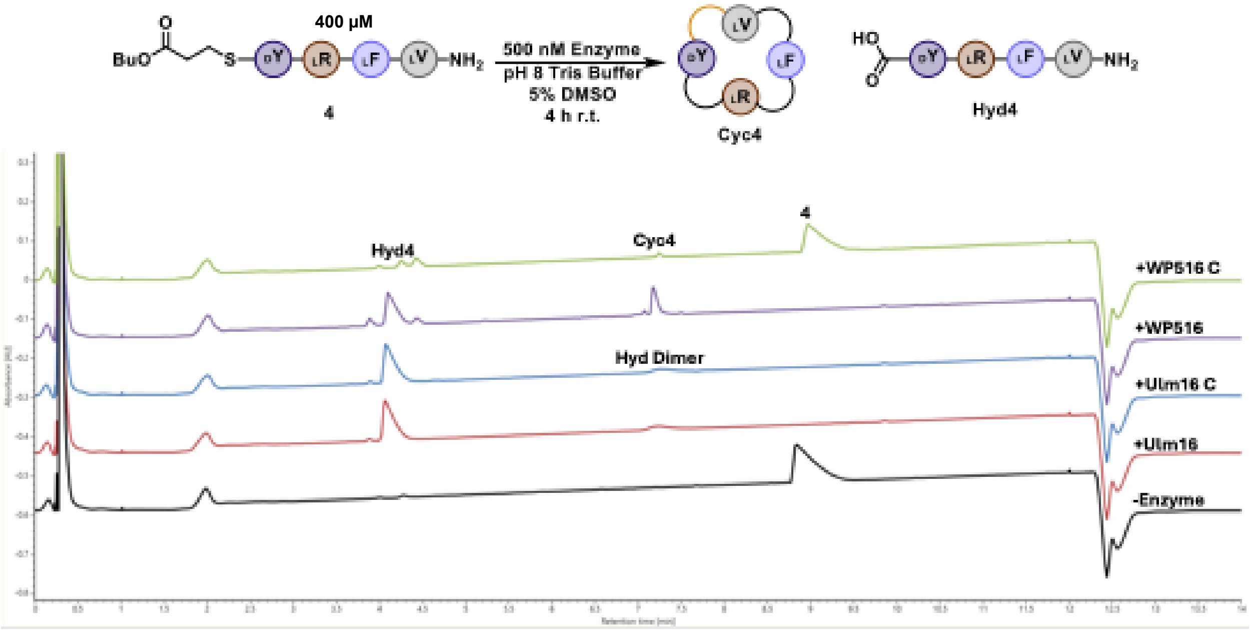

**Figure S79. UPLC UV trace (214 nm) is shown for the assay of WT enzymes and chimeric mutants with peptide substrate 4.** Enzyme and substrate concentration are indicated in the reaction scheme. The substrate is labeled with its corresponding number from the paper where 'Cyc' and 'Hyd' represent head-to-tail cyclic and hydrolyzed peptide, respectively. The shown UPLC traces are representative of three replicates.

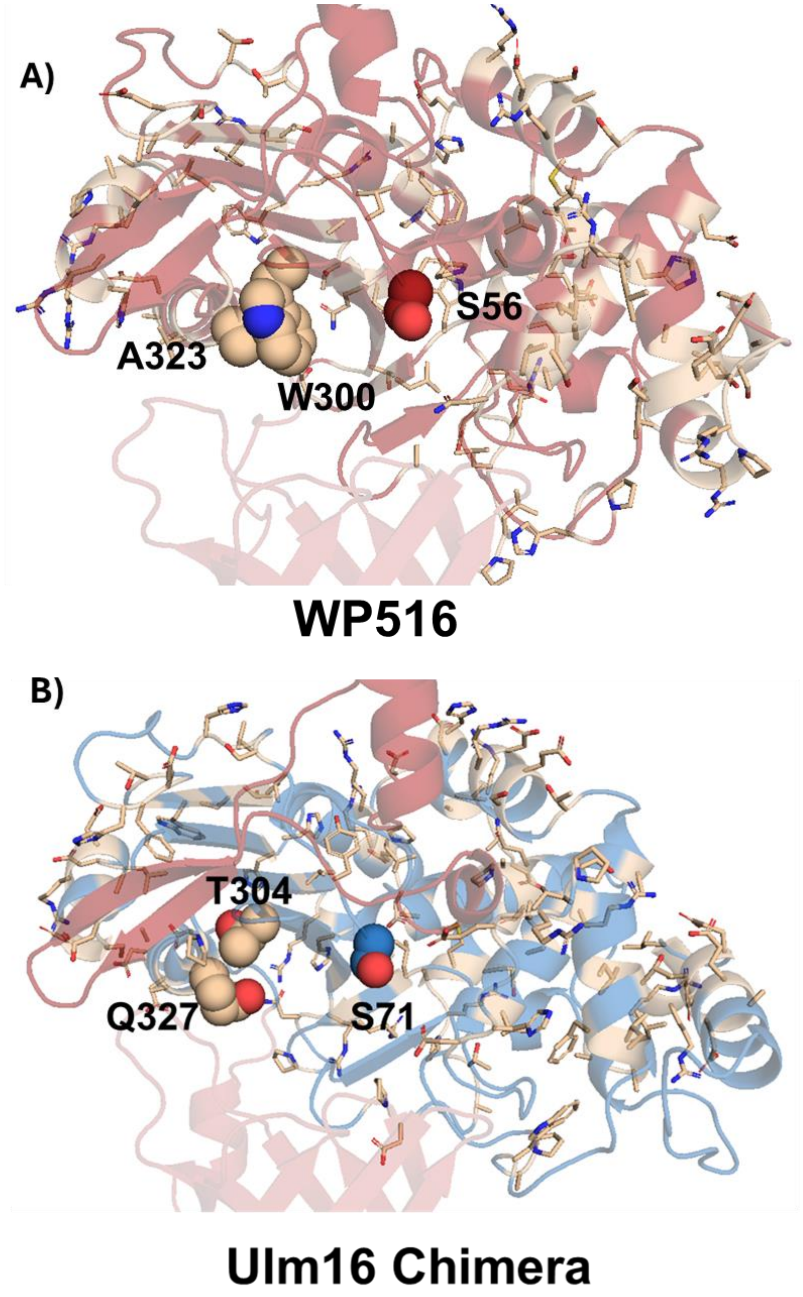

**Figure S80. α/β-hydrolase domain residue analysis.** Residue differences between (A) WP516 and (B) Ulm16 Chimera are highlighted in sphere representation. In WP516, Tryptophan 300 (W300) is substituted with a threonine (T304) in Ulm16. We hypothesized that this change from a bulky to smaller residue reduces Ulm16's ability to cyclize tetrapeptides. The residue directly behind each of these amino acids are also shown in sphere representation. Leucine 300 in Ulm16 was omitted from this figure as it has previously been identified.

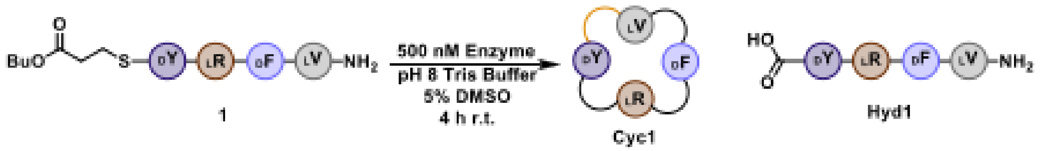

| Enzyme | Peptide | Area Cyclic Peptide | Area Hydrolyzed | Product/Hydrolysis |
| --- | --- | --- | --- | --- |
| 500 nM | 400 uM | 780173 | 211256 | 3.693021737 |

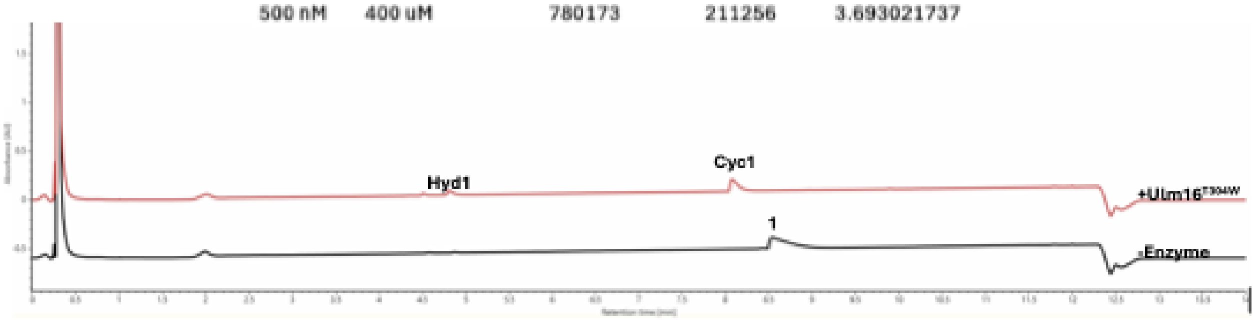

**Figure S81. UPLC UV trace (214 nm) is shown for the assay of Ulm16T304W with peptide substrate 1.** Enzyme and substrate concentration are indicated in the reaction scheme. The substrate is labeled with its corresponding number from the paper where 'Cyc' and 'Hyd' represent head-to-tail cyclic and hydrolyzed peptide, respectively. The shown UPLC traces are representative of three replicates.

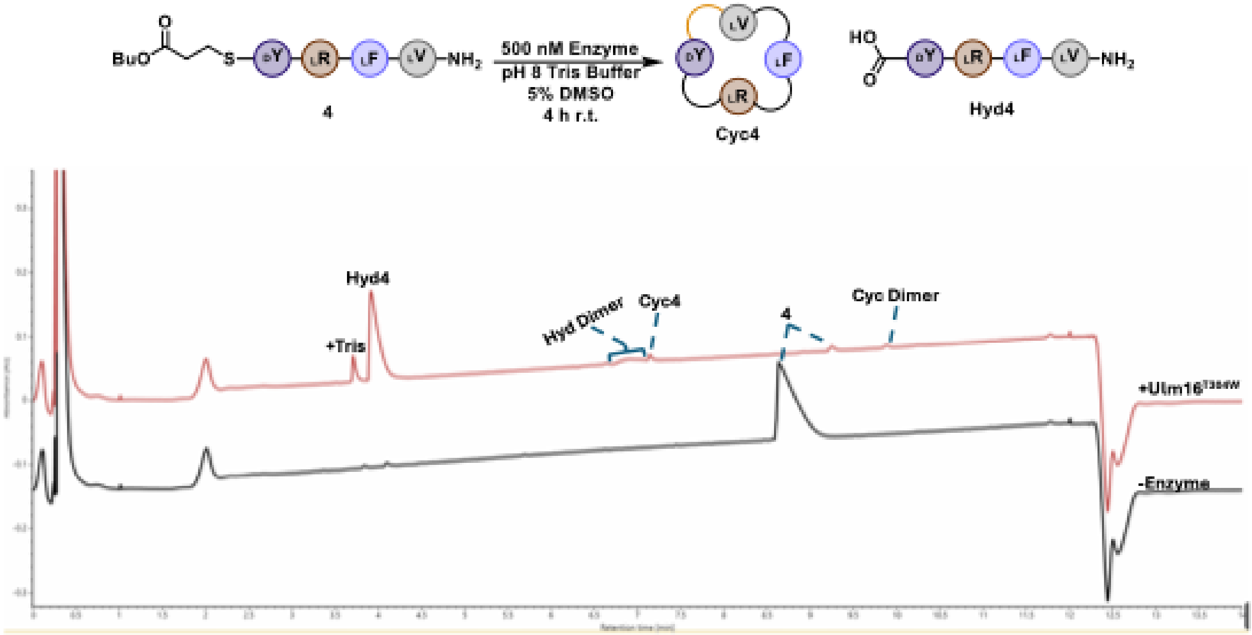

**Figure S82. UPLC UV trace (214 nm) is shown for the assay of Ulm16T304W with peptide substrate 4.** Reaction was run with 500 nM Ulm16T304W, and 400 µM 4. The substrate is labeled with its corresponding number from the paper where 'Cyc' and 'Hyd' represent head-to-tail cyclic and hydrolyzed peptide, respectively. The shown UPLC traces are representative of three replicates.

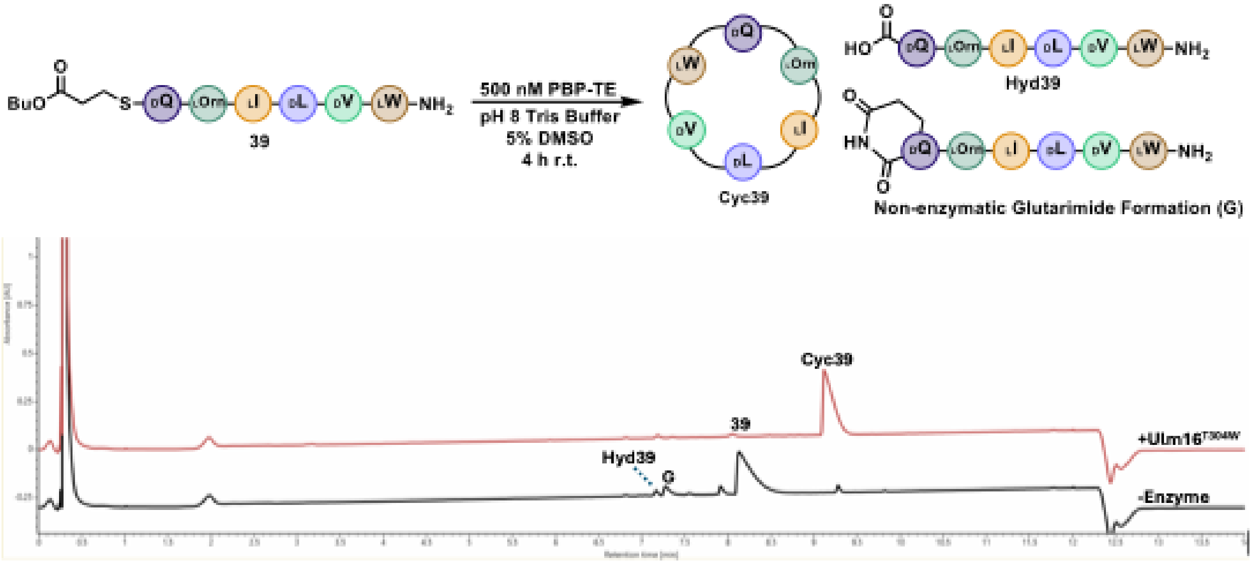

**Figure S83. UPLC UV trace (214 nm) is shown for the assay of Ulm16T304W with peptide substrate 39.** Reaction was run with 500 nM Ulm16T304W, and 400 µM 39. The substrate is labeled with its corresponding number from the paper where ‘Cyc’ and ‘Hyd’ represent head-to-tail cyclic and hydrolyzed peptide, respectively. The shown UPLC traces are representative of three replicates.

-Enzyme

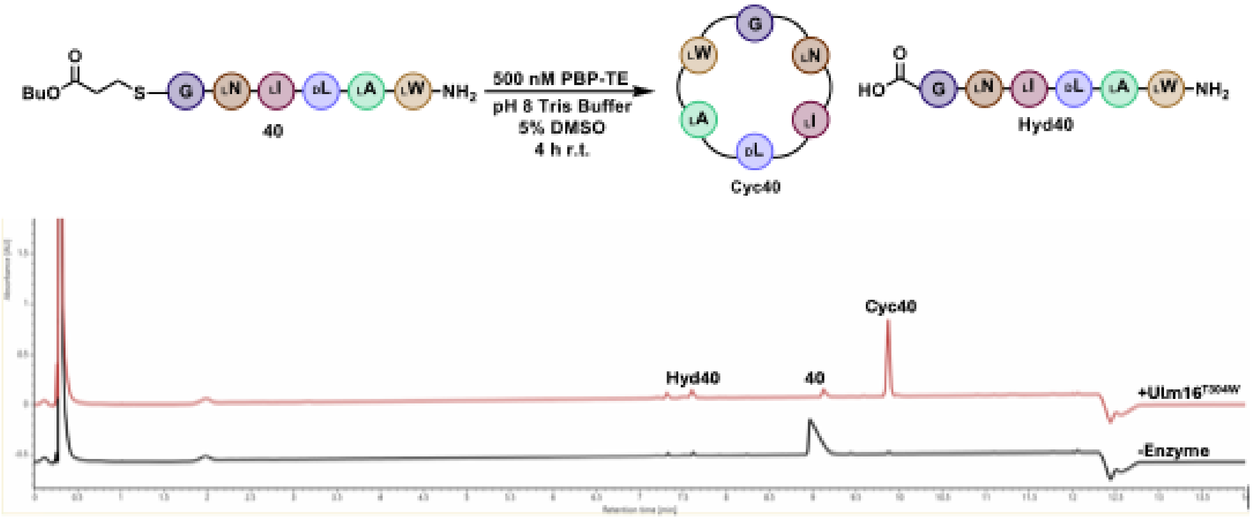

**Figure S84. UPLC UV trace (214 nm) is shown for the assay of Ulm16T304W with peptide substrate 40.** Reaction was run with 500 nM Ulm16T304W, and 400 µM 40. The substrate is labeled with its corresponding number from the paper where ‘Cyc’ and ‘Hyd’ represent head-to-tail cyclic and hydrolyzed peptide, respectively. The shown UPLC traces are representative of three replicates.

### SI Figures S62-S90. MSMS Spectra of Cyclic Peptides Identified in this study.

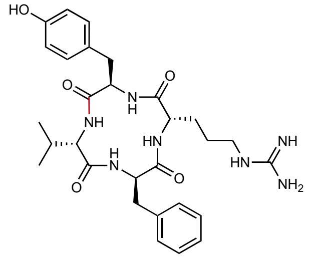

Chemical Formula: $C_{29}H_{39}N_7O_5$
Expected Neutral Mass: 565.3013
Observed Neutral Mass: 565.3016
Mass Error (ppm): 0.5

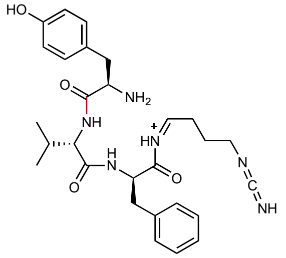

**1**

Chemical Formula: $C_{28}H_{37}N_6O_4^+$
Expected m/z: 521.287
Observed m/z: 521.287
Mass Error (mDa): < 1.0

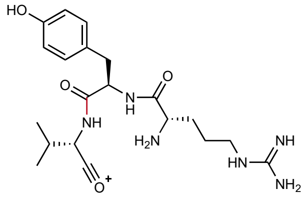

**2**

Chemical Formula: $C_{20}H_{31}N_6O_4$
Expected m/z: 419.240
Observed m/z: 419.245
Mass Error (mDa): 5.0

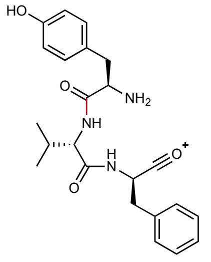

**3**

Chemical Formula: $C_{23}H_{28}N_3O_4^+$
Expected m/z: 410.207
Oberserved m/z: 410.208
Mass Error (mDa): 1.0

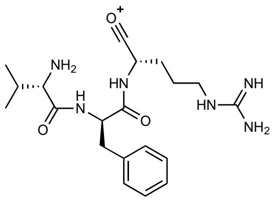

**4**

Chemical Formula: $C_{20}H_{31}N_6O_3^+$
Expected m/z: 403.245
Observed m/z: 403.240
Mass Error (mDa): 5.0

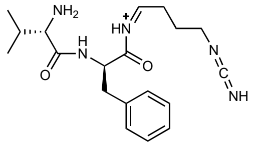

**5**

Chemical Formula: $C_{19}H_{28}N_5O_2^+$
Expected m/z: 358.224
Observed m/z: 358.224
Mass Error (mDa): < 1.0

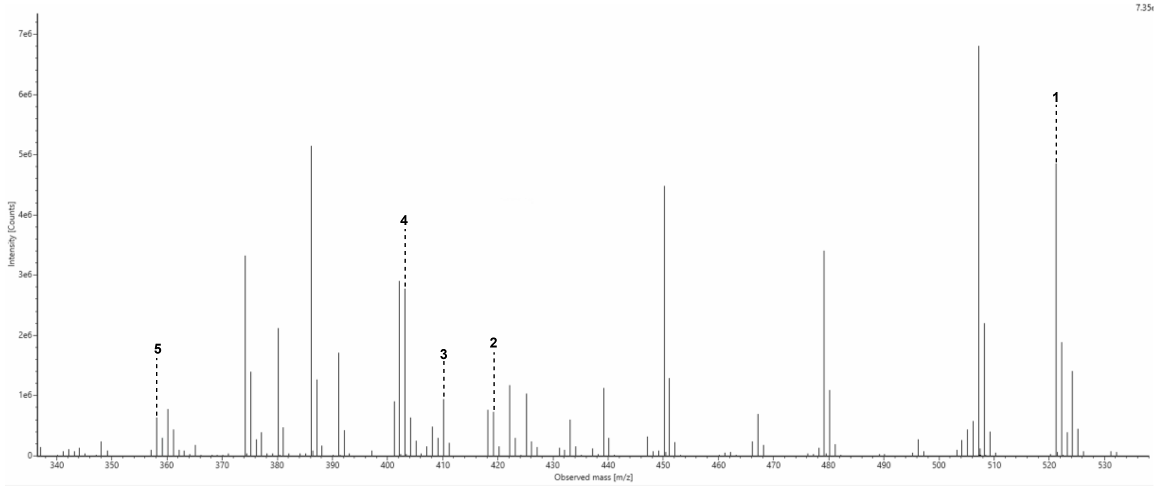

**Figure S85: MS2 spectrum for cyclic peptide 1 (Cyc1).**

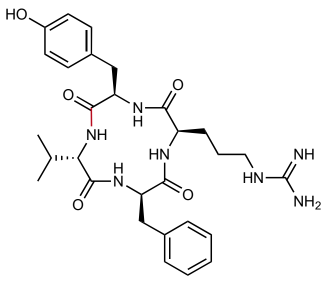

Chemical Formula: $C_{29}H_{39}N_7O_5$
Expected Neutral Mass: 565.3013
Observed Neutral Mass: 565.3027
Mass Error (ppm): 2.5

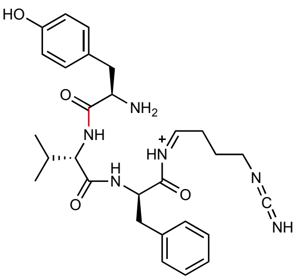

1

Chemical Formula: $C_{28}H_{37}N_6O_4^+$
Expected m/z: 521.287
Observed m/z: 521.287
Mass Error (mDa): < 1.0

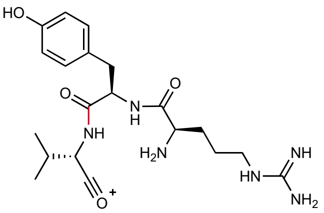

2

Chemical Formula: $C_{20}H_{31}N_6O_4$
Expected m/z: 419.240
Observed m/z: 419.245
Mass Error (mDa): 5.0

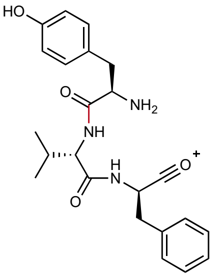

3

Chemical Formula: $C_{23}H_{28}N_3O_4^+$
Expected m/z: 410.207
Oberserved m/z: 410.208
Mass Error (mDa): 1.0

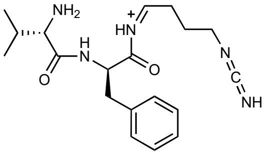

4

Chemical Formula: $C_{19}H_{28}N_5O_2^+$
Expected m/z: 358.224
Observed m/z: 358.223
Mass Error (mDa): < 1.0

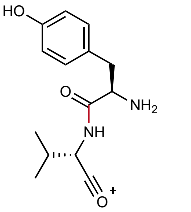

5

Chemical Formula: $C_{14}H_{19}N_2O_3^+$
Expected m/z: 263.139
Observed m/z: 263.141
Mass Error (mDa): 2.0

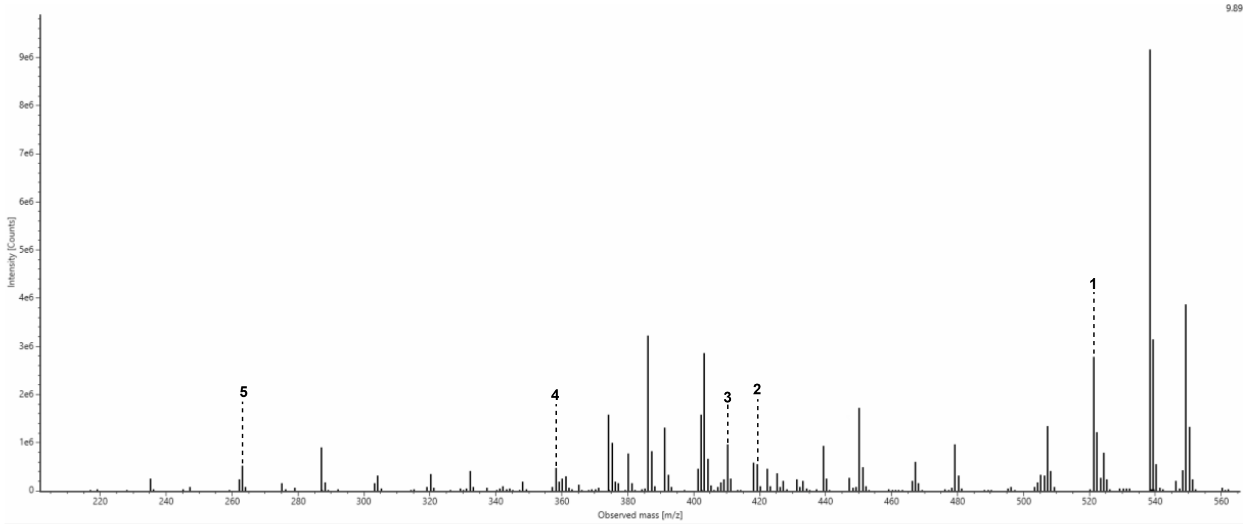

**Figure S86: MS2 spectrum for cyclic peptide 2 (Cyc2).**

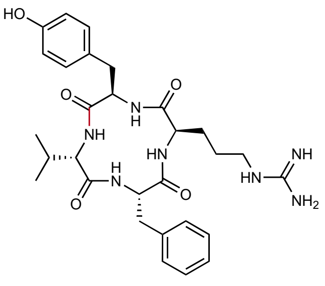

Chemical Formula: $C_{29}H_{39}N_7O_5$
Expected Neutral Mass: 565.3013
Observed Neutral Mass: 565.3023
Mass Error (ppm): 1.8

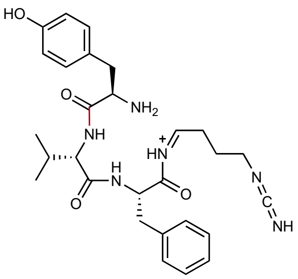

**1**

Chemical Formula: $C_{28}H_{37}N_6O_4^+$
Expected m/z: 521.287
Observed m/z: 521.288
Mass Error (mDa): < 1.0

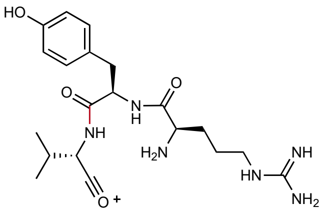

**2**

Chemical Formula: $C_{20}H_{31}N_6O_4$
Expected m/z: 419.240
Observed m/z: 419.246
Mass Error (mDa): 6.0

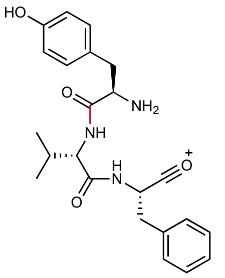

**3**

Chemical Formula: $C_{23}H_{28}N_3O_4^+$
Expected m/z: 410.207
Oberserved m/z: 410.208
Mass Error (mDa): < 1.0

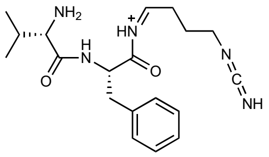

**4**

Chemical Formula: $C_{19}H_{28}N_5O_2^+$
Expected m/z: 358.224
Observed m/z: 358.223
Mass Error (mDa): < 1.0

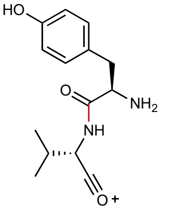

**5**

Chemical Formula: $C_{14}H_{19}N_2O_3^+$
Expected m/z: 263.139
Observed m/z: 263.142
Mass Error (mDa): 3.0

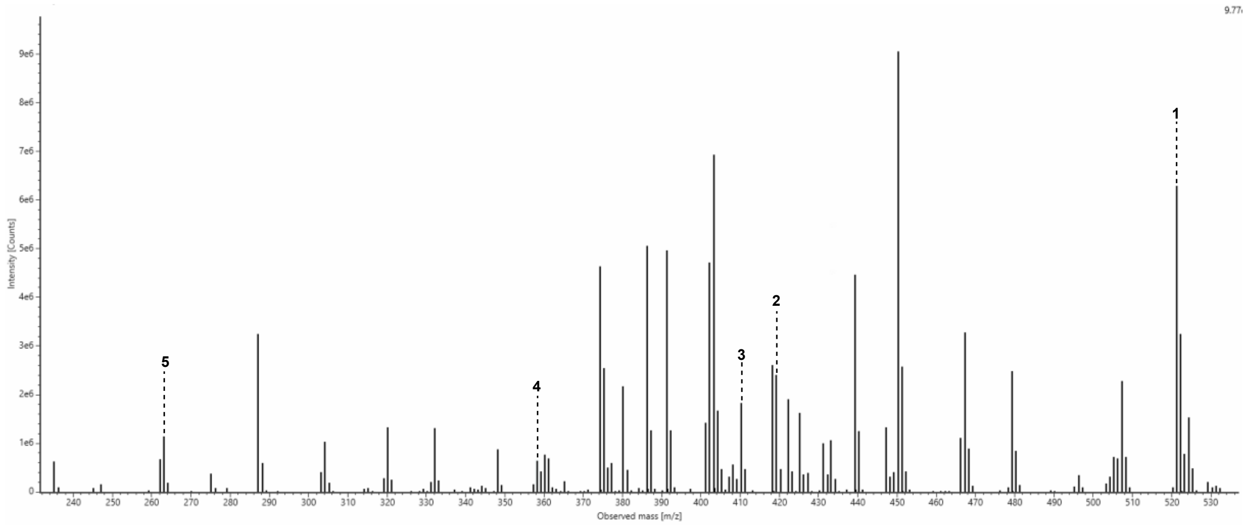

**Figure S87: MS2 spectrum for cyclic peptide 3 (Cyc3).**

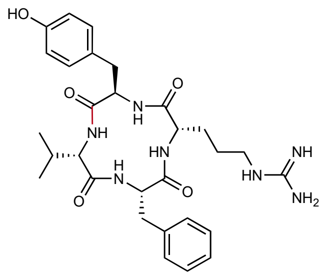

Chemical Formula: $C_{29}H_{39}N_7O_5$
Expected Neutral Mass: 565.3013
Observed Neutral Mass: 565.3027
Mass Error (ppm): 2.6

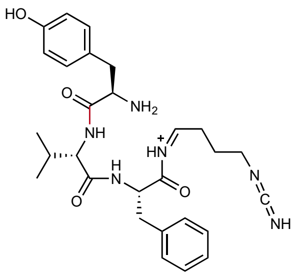

1

Chemical Formula: $C_{28}H_{37}N_6O_4^+$
Expected m/z: 521.287
Observed m/z: 521.289
Mass Error (mDa): 2.0

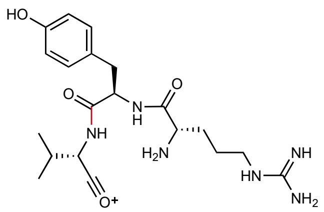

2

Chemical Formula: $C_{20}H_{31}N_6O_4$
Expected m/z: 419.240
Observed m/z: 419.247
Mass Error (mDa): 7.0

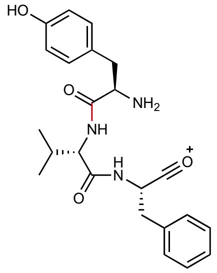

3

Chemical Formula: $C_{23}H_{28}N_3O_4^+$
Expected m/z: 410.207
Oberserved m/z: 410.209
Mass Error (mDa): 1.0

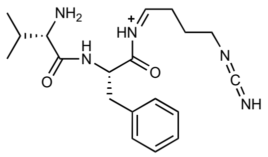

4

Chemical Formula: $C_{19}H_{28}N_5O_2^+$
Expected m/z: 358.224
Observed m/z: 358.226
Mass Error (mDa): 2.0

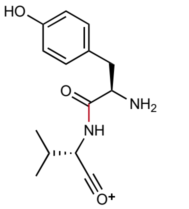

5

Chemical Formula: $C_{14}H_{19}N_2O_3^+$
Expected m/z: 263.139
Observed m/z: 263.143
Mass Error (mDa): 4.0

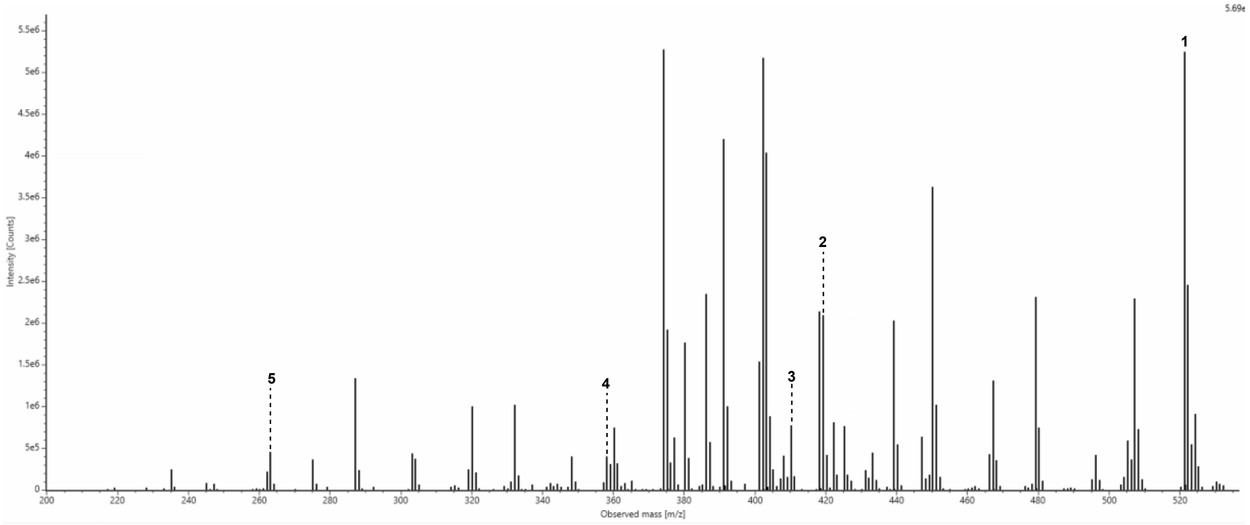

**Figure S88: MS2 spectrum for cyclic peptide 4 (Cyc4).**

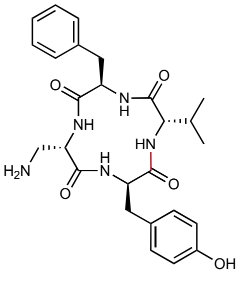

Chemical Formula: $C_{26}H_{33}N_5O_5$
Expected Neutral Mass: 495.2482
Observed Neutral Mass: 495.2478
Mass Error (ppm): 0.7

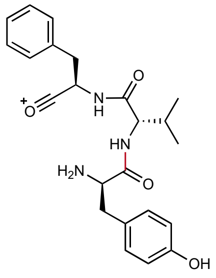

**1**
Chemical Formula: $C_{23}H_{28}N_3O_4^+$
Expected m/z: 410.207
Observed m/z: 410.208
Mass Error (mDa): < 1.0

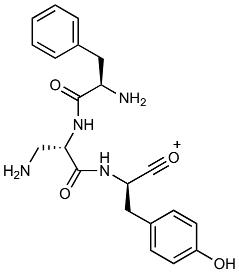

**2**
Chemical Formula: $C_{21}H_{25}N_4O_4^+$
Expected m/z: 397.187
Observed m/z: 397.188
Mass Error (mDa): 1.0

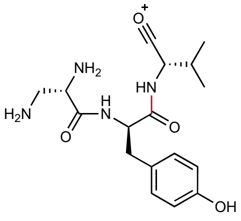

**3**
Chemical Formula: $C_{17}H_{25}N_4O_4^+$
Expected m/z: 349.187
Observed m/z: 349.189
Mass Error (mDa): 2.0

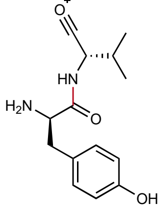

**4**
Chemical Formula: $C_{14}H_{19}N_2O_3^+$
Expected m/z: 263.139
Observed m/z: 263.141
Mass Error (mDa): 2.0

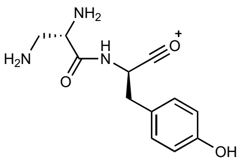

**5**
Chemical Formula: $C_{12}H_{16}N_3O_3^+$
Expected m/z: 250.119
Observed m/z: 250.121
Mass Error (mDa): 2.0

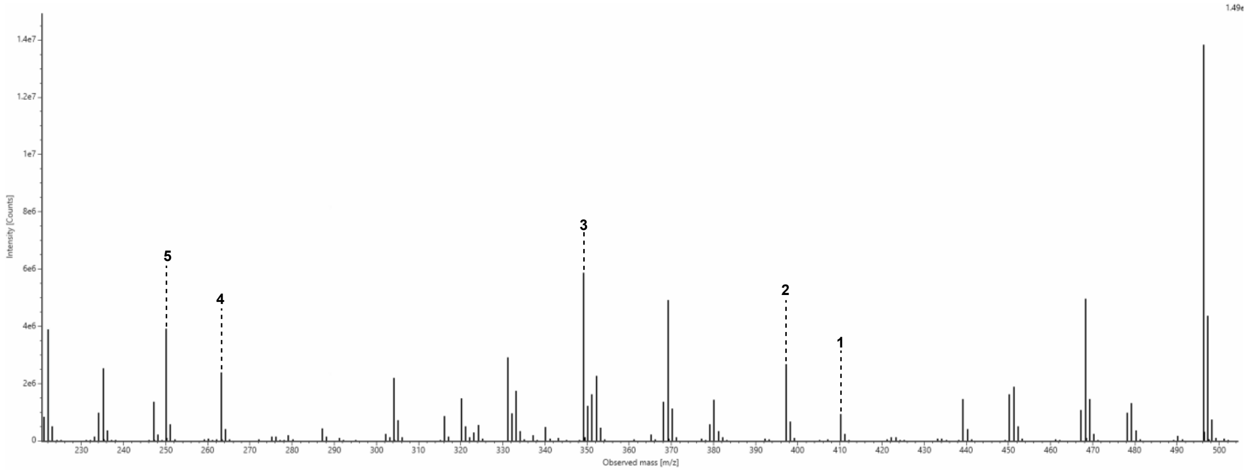

**Figure S89: MS2 spectrum for cyclic peptide 5 (Cyc5).**

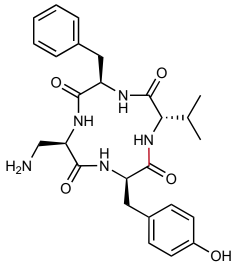

Chemical Formula: $C_{26}H_{33}N_5O_5$
Expected Neutral Mass: 495.2482
Observed Neutral Mass: 495.2558
Mass Error (ppm): 0.8

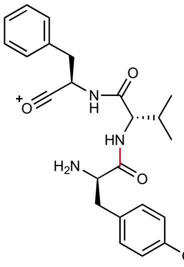

**1**
Chemical Formula: $C_{23}H_{28}N_3O_4^+$
Expected m/z: 410.207
Observed m/z: 410.208
Mass Error (mDa): 1.0

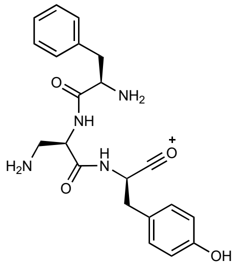

**2**
Chemical Formula: $C_{21}H_{25}N_4O_4^+$
Expected m/z: 397.187
Observed m/z: 397.189
Mass Error (mDa): 2.0

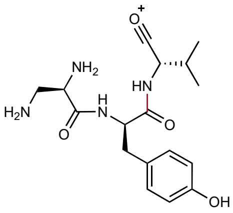

**3**
Chemical Formula: $C_{17}H_{25}N_4O_4^+$
Expected m/z: 349.187
Observed m/z: 349.189
Mass Error (mDa): 2.0

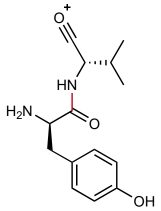

**4**
Chemical Formula: $C_{14}H_{19}N_2O_3^+$
Expected m/z: 263.139
Observed m/z: 263.141
Mass Error (mDa): 2.0

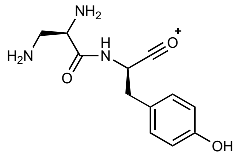

**5**
Chemical Formula: $C_{12}H_{16}N_3O_3^+$
Expected m/z: 250.119
Observed m/z: 250.121
Mass Error (mDa): 2.0

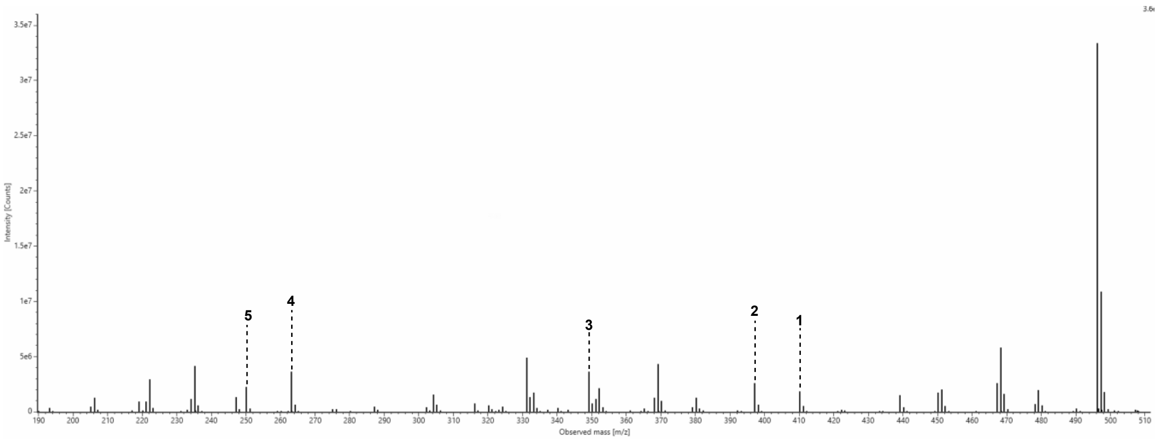

**Figure S90: MS2 spectrum for cyclic peptide 6 (Cyc6).**

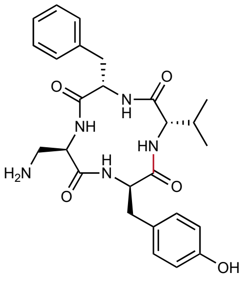

Chemical Formula: $C_{26}H_{33}N_5O_5$
Expected Neutral Mass: 495.2482
Observed Neutral Mass: 495.2482
Mass Error (ppm): 0.1

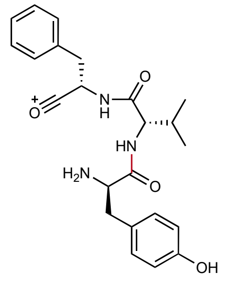

**1**
Chemical Formula: $C_{23}H_{28}N_3O_4^+$
Expected m/z: 410.207
Observed m/z: 410.208
Mass Error (mDa): 1.0

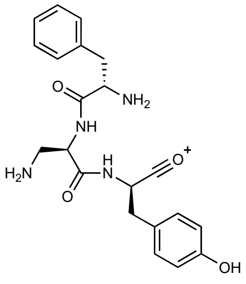

**2**
Chemical Formula: $C_{21}H_{25}N_4O_4^+$
Expected m/z: 397.187
Observed m/z: 397.188
Mass Error (mDa): 1.0

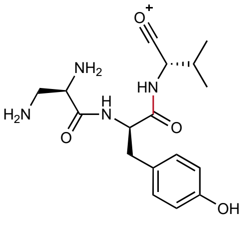

**3**
Chemical Formula: $C_{17}H_{25}N_4O_4^+$
Expected m/z: 349.187
Observed m/z: 349.188
Mass Error (mDa): 1.0

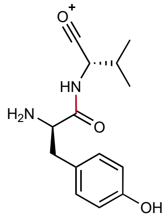

**4**
Chemical Formula: $C_{14}H_{19}N_2O_3^+$
Expected m/z: 263.139
Observed m/z: 263.141
Mass Error (mDa): 2.0

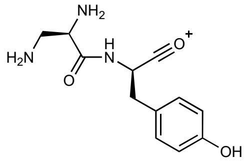

**5**
Chemical Formula: $C_{12}H_{16}N_3O_3^+$
Expected m/z: 250.119
Observed m/z: 250.121
Mass Error (mDa): 2.0

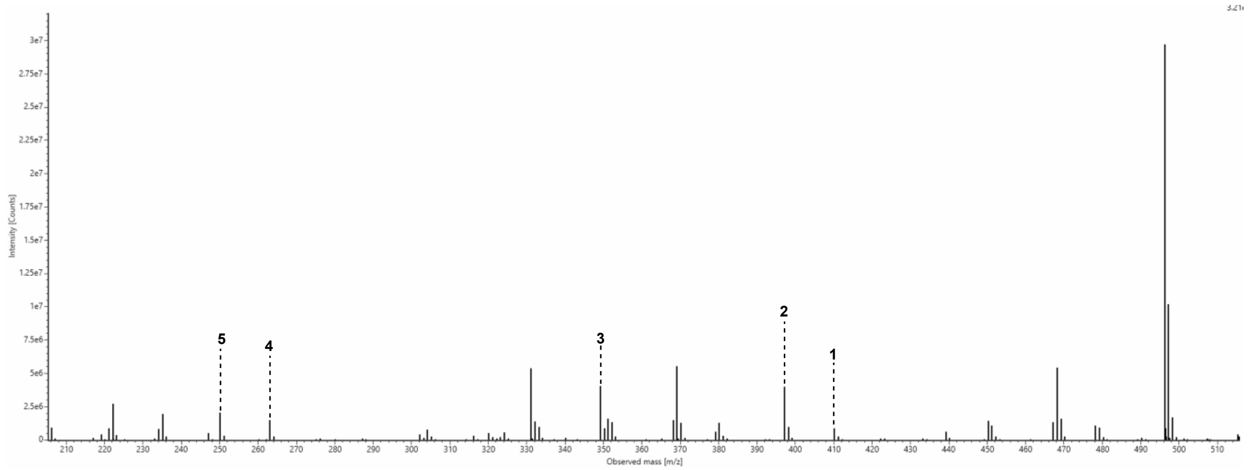

**Figure S91: MS2 spectrum for cyclic peptide 7 (Cyc7).**

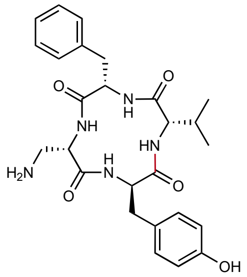

Chemical Formula: $C_{26}H_{33}N_5O_5$
Expected Neutral Mass: 495.2482
Observed Neutral Mass: 495.2488
Mass Error (ppm): 1.2

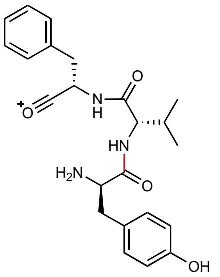

**1**
Chemical Formula: $C_{23}H_{28}N_3O_4^+$
Expected m/z: 410.207
Observed m/z: 410.208
Mass Error (mDa): < 1.0

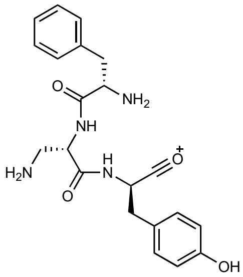

**2**
Chemical Formula: $C_{21}H_{25}N_4O_4^+$
Expected m/z: 397.187
Observed m/z: 397.188
Mass Error (mDa): 1.0

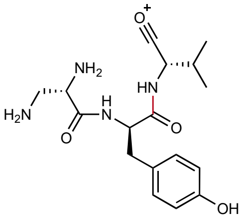

**3**
Chemical Formula: $C_{17}H_{25}N_4O_4^+$
Expected m/z: 349.187
Observed m/z: 349.188
Mass Error (mDa): 1.0

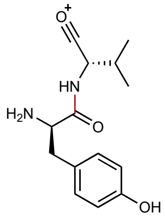

**4**
Chemical Formula: $C_{14}H_{19}N_2O_3^+$
Expected m/z: 263.139
Observed m/z: 263.140
Mass Error (mDa): 1.0

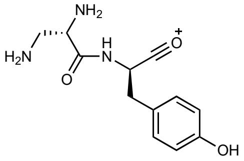

**5**
Chemical Formula: $C_{12}H_{16}N_3O_3^+$
Expected m/z: 250.119
Observed m/z: 250.120
Mass Error (mDa): 1.0

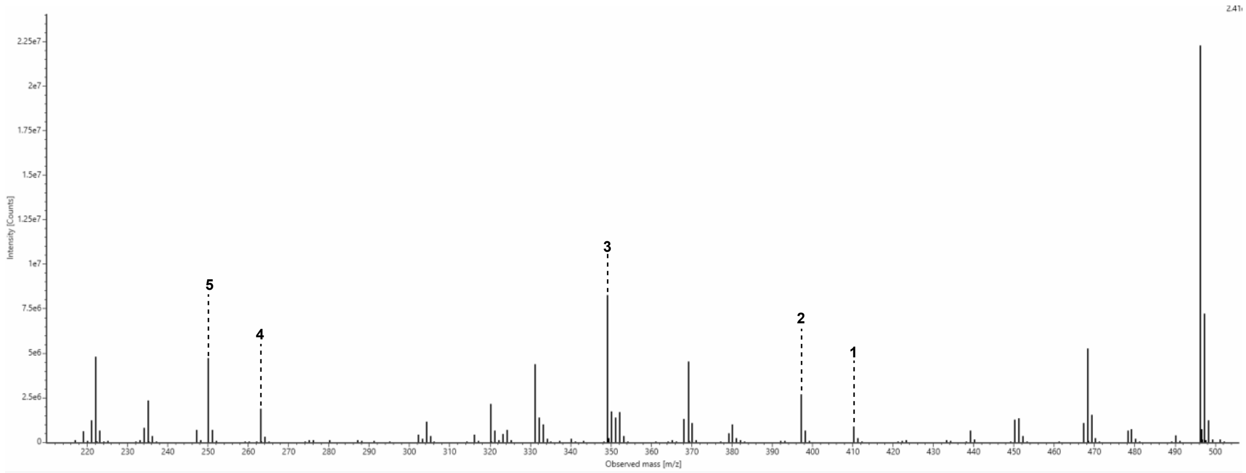

**Figure S92: MS2 spectrum for cyclic peptide 8 (Cyc8).**

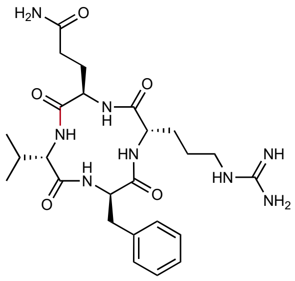

Chemical Formula: $C_{25}H_{38}N_8O_5$
Expected Neutral Mass: 530.2965
Observed Neutral Mass: 530.2965
Mass Error (ppm): < 0.1

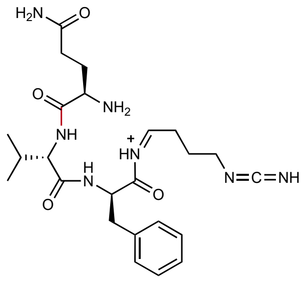

**1**
Chemical Formula: $C_{24}H_{36}N_7O_4^+$
Expected m/z: 486.282
Observe m/z: 486.283
Mass Error (mDa): 1.0

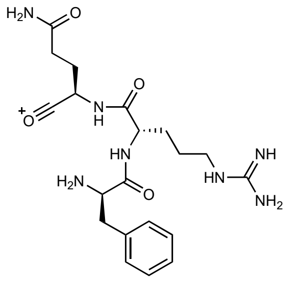

**2**
Chemical Formula: $C_{20}H_{30}N_7O_4^+$
Expected m/z: 432.235
Observed m/z: 432.237
Mass Error (mDa): 2.0

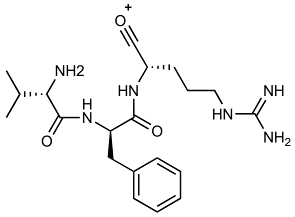

**3**
Chemical Formula: $C_{20}H_{31}N_6O_3^+$
Expected m/z: 403.245
Observed m/z: 403.246
Mass Error (mDa): 1.0

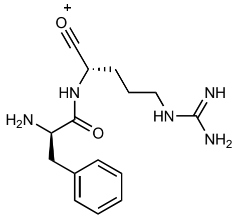

**4**
Chemical Formula: $C_{15}H_{22}N_5O_2^+$
Expected m/z: 304.177
Observed m/z: 304.179
Mass Error (mDa): 2.0

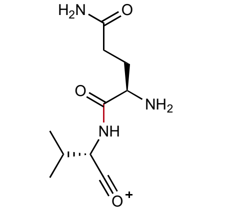

**5**
Chemical Formula: $C_{10}H_{18}N_3O_3$
Expected m/z: 228.135
Observed m/z: 228.132
Mass Error (mDa): 3.0

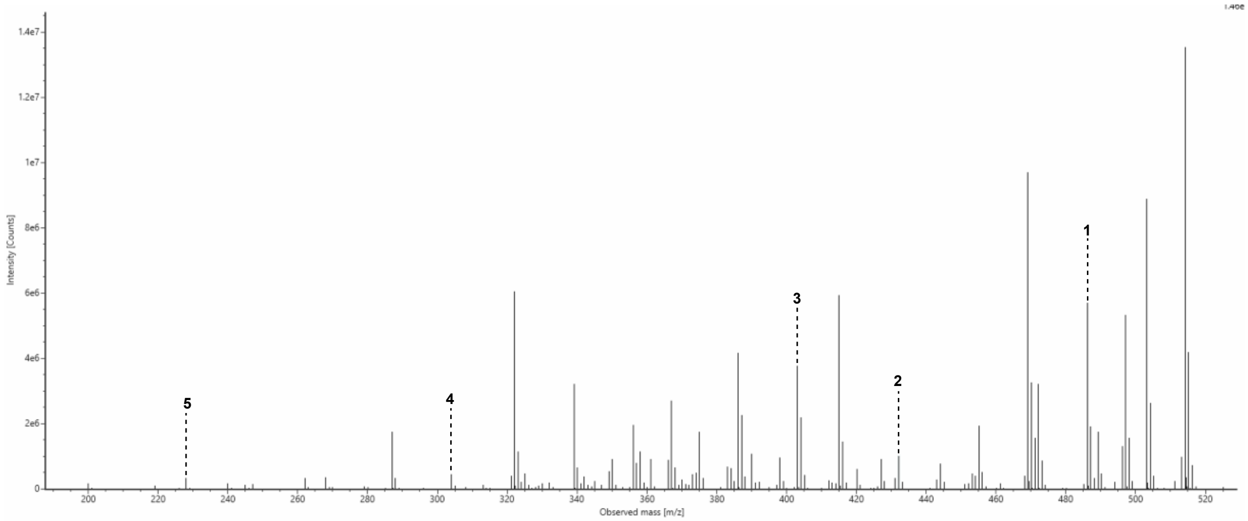

**Figure S93: MS2 spectrum for cyclic peptide 9 (Cyc9).**

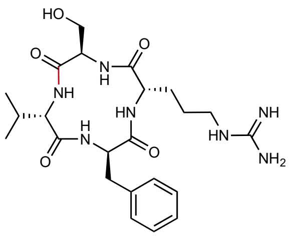

Chemical Formula: $C_{23}H_{35}N_7O_5$
Expected Neutral Mass: 489.2700
Observed Neutral Mass: 489.2703
Mass Error (ppm): 0.6

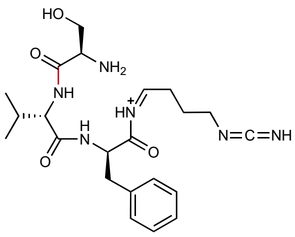

**1**
Chemical Formula: $C_{22}H_{33}N_6O_4^+$
Expected m/z: 445.256
Observed m/z: 445.257
Mass Error (mDa): 1.0

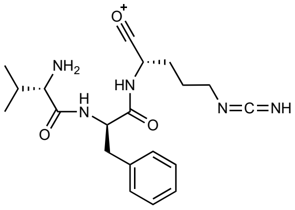

**2**
Chemical Formula: $C_{20}H_{28}N_5O_3^+$
Expected m/z: 386.219
Observed m/z: 386.219
Mass Error (mDa): < 1.0

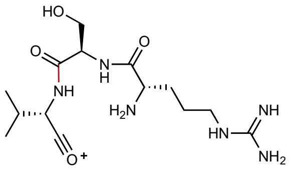

**3**
Chemical Formula: $C_{14}H_{27}N_6O_4^+$
Expected m/z: 343.209
Observed m/z: 343.213
Mass Error (mDa): 4.0

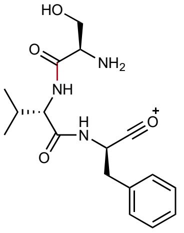

**4**
Chemical Formula: $C_{17}H_{24}N_3O_4^+$
Expected m/z: 334.176
Observed m/z: 334.179
Mass Error (mDa): 3.0

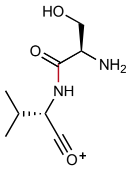

**5**
Chemical Formula: $C_8H_{15}N_2O_3^+$
Expected m/z: 187.108
Observed m/z: 187.108
Mass Error (mDa): < 1.0

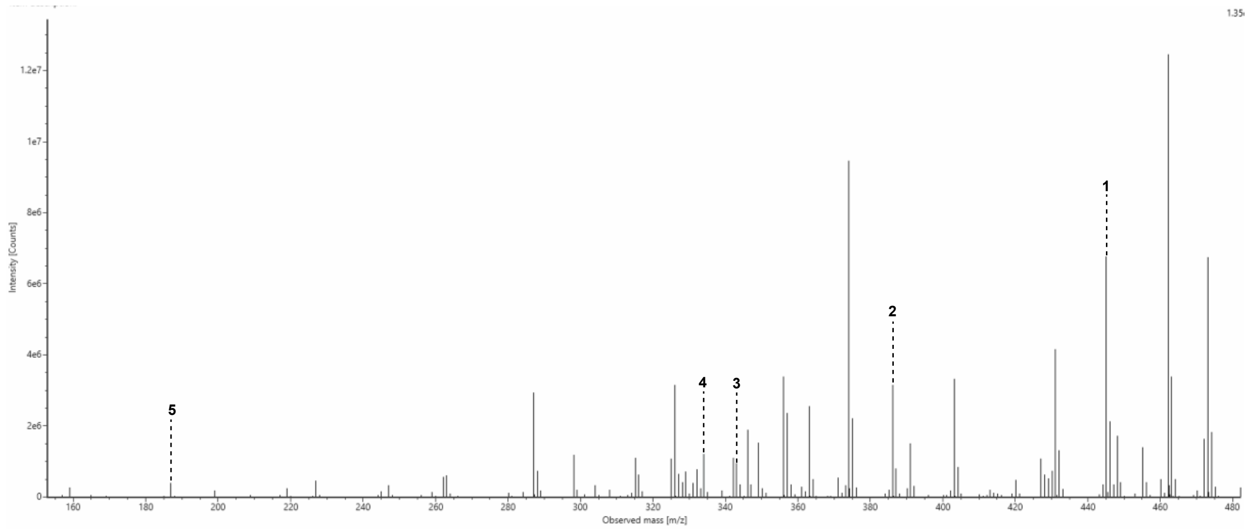

**Figure S94: MS2 spectrum for cyclic peptide 10 (Cyc10).**

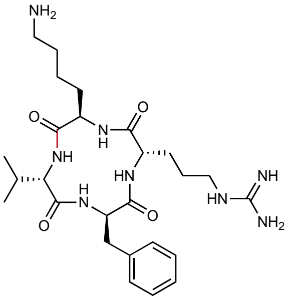

Chemical Formula: $C_{26}H_{42}N_8O_4$
Expected Neutral Mass: 530.3329
Observed Neutral Mass: 530.3336
Mass Error (ppm): 1.3

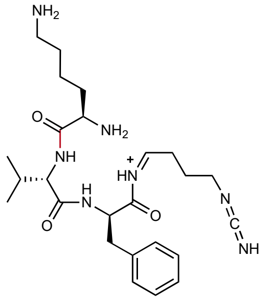

**1**
Chemical Formula: $C_{25}H_{40}N_7O_3^+$
Expected m/z: 486.319
Observed m/z: 486.319
Mass Error (mDa): < 1.0

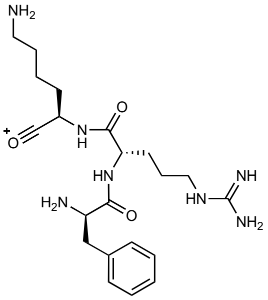

**2**
Chemical Formula: $C_{21}H_{34}N_7O_3^+$
Expected m/z: 432.272
Observed m/z: 432.272
Mass Error (mDa): < 1.0

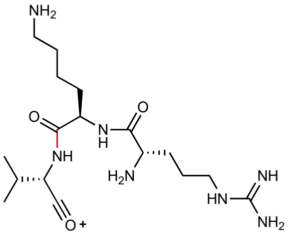

**3**
Chemical Formula: $C_{17}H_{34}N_7O_3^+$
Expected m/z: 384.272
Observed m/z: 384.277
Mass Error (mDa): 6.0

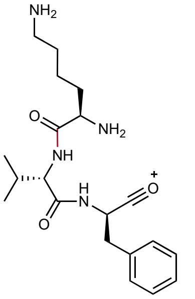

**4**
Chemical Formula: $C_{20}H_{31}N_4O_3^+$
Expected m/z: 375.239
Observed m/z: 375.245
Mass Error (mDa): 6.0

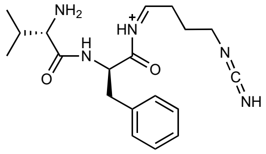

**5**
Chemical Formula: $C_{19}H_{28}N_5O_2^+$
Expected m/z: 358.224
Observed m/z: 358.224
Mass Error (mDa): < 1.0

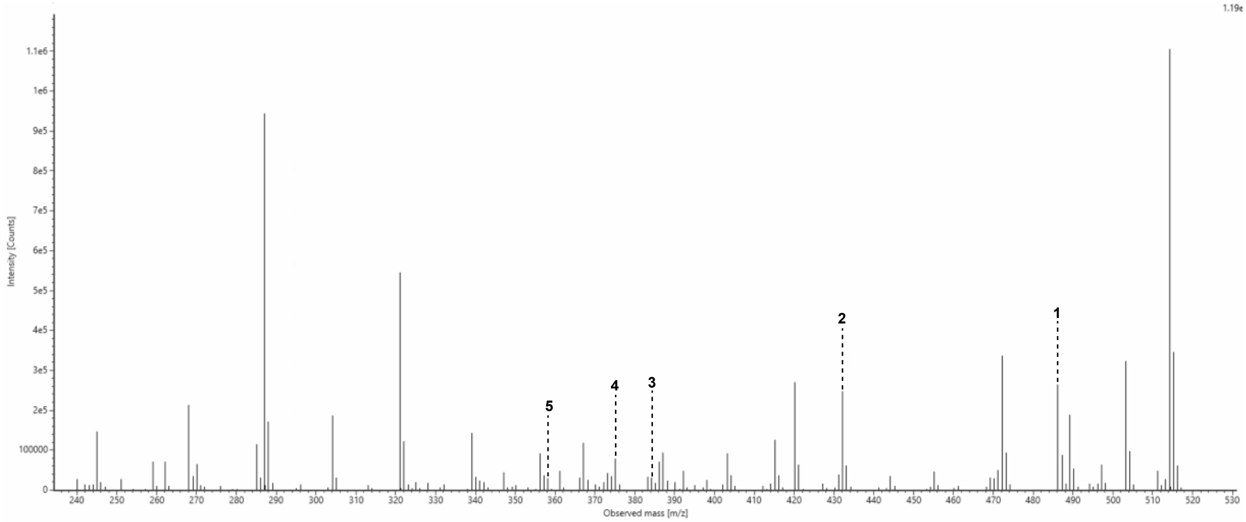

**Figure S94: MS2 spectrum for cyclic peptide 11 (Cyc11).**

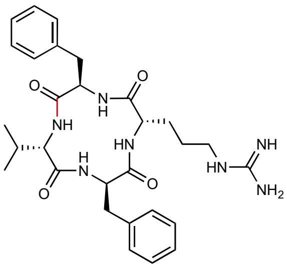

Chemical Formula: $C_{29}H_{39}N_7O_4$
Expected Neutral Mass: 549.3064
Observed Neutral Mass: 549.3068
Mass Error (ppm): 0.8

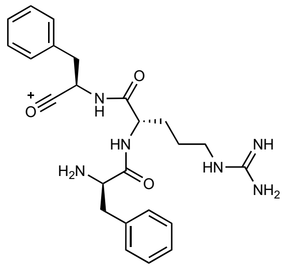

**1**
Chemical Formula: $C_{24}H_{31}N_6O_3$
Expected m/z: 451.246
Observed m/z: 451.247
Mass Error (mDa): 2.0

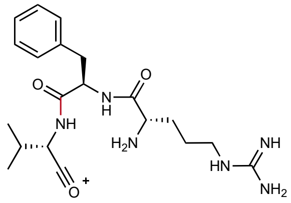

**2**
Chemical Formula: $C_{20}H_{31}N_6O_3^+$
Expected m/z: 403.245
Observed m/z: 403.247
Mass Error (mDa): 2.0

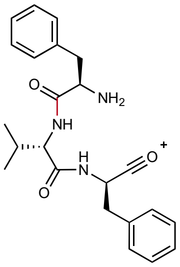

**3**
Chemical Formula: $C_{23}H_{28}N_3O_3^+$
Expected m/z: 394.213
Observed m/z: 394.213
Mass Error (mDa): < 1.0

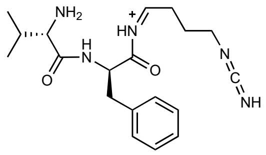

**4**
Chemical Formula: $C_{19}H_{28}N_5O_2^+$
Expected m/z: 358.224
Observed m/z: 358.224
Mass Error (mDa): < 1.0

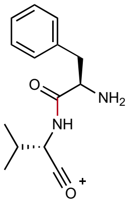

**5**
Chemical Formula: $C_{14}H_{19}N_2O_2^+$
Expected m/z: 247.144
Observed m/z: 247.145
Mass Error (mDa): 1.0

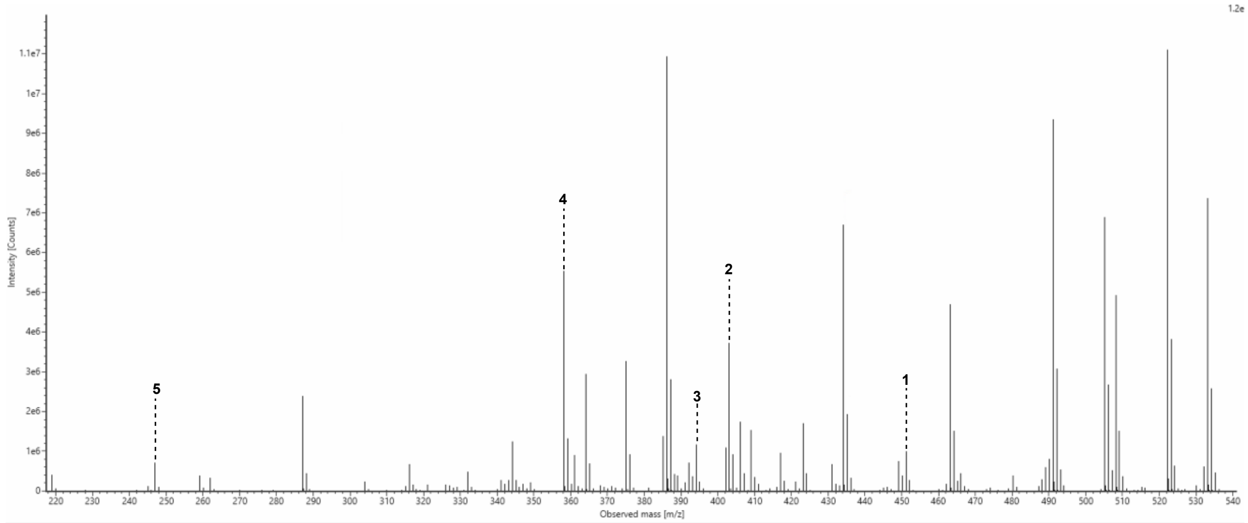

**Figure S95: MS2 spectrum for cyclic peptide 12 (Cyc12).**

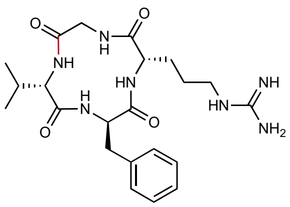

Chemical Formula: $C_{22}H_{33}N_7O_4$
Expected Neutral Mass: 459.2594
Observed Neutral Mass: 459.2604
Mass Error (ppm): 2.2

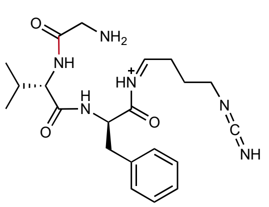

**1**
Chemical Formula: $C_{21}H_{31}N_6O_3^+$
Expected m/z: 415.245
Observed m/z: 415.246
Mass Error (mDa): 1.0

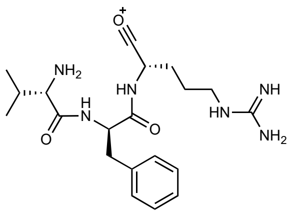

**2**
Chemical Formula: $C_{20}H_{31}N_6O_3^+$
Expected m/z: 403.245
Observed m/z: 403.244
Mass Error (mDa): 1.0

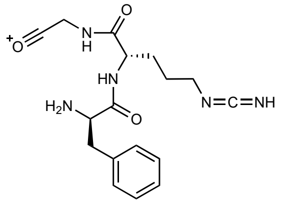

**3**
Chemical Formula: $C_{17}H_{22}N_5O_3^+$
Expected m/z: 344.172
Observed m'z: 344.173
Mass Error (mDa): 1.0

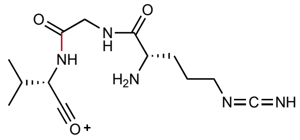

**4**
Chemical Formula: $C_{13}H_{22}N_5O_3^+$
Expected m/z: 296.172
Observed m/z: 296.174
Mass Error (mDa): 2.0

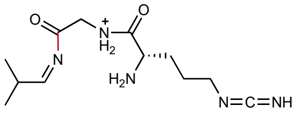

**5**
Chemical Formula: $C_{12}H_{22}N_5O_2^+$
Expected m/z: 268.177
Observed m/z: 268.178
Mass Error (mDa): 1.0

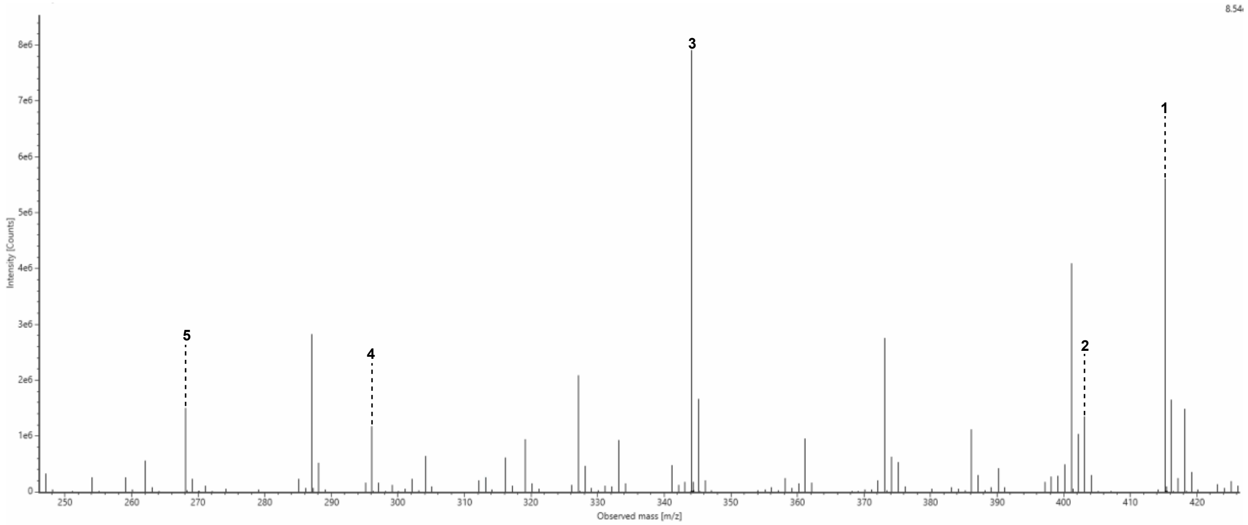

**Figure S96: MS2 spectrum for cyclic peptide 15 (Cyc15).**

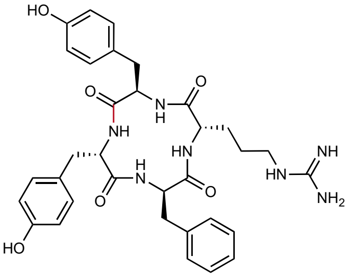

Chemical Formula: $C_{33}H_{39}N_7O_6$
Expected Neutral Mass: 629.2962
Observed Neutral Mass: 629.2974
Mass Error (ppm): 1.9

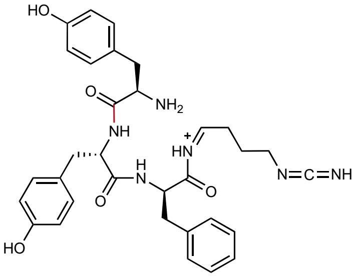

**1**
Chemical Formula: $C_{32}H_{37}N_6O_5^+$
Expected m/z: 585.282
Observed m/z: 585.284
Mass Error (mDa): 2.0

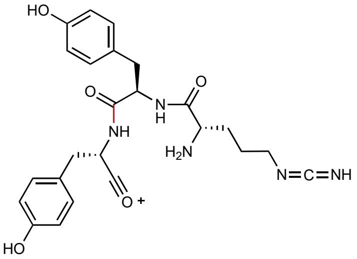

**2**
Chemical Formula: $C_{24}H_{31}N_6O_5^+$
Expected m/z: 483.235
Observed m/z: 483.239
Mass Error (mDa): 4.0

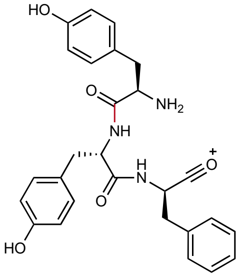

**3**
Chemical Formula: $C_{27}H_{28}N_3O_5^+$
Expected m/z: 474.202
Observed m/z: 474.204
Mass Error (mDa): 2.0

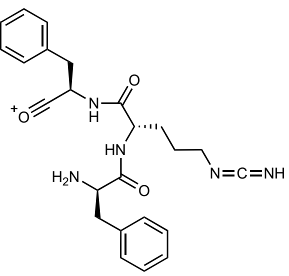

**4**
Chemical Formula: $C_{24}H_{28}N_5O_4^+$
Expected m/z: 450.214
Observed m/z: 450.215
Mass Error (mDa): 1.0

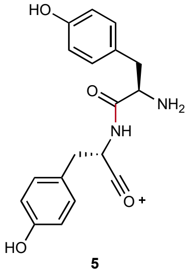

**5**
Chemical Formula: $C_{18}H_{19}N_2O_4^+$
Expected m/z: 327.134
Observed m/z: 327.136
Mass Error (mDa): 2.0

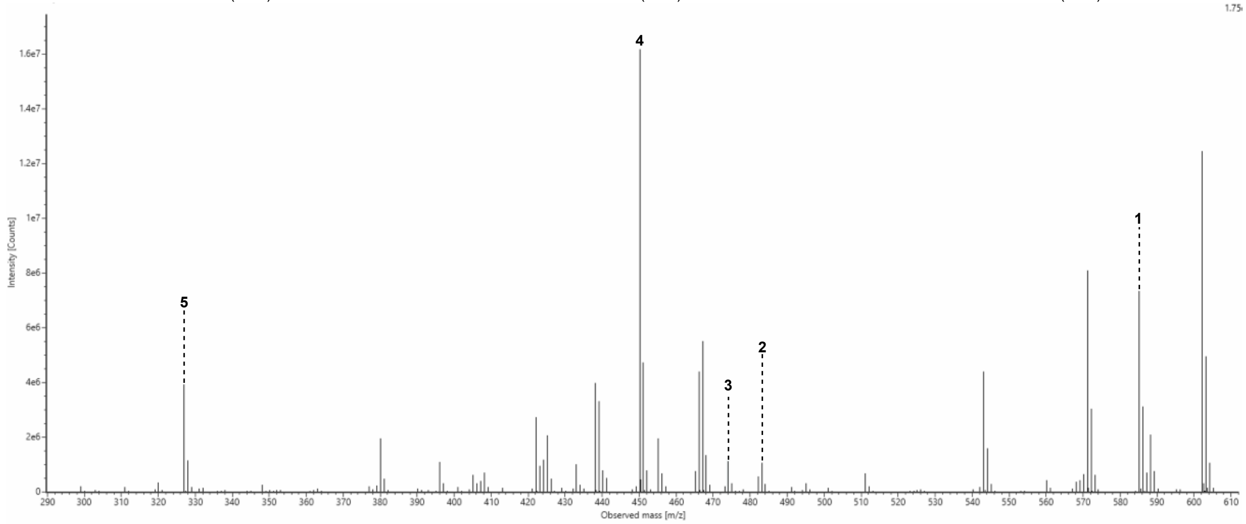

**Figure S97: MS2 spectrum for cyclic peptide 20 (Cyc20).**

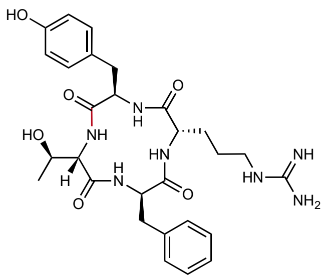

Chemical Formula: $C_{28}H_{37}N_{7}O_{6}$
Expected Neutral Mass: 567.2805
Observed Neutral Mass: 567.2822
Mass Error (ppm): 2.9

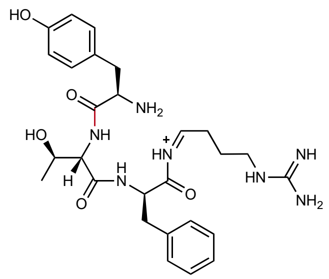

**1**
Chemical Formula: $C_{27}H_{38}N_{7}O_{5}^{+}$
Expected m/z: 540.293
Observed m/z: 540.296
Mass Error (mDa): 3.0

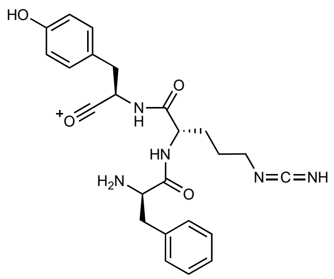

**2**
Chemical Formula: $C_{24}H_{28}N_{5}O_{4}^{+}$
Expected m/z: 450.214
Observed m/z: 450.217
Mass Error (mDa): 3.0

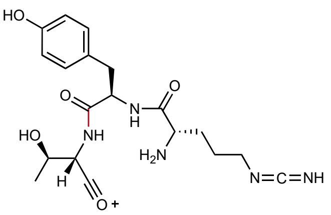

**3**
Chemical Formula: $C_{19}H_{26}N_{5}O_{5}^{+}$
Expected m/z: 404.193
Observed m/z: 404.197
Mass Error (mDa): 4.0

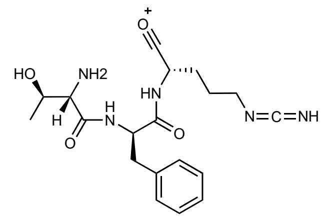

**4**
Chemical Formula: $C_{19}H_{26}N_{5}O_{4}^{+}$
Expected m/z: 388.198
Observed m/z: 388.201
Mass Error (mDa): 3.0

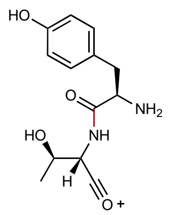

**5**
Chemical Formula: $C_{13}H_{17}N_{2}O_{4}^{+}$
Expected m/z: 265.118
Observed m/z: 265.122
Mass Error (mDa): 4.0

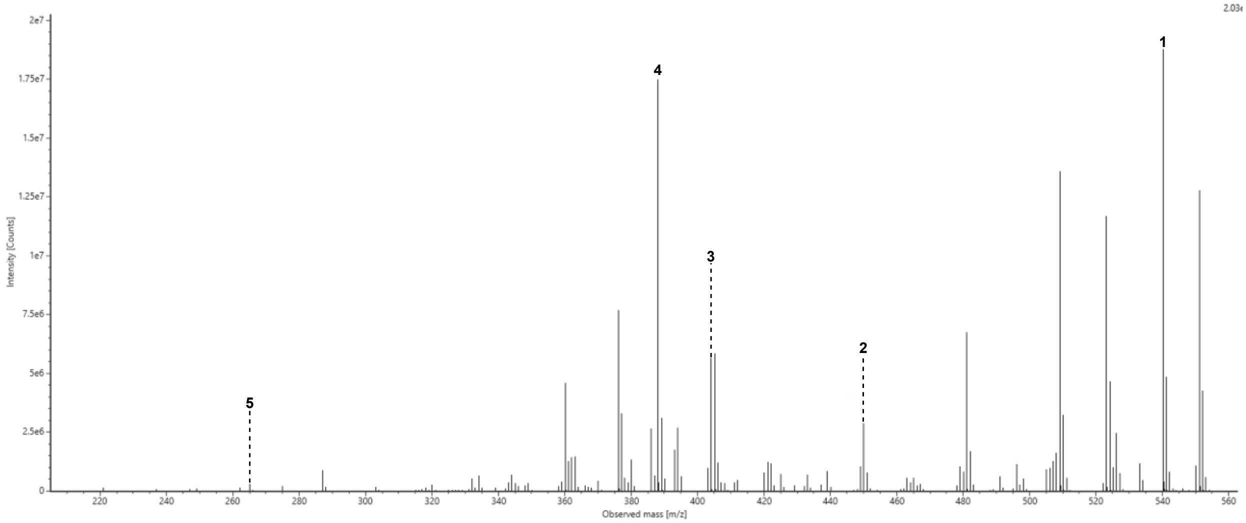

**Figure S98: MS2 spectrum for cyclic peptide 21 (Cyc21).**

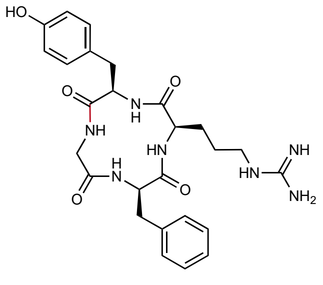

Chemical Formula: $C_{26}H_{33}N_7O_5$
Expected Neutral Mass: 523.2543
Observed Neutral Mass: 523.2562
Mass Error (ppm): 3.6

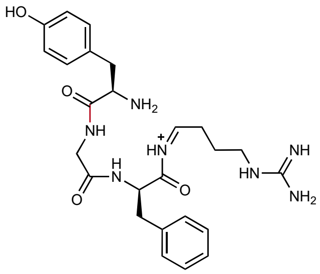

**1**
Chemical Formula: $C_{25}H_{34}N_7O_4^+$
Expected m/z: 496.267
Observed m/z: 496.268
Mass Error (mDa): 1.0

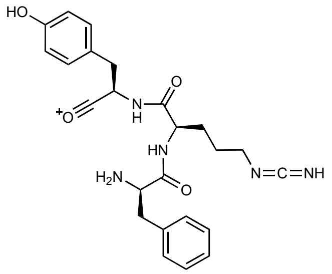

**2**
Chemical Formula: $C_{24}H_{28}N_5O_4^+$
Expected m/z: 450.214
Observed m/z: 450.215
Mass Error (mDa): 1.0

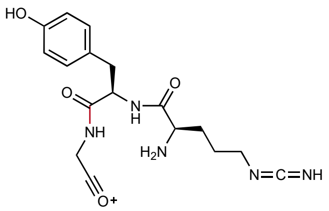

**3**
Chemical Formula: $C_{17}H_{22}N_5O_4^+$
Expected m/z: 360.167
Observed m/z: 360.170
Mass Error (mDa): 3.0

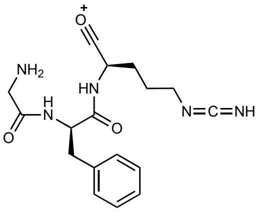

**4**
Chemical Formula: $C_{17}H_{22}N_5O_3^+$
Expected m/z: 344.172
Observed m/z: 344.174
Mass Error (mDa): 2.0

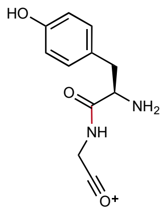

**5**
Chemical Formula: $C_{11}H_{13}N_2O_3^+$
Expected m/z: 221.092
Observed m/z: 221.094
Mass Error (mDa): 2.0

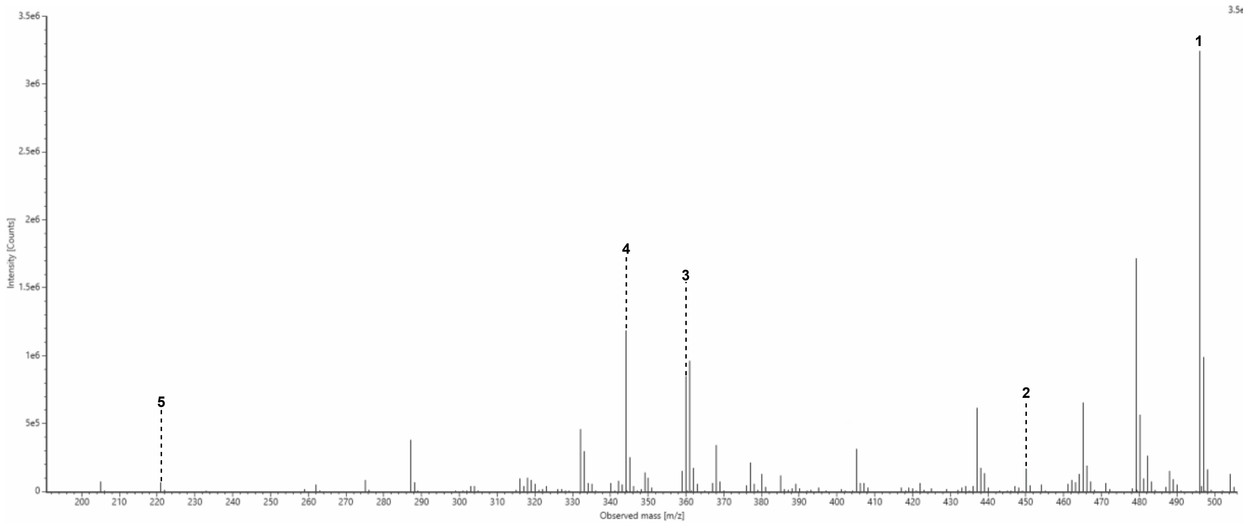

**Figure S99: MS2 spectrum for cyclic peptide 22 (Cyc22).**

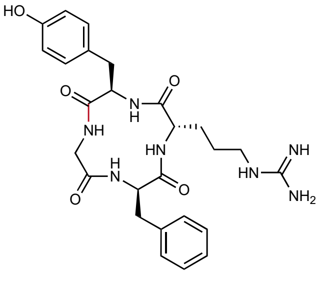

Chemical Formula: $C_{26}H_{33}N_7O_5$
Expected Neutral Mass: 523.2543
Observed Neutral Mass: 523.2559
Mass Error (ppm): 3.1

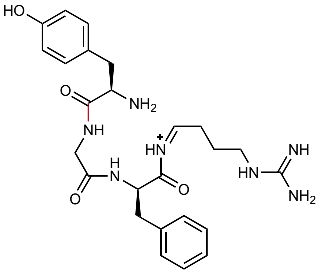

1
Chemical Formula: $C_{25}H_{34}N_7O_4^+$
Expected m/z: 496.267
Observed m/z: 496.268
Mass Error (mDa): 1.0

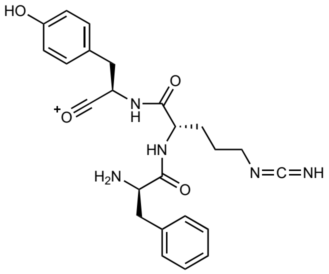

2
Chemical Formula: $C_{24}H_{28}N_5O_4^+$
Expected m/z: 450.214
Observed m/z: 450.215
Mass Error (mDa): 1.0

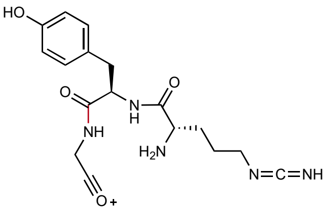

3
Chemical Formula: $C_{17}H_{22}N_5O_4^+$
Expected m/z: 360.167
Observed m/z: 360.170
Mass Error (mDa): 3.0

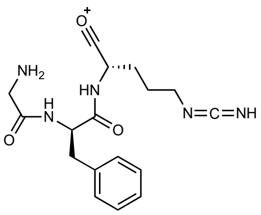

4
Chemical Formula: $C_{17}H_{22}N_5O_3^+$
Expected m/z: 344.172
Observed m/z: 344.174
Mass Error (mDa): 2.0

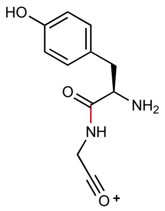

5
Chemical Formula: $C_{11}H_{13}N_2O_3^+$
Expected m/z: 221.092
Observed m/z: 221.095
Mass Error (mDa): 3.0

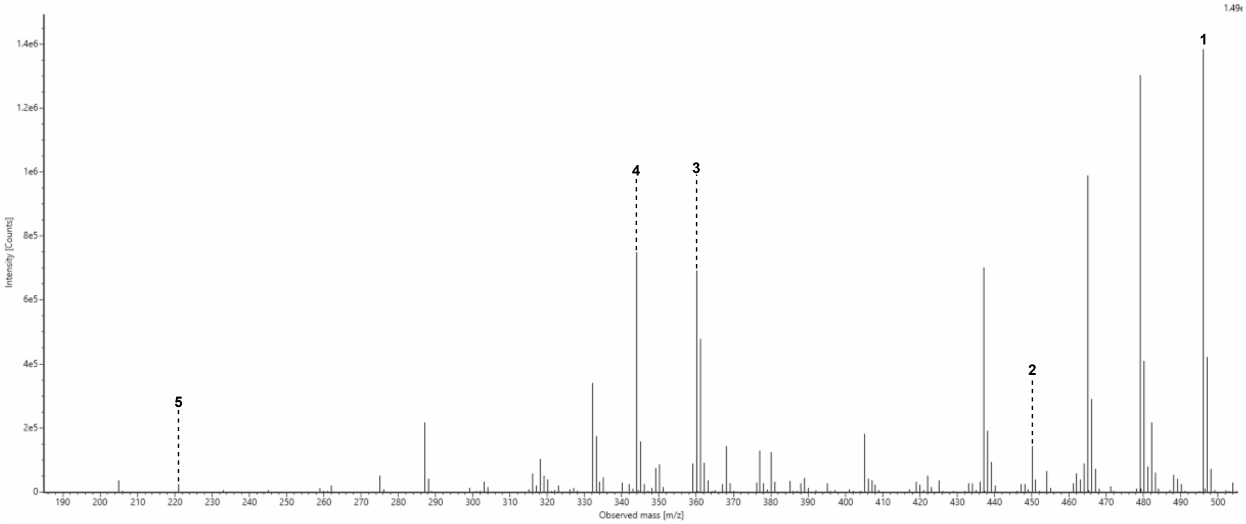

**Figure S100: MS2 spectrum for cyclic peptide 23 (Cyc23).**

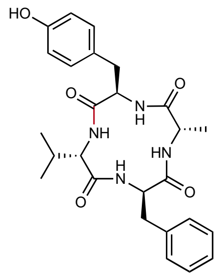

Chemical Formula: $C_{26}H_{32}N_4O_5$
Expected Neutral Mass: 480.2373
Observed Neutral Mass: 480.2379
Mass Error (ppm): 1.2

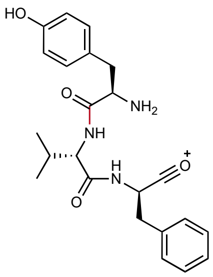

**1**
Chemical Formula: $C_{23}H_{28}N_3O_4^+$
Expected m/z: 410.207
Observed m/z: 410.210
Mass Error (mDa): 3.0

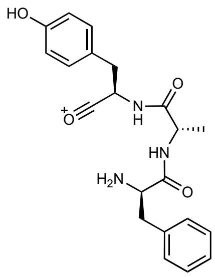

**2**
Chemical Formula: $C_{21}H_{24}N_3O_4^+$
Expected m/z: 382.176
Observed m/z: 382.181
Mass Error (mDa): 5.0

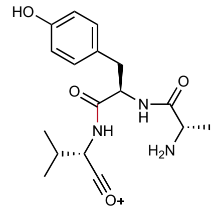

**3**
Chemical Formula: $C_{17}H_{24}N_3O_4^+$
Expected m/z: 334.176
Observed m/z: 334.179
Mass Error (mDa): 3.0

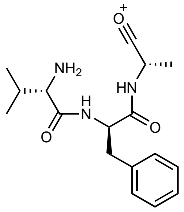

**4**
Chemical Formula: $C_{17}H_{24}N_3O_3^+$
Expected m/z: 318.181
Observed m/z: 318.184
Mass Error (mDa): 3.0

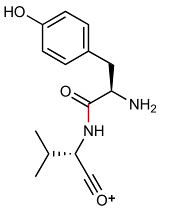

**5**
Chemical Formula: $C_{14}H_{19}N_2O_3^+$
Expected m/z: 263.139
Observed m/z: 263.142
Mass Error (mDa): 3.0

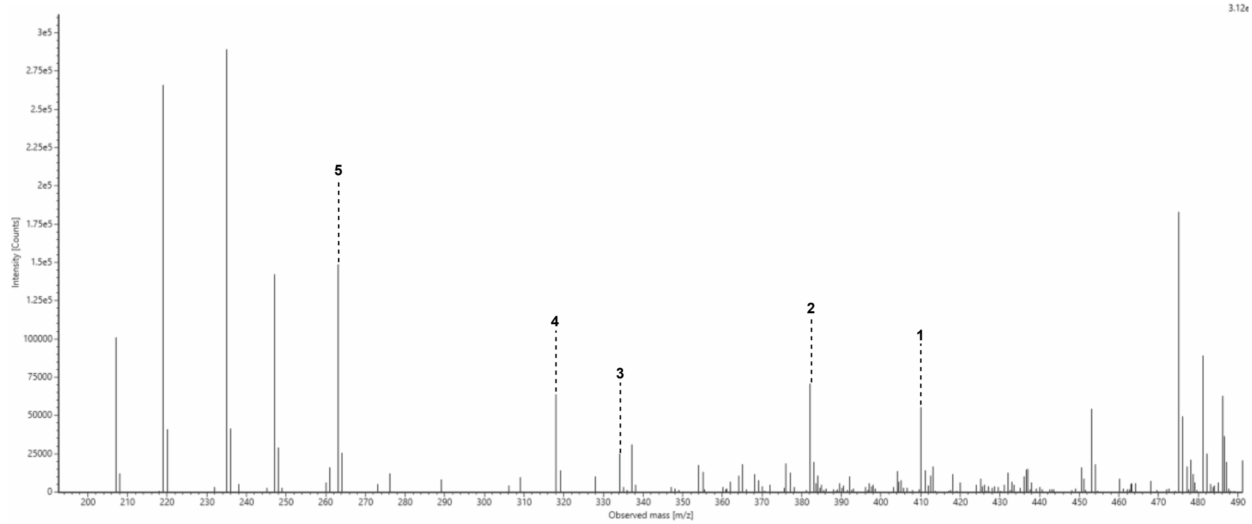

**Figure S101: MS2 spectrum for cyclic peptide 24 (Cyc24).**

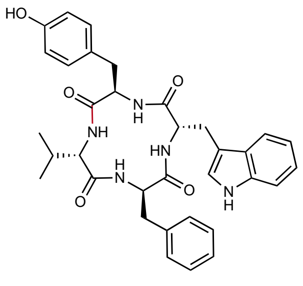

Chemical Formula: $C_{34}H_{37}N_5O_5$
Expected Neutral Mass: 595.2795
Observed Neutral Mass: 595.2779
Mass Error (ppm): 2.7

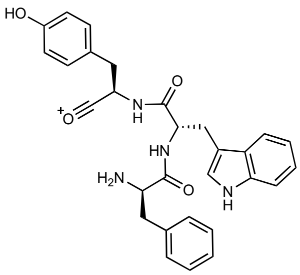

1
Chemical Formula: $C_{29}H_{29}N_4O_4^+$
Expected m/z: 497.218
Observed m/z: 497.217
Mass Error (mDa): 1.0

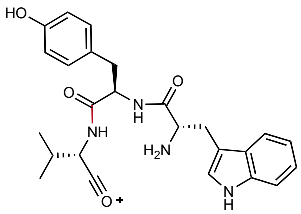

2
Chemical Formula: $C_{25}H_{29}N_4O_4^+$
Expected m/z: 449.218
Observed m/z: 449.219
Mass Error (mDa): 1.0

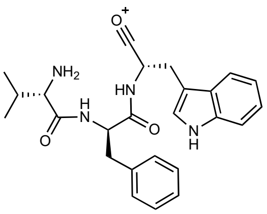

3
Chemical Formula: $C_{25}H_{29}N_4O_3^+$
Expected m/z: 433.223
Observed m/z: 433.222
Mass Error (mDa): 1.0

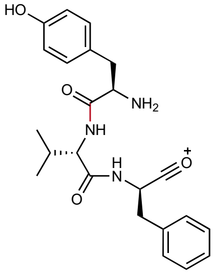

4
Chemical Formula: $C_{23}H_{28}N_3O_4^+$
Expected m/z: 410.207
Observed m/z: 410.209
Mass Error (mDa): 1.0

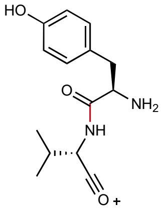

5
Chemical Formula: $C_{14}H_{19}N_2O_3^+$
Expected m/z: 263.139
Observed m/z: 263.141
Mass Error (mDa): 2.0

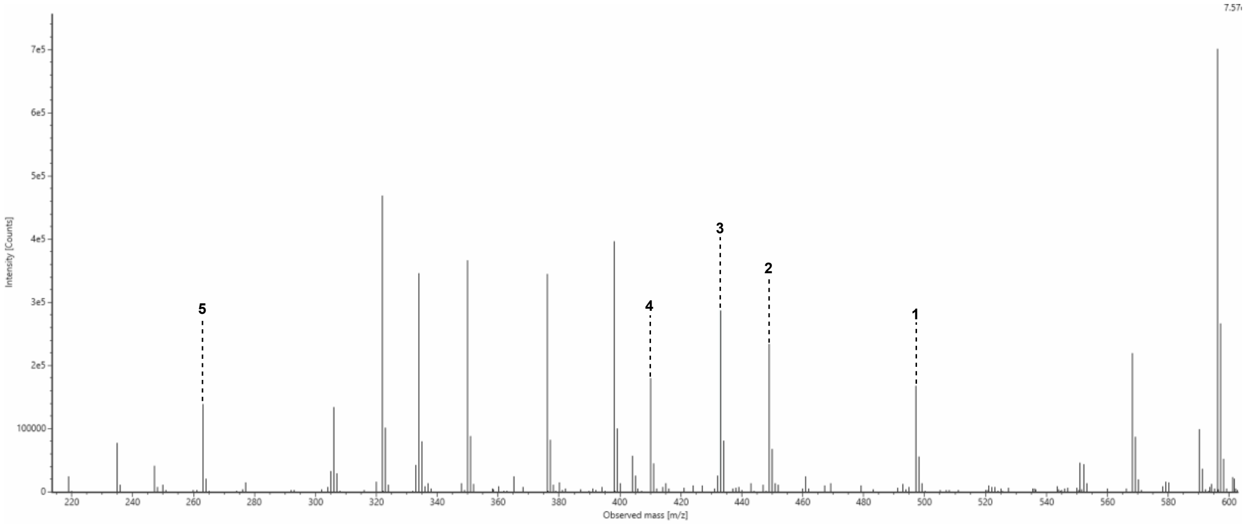

**Figure S102: MS2 spectrum for cyclic peptide 25 (Cyc25).**

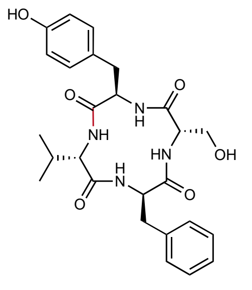

Chemical Formula: $C_{26}H_{32}N_4O_6$
Expected Neutral Mass: 496.2322
Observed Neutral Mass: 496.2317
Mass Error (ppm): 1.1

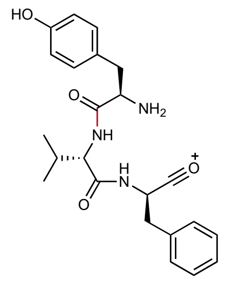

**1**
Chemical Formula: $C_{23}H_{28}N_3O_4^+$
Expected m/z: 410.207
Observed m/z: 410.209
Mass Error (mDa): 2.0

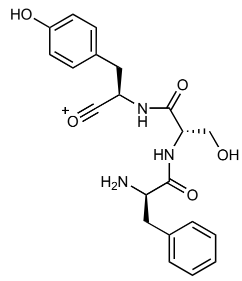

**2**
Chemical Formula: $C_{21}H_{24}N_3O_5^+$
Expected m/z: 398.171
Observed m/z: 398.174
Mass Error (mDa): 3.0

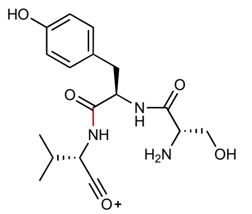

**3**
Chemical Formula: $C_{17}H_{24}N_3O_5^+$
Expected m/z: 350.171
Observed m/z: 350.170
Mass Error (mDa): 1.0

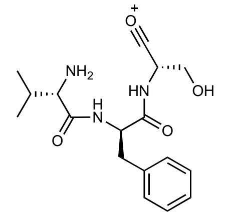

**4**
Chemical Formula: $C_{17}H_{24}N_3O_4^+$
Expected m/z: 334.176
Observed m/z: 334.179
Mass Error (mDa): 3.0

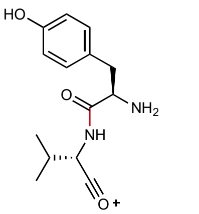

**5**
Chemical Formula: $C_{14}H_{19}N_2O_3^+$
Expected m/z: 263.139
Observed m/z: 263.141
Mass Error (mDa): 2.0

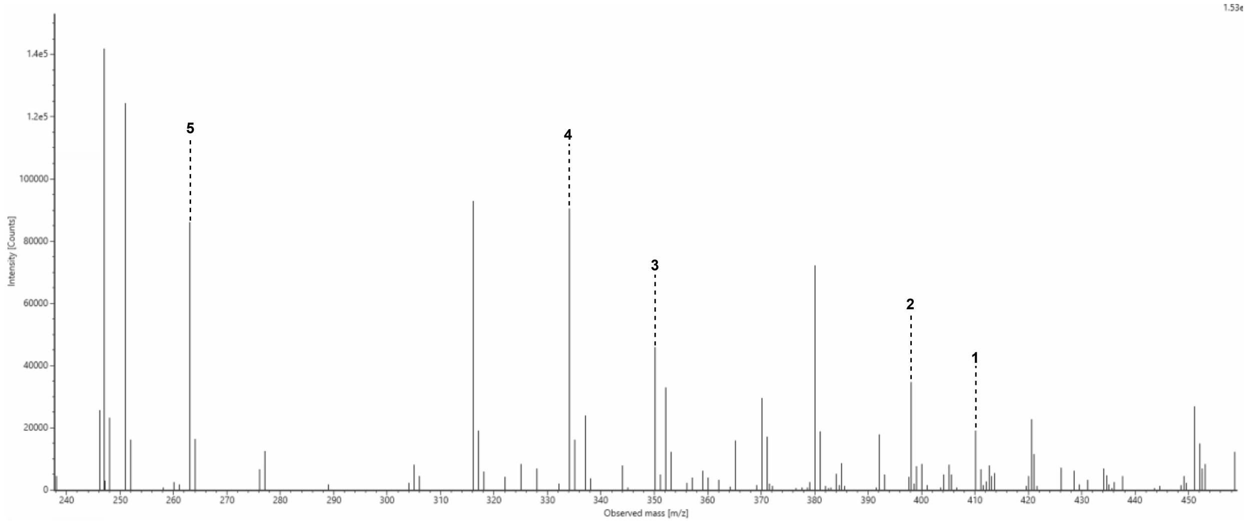

**Figure S103: MS2 spectrum for cyclic peptide 26 (Cyc26).**

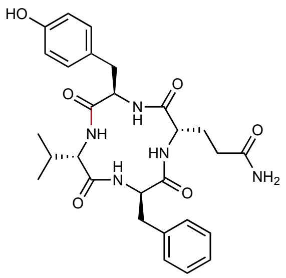

Chemical Formula: $C_{28}H_{35}N_5O_6$
Expected Neutral Mass: 537.2587
Observed Neutral Mass: 537.2584
Mass Error (ppm): 0.7

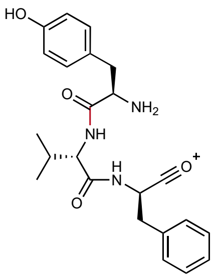

**1**
Chemical Formula: $C_{23}H_{28}N_3O_4^+$
Expected m/z: 410.207
Observed m/z: 410.208
Mass Error (mDa): 1.0

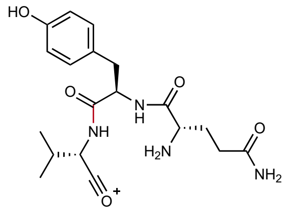

**2**
Chemical Formula: $C_{19}H_{27}N_4O_5^+$
Expected m/z: 391.198
Observed m/z: 391.199
Mass Error (mDa): 1.0

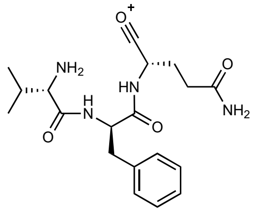

**3**
Chemical Formula: $C_{19}H_{27}N_4O_4^+$
Expected m/z: 375.203
Observed m/z: 375.203
Mass Error (mDa): < 1.0

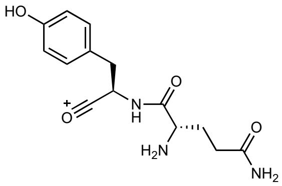

**4**
Chemical Formula: $C_{14}H_{18}N_3O_4^+$
Expected m/z: 292.129
Observed m/z: 292.131
Mass Error (mDa): 2.0

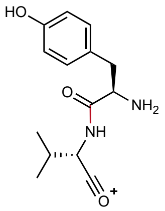

**5**
Chemical Formula: $C_{14}H_{19}N_2O_3^+$
Expected m/z: 263.139
Observed m/z: 263.141
Mass Error (mDa): 2.0

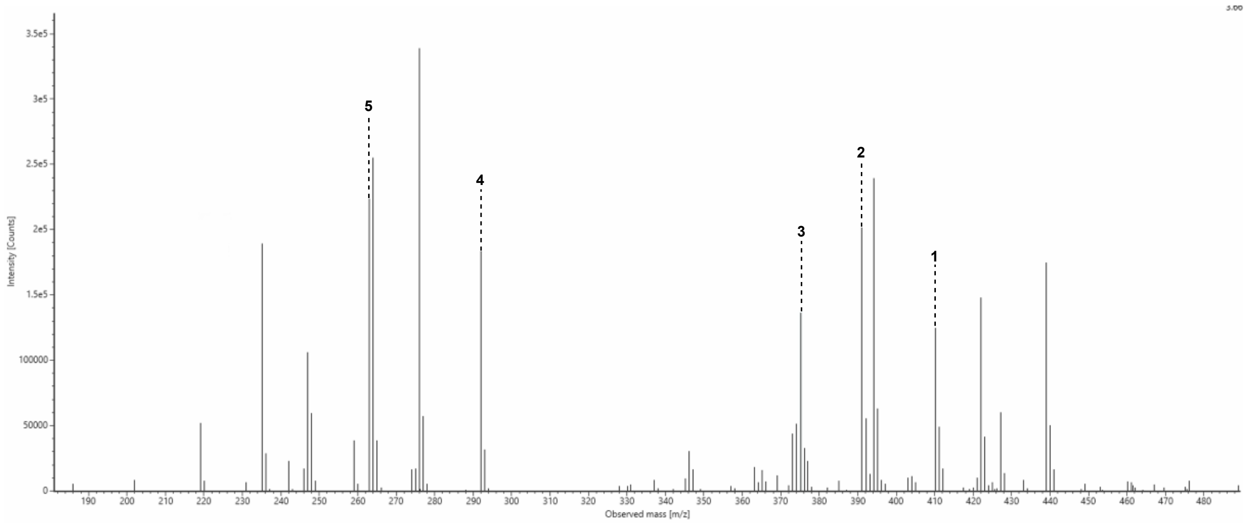

**Figure S104: MS2 spectrum for cyclic peptide 27 (Cyc27).**

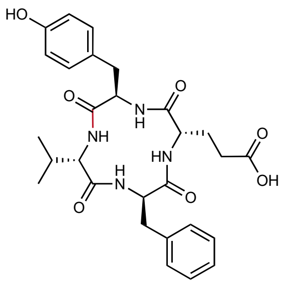

Chemical Formula: $C_{28}H_{34}N_4O_7$
Expected Neutral Mass: 538.2428
Observed Neutral Mass: 538.2439
Mass Error (ppm): 2.1

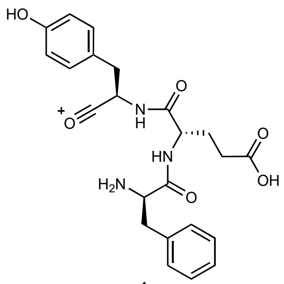

**1**
Chemical Formula: $C_{23}H_{26}N_3O_6^+$
Expected m/z: 440.182
Observed m/z: 440.182
Mass Error (mDa): < 1.0

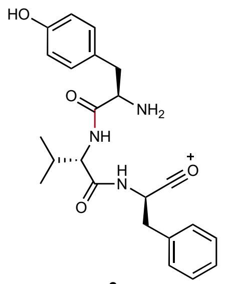

**2**
Chemical Formula: $C_{23}H_{28}N_3O_4^+$
Expected m/z: 410.207
Observed m/z: 410.207
Mass Error (mDa): < 1.0

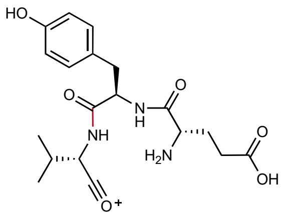

**3**
Chemical Formula: $C_{19}H_{26}N_3O_6^+$
Expected m/z: 392.182
Observed m/z: 392.184
Mass Error (mDa): 2.0

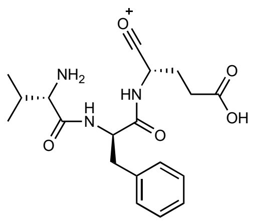

**4**
Chemical Formula: $C_{19}H_{26}N_3O_5^+$
Expected m/z: 376.187
Observed m/z: 376.187
Mass Error (mDa): < 1.0

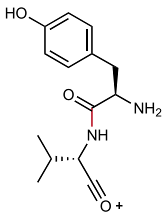

**5**
Chemical Formula: $C_{14}H_{19}N_2O_3^+$
Expected m/z: 263.139
Observed m/z: 263.140
Mass Error (mDa): 1.0

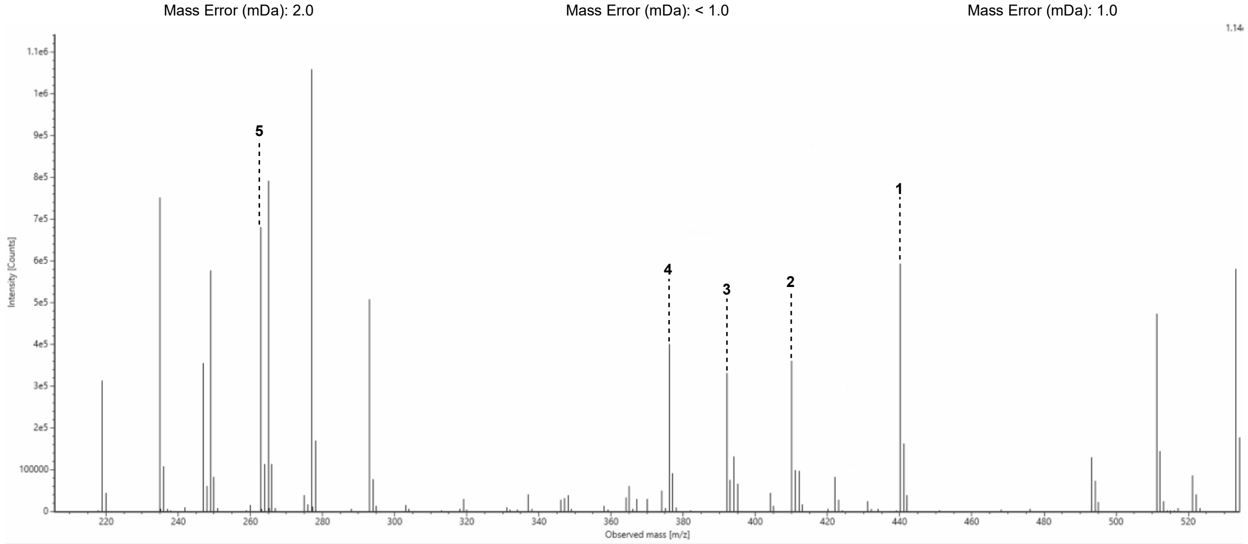

**Figure S105: MS2 spectrum for cyclic peptide 28 (Cyc28).**

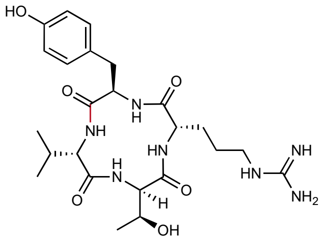

Chemical Formula: $C_{24}H_{37}N_7O_6$
Expected Neutral Mass: 519.2805
Observed Neutral Mass: 519.2817
Mass Error (ppm): 2.2

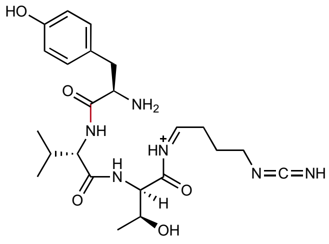

1
Chemical Formula: $C_{23}H_{35}N_6O_5^+$
Expected m/z: 475.266
Observed m/z: 475.268
Mass Error (mDa): 2.0

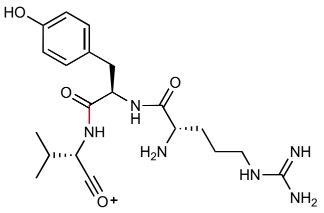

2
Chemical Formula: $C_{20}H_{31}N_6O_4^+$
Expected m/z: 419.240
Observed m/z: 419.246
Mass Error (mDa): 6.0

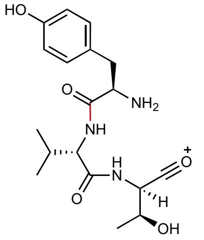

3
Chemical Formula: $C_{18}H_{26}N_3O_5^+$
Expected m/z: 364.187
Observed m/z: 364.190
Mass Error (mDa): 3.0

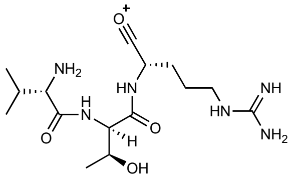

4
Chemical Formula: $C_{15}H_{29}N_6O_4^+$
Expected m/z: 357.224
Observed m/z: 357.226
Mass Error (mDa): 2.0

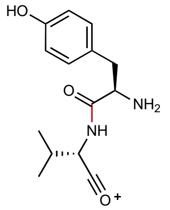

5
Chemical Formula: $C_{14}H_{19}N_2O_3^+$
Expected m/z: 263.139
Observed m/z: 263.141
Mass Error (mDa): 2.0

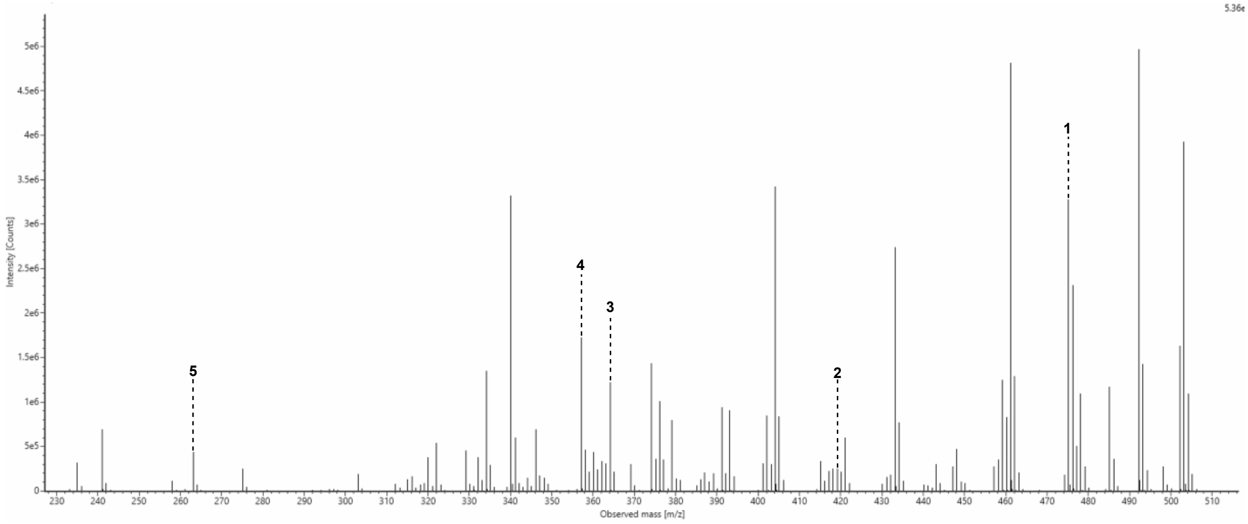

**Figure S106: MS2 spectrum for cyclic peptide 29 (Cyc29).**

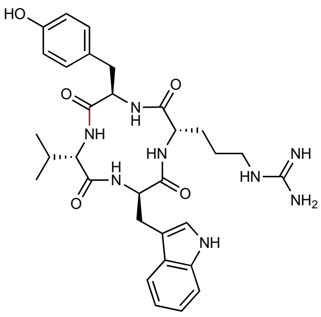

Chemical Formula: $C_{31}H_{40}N_8O_5$
Expected Neutral Mass: 604.3122
Observed Neutral Mass: 604.3138
Mass Error (ppm): 2.6

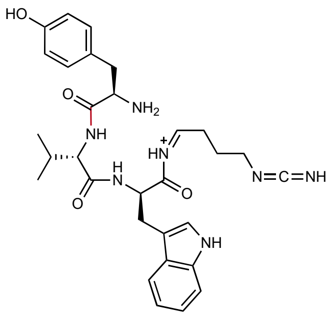

**1**
Chemical Formula: $C_{30}H_{38}N_7O_4^+$
Expected m/z: 560.298
Observed m/z: 560.299
Mass Error (mDa): 1.0

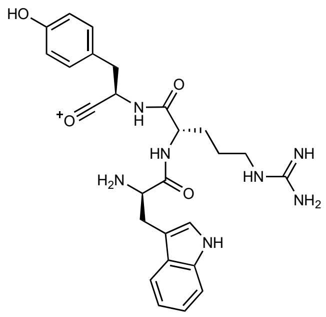

**2**
Chemical Formula: $C_{26}H_{32}N_7O_4^+$
Expected m/z: 506.251
Observed m/z: 506.254
Mass Error (mDa): 3.0

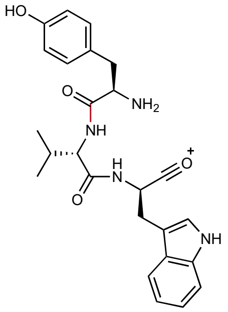

**3**
Chemical Formula: $C_{25}H_{29}N_4O_4^+$
Expected m/z: 449.218
Observed m/z: 449.221
Mass Error (mDa): 3.0

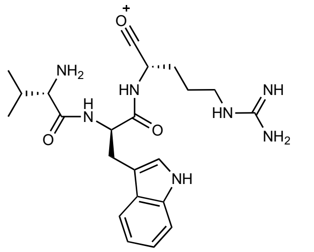

**4**
Chemical Formula: $C_{22}H_{32}N_7O_3^+$
Expected m/z: 442.256
Observed m/z: 442.257
Mass Error (mDa): 1.0

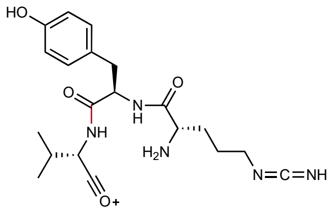

**5**
Chemical Formula: $C_{20}H_{28}N_5O_4^+$
Expected m/z: 402.214
Observed m/z: 402.213
Mass Error (mDa): 1.0

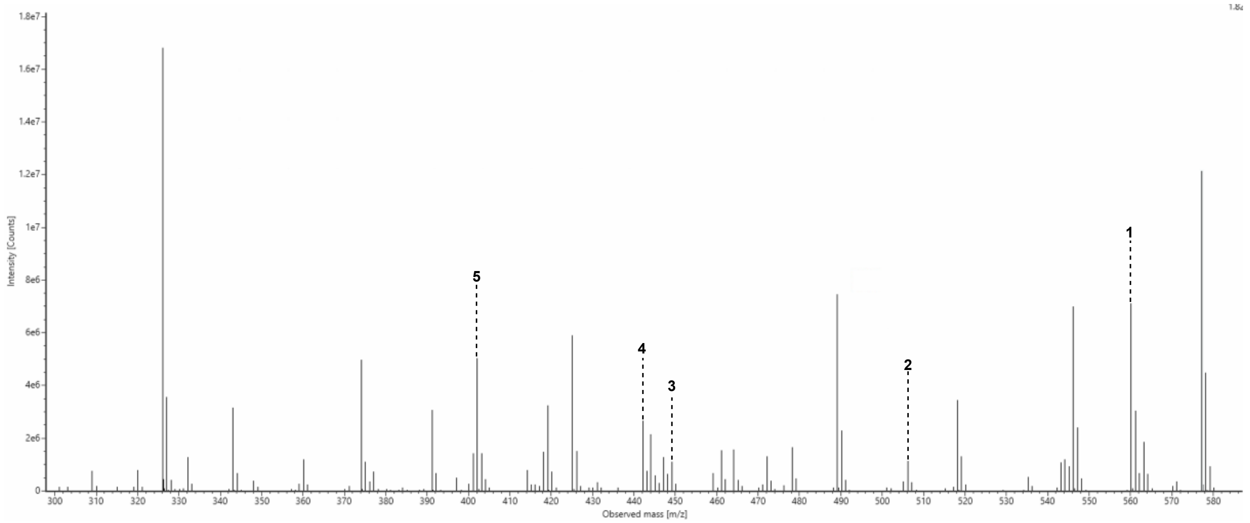

**Figure S107: MS2 spectrum for cyclic peptide 30 (Cyc30).**

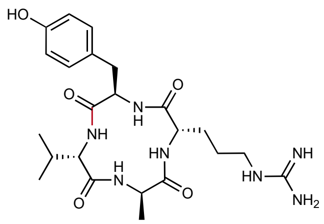

Chemical Formula: $C_{23}H_{35}N_7O_5$
Expected Neutral Mass: 489.2700
Observed Neutral Mass: 489.2700
Mass Error (ppm): 0.1

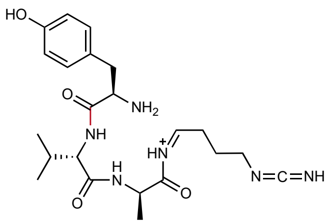

**1**
Chemical Formula: $C_{22}H_{33}N_6O_4^+$
Expected m/z: 445.256
Observed m/z: 445.256
Mass Error (mDa): < 1.0

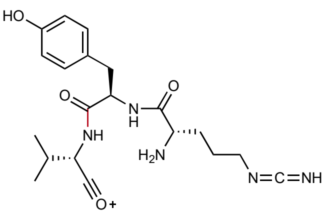

**2**
Chemical Formula: $C_{20}H_{28}N_5O_4^+$
Expected m/z: 402.214
Observed m/z: 402.217
Mass Error (mDa): 3.0

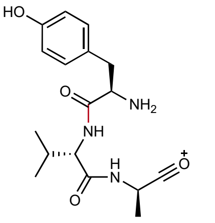

**3**
Chemical Formula: $C_{17}H_{24}N_3O_4^+$
Expected m/z: 334.176
Observed m/z: 334.177
Mass Error (mDa): 1.0

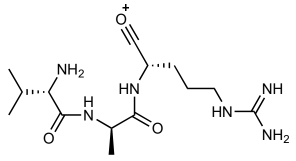

**4**
Chemical Formula: $C_{14}H_{27}N_6O_3^+$
Expected m/z: 327.214
Observed m/z: 327.215
Mass Error (mDa): 1.0

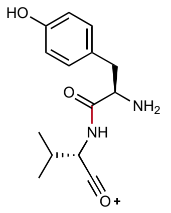

**5**
Chemical Formula: $C_{14}H_{19}N_2O_3^+$
Expected m/z: 263.139
Observed m/z: 263.140
Mass Error (mDa): 1.0

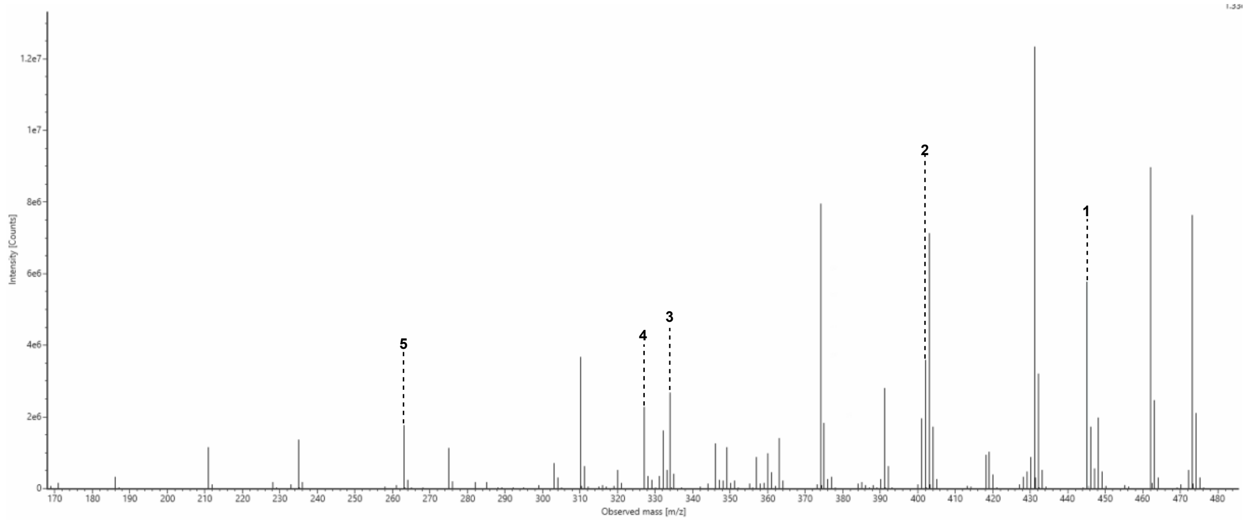

**Figure S108: MS2 spectrum for cyclic peptide 31 (Cyc31).**

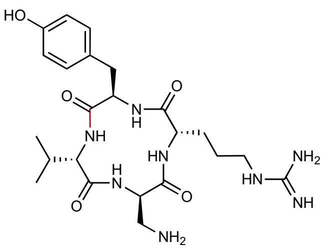

Chemical Formula: $C_{23}H_{36}N_8O_5$
Expected Neutral Mass: 504.281
Observed Neutral Mass: 504.282
Mass Error (ppm): 2.8

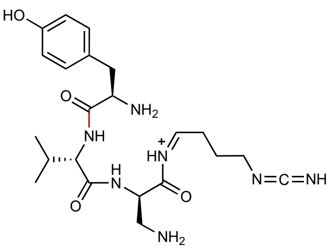

**1**
Chemical Formula: $C_{22}H_{34}N_7O_4^+$
Expected m/z: 460.267
Observed m/z: 460.265
Mass Error (mDa): 2.0

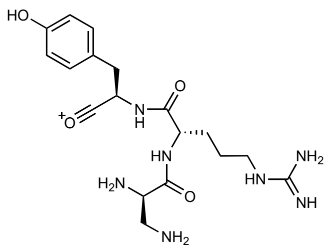

**2**
Chemical Formula: $C_{18}H_{28}N_7O_4^+$
Expected m/z: 406.220
Observed m/z: 406.224
Mass Error (mDa): 4.0

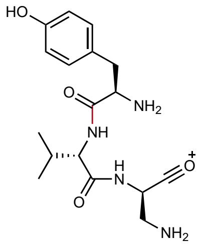

**3**
Chemical Formula: $C_{17}H_{25}N_4O_4^+$
Expected m/z: 349.187
Observed m/z: 349.189
Mass Error (mDa): 2.0

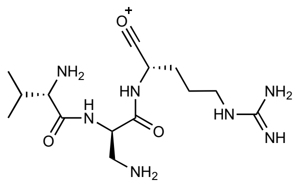

**4**
Chemical Formula: $C_{14}H_{28}N_7O_3^+$
Expected m/z: 342.225
Observed m/z: 342.224
Mass Error (mDa): 1.0

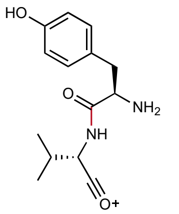

**5**
Chemical Formula: $C_{14}H_{19}N_2O_3^+$
Expected m/z: 263.139
Observed m/z: 263.140
Mass Error (mDa): 1.0

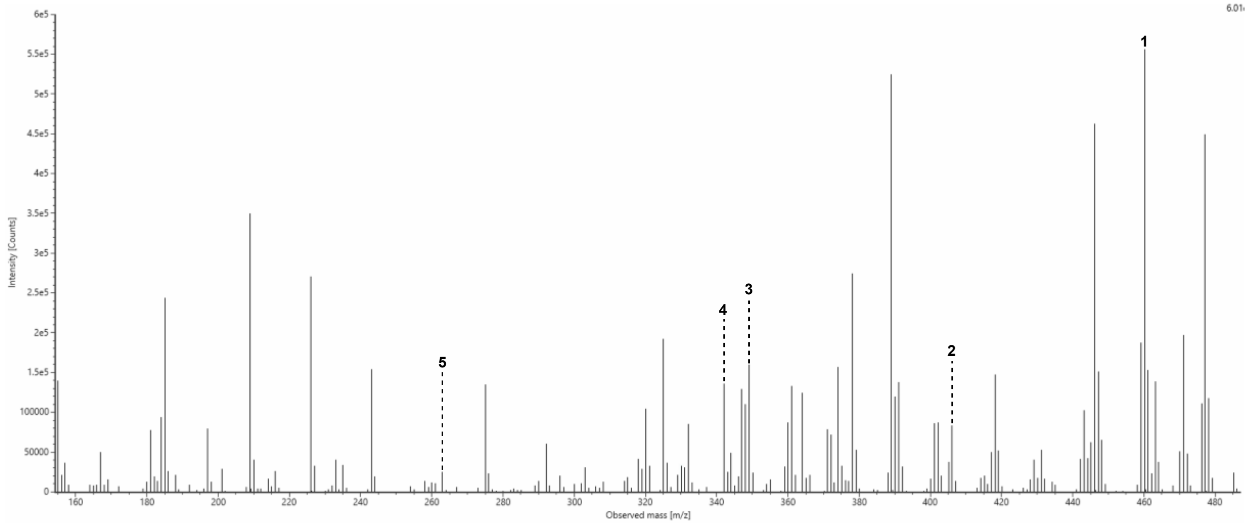

**Figure S109: MS2 spectrum for cyclic peptide 32 (Cyc32).**

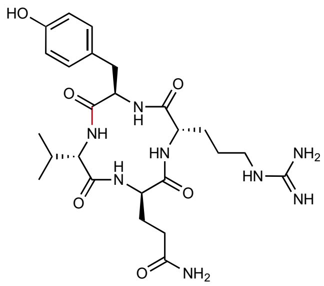

Chemical Formula: $C_{25}H_{38}N_8O_6$
Expected Neutral Mass: 546.2914
Observed Neutral Mass: 546.2912
Mass Error (ppm): 0.4

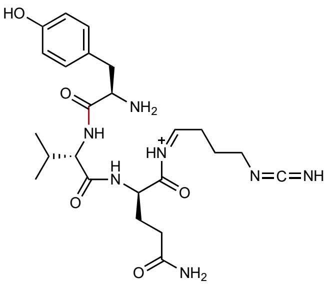

**1**
Chemical Formula: $C_{24}H_{36}N_7O_5^+$
Expected m/z: 502.277
Observed m/z: 502.277
Mass Error (mDa): < 1.0

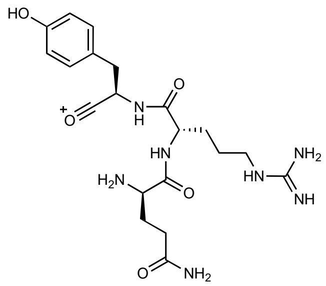

**2**
Chemical Formula: $C_{20}H_{30}N_7O_5^+$
Expected m/z: 448.230
Observed m/z: 448.232
Mass Error (mDa): 2.0

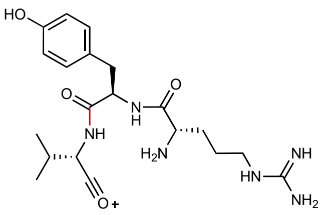

**3**
Chemical Formula: $C_{20}H_{31}N_6O_4$
Expected m/z: 419.240
Observed m/z: 419.241
Mass Error (mDa): 1.0

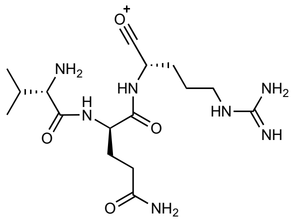

**4**
Chemical Formula: $C_{16}H_{30}N_7O_4^+$
Expected m/z: 384.235
Observed m/z: 384.232
Mass Error (mDa): 3.0

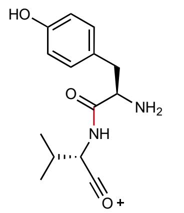

**5**
Chemical Formula: $C_{14}H_{19}N_2O_3^+$
Expected m/z: 263.139
Observed m/z: 263.139
Mass Error (mDa): < 1.0

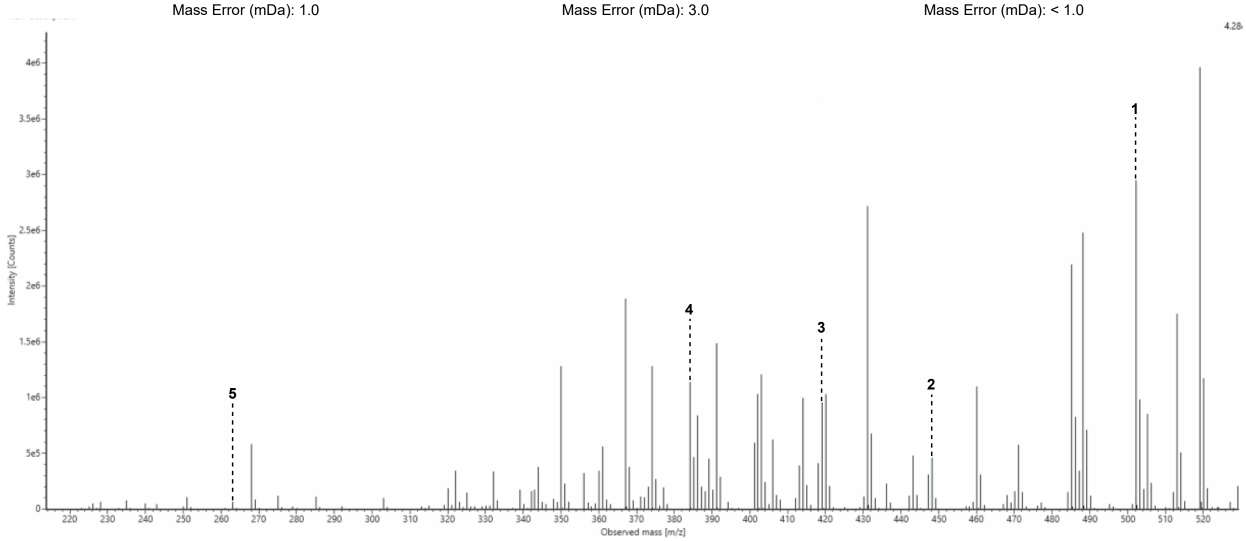

**Figure S110: MS2 spectrum for cyclic peptide 33 (Cyc33).**

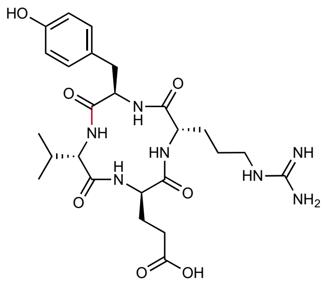

Chemical Formula: $C_{25}H_{37}N_7O_7$
Expected Neutral Mass: 547.2755
Observed Neutral Mass: 547.2773
Mass Error (ppm): 3.4

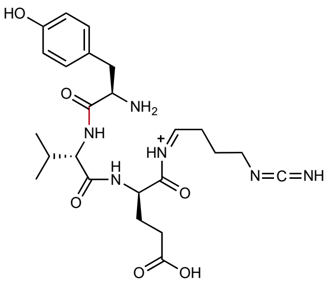

**1**
Chemical Formula: $C_{24}H_{35}N_6O_6^+$
Expected m/z: 503.261
Observed m/z: 503.264
Mass Error (mDa): 3.0

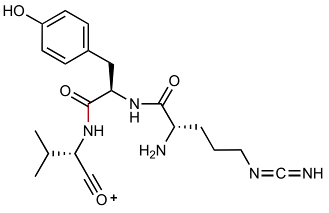

**2**
Chemical Formula: $C_{20}H_{28}N_5O_4^+$
Expected m/z: 402.214
Observed m/z: 402.218
Mass Error (mDa): 4.0

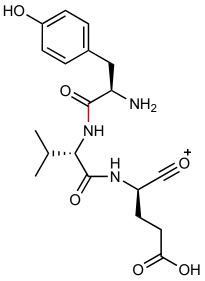

**3**
Chemical Formula: $C_{19}H_{26}N_3O_6^+$
Expected m/z: 392.182
Observed m/z: 392.185
Mass Error (mDa): 3.0

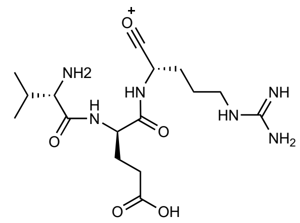

**4**
Chemical Formula: $C_{16}H_{29}N_6O_5^+$
Expected m/z: 385.219
Observed m/z: 385.221
Mass Error (mDa): 2.0

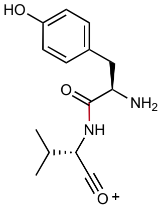

**5**
Chemical Formula: $C_{14}H_{19}N_2O_3^+$
Expected m/z: 263.139
Observed m/z: 263.142
Mass Error (mDa): 3.0

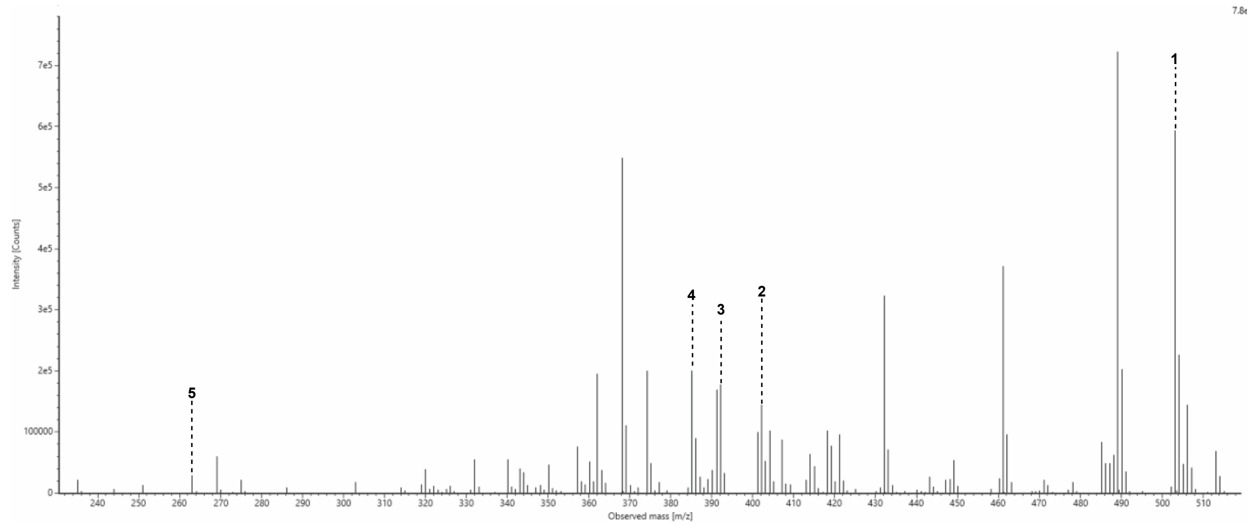

**Figure S111: MS2 spectrum for cyclic peptide 34 (Cyc34).**

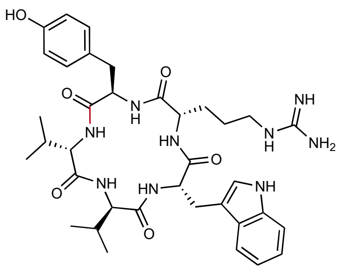

Chemical Formula: $C_{36}H_{49}N_9O_6$
Expected Neutral Mass: 703.3806
Observed Neutral Mass: 703.3827
Mass Error (ppm): 3.0

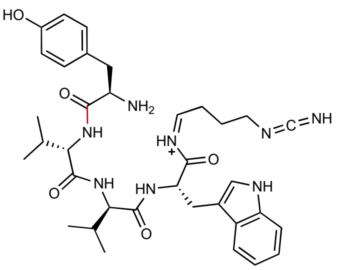

1
Chemical Formula: $C_{35}H_{47}N_8O_5^+$
Expected m/z: 659.366
Observed m/z: 659.368
Mass Error (mDa): 2.0

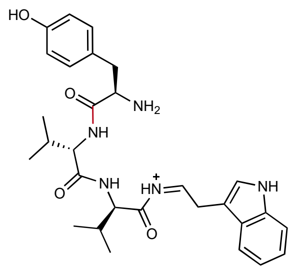

2
Chemical Formula: $C_{29}H_{38}N_5O_4^+$
Expected m/z: 520.292
Observed m/z: 520.295
Mass Error (mDa): 3.0

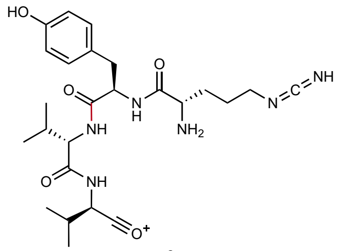

3
Chemical Formula: $C_{25}H_{37}N_6O_5^+$
Expected m/z: 501.282
Observed m/z: 501.284
Mass Error (mDa): 2.0

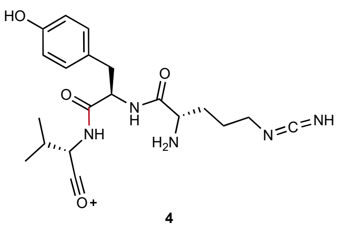

4
Chemical Formula: $C_{20}H_{28}N_5O_4^+$
Expected m/z: 402.214
Observed m/z: 402.216
Mass Error (mDa): 2.0

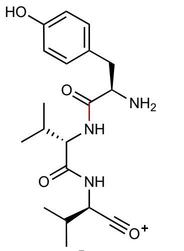

5
Chemical Formula: $C_{19}H_{28}N_3O_4^+$
Expected m/z: 362.207
Observed m/z: 362.209
Mass Error (mDa): 2.0

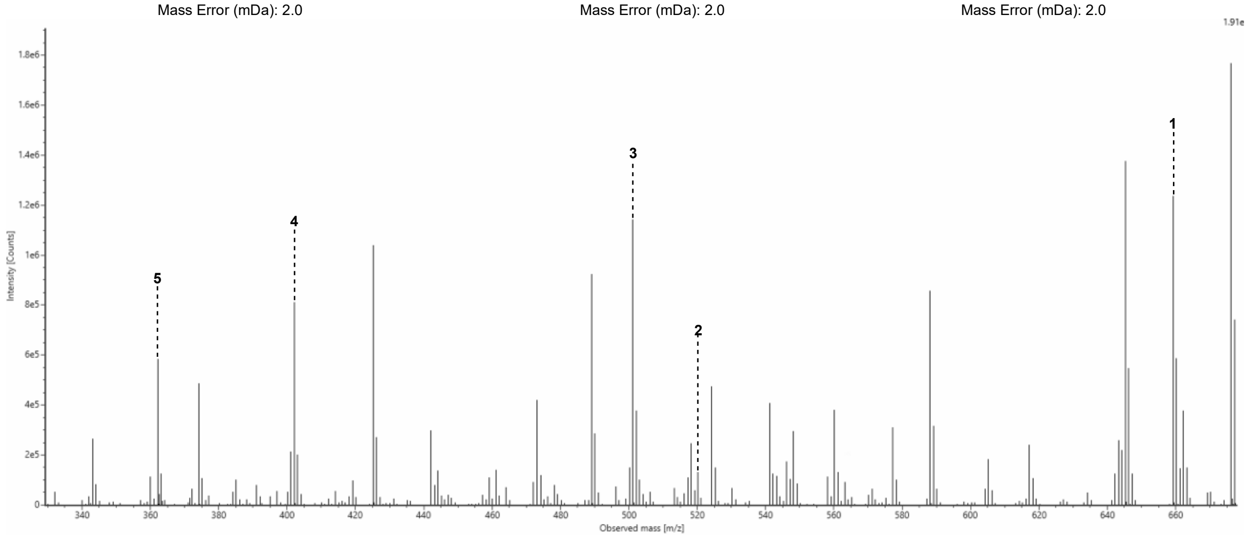

**Figure S112: MS2 spectrum for cyclic peptide 38 (Cyc38).**

**SI Figures S113 to S146. 1H and 13C NMR spectra of peptide thioesters.**

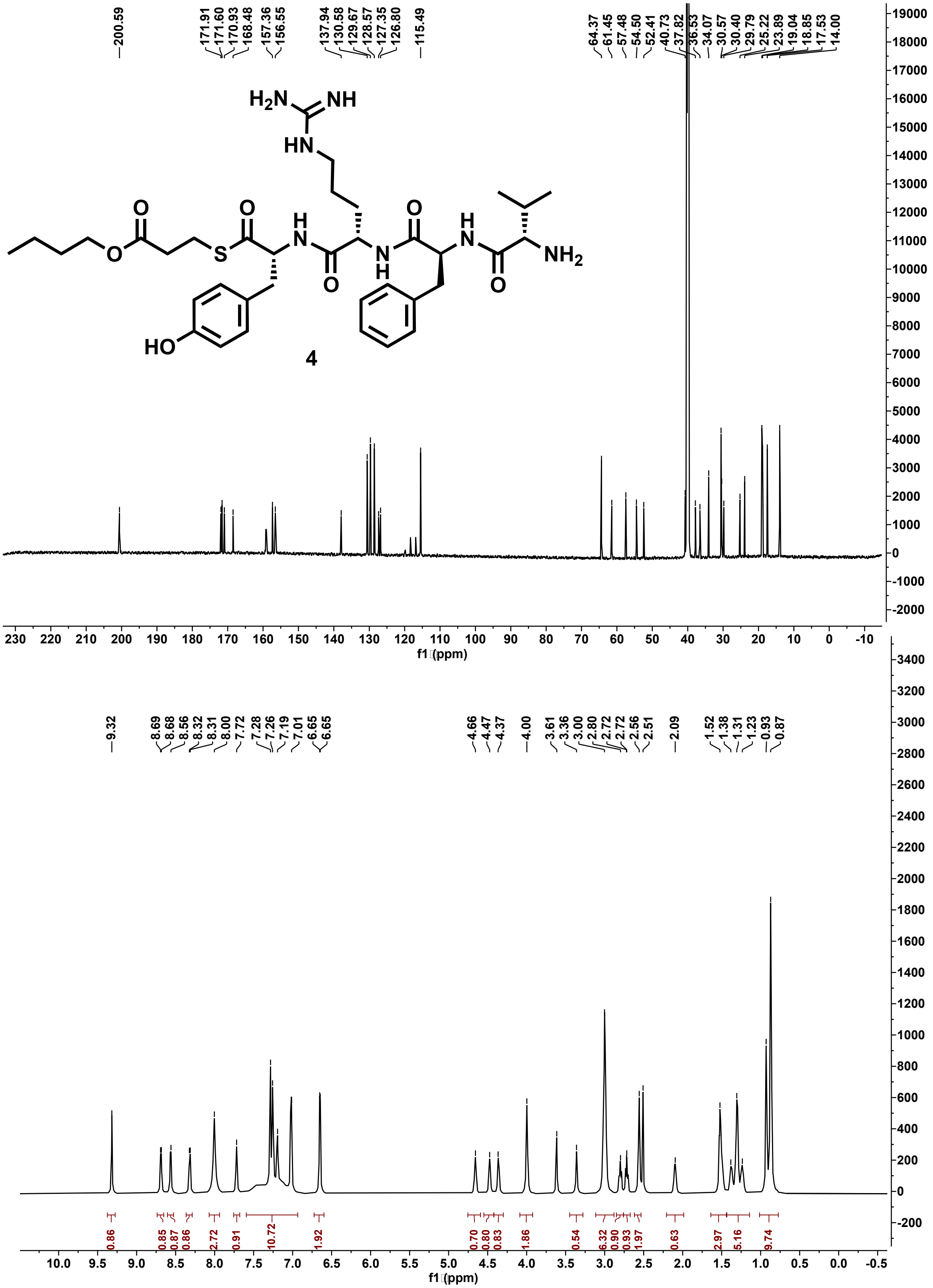

**Figure S113: $^{13}$C (201 MHz) and $^{1}$H (800 MHz) NMR spectra of 4 in DMSO-d$_6$.**

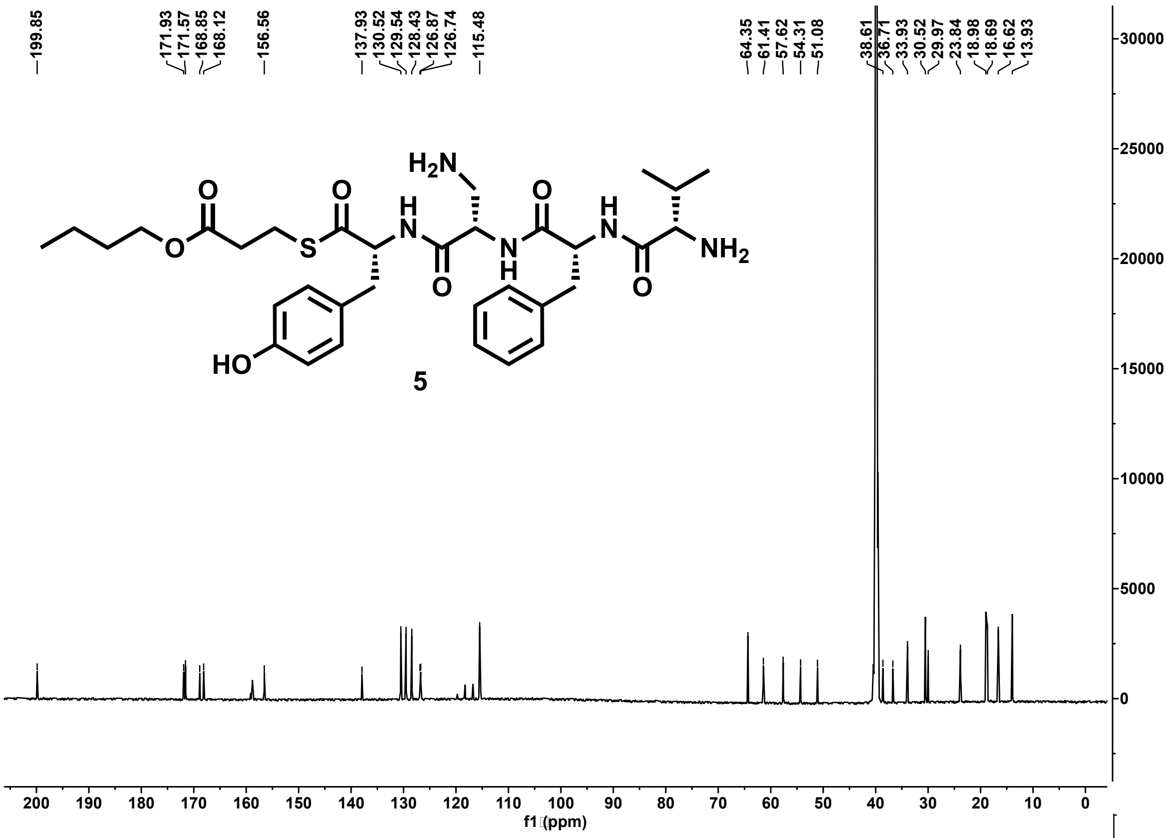

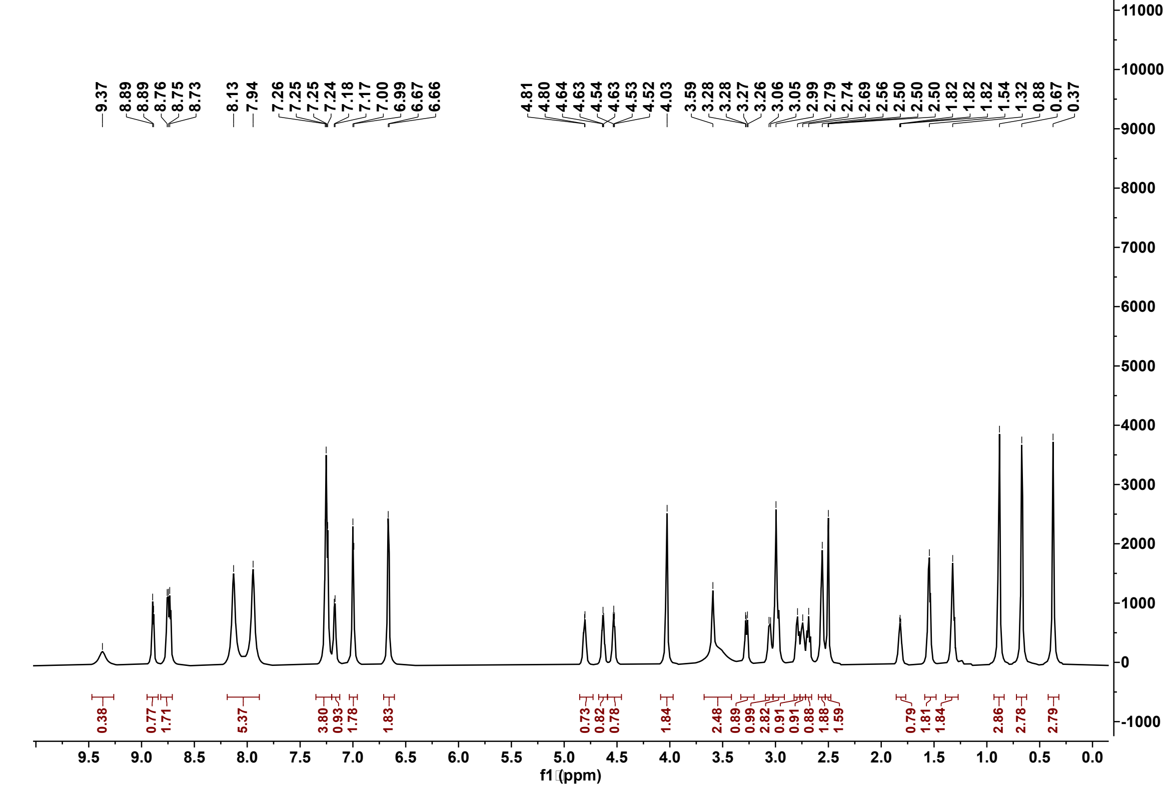

**Figure S114: $^{13}C$ (201 MHz) and $^{1}H$ (800 MHz) NMR spectra of 5 in DMSO-d$_6$.**

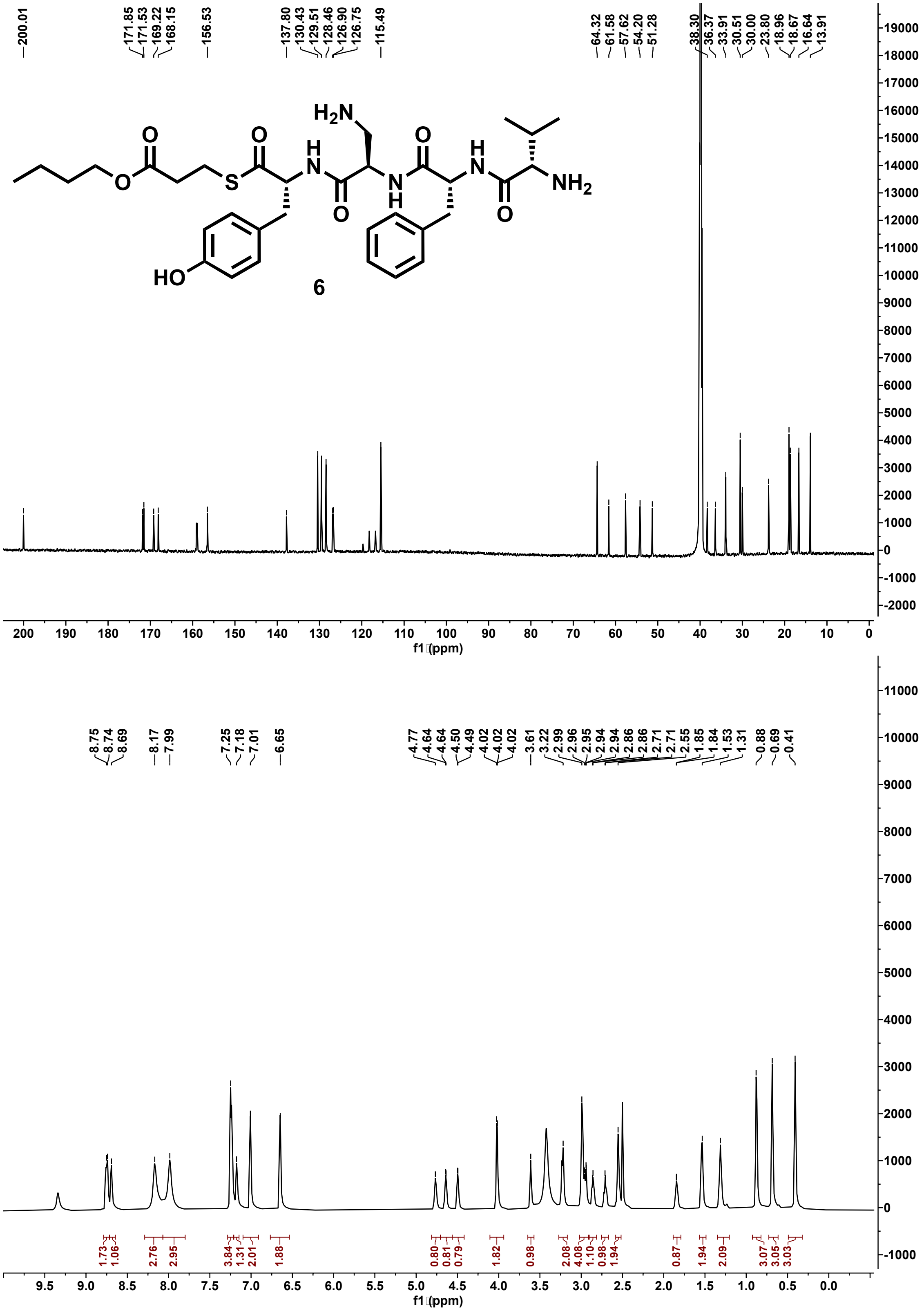

**Figure S115:** $^{13}$C (201 MHz) and $^{1}$H (800 MHz) NMR spectra of 6 in DMSO-$d_6$.

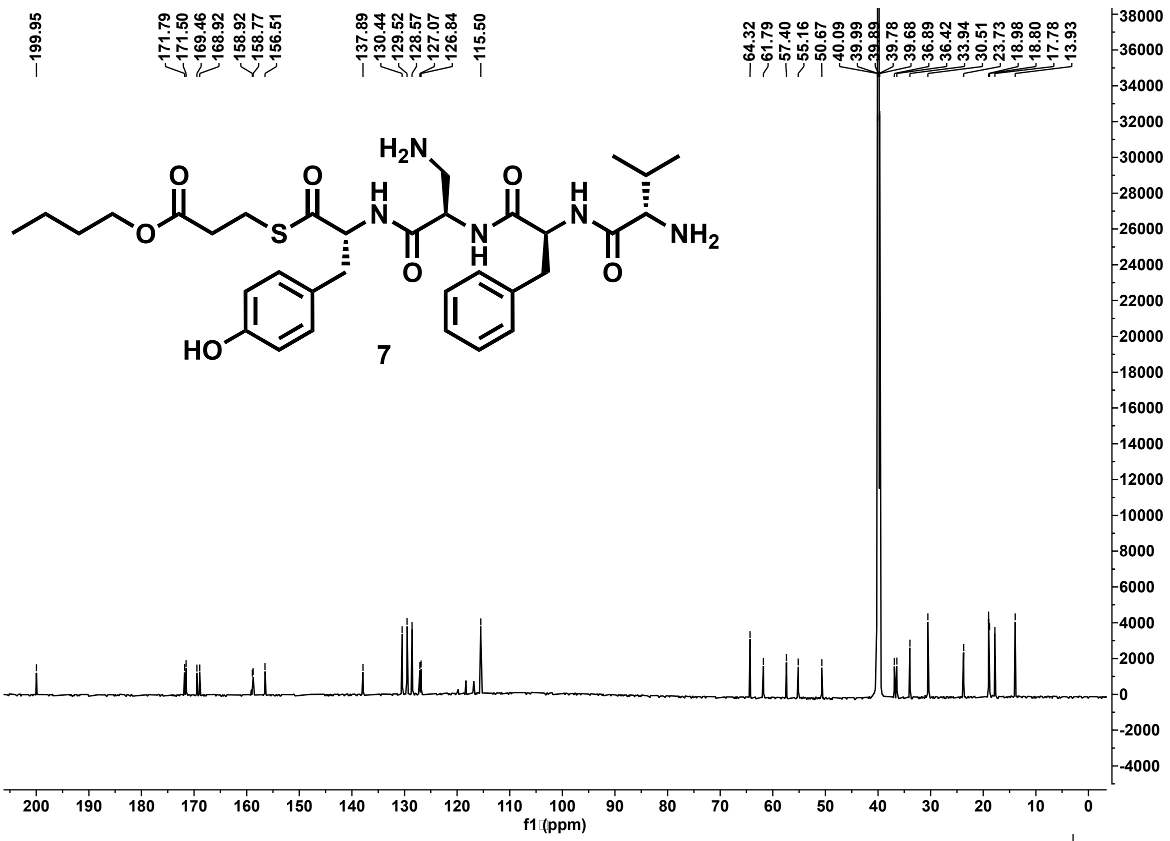

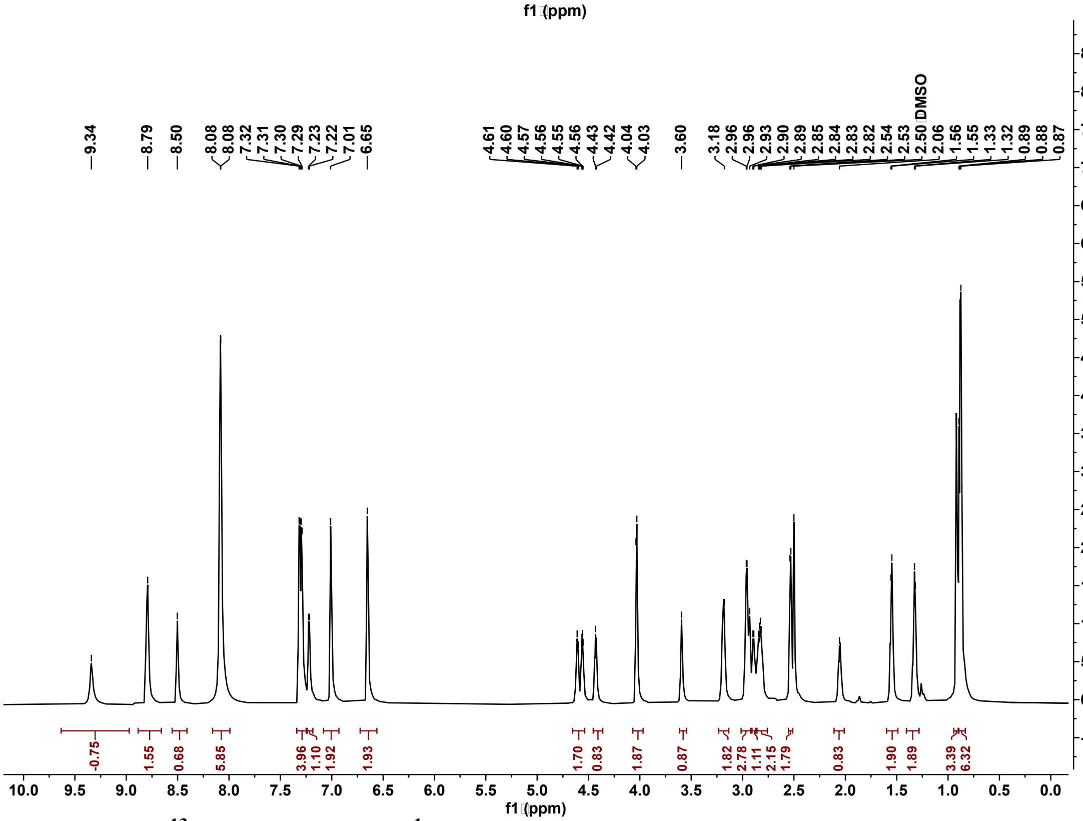

**Figure S116:** $^{13}C$ (201 MHz) and $^{1}H$ (800 MHz) NMR spectra of 7 in DMSO-$d_6$.

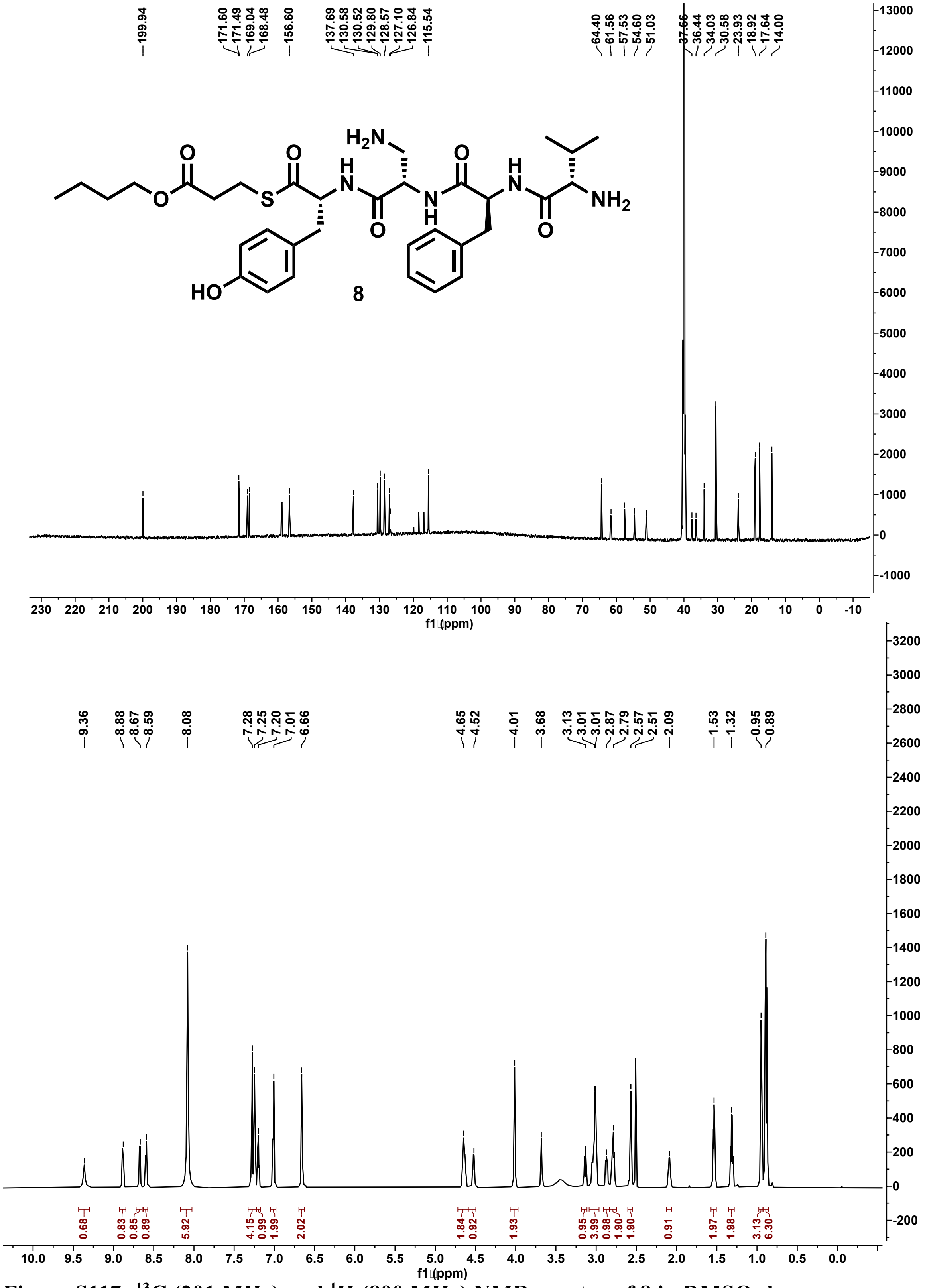

**Figure S117: $^{13}$C (201 MHz) and $^{1}$H (800 MHz) NMR spectra of 8 in DMSO-d$_6$.**

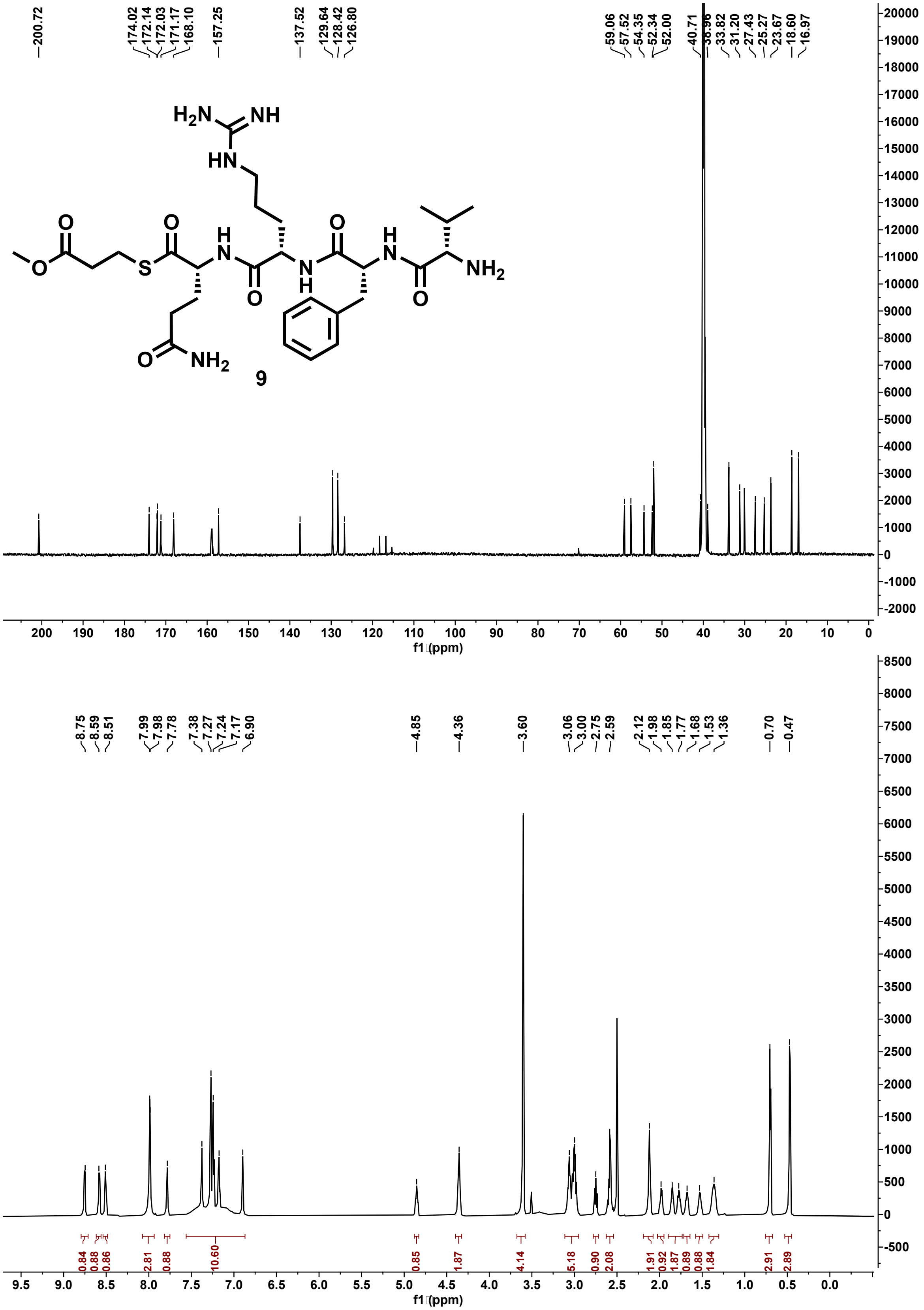

**Figure S118: $^{13}$C (201 MHz) and $^{1}$H (800 MHz) NMR spectra of 9 in DMSO-$d_6$.**

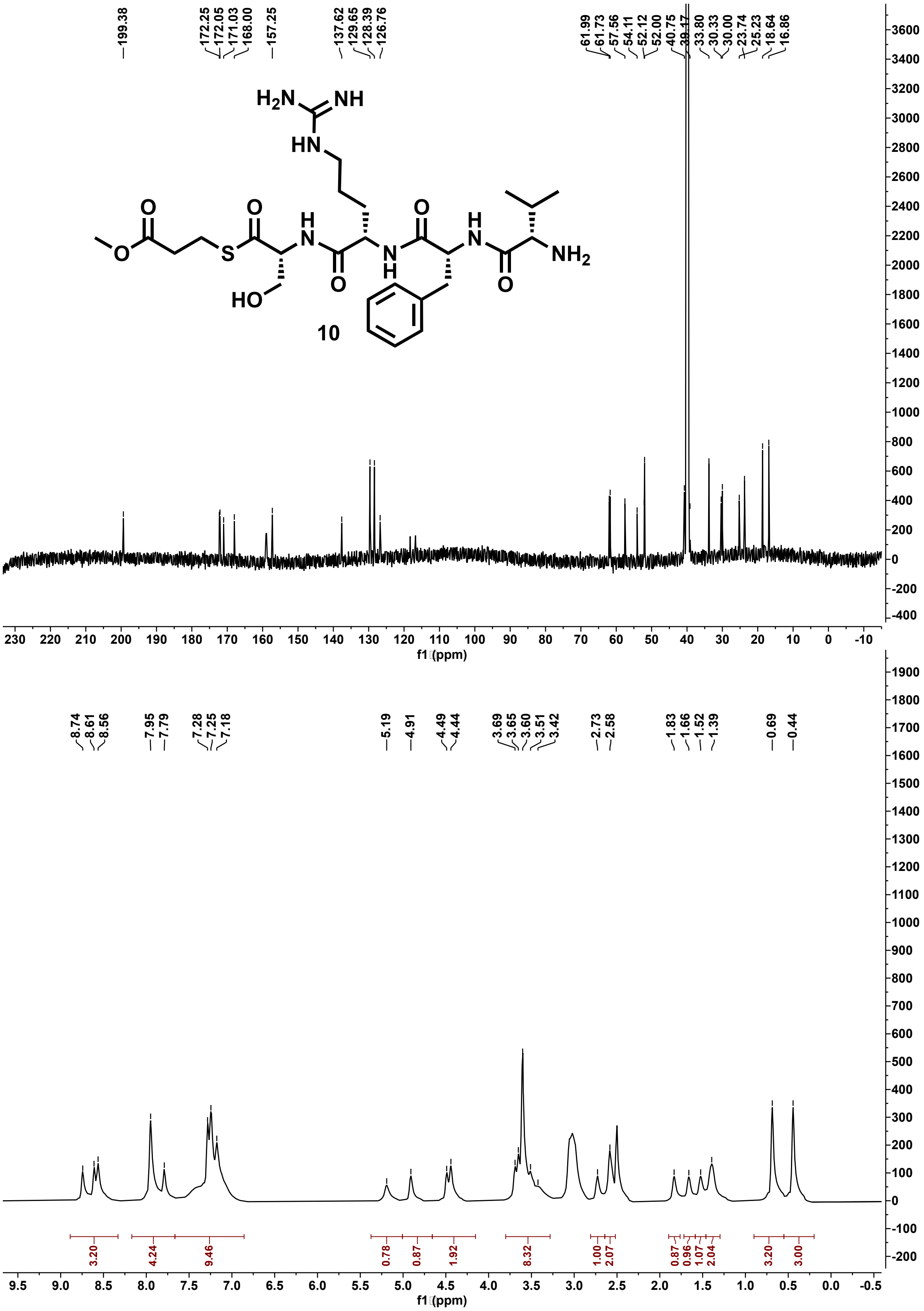

**Figure S119: $^{13}C$ (201 MHz) and $^{1}H$ (800 MHz) NMR spectra of 10 in DMSO-$d_6$.**

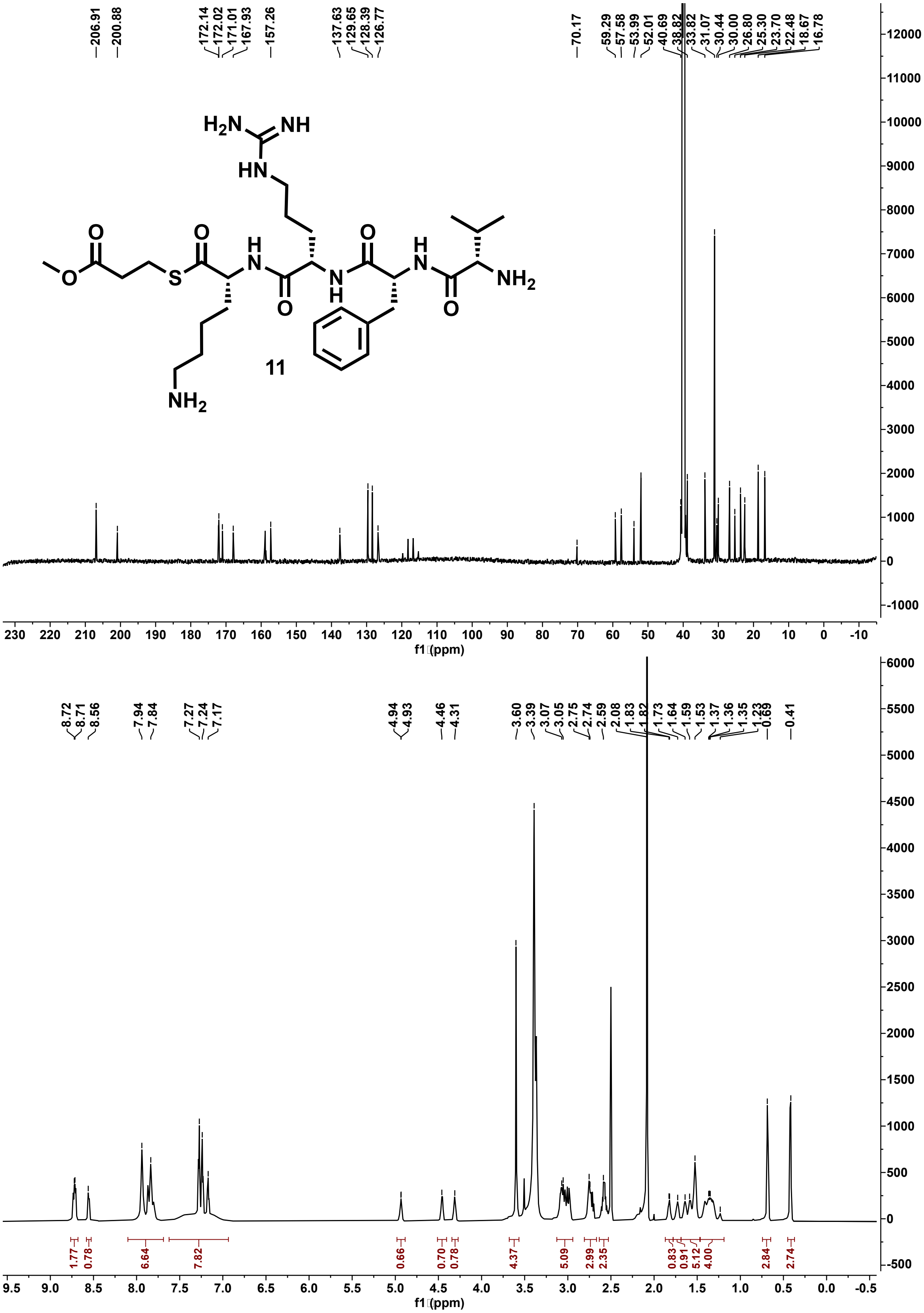

**Figure S120: $^{13}$C (201 MHz) and $^{1}$H (800 MHz) NMR spectra of 11 in DMSO-d$_6$.**

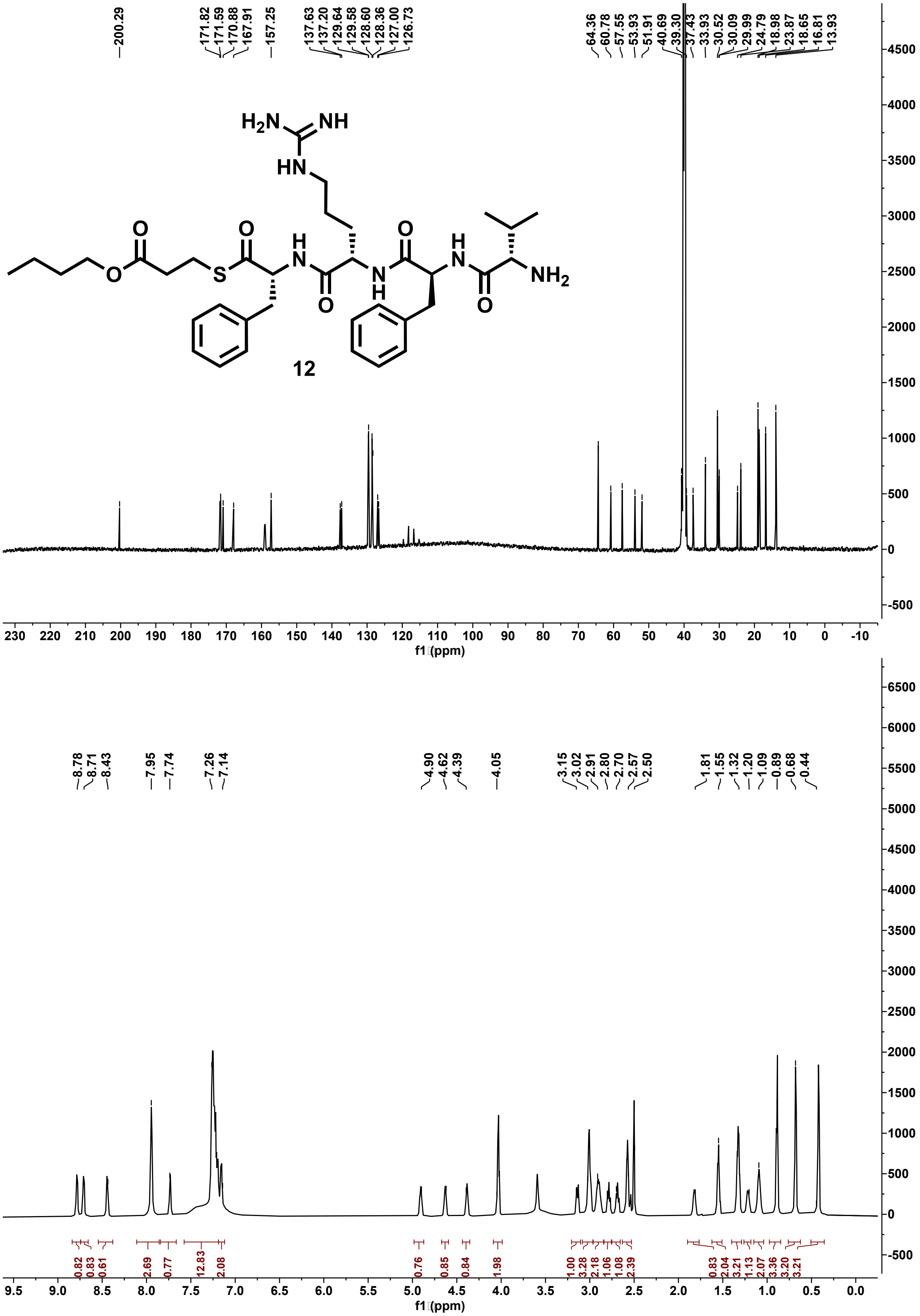

**Figure S121:** $^{13}$C (201 MHz) and $^{1}$H (800 MHz) NMR spectra of 12 in DMSO-$d_6$.

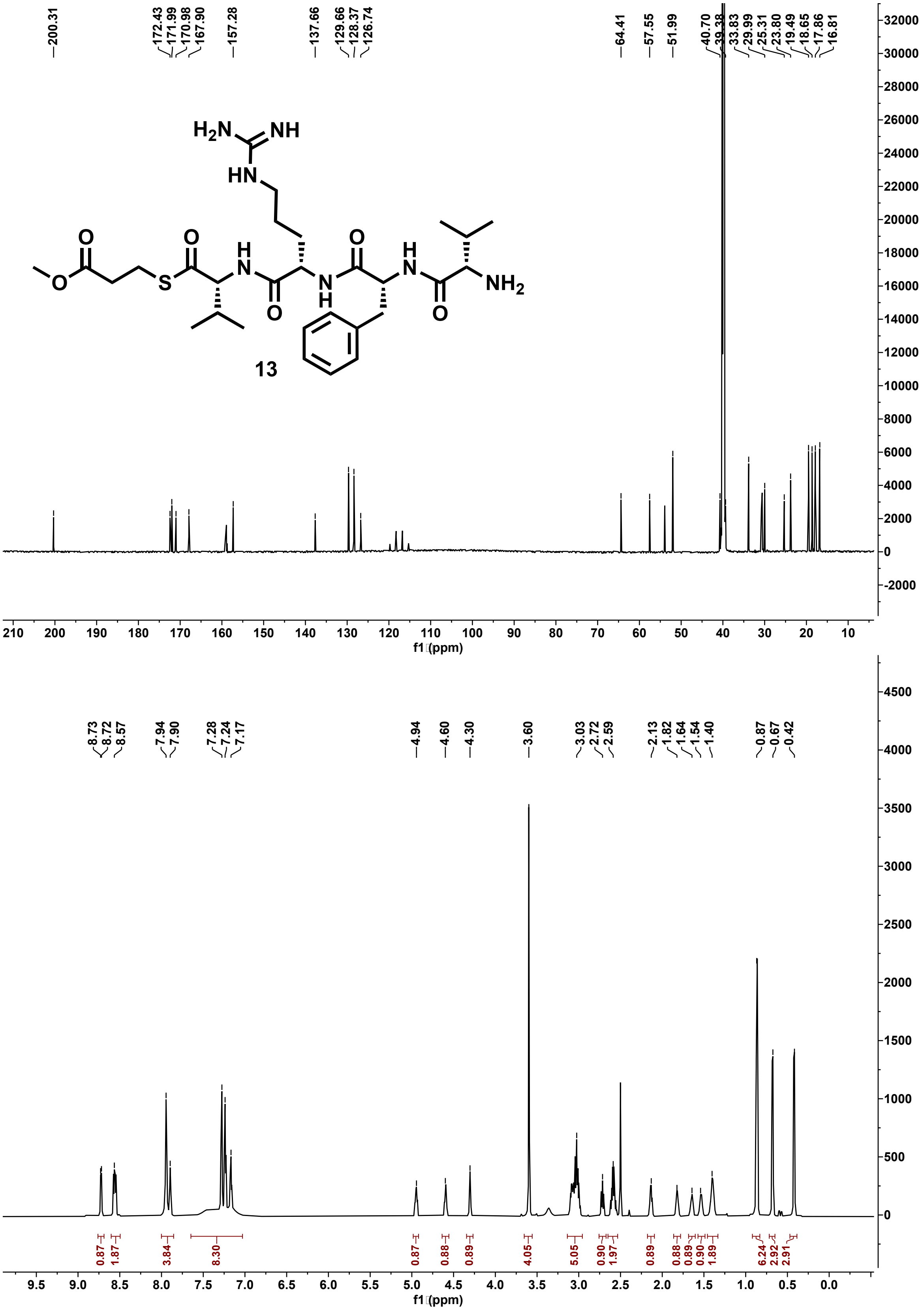

**Figure S122: $^{13}C$ (201 MHz) and $^{1}H$ (800 MHz) NMR spectra of 13 in DMSO-$d_6$.**

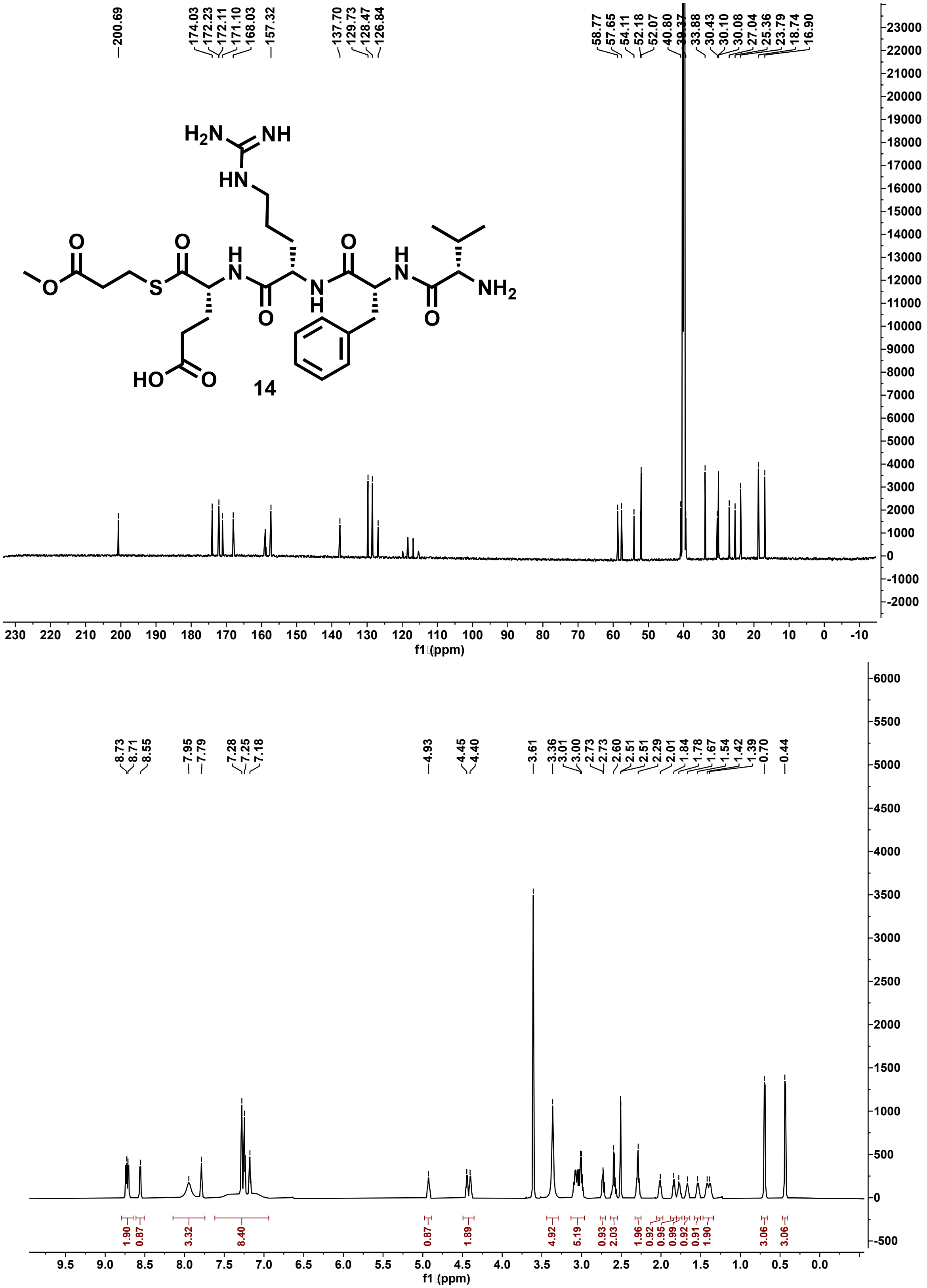

**Figure S123: $^{13}$C (201 MHz) and $^{1}$H (800 MHz) NMR spectra of 14 in DMSO-$d_6$.**

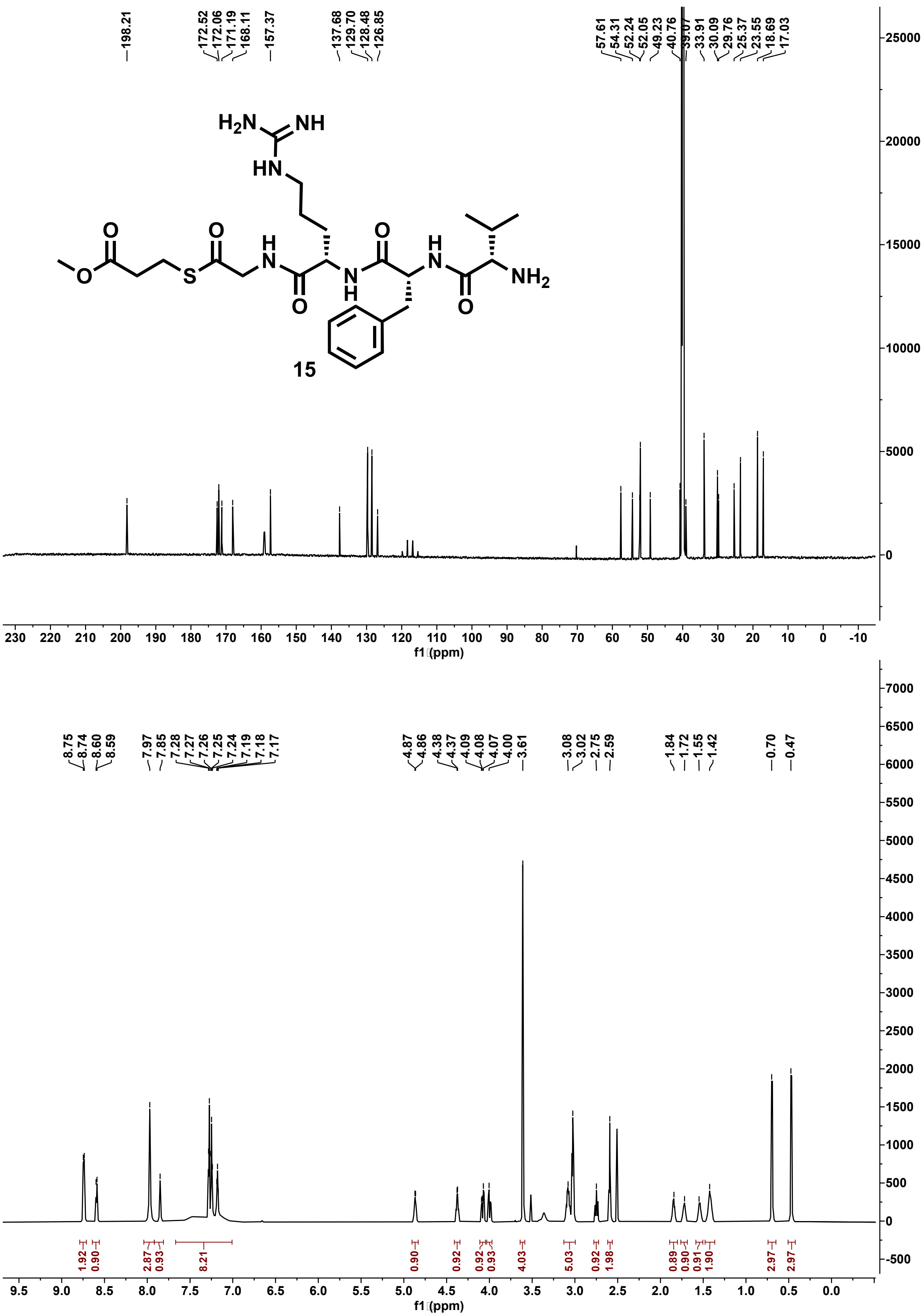

**Figure S124: $^{13}C$ (201 MHz) and $^{1}H$ (800 MHz) NMR spectra of 15 in DMSO-$d_6$.**

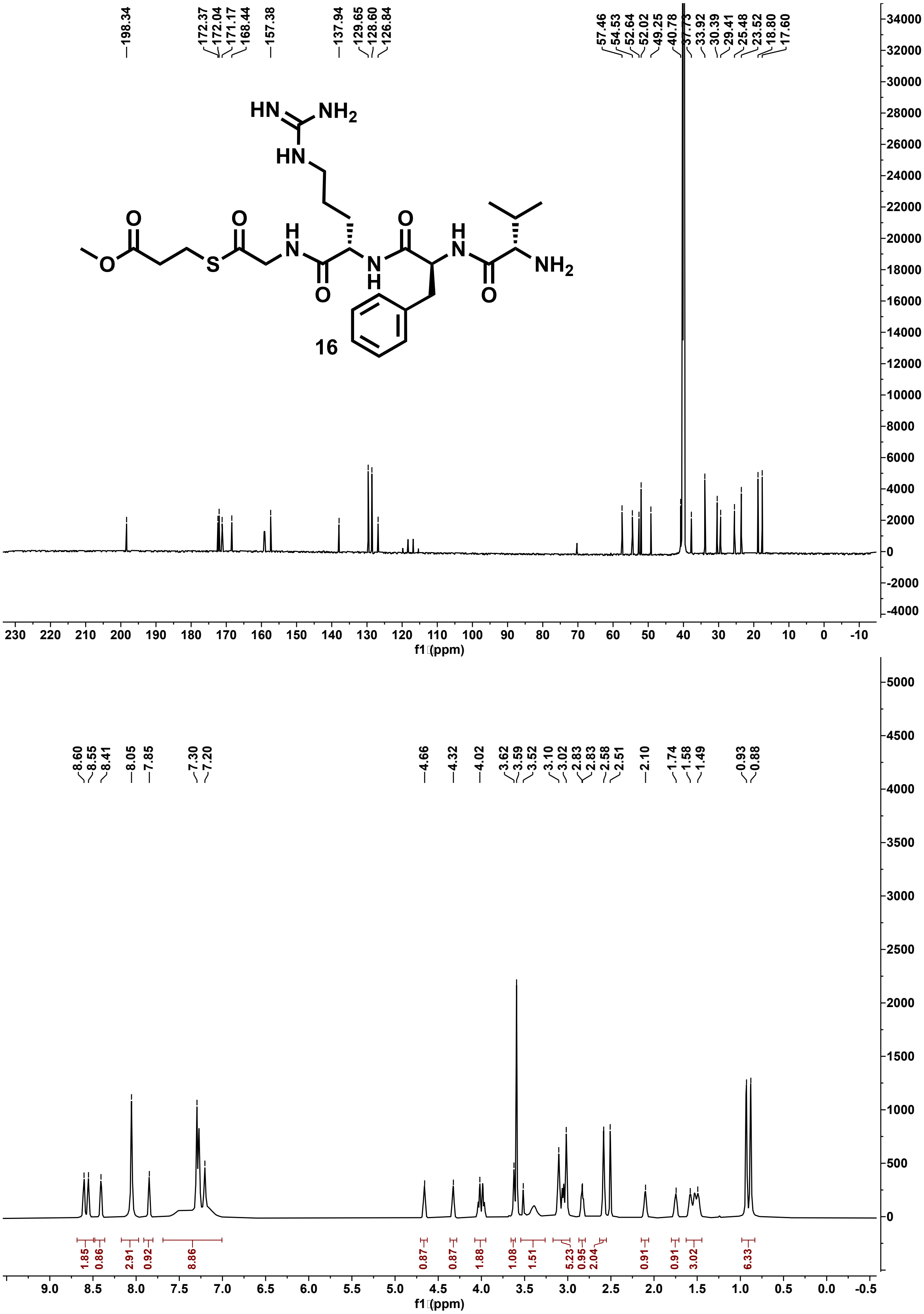

**Figure S125:** $^{13}C$ **(201 MHz) and** $^{1}H$ **(800 MHz) NMR spectra of 16 in DMSO-**$d_6$**.**

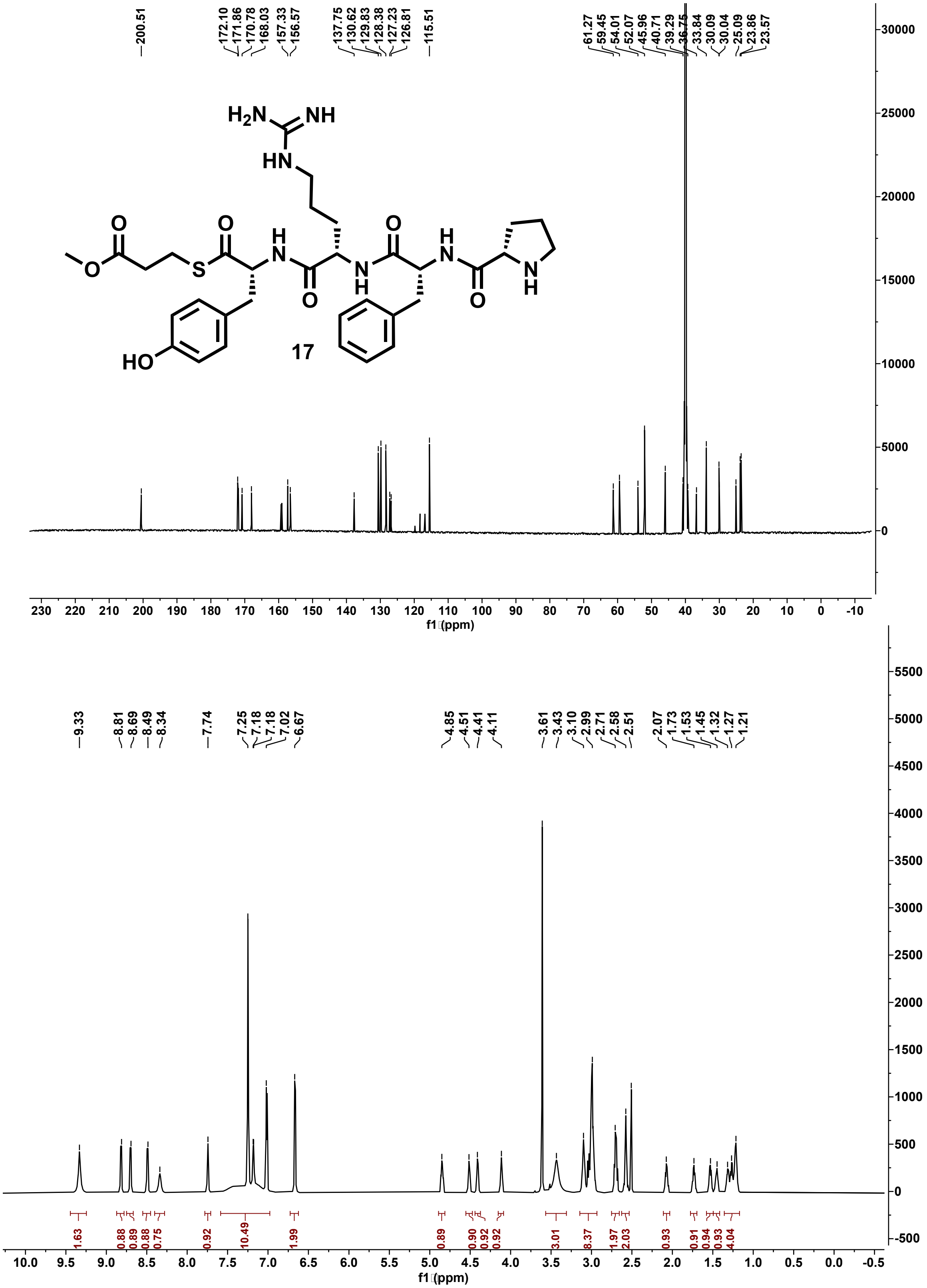

**Figure S126: $^{13}C$ (201 MHz) and $^{1}H$ (800 MHz) NMR spectra of 17 in DMSO-$d_6$.**

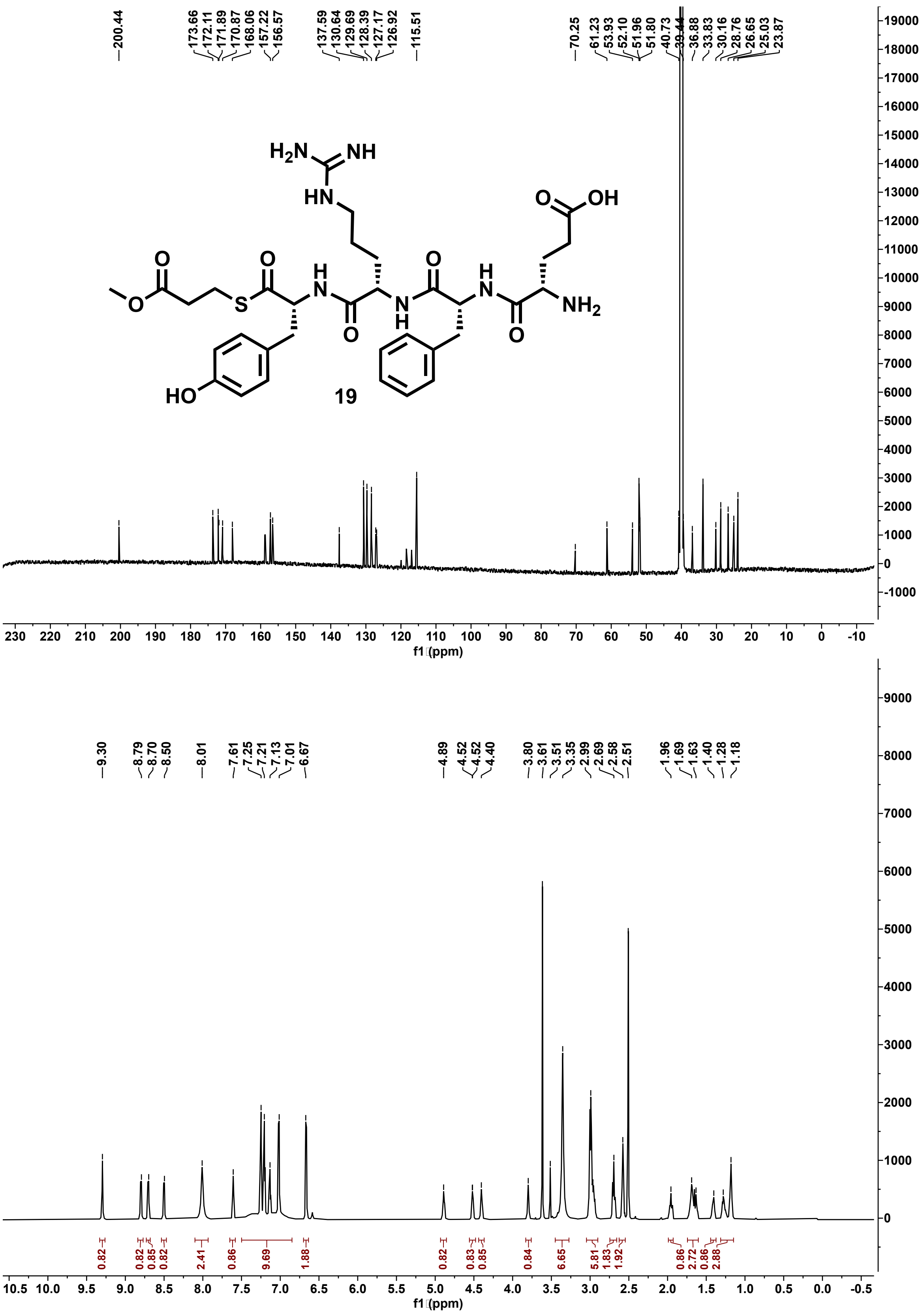

**Figure S127: $^{13}$C (201 MHz) and $^{1}$H (800 MHz) NMR spectra of 19 in DMSO-$d_6$.**

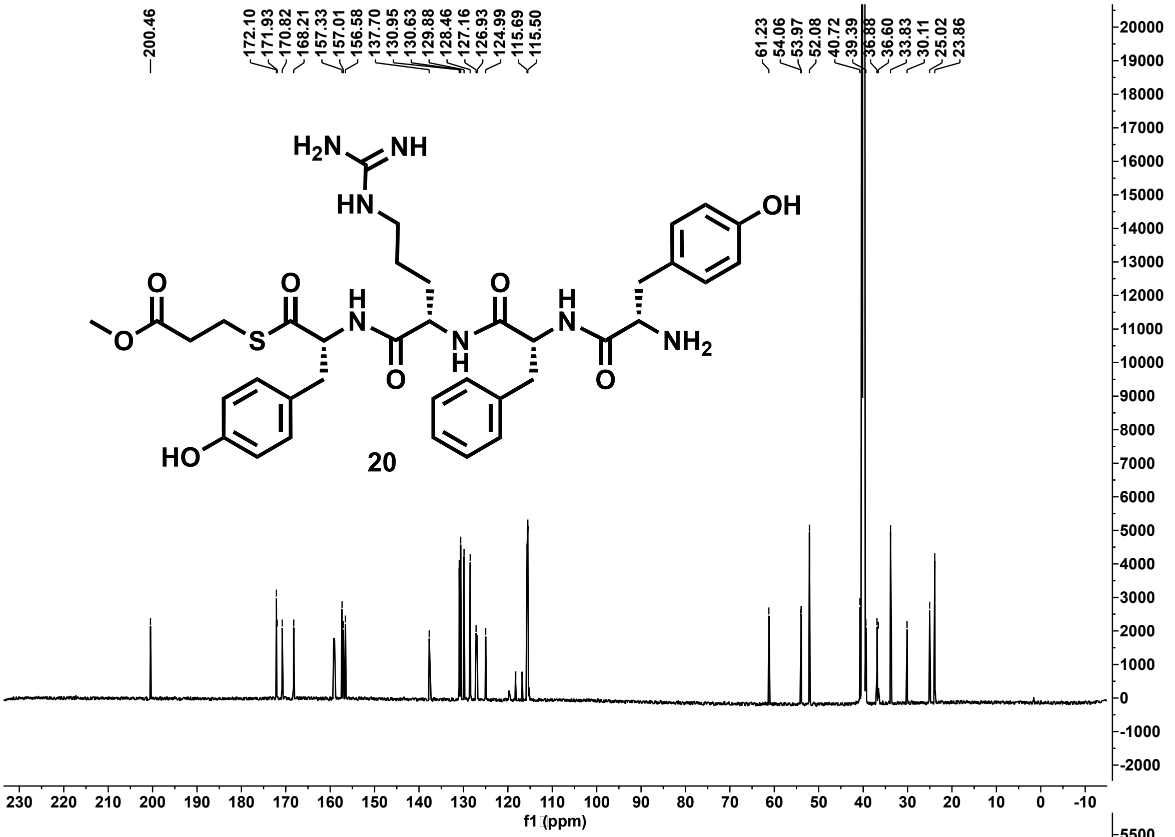

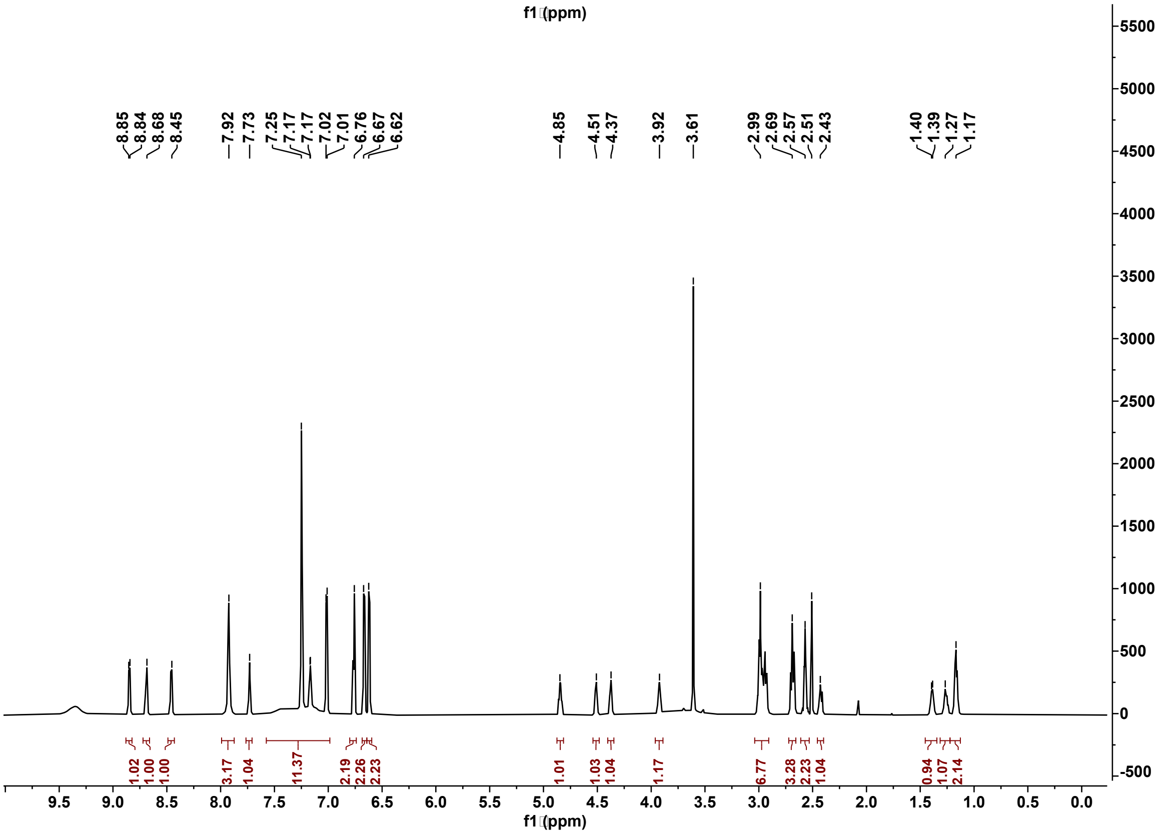

**Figure S128: $^{13}$C (201 MHz) and $^{1}$H (800 MHz) NMR spectra of 20 in DMSO-d$_6$.**

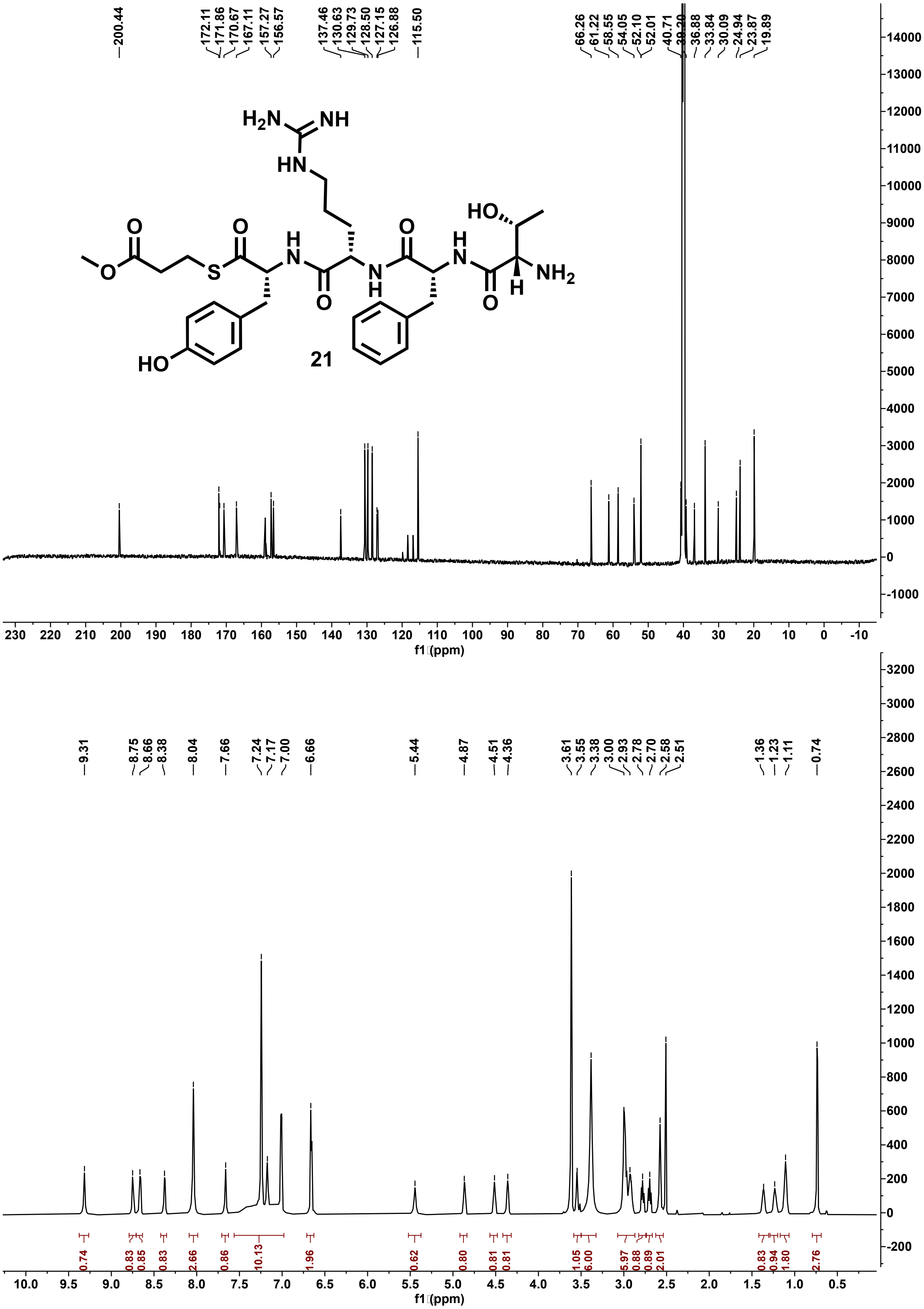

**Figure S129: $^{13}C$ (201 MHz) and $^{1}H$ (800 MHz) NMR spectra of 21 in DMSO-d$_6$.**

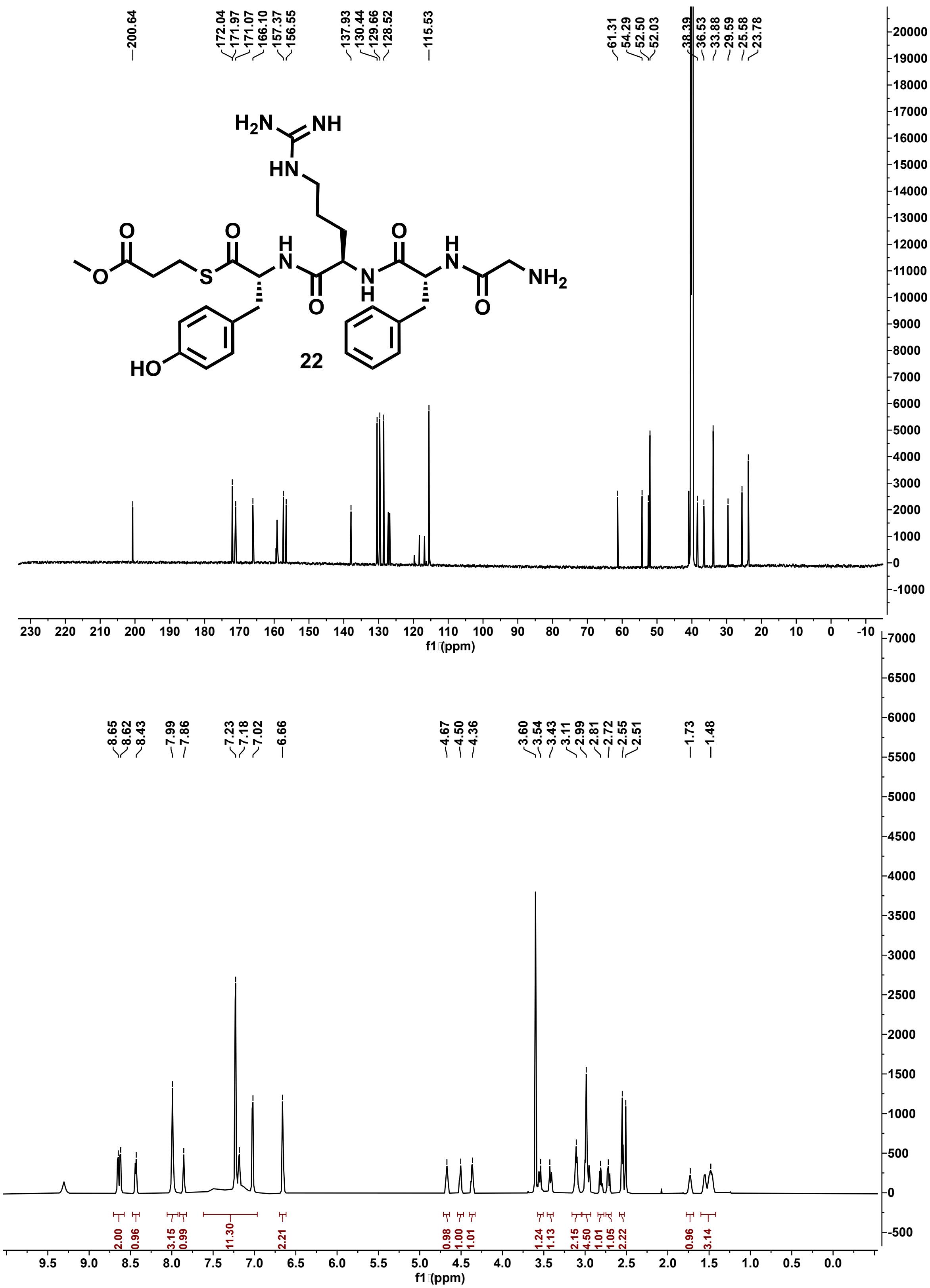

**Figure S130:** $^{13}C$ (201 MHz) and $^{1}H$ (800 MHz) NMR spectra of 22 in DMSO-$d_6$.

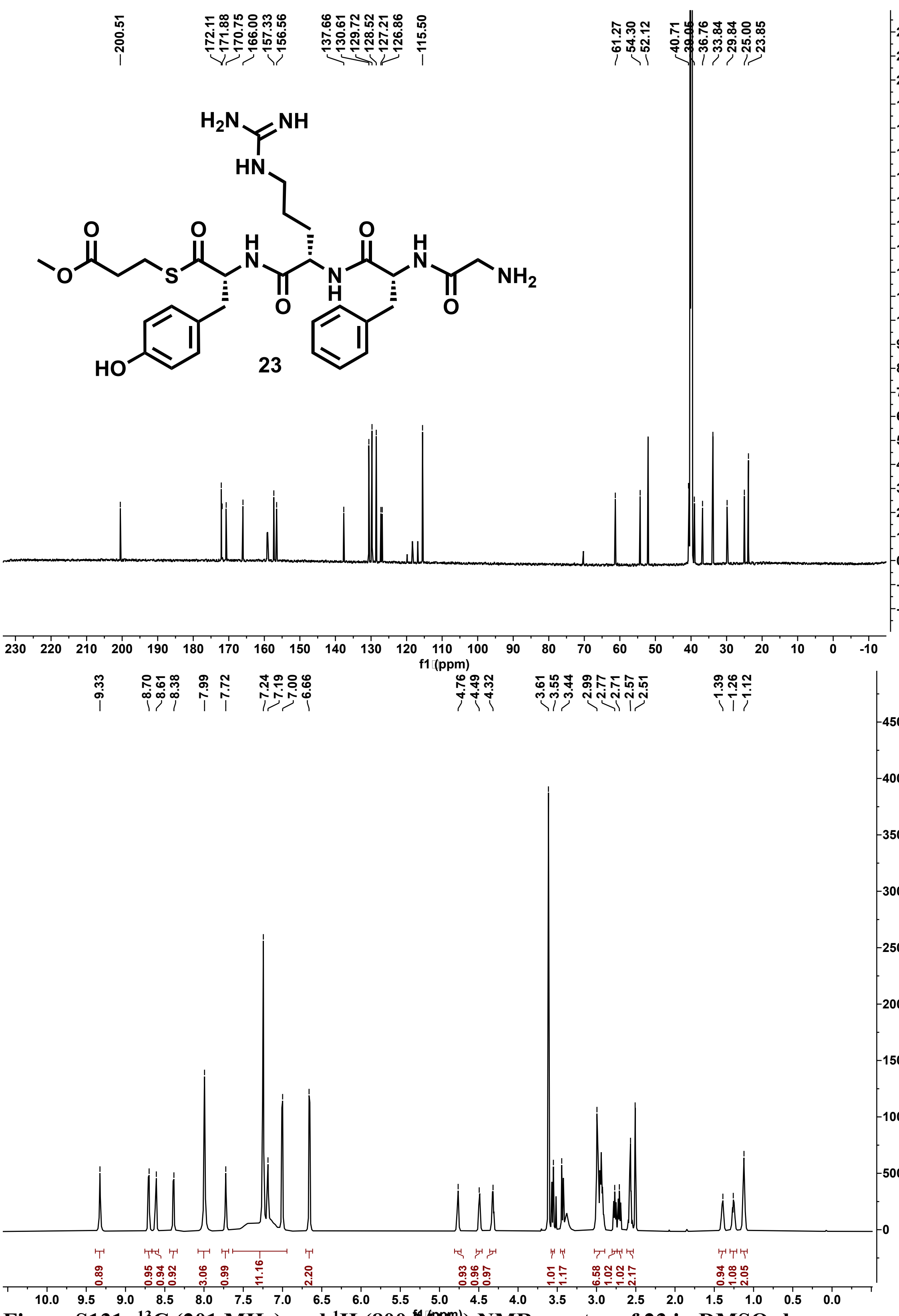

**Figure S131: $^{13}$C (201 MHz) and $^{1}$H (800 MHz) NMR spectra of 23 in DMSO-d$_6$.**

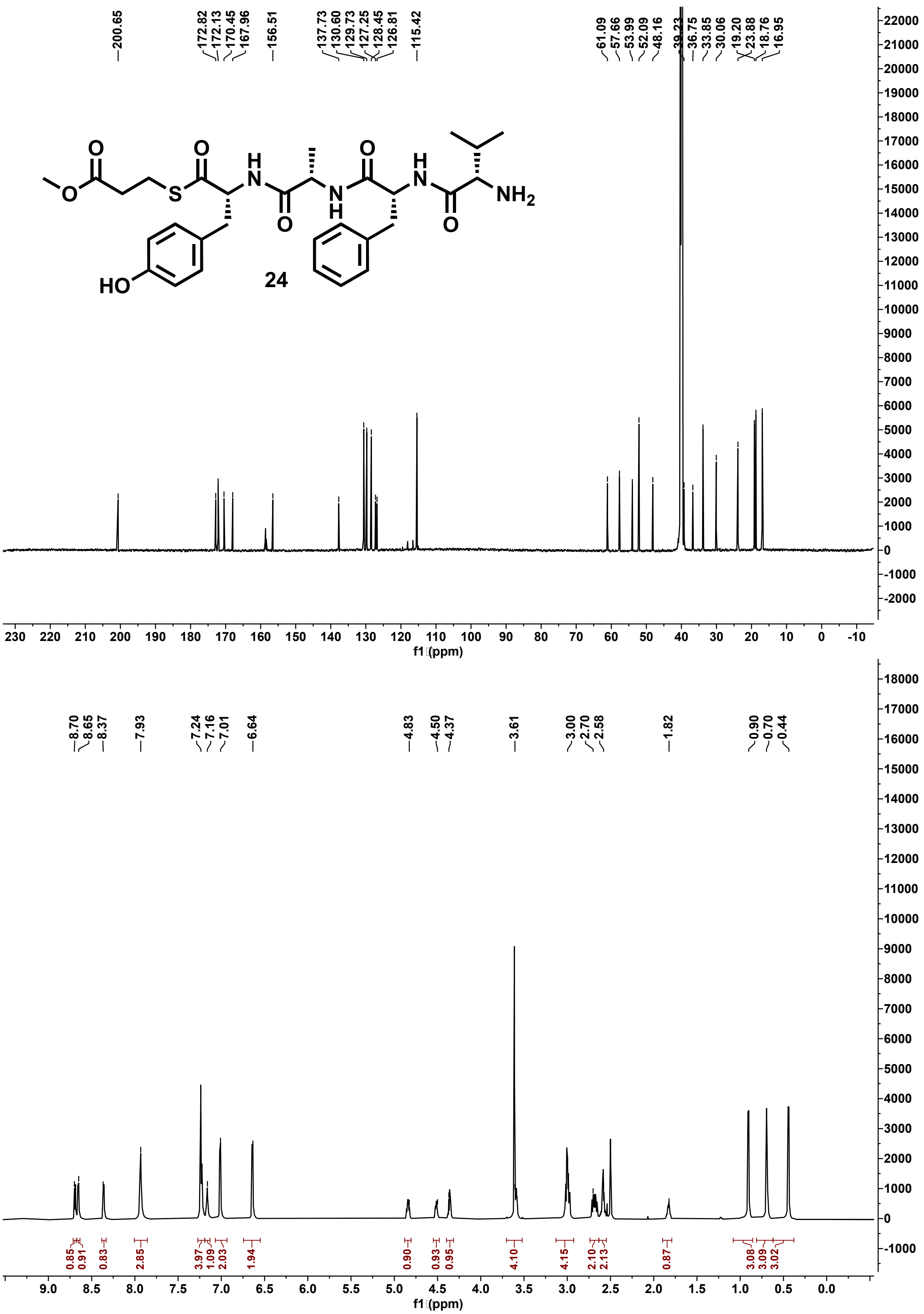

**Figure S132: $^{13}C$ (201 MHz) and $^{1}H$ (800 MHz) NMR spectra of 24 in DMSO-$d_6$.**

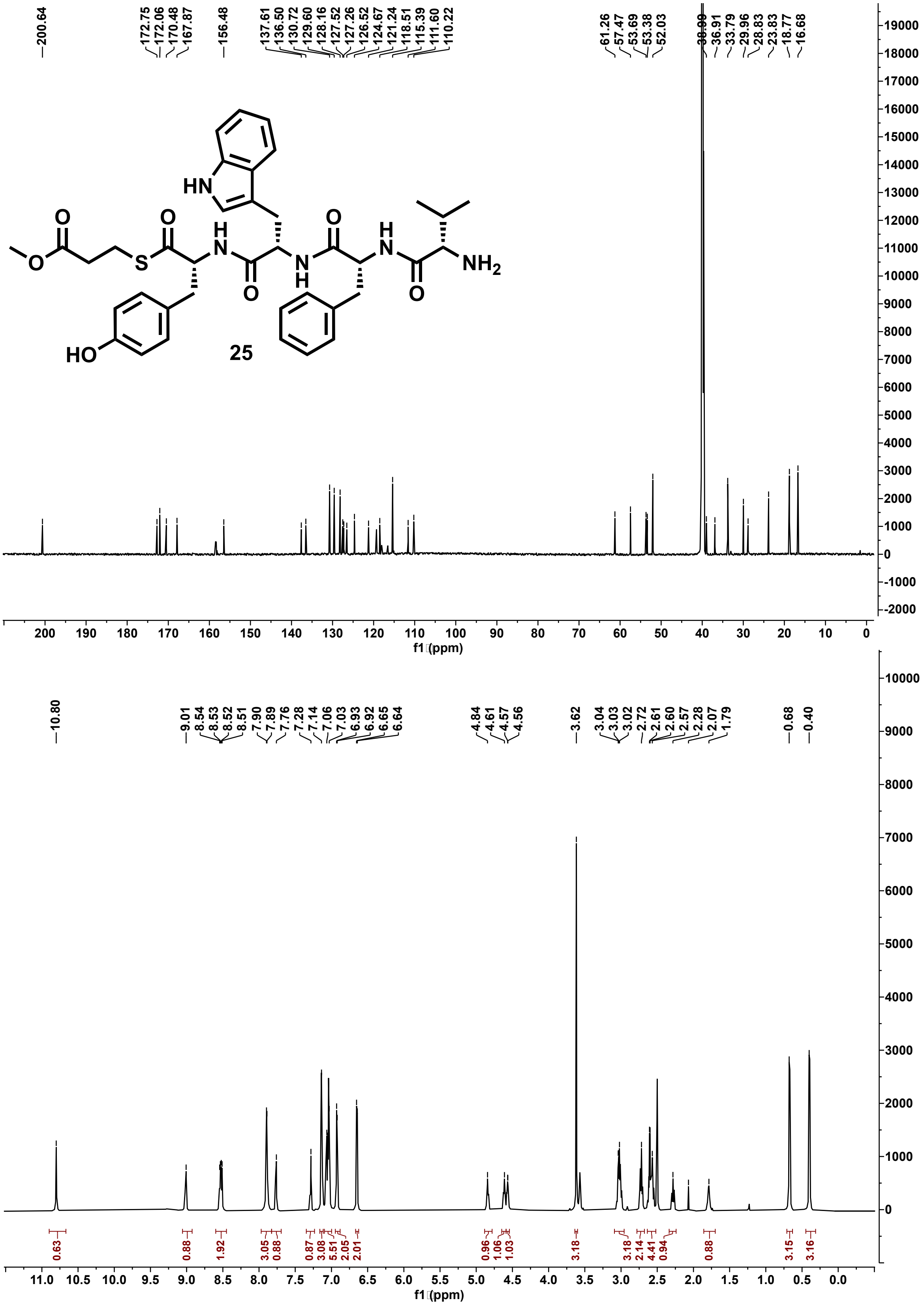

**Figure S133: $^{13}C$ (201 MHz) and $^{1}H$ (800 MHz) NMR spectra of 25 in DMSO-$d_6$.**

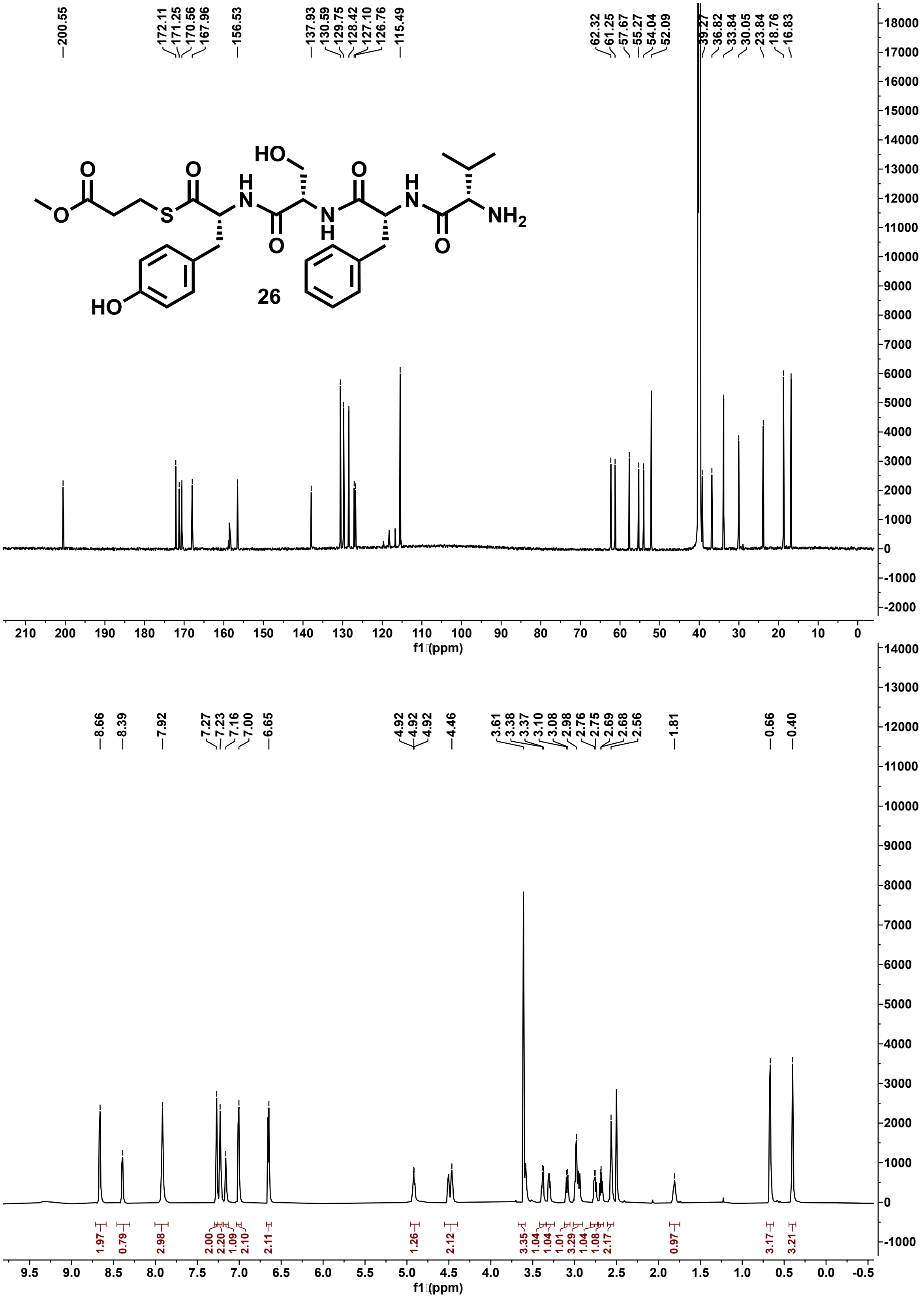

**Figure S134: $^{13}$C (201 MHz) and $^{1}$H (800 MHz) NMR spectra of 26 in DMSO-$d_6$.**

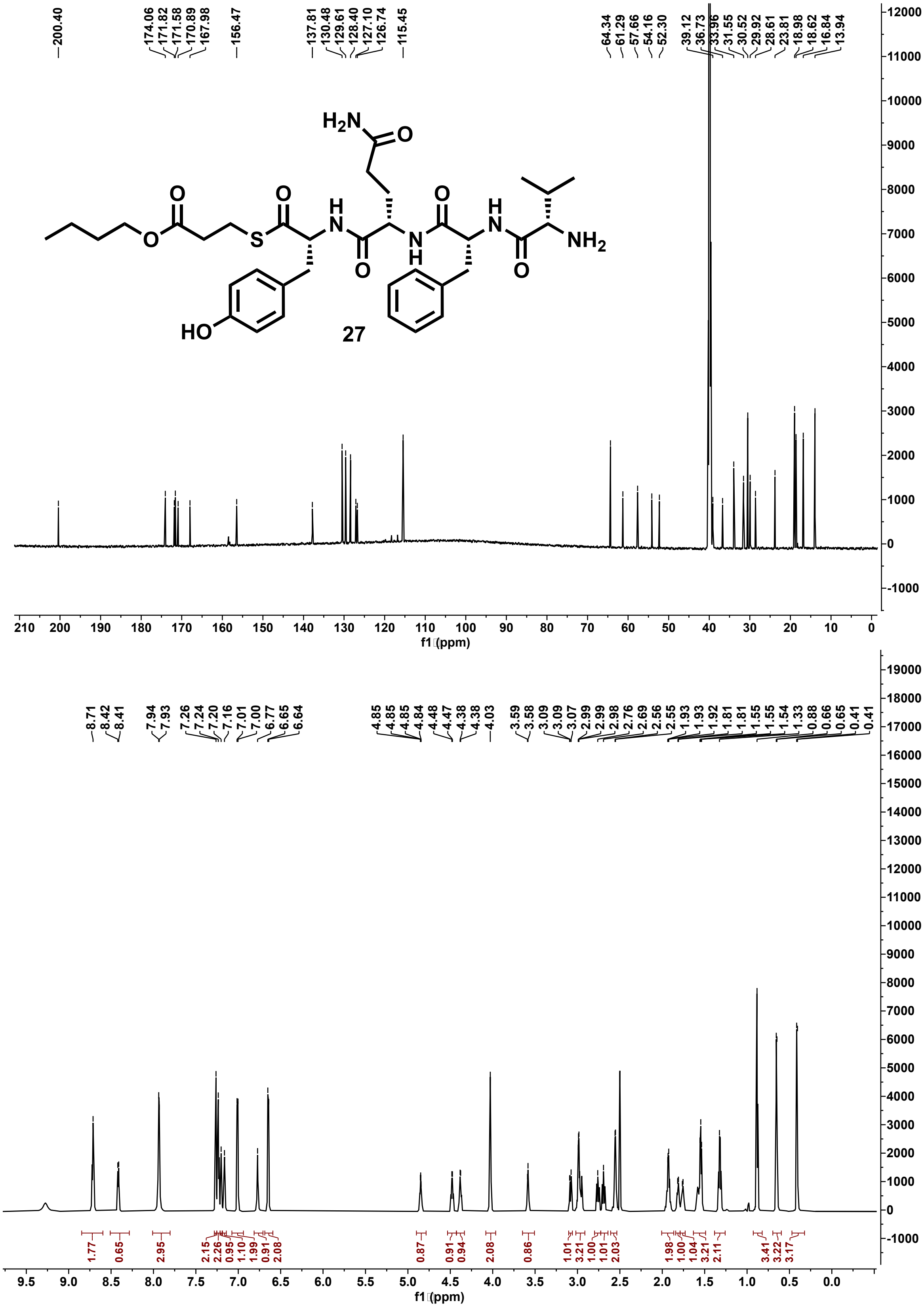

**Figure S135: $^{13}$C (201 MHz) and $^{1}$H (800 MHz) NMR spectra of 27 in DMSO-$d_6$.**

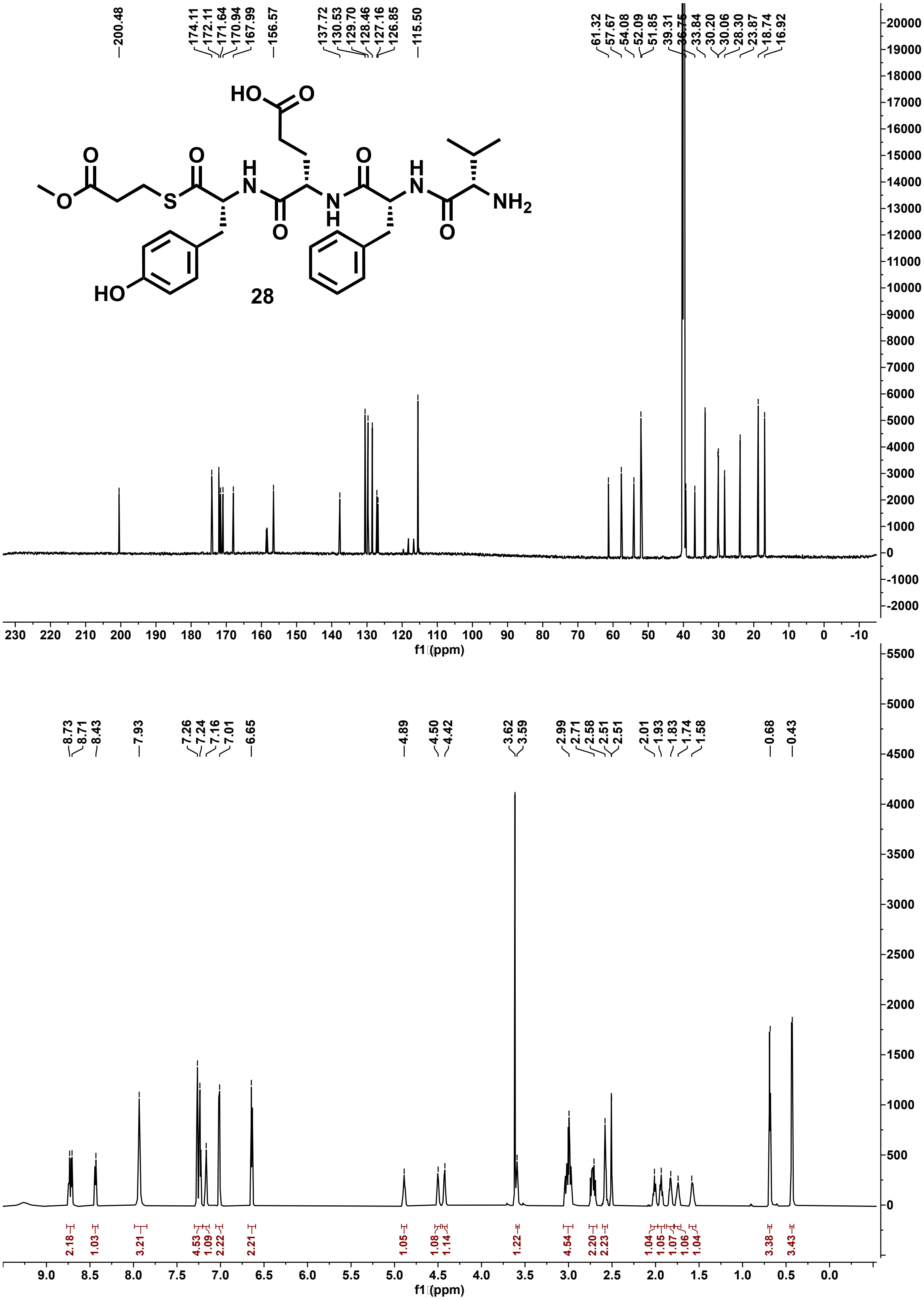

**Figure S136: $^{13}C$ (201 MHz) and $^{1}H$ (800 MHz) NMR spectra of 28 in DMSO-$d_6$.**

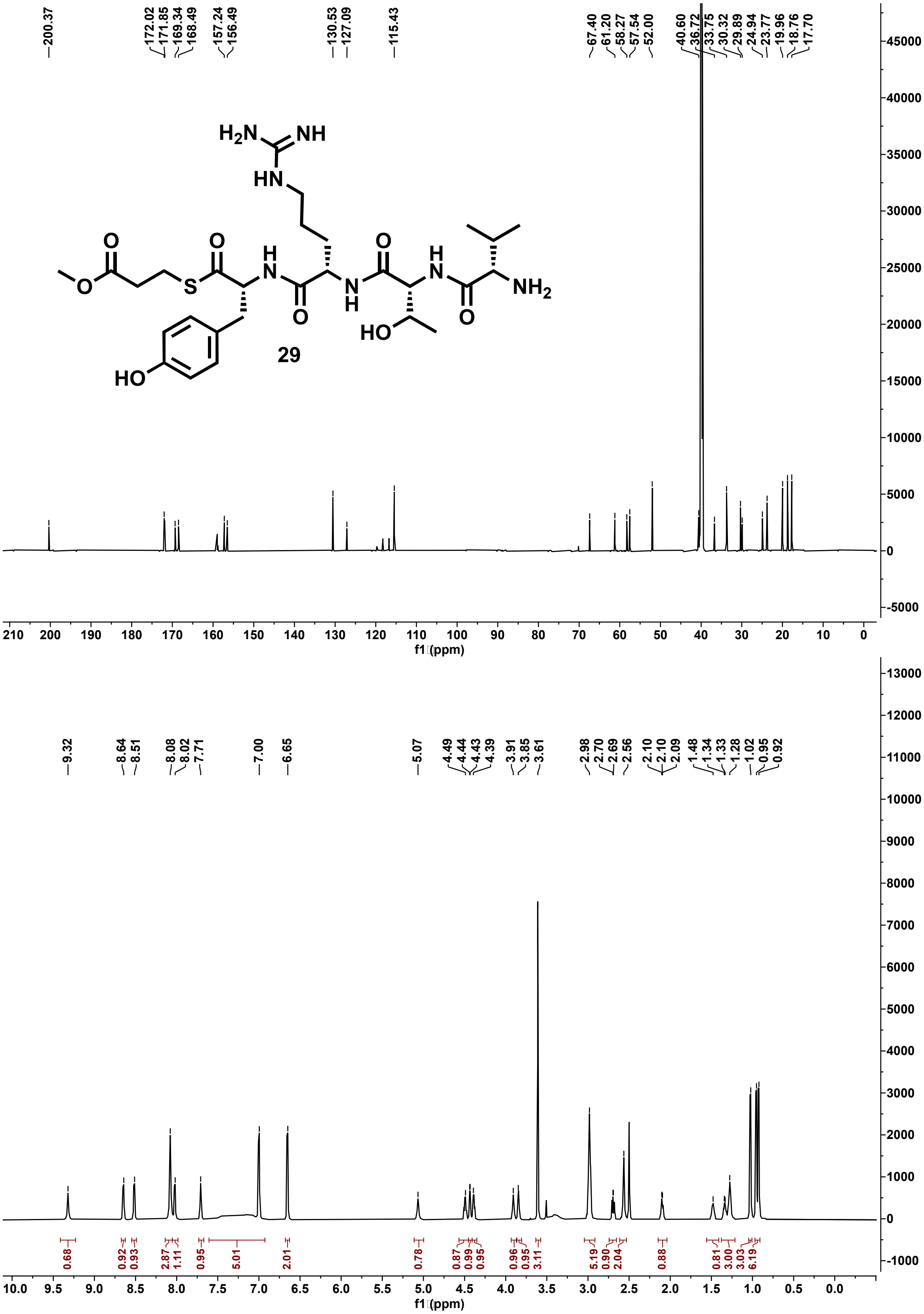

**Figure S137:** $^{13}C$ (201 MHz) and $^{1}H$ (800 MHz) NMR spectra of 29 in DMSO-$d_6$.

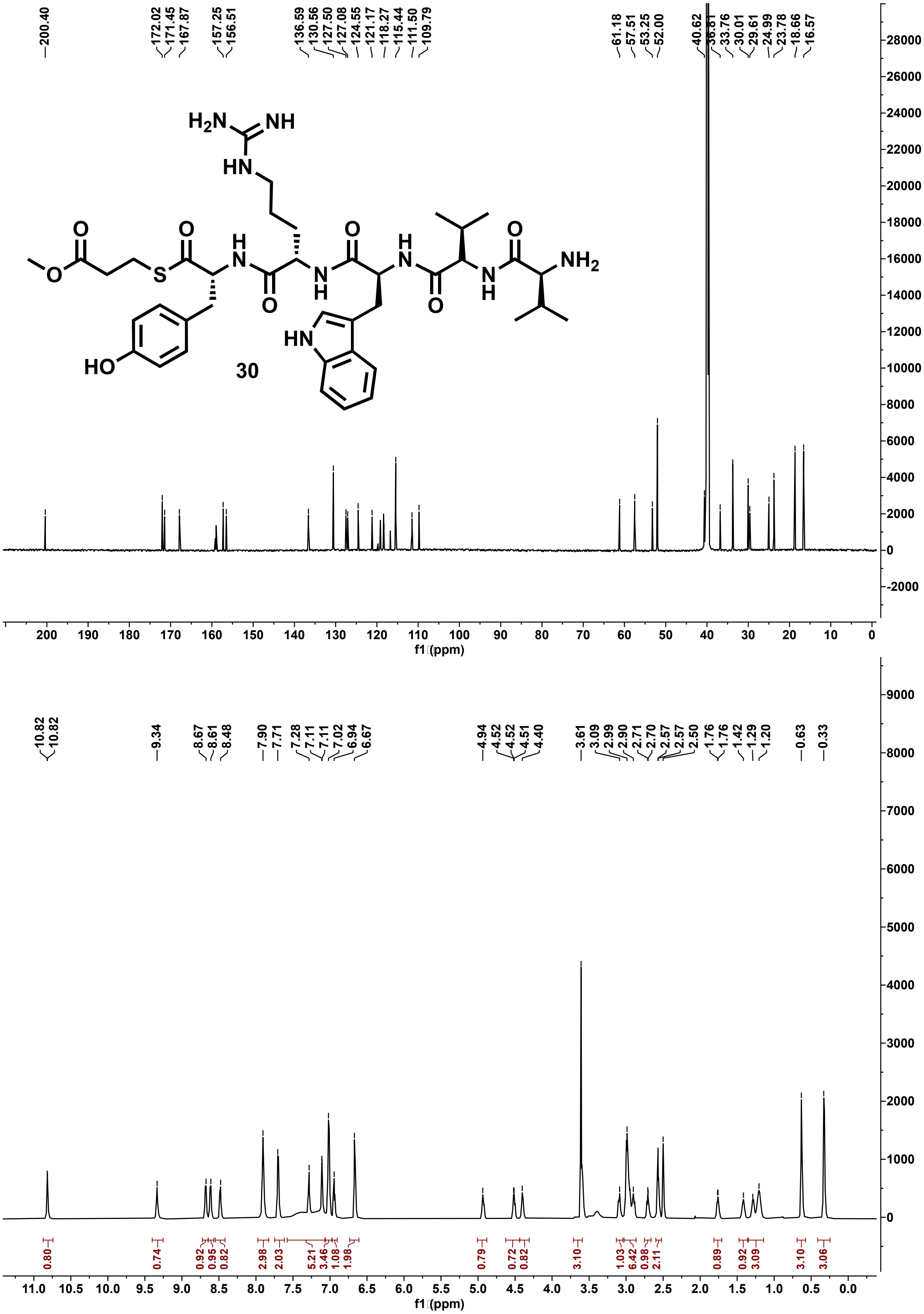

**Figure S138: $^{13}$C (201 MHz) and $^{1}$H (800 MHz) NMR spectra of 30 in DMSO-$d_6$.**

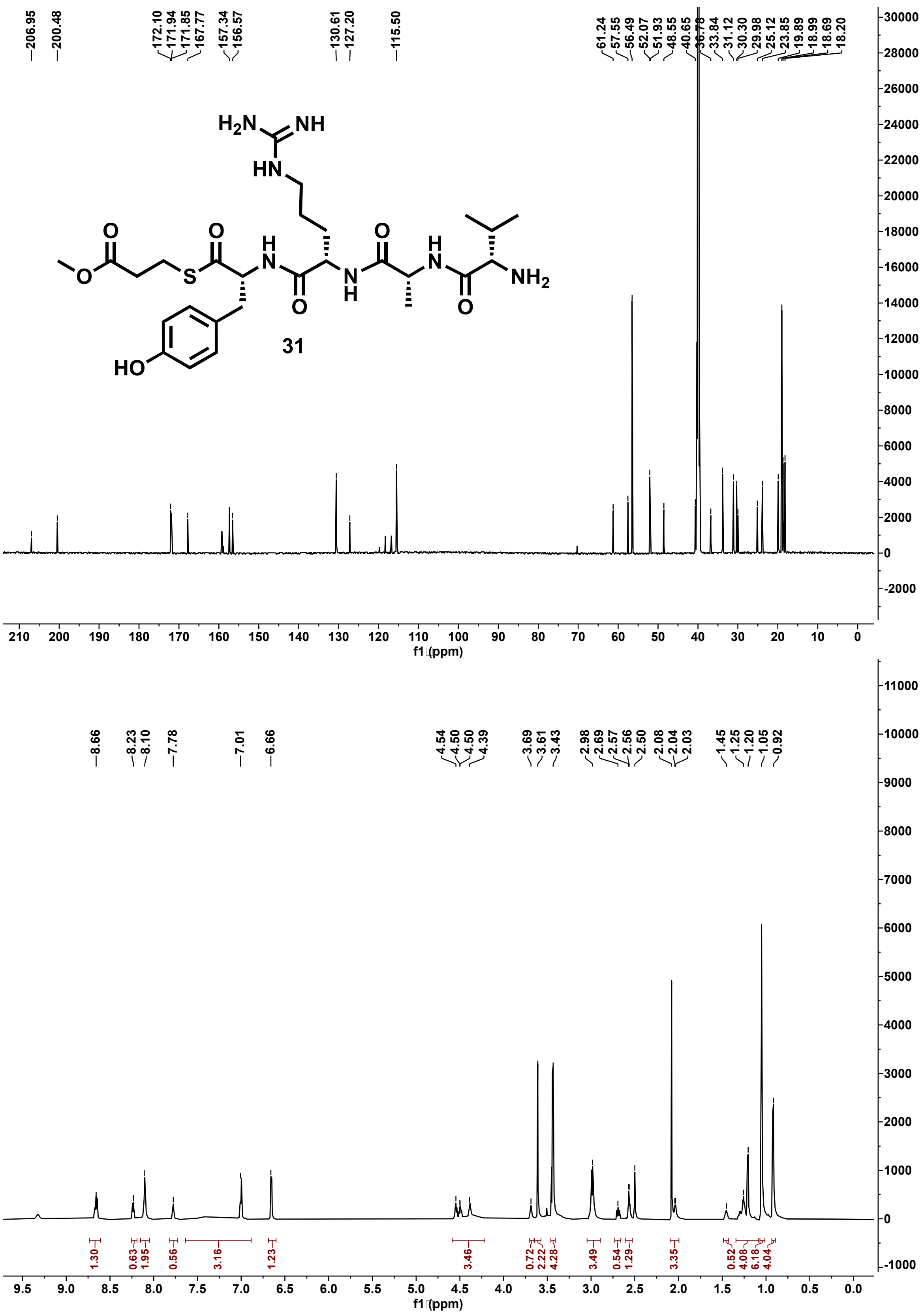

**Figure S139: $^{13}C$ (201 MHz) and $^{1}H$ (800 MHz) NMR spectra of 31 in DMSO-$d_6$.**

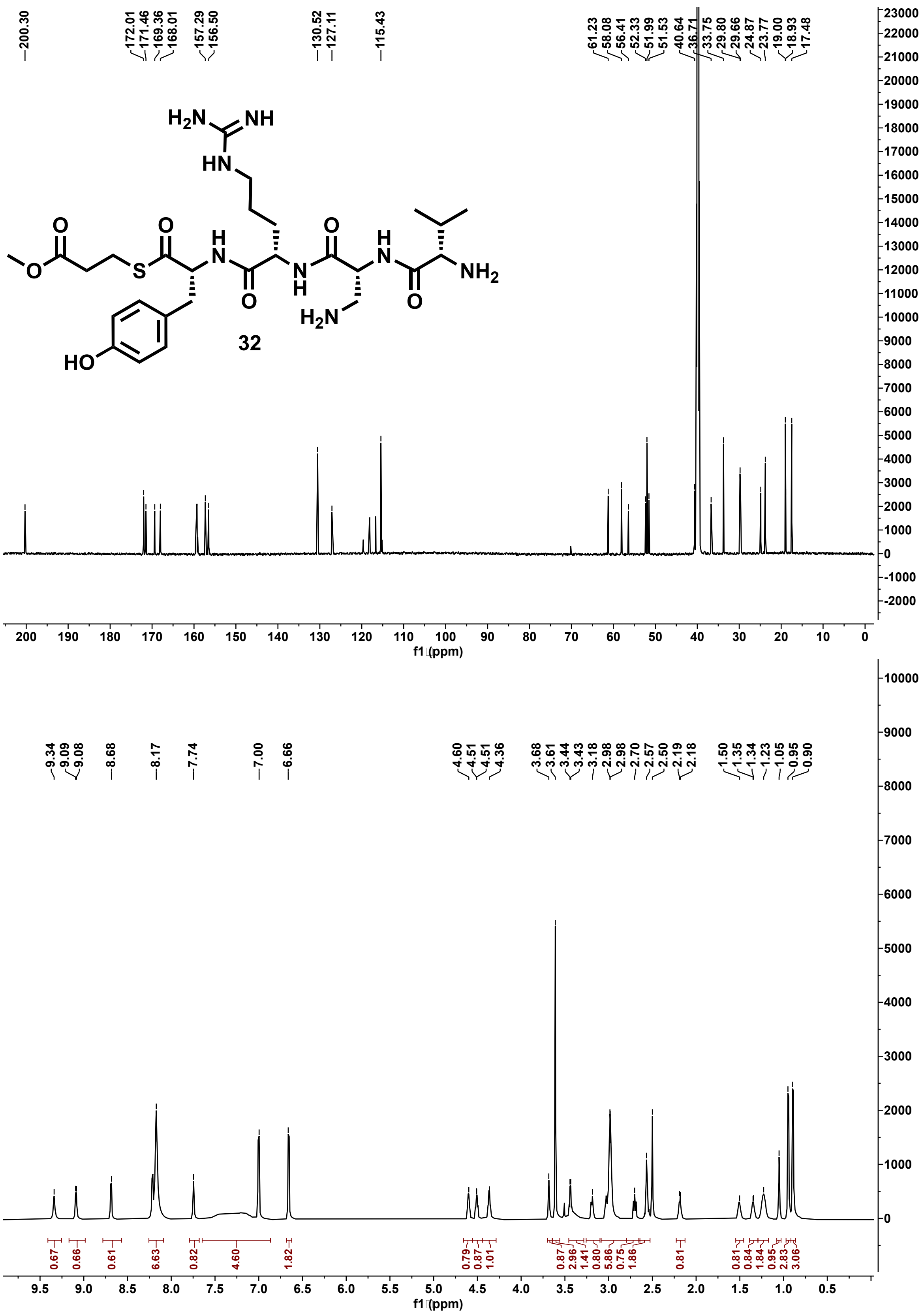

**Figure S140: $^{13}$C (201 MHz) and $^{1}$H (800 MHz) NMR spectra of 32 in DMSO-d$_6$.**

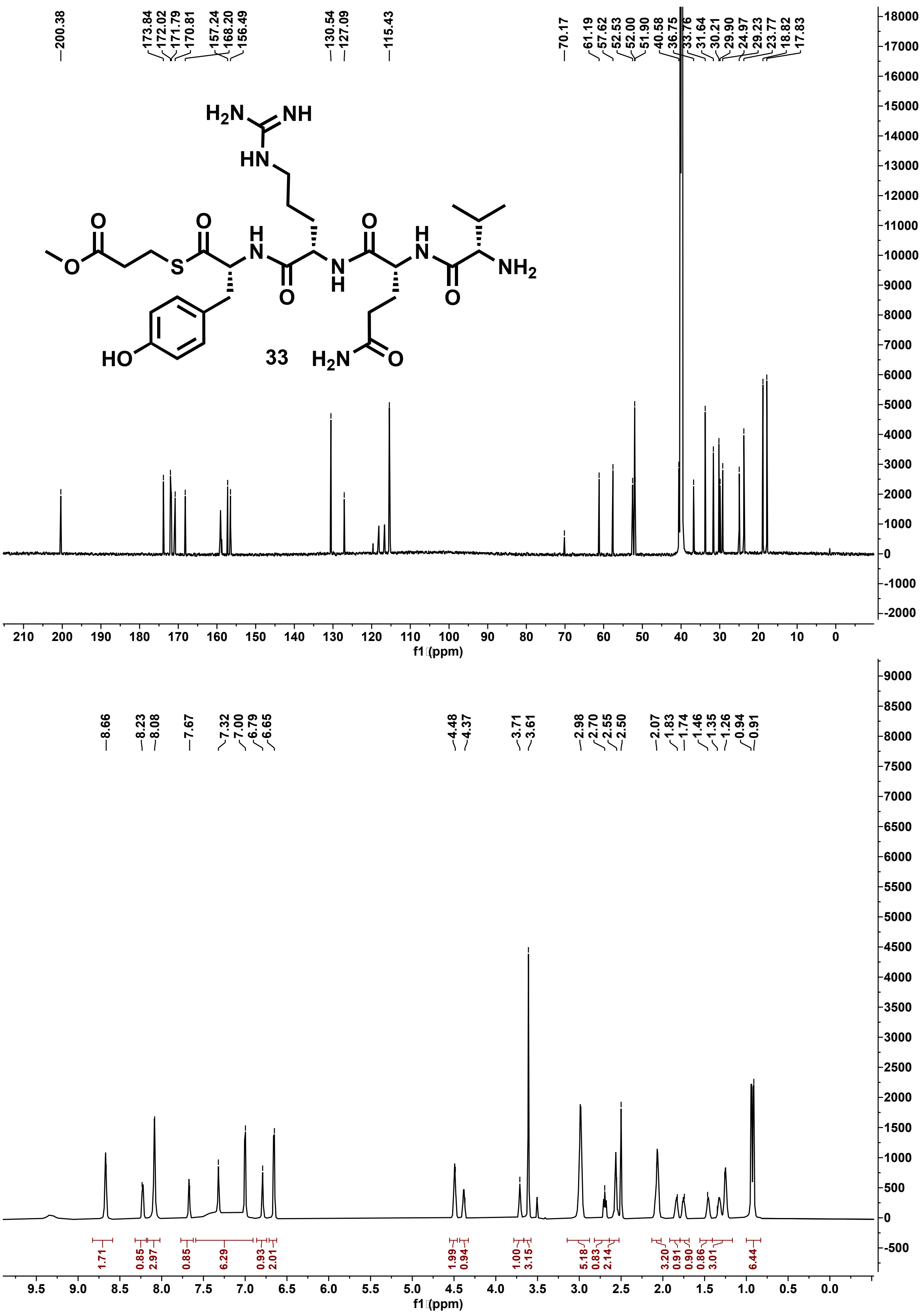

**Figure S141: $^{13}C$ (201 MHz) and $^{1}H$ (800 MHz) NMR spectra of 33 in DMSO-$d_6$.**

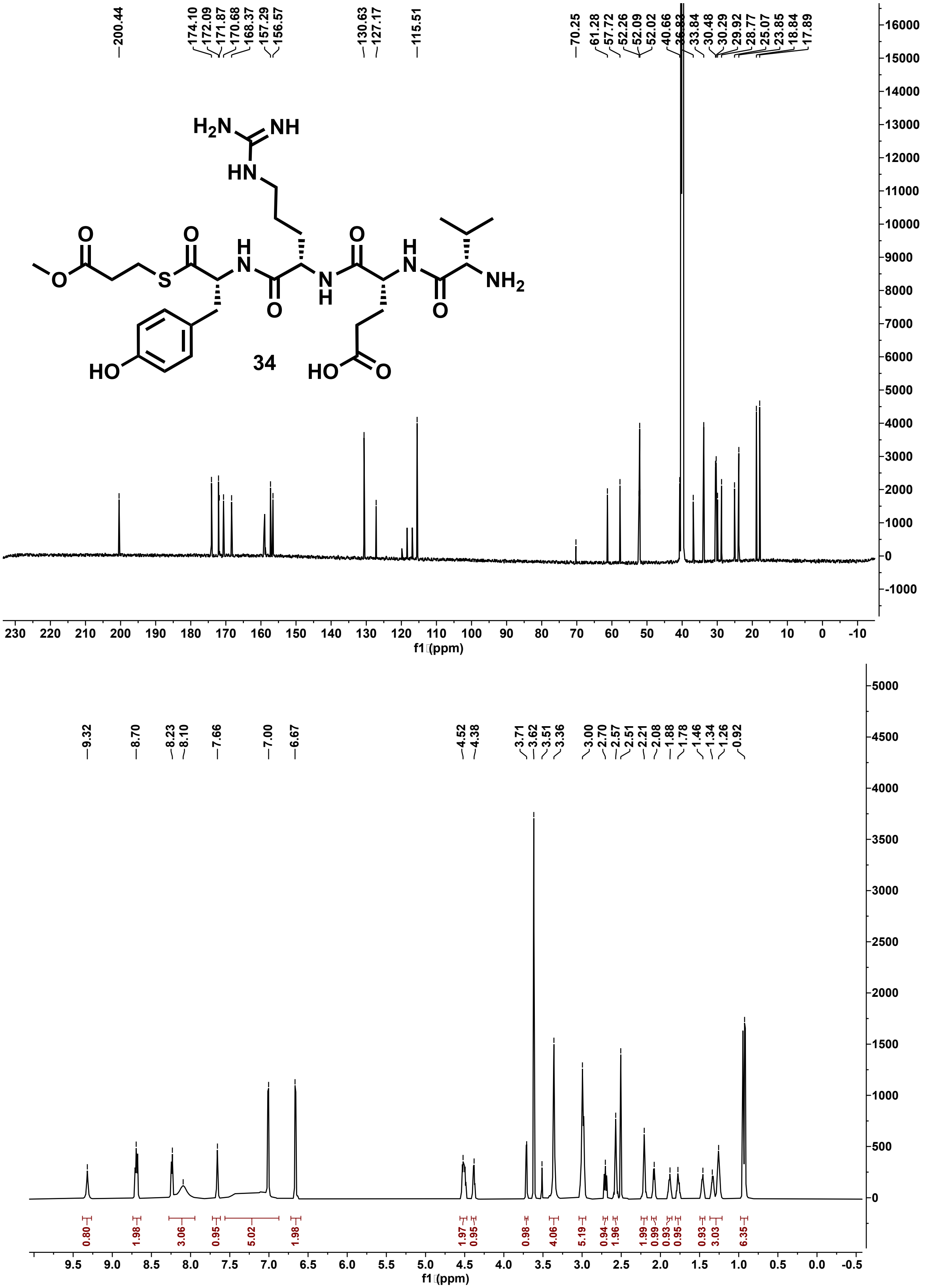

**Figure S142: $^{13}C$ (201 MHz) and $^{1}H$ (800 MHz) NMR spectra of 34 in DMSO-$d_6$.**

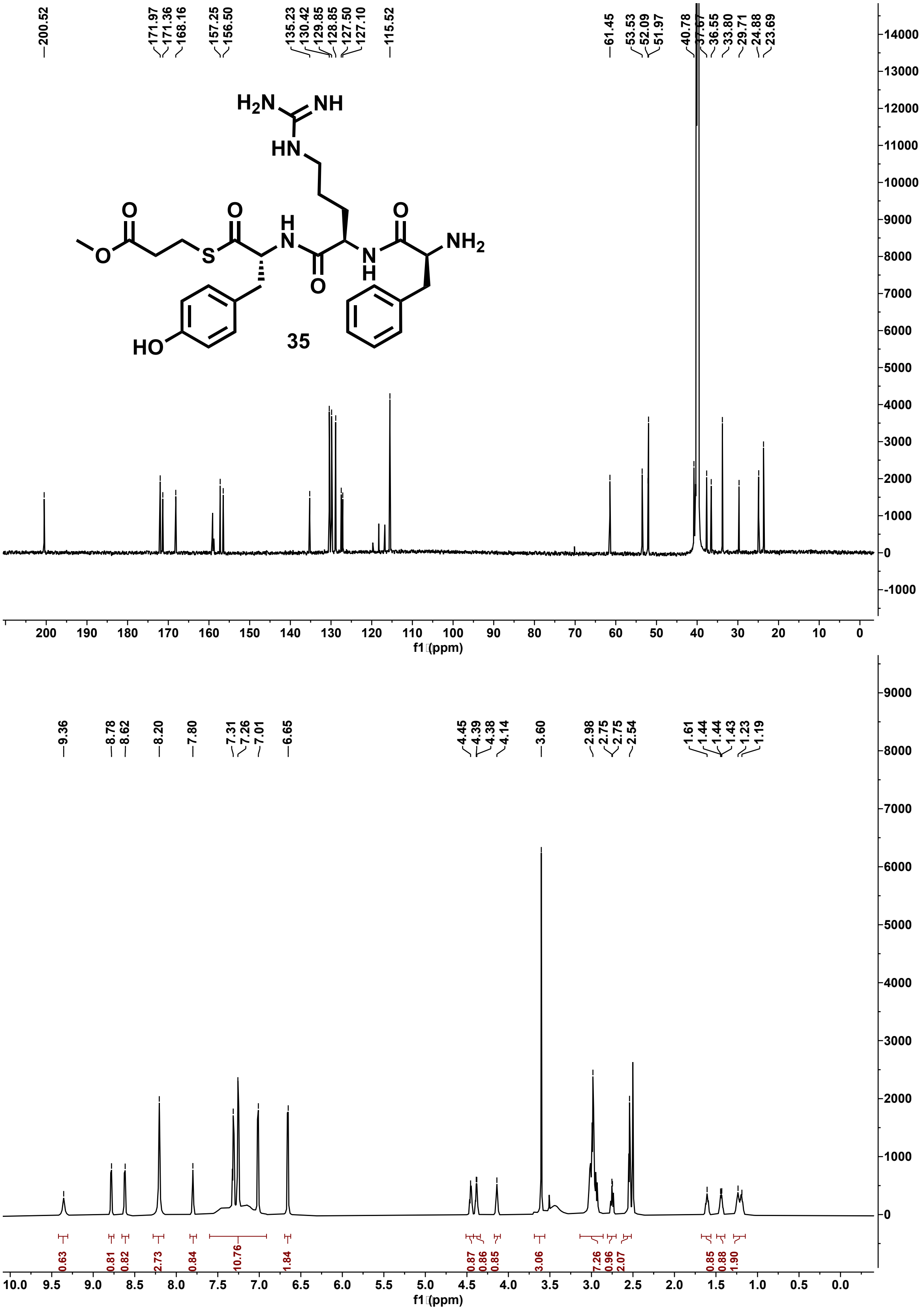

**Figure S143:** $^{13}C$ **(201 MHz) and** $^{1}H$ **(800 MHz) NMR spectra of 35 in DMSO-d**$_6$**.**

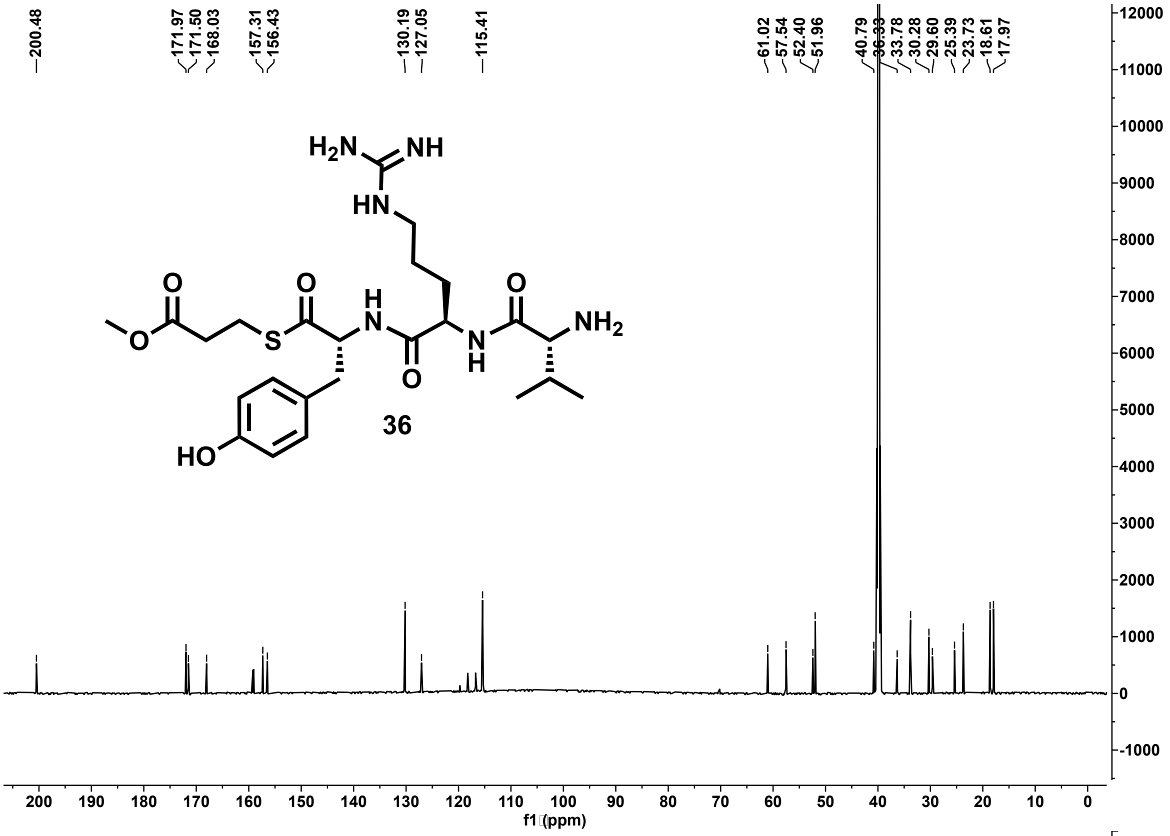

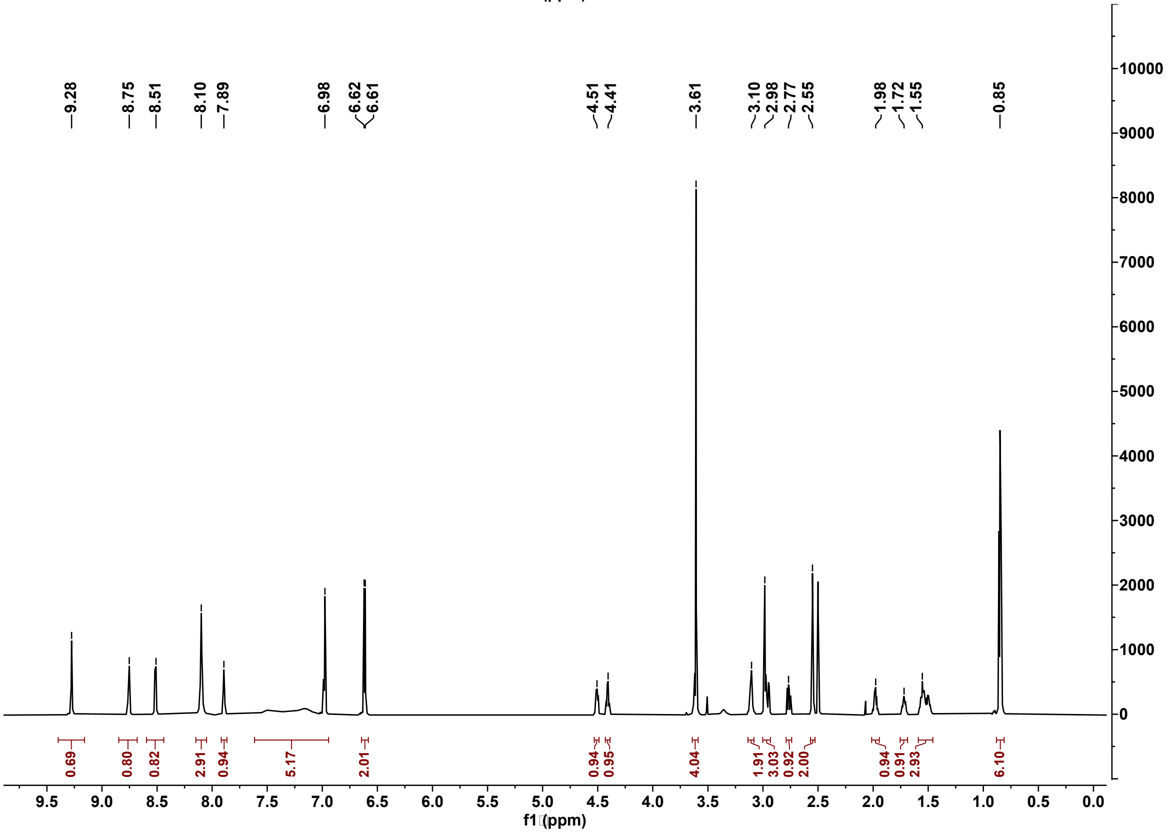

Figure S144: $^{13}$C (201 MHz) and $^{1}$H (800 MHz) NMR spectra of 36 in DMSO-d$_6$.

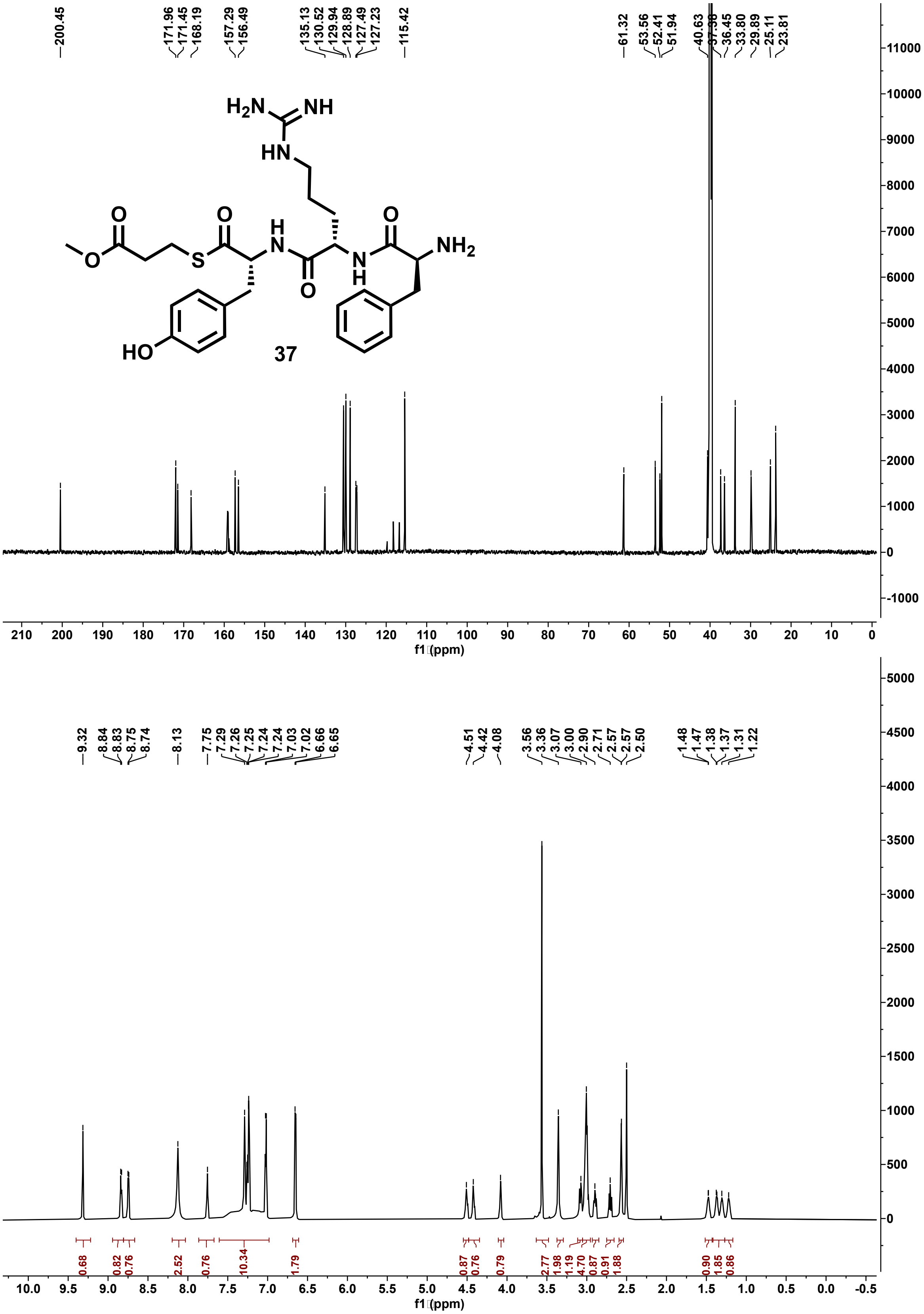

**Figure S145: $^{13}C$ (201 MHz) and $^{1}H$ (800 MHz) NMR spectra of 37 in DMSO-$d_6$.**

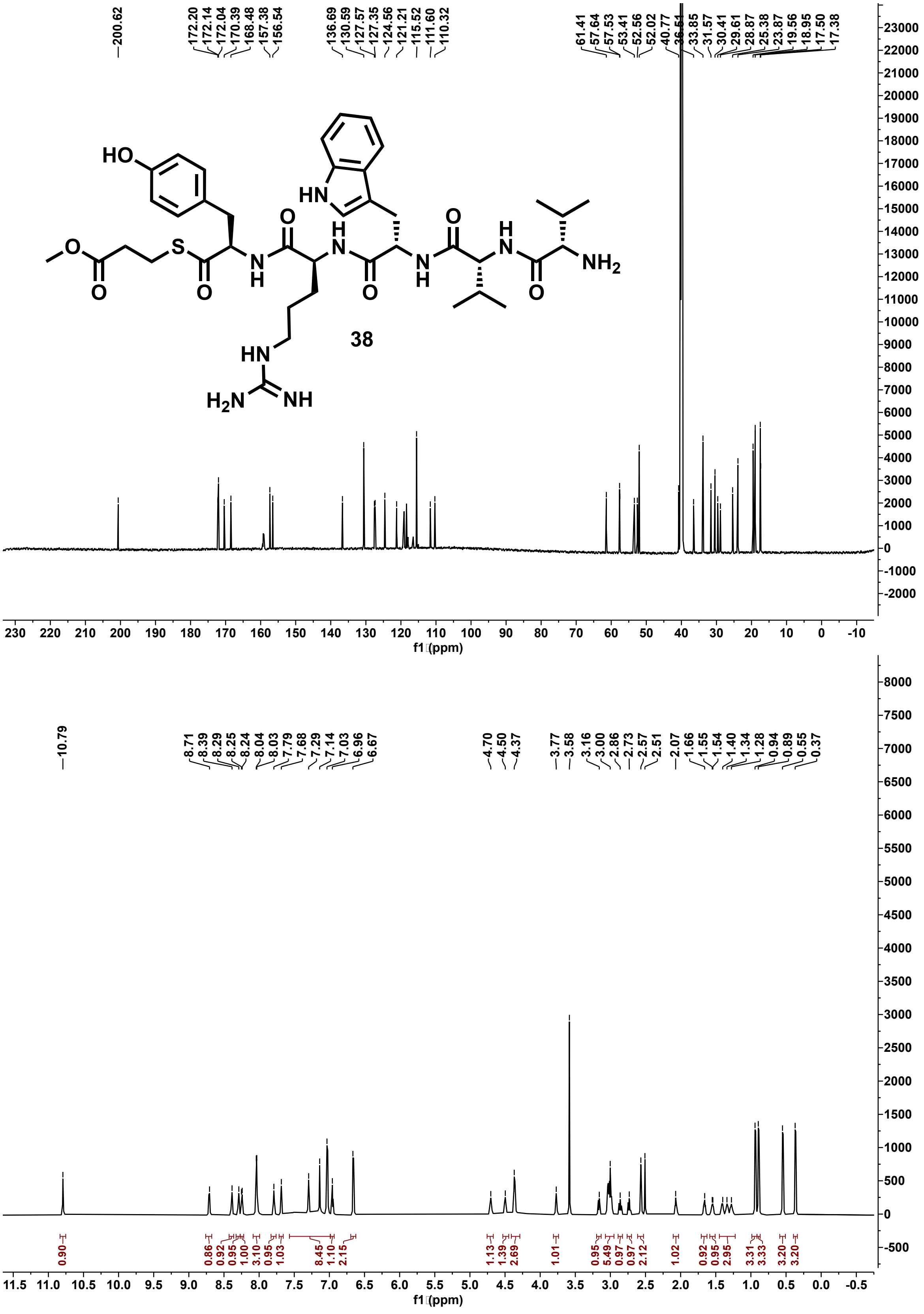

**Figure S146: $^{13}C$ (201 MHz) and $^{1}H$ (800 MHz) NMR spectra of 38 in DMSO-$d_6$.**

**Supplementary note 1: Gene sequence for WP516**

ATGGACGATGTGATCGCACGGCTGGCGCCCCTGCTGCGGCGCCACCGTGTCCCGGGAGCGCA
GCTCGCCCTGCGGTGGCAGGGGCGTACGTACACCGCCGAGGCGGGTGAGGAGAGCGCCGGC
GCCGGCCGGCCGGTGACCGGCGGGACGGCGTTCCCGCTCGGCTCGCTCACCAAGCCGTTCAC
CGCCACGCTGGCGATGATGCTGGCCGCCGACGGCGACCTGGAGCTCGACGAGCCGGTGTCGG
CCTGCCTCCCCGGGCTCCGGCCCGGGGCGGGGCGGGTGACCCTGCGGCAGCTCCTCAGCCAC
ACGGCCGGGCTGCCTGCCAACGTGGAGGAGTCGGCGGCGGGCCTGACCCGCCGCCGGTGGG
CGGAGGAGTCCGCCGCGGCGGTGCACCCGCCCGGCTCCGCCTTCTCCTACTCCAACGCCGGA
TACCTACTGGTGGCGCACCTCGTCGAGGACGTCACCGGGATGAGCTGGCGGGCGGCCGTGGA
GTCGTTCCTGCTGCGTCCGCTGGGCATCGAGCCGCTGTTCGCGGTCGGCGCCCGGCGGGCGG
CGGAGCAGGCCGGCGCCCGGCCGGCCGCCGACGGCCACGTCGTGCGGCCGGACGGGGCGGC
GCTGCCGATCGCCGAGCAGTCGGTGACGGCGCTGGAGGAGCCGGTGGCGGGGCTGGCGGGC
AGCGCGGCCGACCTGCTCGCCTTCGCCGCGCTGCACCTGCCCGGCCACGAGGGGCCCGGCCT
GCTGGACGGGGAGAGCGCCGCGGAGATGCGCCGCGACCAGCTCGGCGGCCTGCCCGCGGGC
GCGTTCGGCCTGGCGGACGGCTGGGGCCTGGGCTGGTCCCTGTACGGCACGGCGTCGGGCAC
CTGGTTCGGCCACGACGGCACCGGCGACGGCGCCTGGTGCCACCTGAGGGTGCAGCCGGAG
ACCGGGACGGCGGTCGCGCTGACCACCAACGGCGGCAACGGGGCCGTCCTGTGGGAGGCCG
TGGTCGCGGAGCTGCGCGCGGCGGGCGTGGACGTCGGCCACTACCCCCTGGGCGCGCTGACC
GGCGGGGGAACGCCCCTGCCGGCCGGCTCGCCCGCGAAGGACGGATGCGCGGGCCGCTACG
CCAACGGCTCCTGGACCTGCGCCGTCGAGGCGTCGGGCGGCGAGTTGTACCTGTCCGTCGGG
CCGGGACCGCGCTGGCGGCTGCGCTGCTTCGAGGACCTGCGCTTCACCACGGAGGGCTCGG
ACGCCGGCTCGATGCCCTACGTCGGCCGCTTCCTGCGCGACCCCGGCTCGGGGACGGTCGAG
CTGGTGCAGATCACCGGGCGGCTGGCCCGGCGGCATGGCTAG

**Supplementary note 2: Gene sequence for SEC28301.1**

ATGAATGATCTGCTGGCAGAACTGGCACTGCTGCATGGTGTTCCGGGTGCACAGGTTGCAGTT
TGGCGTGATGGTCTGCTGCGTACCGCAGAAACCGGTGAAGAAGAAGCAGGTAGCGGTCGTCC
GGTTACCGTTGAAACCGCATTTCCGCTGGGTAGCCTGACCAAACCGTTTACCGCAACACTGGC
AGCACTGCTGGTTGCAGATGGTGATGTTGATCTGGATGAACCGCTGGCCGAACAGCTGGGTG
AACTGCGTGCAGGTCCGGAATTTACCCTGCGTCAGGTTCTGAGCCATACCGCAGGTCTGGCA
GCAAATACAGCAGAACCGCCTGTTGGTACAACCCGTGCACGTTGGCTGGCACGTCATGCAGC
CGAACCGGTTGTGCATGAACCGGGTACAGTTTTTAGCTATAGCAATCCGGGTTATGTTATTGCA
GGTCGTCTGGTTGAAGAAATTACCGGTCTGGATTGGGCTGAAGCAGTTCGTACCATGCTGCTG
CGTCCTCTGGGTCATGATGCAGCAGCACCGACCGCACTGGGTCATCTGGTTCGTCCGGATGCA
CCGCCTCGTCCGATTCGTGAACAGAGCGTTCCGCCTCTGGAAGATCCGGCAGCAGGTCTGCG
TGTTAGCGCACGTGATCTGGCAGTTTTTGGTGCAGCACATTTAGGTTTAGGTCCTGGTGGTGG
CCTGCTGGATGCAGCAACCGCACGTGCAATGCGTGAAGATGTTACCGAAGGCCTGGCAGTTG
GTGCACATGGTCTGGCCGATGGTTGGGGTGCAGGTTGGAGCCGTTATGGTGCATGGTTTGGCC
ATGATGGCACCGGTGATGGTGCCTGGTCACATCTGCGTGTGGATCCGAGCACACGTACCGTTG
TTGCACTGACCGCAAATGGTAGCAGCGGTGCACGTCTGTGGGAAAGCCTGCTGGGTCGTCTG
CGTGGTACAGGTCTGGAAGTTGGTGATCATCCGCGTGATACCGGTGGTGCAGAAAGCGTTCC
GGCACCGCAAGAATGTGCAGGTCATTATGCAAATGGTGATTGGAGCTGTCGTGTTGAAGCAG
AAGATGGCGATCTGCTGCTGAGCGTTGCCGGTGCAGCTCCGGTTCGCCTGCTGGTAGGTGAA
GATCTGAGTTTTCGTACCCCGAGCGGTGGTGGTCCGCGTGCAGCAATGCCGTATCTGGGTCGT
TTTCTGCGTGATCCGGCAACCGGTGCACTGGATCGTGTTCAGATTACTGGTCGTCTGTGTGTTC
GTCGTAAATAA

**Supplementary note 3: Gene sequence for WP516 Chimera**

ATGGATGACGTAATAGCTCGTTTGGCACCTCTCCTGCGCCGCCATCGCGTGCCAGGAGCGCAG
TTGGCTCTCCGATGGCAAGGGCGGACGTATACAGCCGAGGCTGGCGAAGAATCCGCTGGTGC
TGGGAGACCGGTCACTGGTGGGACAGCGTTCCCGCTTGGGTCACTCACGAAGCCGTTCACGG
CCACGCTCGCGATGATGTTAGCTGCGGATGGTGACCTTGAGTTAGACGAGCCTGTCTCCGCCT
GTTTACCCGGATTACGCCCCGGGGCGGGTCGCGTCACATTACGTCAATTGTTAAGTCACACAG
CCGGGTTGCCGGCCAATGTAGAGGAGTCCGCCGCCGGACTGACACGACGACGTTGGGCCGA
GGAGAGCGCAGCAGCGGTACATCCACCAGGGTCGGCCTTTAGCTACTCTAACGCTGGGTATTT
ATTAGTGGCACACTTAGTAGAAGATGTGACAGGTATGTCGTGGCGTGCCGCCGTGGAGTCCTT
CCTTCTTCGTCCGCTGGGTATCGAGCCTCTCTTTGCCGTGGGTGCCAGACGCGCGGCAGAGCA
AGCAGGAGCTCGTCCCGCCGCAGACGGCCATGTAGTACGTCCCGACGGCGCAGCACTTCCAA
TAGCCGAGCAGTCGGTTACCGCCTTGGAAGAGCCCGTGGCGGGATTAGCCGGTTCAGCCGCC
GACCTGCTGGCCTTCGCCGCGTTGCATCTTCCAGGACATGAGGGCCCGGGGCTTTTAGACGGC
GAGAGCGCAGCTGAGATGAGACGTGACCAACTCGGTGGACTGCCTGCAGGGGCTTTCGGAC
TCGCTGACGGGTGGGGCCTGGGATGGAGTCTTTACGGTACGGCTAGCGGTACCTGGTTTGGCC
ACGACGGCACCGGAGACGGCGCATGGTGTCATTTACGCGTCCAGCCAGAAACCGGCACCGCA
GTTGCCCTGACTACAAACGGCGGAAATGGAGCTGTCCTCTGGGAAGCGGTTGTGGCGGAGCT
TAGAGCTGCTGGTGTGGACGTCGGTGTCCACCGTCCAGCTCCGCCGCCGGCGATTGCAGCTG
CAGCGTTTGCCGATTGCACCGGTACATACCGTAACGGGGACCTGGCCGTTACTGTGGGAATTG
ACGGCCCGTATCTGGTTCTGGAGCTGCCGGGTGGAGCGCGTGAACTCGCACAACCTTTAGCC
CACAGAACGTTCTCGAGCCGGGGTGCAGGTTTCTTAGGGCGCTTCGTTACGGACGCTCGCTC
AGACGCTGTACACGCGTTACAGTACTCCGGGCGAACCCTGTTACGTGAGGCGGGTTGA

**Supplementary note 4: Gene sequence for Ulm16 Chimera**

GGGCTTGATTTGGATCGCCTCGCGCGGGATTGTGATGTAGTTGGTGGTCAGTTGGCTTTACAC
CATCAAGGCACTCTGACCACCTGGGAGTTCGGGACTGAGGAACACGCCGGCGGTAGACCAG
TCCACGTCGGGTCAGCTTTTCCATATGGCAGCGTGACAAAGGCATTCACCGCAACCGCTGTTT
TACAGTTGGCCGGTGACGGAGACTTGGACCTCGACCGCCCCGTCCGTGAGCTGCTGCCAGAG
GCAGAAGCCGAAGCTGAAATTGAAGTCGAAGCAGGATCTGGGACCGGCGCACGCGCGGACG
GTGGACACCCTGCGCTTGCGGCGACACTGCGGCAATTGTTATCCCATACGGCTGGCTTACCTA
GCGACCATGATGACGAACGCGCTCCGAGCTTACGACGTTGGCTCACCGGGTTCTTAGCCCTGC
CGGTGGGCCCCTGGCCAGCACCAGGCTCTTTCAGTTACAGTAACGTGGGGTACGGAATCGCA
GGCCGCGTCGTAGAAGCCGTAACGGGCTTAACTTGGTCAGAAGCAGTCCGCGACTTTCTGTT
ACATCCACTGGGCATCGAACCGTTATTTGCTGTCGGAGCTCGACGCGCTGCCGAGCAGGCTG
GAGCACGGCCCGCCGCTGACGGCCACGTGGTGCGTCCTGATGGAGCAGCATTGCCAATAGCC
GAGCAAAGTGTCACTGCGCTCGAGGAACCAGTCGCAGGTCTGGCGGGCAGTGCGGCTGACC
TGGTTCGCTTAGGGCGGTTGCACTTGGACGAACCGGGCGACCCTGACTTGGCTCGCCTGGCG
GATCCTGATGCCTTACGTGAGATGGCTCGTCCGACGGCGGGAGCTGATCCGTTTGGTTTAGCG
GACGGTTGGGGTCCAGGCTTAGGTCGCTTTGGTCCGGCCGGAAACAGATGGCTGGGACACGA
CGGCACCGGTGATGGCGCTACGTGTCATTTACGTATCCATCCTGGACGCGGCACGGTCGTAGC
CCTGACAACGAATTCTCCAACAGGGCAAGCCTTATGGGACGCCGTTGTTGACGCTTTACGTGA
CGCCGACATCGACGTTGGACACTACCCCCTGGGTGCCTTGACCGGGGGGGGAACCCCACTGC
CAGCTGGATCACCCGCTAAAGACGGTTGCGCTGGCAGATATGCGAATGGGTCGTGGACATGC
GCTGTCGAGGCTTCTGGTGGTGAACTGTATTTATCGGTCGGTCCGGGTCCAAGATGGAGACTG
CGTTGTTTCGAAGACTTACGTTTCACGACCGAGGGCTCAGATGCAGGATCGATGCCGTACGTT
GGGAGATTCCTCAGAGACCCCGGTTCCGGTACAGTAGAGTTAGTTCAAATCACAGGCCGCCT
GGCACGCCGCCATGGCTAGAAGCTTGCGGCCGCACTCGAGCACCACCACCACCACCACTGA

**Supplementary note 5: Gene sequence for Ulm16$^{T304W}$**

GGAGCGGGAGACGGAGCACCTGTGGGATTAGACTTAGATCGCCTTGCCCGTGATTGTGACGT
CGTAGGTGGTCAGCTTGCATTGCATCACCAAGGCACTCTGACCACCTGGGAATTCGGTACTGA
GGAACACGCTGGTGGTCGGCCAGTCCATGTGGGTTCTGCCTTTCCGTATGGTTCTGTGACCAA
AGCCTTTACAGCAACTGCTGTCCTTCAATTGGCAGGGGACGGCGACCTTGACCTTGATCGCCC
GGTTAGAGAGCTGCTGCCGGAAGCAGAAGCTGAGGCGGAAATAGAGGTGGAAGCAGGCTCG
GGAACAGGTGCGCGGGCGGACGGCGGGCATCCCGCGTTAGCGGCAACATTGCGGCAGCTCCT
GTCACACACAGCGGGCCTTCCTTCCGACCACGACGATGAACGCGCTCCGTCTCTGCGCCGTT
GGTTAACCGGTTTCTTAGCTCTTCCGGTAGGCCCGTGGCCGGCTCCTGGCTCGTTTAGTTATTC
CAATGTAGGCTATGGCATAGCCGGTCGCGTTGTTGAGGCAGTAACGGGACTGACTTGGTCGGA
AGCGGTTCGCGACTTTCTGCTGCACCCGCTCGGCACCGCGATTACTGTGCTTCCGACTGATCC
AGGTAGTCTTCCGGCAGGCGGACTGGCCGGATCAGCAGCTGATCTGGTCAGATTGGGCCGCC
TGCACCTTGACGAACCAGGCGATCCAGACCTGGCACGACTTGCAGATCCGGATGCTTTACGG
GAAATGGCACGGCCCACCGCCGGTGCCGATCCATTCGGGTTGGCGGATGGATGGGGCCCGGG
ACTCGGACGTTTCGGTCCCGCCGGAAATCGGTGGCTGGGGCATGACGGGACTTTAGATGGCG
CTTGGTGCCATCTGCGTATTCATCCCGGGCGCGGAACAGTGGTAGCACTGACCACTAACTCAC
CTACAGGCCAGGCTCTGTGGGATGCCGTGGTAGACGCGCTTCGGGACGCGGATATAGATGTCG
GTGTACACCGGCCTGCACCTCCTCCGGCGATTGCGGCAGCGGCTTTTGCGGACTGCACCGGA
ACCTACCGCAATGGTGATCTTGCCGTGACTGTTGGGATAGATGGGCCATATCTGGTTTTGGAAC
TGCCGGGCGGGGCCCGAGAGCTTGCACAGCCTCTGGCTCACCGCACTTTTAGTTCTCGGGGC
GCGGGCTTTCTGGGGCGGTTTGTGACAGATGCACGCTCTGACGCTGTGCACGCACTGCAGTAT
TCAGGCCGGACGTTGTTACGAGAAGCAGGAGAGTCTTAG
